# FastqPuri: high-performance preprocessing of RNA-seq data

**DOI:** 10.1101/480707

**Authors:** Paula Pérez-Rubio, Claudio Lottaz, Julia C Engelmann

## Abstract

**Background:** RNA sequencing (RNA-seq) has become the standard means of analyzing gene and transcript expression in high-throughput. While previously sequence alignment was a time demanding step, fast alignment methods and even more so transcript counting methods which avoid mapping and quantify gene and transcript expression by evaluating whether a read is compatible with a transcript, have led to significant speed-ups in data analysis. Now, the most time demanding step in the analysis of RNA-seq data is preprocessing the raw sequence data, such as running quality control and adapter, contamination and quality filtering before transcript or gene quantification. To do so, many researchers chain different tools, but a comprehensive, flexible and fast software that covers all preprocessing steps is currently missing.

**Results:** We here present **FastqPuri**, a light-weight and highly efficient preprocessing tool for fastq data. **FastqPuri** provides sequence quality reports on the sample and dataset level with new plots which facilitate decision making for subsequent quality filtering. Moreover, **FastqPuri** efficiently removes adapter sequences and sequences from biological contamination from the data. It accepts both single- and paired-end data in uncompressed or compressed fastq files. **FastqPuri** can be run stand-alone and is suitable to be run within pipelines. We benchmarked **FastqPuri** against existing tools and found that **FastqPuri** is superior in terms of speed, memory usage, versatility and comprehensiveness. **Conclusions: FastqPuri** is a new tool which covers all aspects of short read sequence data preprocessing. It was designed for RNA-seq data to meet the needs for fast preprocessing of fastq data to allow transcript and gene counting, but it is suitable to process any short read sequencing data of which high sequence quality is needed, such as for genome assembly or SNV (single nucleotide variant) detection. **FastqPuri** is most flexible in filtering undesired biological sequences by offering two approaches to optimize speed and memory usage dependent on the total size of the potential contaminating sequences. **FastqPuri** is available at https://github.com/jengelmann/FastqPuri. It is implemented in C and R and licensed under GPL v3.

## Background

Quality control (QC) and filtering of sequence data are important preprocessing steps to generate accurate results from RNA-seq experiments. The work-flow usually proceeds as follows: initial check of sequence quality based on diagnostic quality plots followed by sequence filtering to remove adapters and low quality bases. Then, contaminations from other organisms are removed, and finally, another quality control run is performed to confirm that the sequence data is now acceptable.

Although tools exist that perform sequence data quality control, and others that do filtering or trimming, there is no adequate and comprehensive tool that would cover all preprocessing steps commonly used on RNA-seq data. Considering QC, FastQC [1] is widely used for RNA-seq data, but because it was designed for genomic data, several of its quality checking modules are not suitable for RNA-seq data (e.g., overrepresented sequences, sequence duplication level, GC content). While RSeQC [16] and RNA-SeQC [8] were written for RNA-seq data, they only take alignment files (BAM) as input, which renders them inappropriate when working with alignment-free transcript counters such as kallisto [4] and salmon [14]. AfterQC [5] performs quality control and global quality filtering, but does not specifically address RNA-seq data. Its strand bias detection and overlapping pair analysis is not useful for RNA-seq data, and contamination filtering is not included. AfterQC is also limited in its automatic filtering capabilities based on quality scores. It can only globally trim, that is remove a fixed number of bases from each read. While RNA-QC-Chain [18] claims to provide comprehensive quality control for RNA-seq data, it lacks informative graphics of the raw read (fastq) data and can only filter rRNA contaminations.

Moreover, while sequence alignment used to be the most time-demanding step in RNA-seq data analysis, this has changed since alignment free transcript counters were introduced. Now, quality control and filtering are the time-consuming bottlenecks. **FastqPuri** provides an automated and most efficient implementation for these first steps needed in all RNA-seq work-flows. It includes general quality control as well as filtering of low quality bases, calls marked as N, adapter remnants and reads originating from contaminating organisms. Our software handles both uncompressed and compressed fastq files from single- or paired end sequencing, and provides superior diagnostic plots in a per sample quality report and a summary report over all samples in the dataset.

## Implementation

**FastqPuri** consists of six executables which can be run sequentially to assess sequence quality and perform sequence filtering. `Qreport` assesses sequence quality at the sample level, while `Sreport` generates a summary quality report for a collection of samples, e.g. the complete dataset. For contamination filtering, **FastqPuri** offers two different methods, a tree-based and a bloom filter-based method. The executables `trimFilter` and `trimFilterPE` filter contaminations, adapters and low quality bases from single-end and paired-end data, respectively. The work-flow of fastq sequence data preprocessing with **FastqPuri** is depicted in Figure 1.

**Figure 1:**
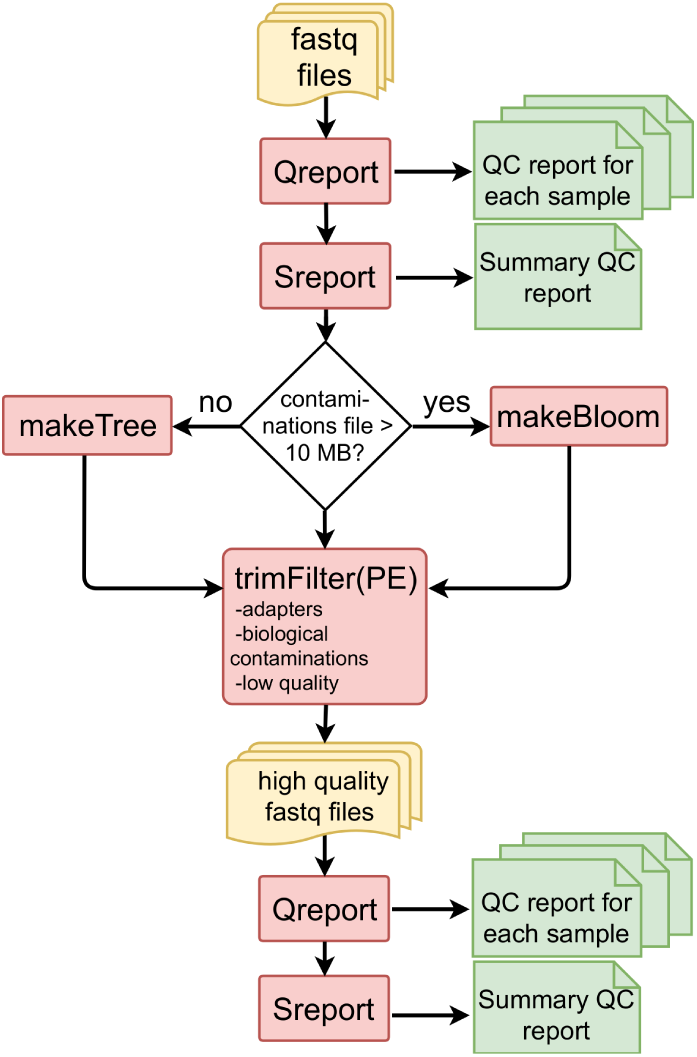
Workflow for preprocessing fastq files with FastqPuri. `Qreport` generates a quality report in html format for each sample, while `Sreport` generates one summary quality report for all samples. Depending on the size of the sequence file with potential contaminations, `makeTree` or `makeBloom` generates a data structure for filtering contaminations. `trimFilter` (or trimFilterPE for paired-end data) filters and trims reads containing adapters or adapter remnants, biological contaminations and low quality bases. On the filtered reads, `Qreport` and `Sreport` can be run again to ensure that the filtered data meets the user’s expectations. Legend: yellow: fastq files, red: **FastqPuri** executables, green: **FastqPuri** quality reports in html format.

### Assessing sequence quality

Assessing sequence quality thoroughly is essential to be able to detect problems during sample handling, RNA extraction, library preparation and sequencing. None of the existing tools fulfilled our requirements to comprehensively assess sequence quality and estimate the impact on data loss by applying different quality filters. Therefore, we designed novel graphics which allow to estimate how many sequences will be discarded at a specific quality threshold, for a range of thresholds. With existing tools, this would require several runs of filtering with different thresholds and calculating the number of kept reads, while we get this information with just one run of `Qreport`. The resulting html report contains general information about the dataset (Figure 2A), the common plots of average sequence quality per base position (Figure 2B), average quality per position per tile per lane (Figure 2C) and nucleotide content per position (Figure 2D). In addition, **FastqPuri** quality reports include plots to facilitate decision making about thresholds to be used for quality filtering, especially for the purpose of using transcript counting approaches for transcript and gene expression analyses. Therefore, Figure 2E displays the proportion of nucleotides per position per tile which fall below the high quality threshold required. This plot better highlights problematic tiles and nucleotide positions than the one showing average quality values per position and tile (Figure 2C), which is shown e.g. in FastQC reports. For example, from Figure 2C, we cannot see if the bases of all the reads have lower qualities at positions 1-5, or if there is only a subset with very low qualities that would decrease the mean. From Figure 2E it becomes clear that most of the reads (> 95%) have quality scores above the required quality threshold across all tiles. Figure 2F shows the proportion of low quality nucleotides per base A, C, G, T and per tile. Figure 2G shows the proportion of reads meeting a certain quality threshold, allowing a quick assessment of the data that would be discarded at a given threshold. This information is lost in plots showing averages, as for example Figure 2B. Moreover, for transcript counting methods such as kallisto and salmon, it is important to get an overview over how many reads contain many low quality bases. They should be filtered out to avoid false-positive mappings, because these methods do not take quality scores into account. If many reads carry only one low quality base, this could be tolerated. Therefore, we show the number of reads with *m* low quality bases in a histogram to allow the user to make an estimate about how many sequences will be discarded when requiring a certain percentage of high quality nucleotides per read (Figure 2H). Quality reports for each sample are generated by the executable `Qreport`, while the executable `Sreport` provides a summary quality report over all samples in the dataset. There are two types of summary quality reports: the first one is a quality summary report and consists of an html report with a table of the number of reads, number of tiles, percentage of reads with low quality bases, percentage of reads with bases tagged as N for all samples, and a heatmap showing the average quality per position for all samples. The second type of summary report provides an overview over the filtering which was performed with `trimFilter(PE)` (see following section). It contains a table specifying the filter options used, and a table containing, for all samples (rows), the total number of reads, the number of accepted reads, the percentage of reads discarded due to adapter contaminations, undesired genome contaminations, low quality issues, presence of Ns, and the percentage of reads trimmed due to adapter contaminations, low quality issues and presence of N’s.

**Figure 2:**
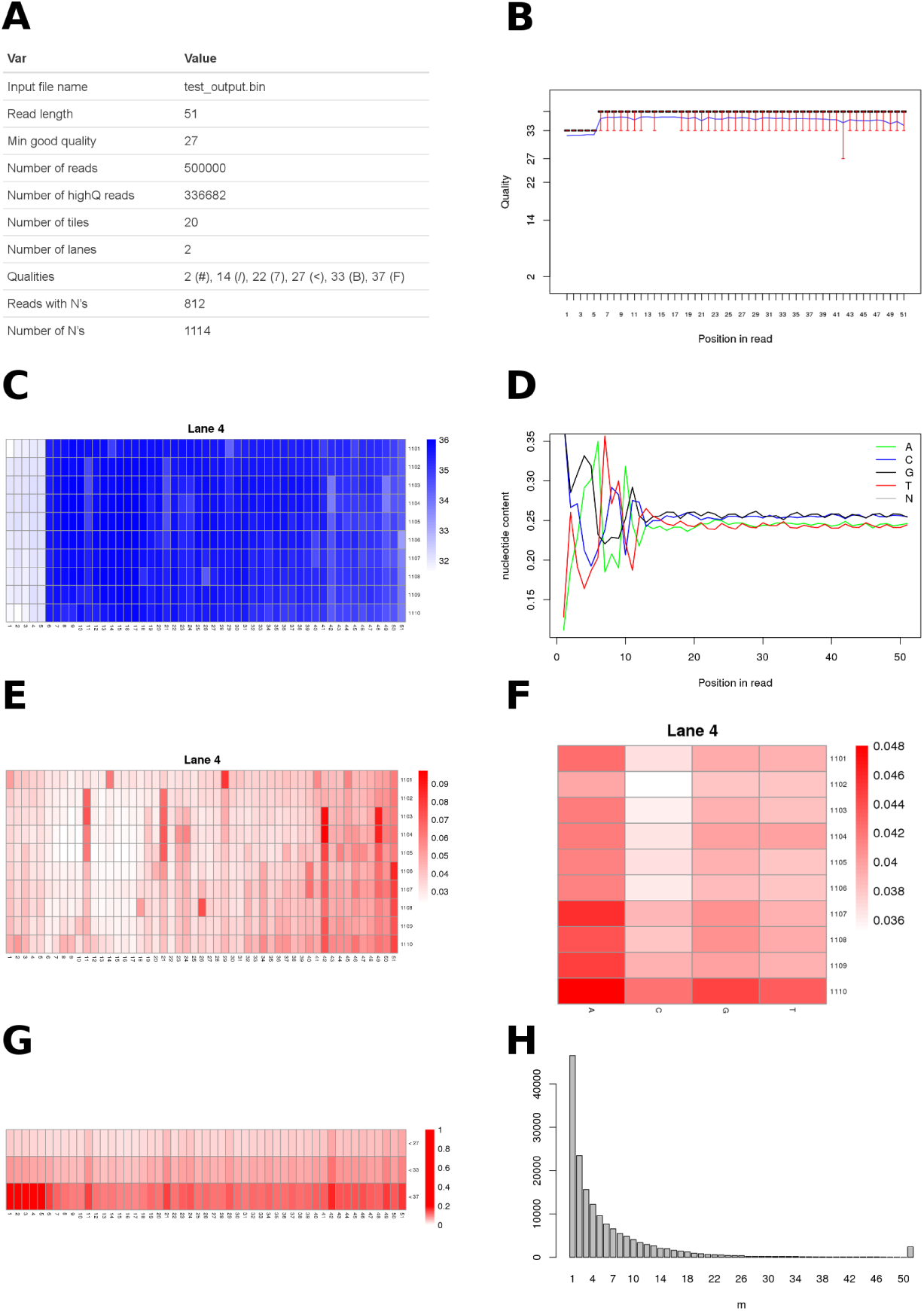
Graphics shown in Qreport. A) Data set overview and basic statistics. B) Per base sequence quality box plots. The blue line corresponds to the mean quality value. C) Cycle average quality, per tile, per lane. D) Nucleotide content per position. E) Proportion of low quality bases, per tile, per lane. F) Fraction of low quality bases {A, C, G, T} per position, per tile and per lane. G) Proportion of bases with quality scores below different thresholds, for all tiles, all lanes. H) Number of reads with *m* low quality bases.

### Filtering contaminations

We first filter out technical (e.g. adapters, primers) and biological undesired sequences and then bases and reads with low quality scores. We purposely do it in this order to make sure we do not overlook contaminating sequences that were trimmed due to quality issues. The actual filtering is performed by `trimFilter` for single-end reads and `trimFilterPE` for paired-end reads. Optionally, the executables `makeTree` and `makeBloom` are used to prepare the filtering (Figure 1), they are described below.

#### Contamination with adapter sequences

**FastqPuri** can remove adapters, adapter remnants or any other kind of technical sequence that is introduced during sequence library preparation from single and paired end data. We use an approach similar to trimmomatic [3], scanning reads from the 3’ to 5’ end with a 16 nt seed and performing local alignment if the seed is accepted. If the alignment score exceeds the threshold, the adapter is removed. If the remaining read is shorter than the minimum allowed sequence length, it is discarded. For paired-end data, both reads of a pair are discarded when one becomes too short after adapter-trimming.

#### Contamination with biological sequences

RNA-seq data can contain substantial numbers of reads which did not originate from mRNAs of interest. Even if an mRNA enrichment or rRNA depletion library preparation protocol was used, reads representing rRNA may be found [15, 17]. In addition, biological contaminations from spill-over, pathogen or host genomes, or bench contamination can result in sequence reads of different organisms than the one under study and in the worst case lead to distorted (false-positive) gene/transcript counts [2]. Therefore, it is good practice to check for potential sequence contaminations and remove them if needed. This functionality is provided by `trimFilter` for single-end reads and `trimFilterPE` for paired-end reads. We offer two options depending on the length of sequences to be removed exceeding 10 MB or not.

#### Short contaminating sequence

**4-ary tree.** If the fasta file of potential contaminations is smaller than 10 MB, we suggest to construct a 4-ary tree from the fasta file and use this to search for contaminations. The executable `makeTree` constructs a tree and saves it to disk for subsequent filtering with `trimFilter(PE)`. This is convenient for running the same contamination search on many samples. However, since constructing the tree is a relatively cheap computational task for the sequence lengths under consideration, per default the tree is not stored but generated each time `trimFilter(PE)` is called with `--method TREE. Searching` the tree is very fast but memory intensive. Therefore we limit the size of the potential contaminating species sequence file to be used with this filtering method.

#### Long contaminating sequences

**bloom filter. FastqPuri** offers a bloom filter approach to search for contaminations coming from large sequence files, e.g genomes from potential contaminating organisms with sizes up to 4 GB. For these applications, it is sensible to construct the bloom filter and store it in a file. This is done by `makeBloom`. A bloom filter is a probabilistic data structure which can be used to test if an element (here: a read) is an element of a set (here: the set of potential contaminating sequences). `trimFilter(PE)` with the option `--method BLOOM` then classifies each read as being contained in the bloom filter (representing contamination) or not. False positive hits are possible and by default, we accept 5% false positives. False negatives are not possible, except for cases where the contaminating sequences are different from the reference sequence due to individual variation, incomplete reference sequences or sequencing errors. Details about creating the bloom filter can be found in the supplement.

### Filtering based on base quality

We offer the following quality-based filtering options with `trimFilter(PE)`, which are specified with the `trimQ` argument:

- `NO`: (or flag absent): nothing is done to the reads with low quality,
- `ALL`: all reads containing at least one low quality nucleotide are discarded,
- `ENDS`: look for low quality base callings at the beginning and at the end of the read. Trim them at both ends until the quality is above the threshold. Keep the read if the length of the remaining part is at least the minimum allowed. Discard it otherwise,
- `FRAC [--percent p]`: discard the read if there are more than p% nucleotides with quality scores below the threshold,
- `ENDSFRAC [--percent p]`: first trim the ends as in the ENDS option. Accept the trimmed read if the number of low quality nucleotides does not exceed p%, discard it otherwise.
- `GLOBAL --global n1:n2`: cut all reads globally n1 nucleotides from the left and n2 from the right.

Independent of filtering based on quality scores, `trimFilter(PE)` can discard or trim reads containing ‘N’ nucleotides. This is done by passing the argument `--trimN` and one of the following options,

- `NO`: (or flag absent): nothing is done to the reads containing N’s,
- `ALL`: all reads containing at least one N are discarded,
- `ENDS`: N’s are trimmed if found at the ends, left “as is” otherwise. If the trimmed read length is smaller than the minimal allowed read length, it is discarded.
- `STRIP`: Obtain the largest N free subsequence of the read. Accept it if its length is at least the minimum allowed length, discard it otherwise.

## Results

### Comparison with other tools and evaluation

Several short read sequencing data tools address quality control and/or filtering. However, none of them integrates all preprocessing steps and meets our needs in terms of versatility, efficiency and visualization. Notably, none of the tools for quality analyses accepts bz2 files, the currently most common compression mode used by sequencing facilities to deliver Illumina fastq files. In Table 1, we compare the options of **FastqPuri** with several existing tools. With respect to the performance, efficiency and memory usage, we performed benchmarking on simulated and real data.

**Table 1:** Provided functionality of **FastqPuri** and existing tools. **lang**: programming language, **QC**: quality control, **QF**: low quality filtering, **Ad**: removes technical sequences such as adapters, **cont**: removes contaminations, **PE**: handles paired end data, **Year**: year of publication. fq* stands for un-compressed fastq or fastq compressed in gz, bz2, xz and for FastqPuri also Z format. For both FastqPuri and Biobloom, input may be tarred. ^1^functionality was added later.

| Tool name | lang | input | QC | QF | Ad | cont | PE | Year |
| --- | --- | --- | --- | --- | --- | --- | --- | --- |
| **FastqPuri** | C, R | fq* | ✓ | ✓ | ✓ | ✓ | ✓ | 2018 |
| RNA-QC-Chain [18] | C++ | fq,fa | ~ | ✓ | ✓ | ~ | ✓ | 2018 |
| afterQC [5] | C,python | fq | ✓ | ~ | ✓ | × | ✓ | 2017 |
| Cutadapt [12] | C,python | fq,fa,gz | × | ~ | ✓ | × | ✓<sup>1</sup> | 2011 |
| trimmomatic [3] | java | fq,gz | × | ✓ | ✓ | × | ✓ | 2014 |
| Biobloom [6] | C++ | BAM/SAM,fa,fq* | × | × | × | ✓ | ✓ | 2014 |
| FastQC [1] | java | fq,gz | ✓ | × | × | × | × | 2010 |
| SolexaQA++ [7] | C++,R | fq,gz | ✓ | ✓ | × | × | × | 2010 |
| RSeQC [16] | C,python | BAM/SAM | ✓ | × | × | × | × | 2012 |
| RNA-SeQC [8] | java | BAM | ✓ | × | × | ~ | × | 2012 |
| QoRTs [10] | java,R | BAM | ✓ | × | × | × | × | 2015 |

#### FastqPuri efficiently generates comprehensive sequence quality reports

Only a fraction of the tools that deliver quality control plots on RNA-seq data do so before read alignment, that is on fastq files: afterQC, FastQC and Solex-aQA++. RNA-QC-Chain has a quality control executable, but does not generate any plots. In terms of computer performance and memory usage, we compared **FastqPuri** with afterQC, FastQC, RNA-QC-Chain and SolexaQA++.

We ran the above mentioned tools on fastq files from three different datasets representing different sequence name formats and quality encodings. We also ran them with different input formats in parallel: fastq, gz and bz2. We compare the performance of running all programs on the uncompressed file, the gz-compressed file and `Qreport` running on the bz2-compressed file. For benchmarking tools which do not accept compressed input, we ran the tool on uncompressed data and added the time for decompressing the file to their timings. The performances in terms of time are shown in Figure 3. FastqPuri’s `Qreport` was substantially faster than all of the other tools when using bz2 files, by a factor of at least 2. `Qreport` and AfterQC were always faster than the other tools, but AfterQC failed to analyze fastq data in Illumina 1.3+ format with quality scores encoded with Phred+64. RNA-QC-Chain failed whenever data was in paired-end format. We profiled peak memory usage with the same datasets and show the results in Figure 4. While some QC tools have quite high peak memory demands, FastqPuri’s `Qreport` and AfterQC had the lowest peak memory usage, with `Qreport` outperforming all other tools on all datasets.

**Figure 3:**
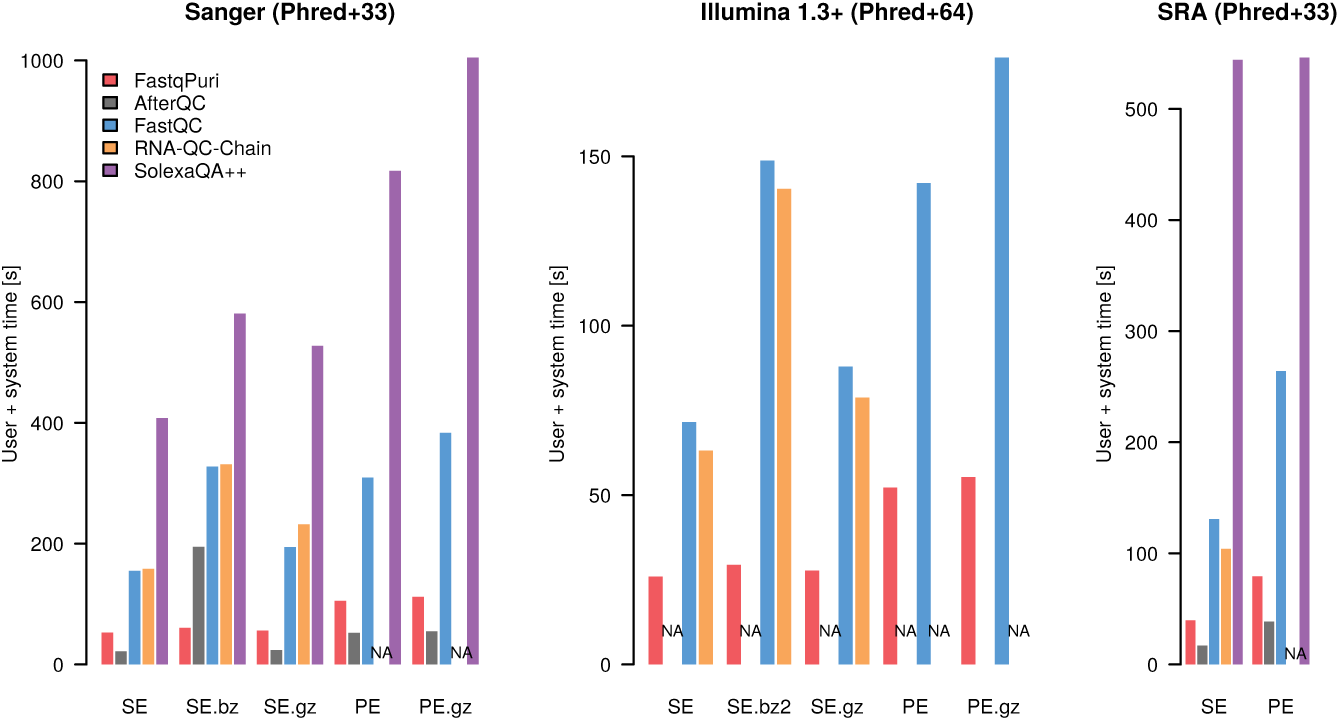
Run times (user plus CPU time in seconds) of FastqPuri’s Qreport versus other tools for three different datasets. The datasets represent different quality encodings (Phred+33 and Phred+64) as well as different sequence name formats. Timings for SolexaQA++ on Illumina 1.3+ data are not shown because the smallest value was around 10 minutes and all other values became invisibly small on that scale.

**Figure 4:**
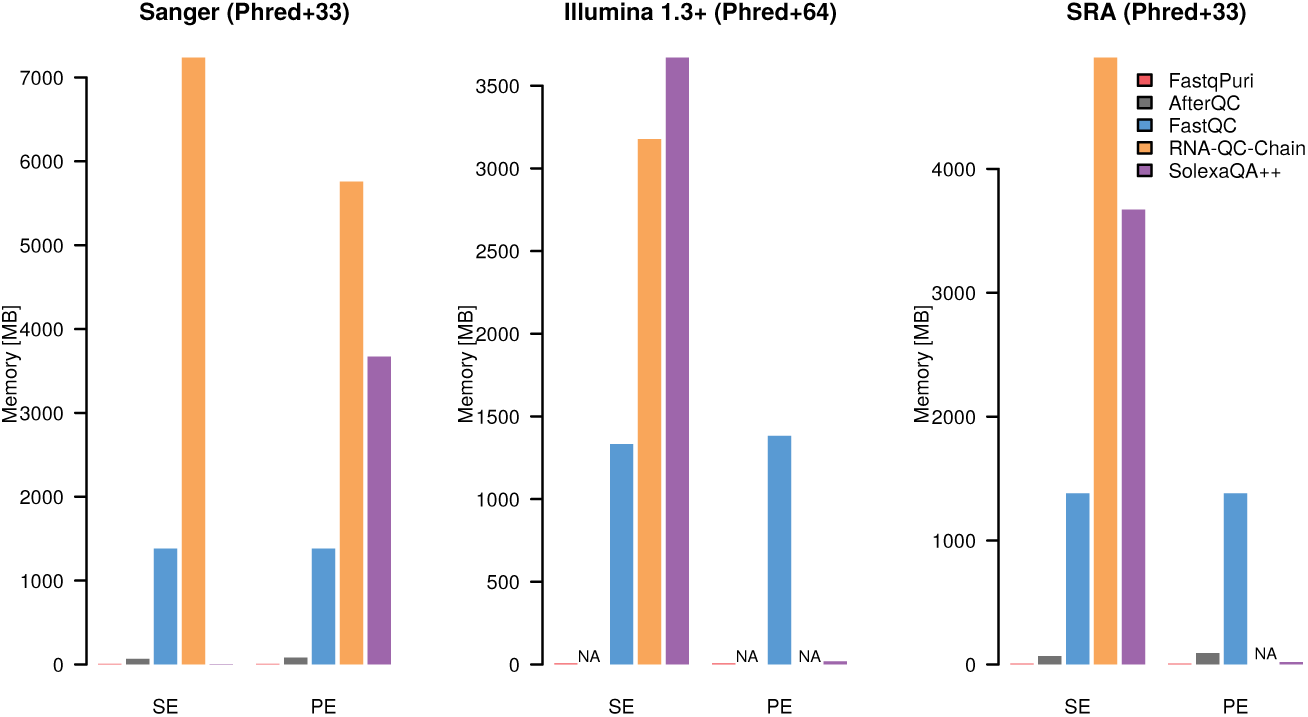
Memory usage (in MB) of FastqPuri’s Qreport versus other tools for three different datasets. The datasets represent different quality encodings (Phred+33 and Phred+64) as well as different sequence name formats.

#### FastqPuri outperforms trimmomatic in adapter trimming

We benchmarked adapter trimming with **FastqPuri** and with trimmomatic, the adapter trimming tool that performed best on paired- and single-end data in terms of speed and PPV (positive predictive value), albeit at the cost of large peak memory requirements [11]. We ran both tools on dataset 3 (see Table 3), once on the forward reads representing single-end data and once on both forward and reverse reads representing a paired-end dataset. The time spent for both compressed and uncompressed output is shown in Figure 5. FastqPuri’s `trimFilter(PE)` was substantially faster than trimmomatic for both single-end and paired-end data, with running times of 4-22% of the ones of trimmomatic. For bz2 files, the speed-up was most pronounced and `trimFilter` needed only 4% of the time of trimmomatic to process a single-end read file. The peak memory used by trimmomatic was about 32 GB, while `trimFilter(PE)` needed only between 8 and 9 MB, which is less than 3% of the peak memory of trimmomatic. Thus, **FastqPuri** outperformed trimmomatic in both consumed time and peak memory usage.

**Figure 5:**
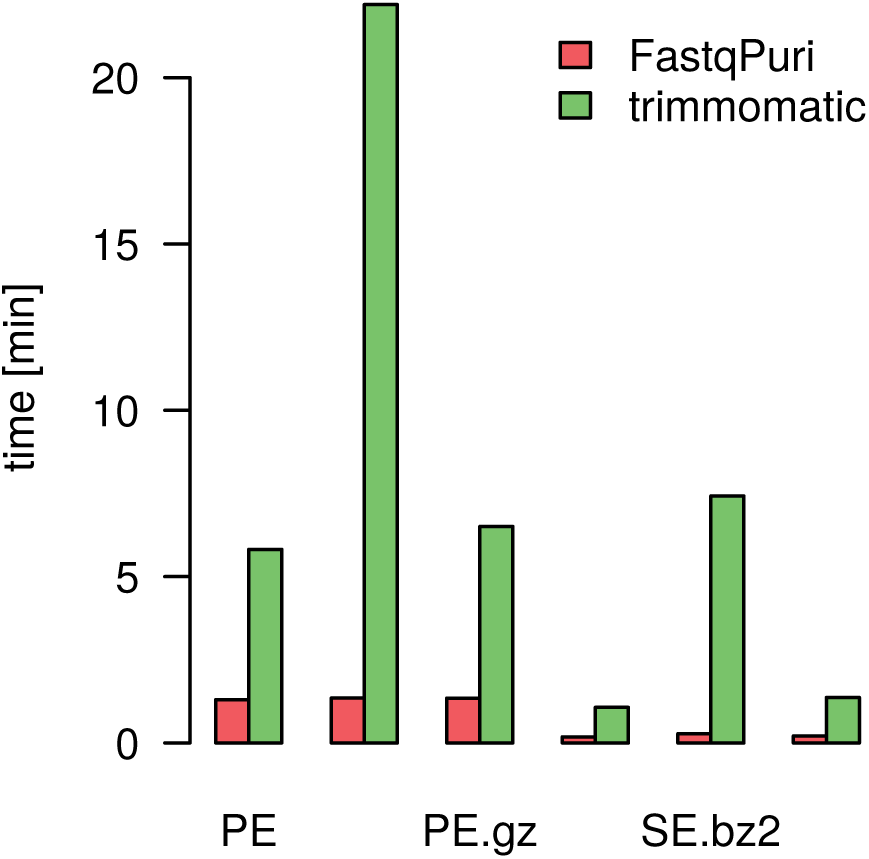
Run times (user plus CPU time in seconds) of FastqPuri’s trimFilter and trimFilterPE to remove adapter sequences versus trim-momatic.

#### FastqPuri efficiently filters contaminations with the tree method

We ran `trimFilter` on a human RNA-seq dataset (dataset 1) and `trimFilterPE` on a microalgae (*Nannochlorpsis oceanica*) dataset (dataset 3), searching for human rRNA contamination. We ran RNA-QC-Chain on the same datasets, as this tool specifically identifies and removes rRNA. The time taken and peak memory usage of both tools on the two datasets is shown in Figure 6. FastqPuri’s `trimFilter(PE)` clearly outperformed RNA-QC-Chain for both fastq and compressed input formats in terms of time (upper panel) and peak memory (lower panel) usage. In dataset 1, `trimFilter` detected 1 334 045 rRNA reads while RNA-QC-Chain found only 192 839 reads which were predicted to originate from 28 S rRNA transcripts. RNA-QC-Chain searches against an in-built database of 16/18S and 23/28S sequences, while we used the complete human rRNA gene cassette for filtering. Therefore, it is highly likely that RNA-QC-Chain missed many sequence reads originating from human rRNA.

**Figure 6:**
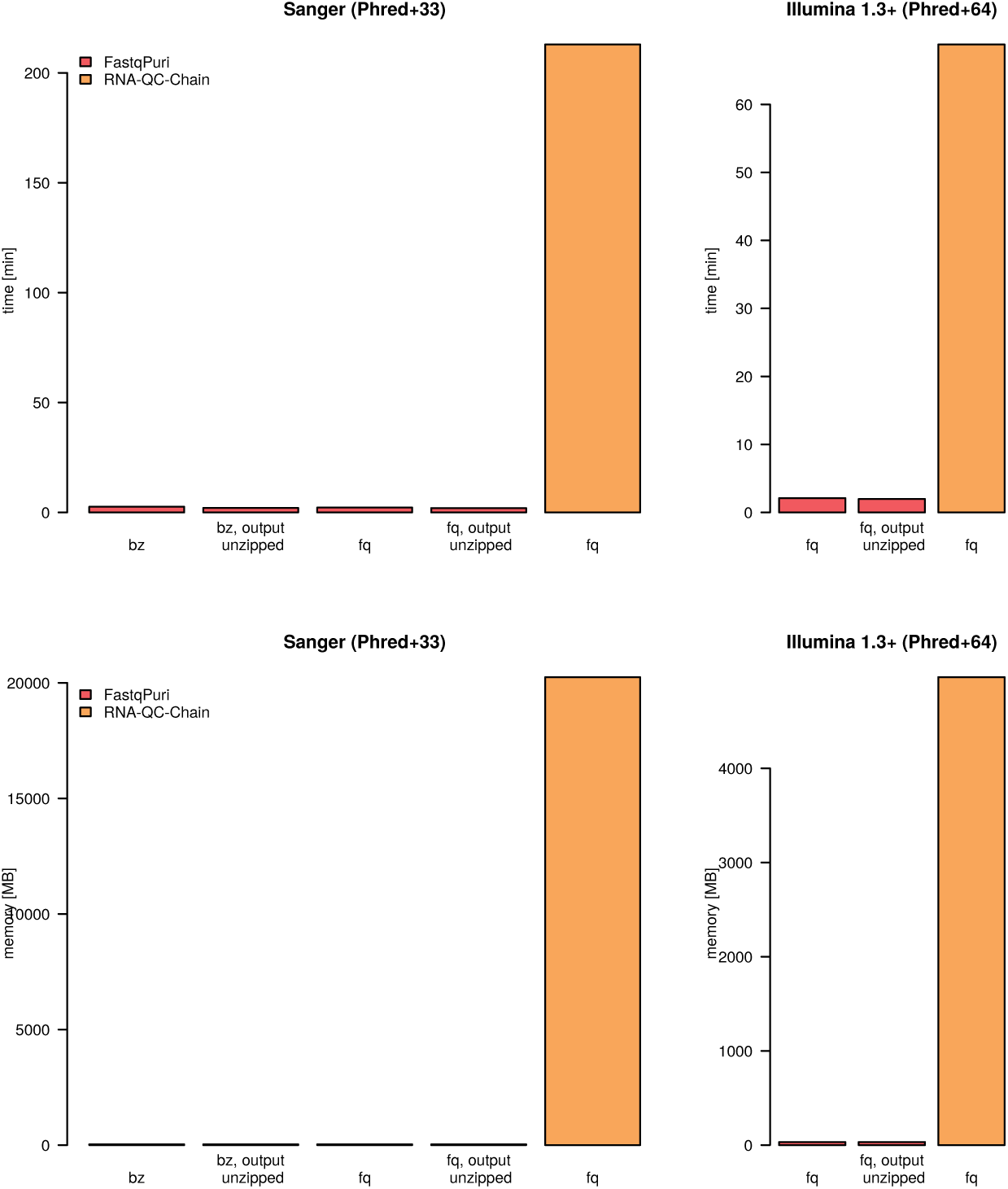
Run times (user plus CPU time in seconds) and memory usage (in MB) of FastqPuri’s trimFilter and RNA-QC-Chain to remove reads from human rRNA transcripts.

In dataset 3, FastqPuri attributed 8 519 sequence reads to human rRNA transcripts, while RNA-QC-Chain predicted 21 012 transcripts derived from 28 S rRNA and 18 626 reads from 18 S rRNA. This difference can again be explained by the different reference sequences being used to detect rRNA contamination.

#### Filtering contaminations with the bloom filter method are on a level with existing methods

We compared the computer performance of **FastqPuri** with BioBloom [6] for the bloom filter creation and removal of long contaminating sequences. First we simulated a contaminated human dataset by sampling reads from the human transcriptome and adding simulated reads from the mouse transcriptome (details in methods). Then, we created a bloom filter on the mouse genome to filter out the contaminating mouse reads. The performance and memory peak usage of creating the bloom filter and classifying reads as contamination are summarized in Table 2. **FastqPuri** was faster in generating the bloom filter, but slower in classifying reads than BioBloom. Since making the bloom filter took longer than classifying the reads, **FastqPuri** was faster when summing up the time of these two steps. In terms of peak memory usage, BioBloom used less memory than **FastqPuri** when generating the bloom filter, and the same peak memory when classifying reads. In terms of sensitivity and specificity of **FastqPuri** and BioBloom, both methods performed equally well, with **FastqPuri** being slightly better in terms of sensitivity (0.998 versus 0.993) and BioBloom in terms of specificity (0.932 versus 0.937).

**Table 2:** Timings on removing biological contaminations with **FastqPuri** and BioBloom. ‘Bloom maker’ refers to generating the bloom filter, ‘Contaminations’ refers to classifying the reads using the bloom filter.

| user time | Bloom maker | Contaminations |
| --- | --- | --- |
| **FastqPuri** | 32m45s | 4m10s |
| **BioBloom** | 41m28s | 2m58s |
| CPU time | Bloom maker | Contaminations |
| **FastqPuri** | 0m9.2s | 0m4.4s |
| **BioBloom** | 0m7.1s | 0m5.4s |
| peak mem | Bloom maker | Contaminations |
| **FastqPuri** | 5.78GB | 3.24GB |
| **BioBloom** | 3.24GB | 3.24GB |

### Discussion

RNA-seq is currently widely used to assess transcript and gene expression levels. Fast transcript counting methods render sequence data quality control and preprocessing the most time demanding steps in data analysis. Moreover, since transcript counting methods such as salmon and kallisto do not take quality scores into account when searching k-mers in reads, sensible quality-control is necessary. The novel quality plots allow the user to make informed choices about quality filtering and data discarded at different quality thresholds. The QC report generated by **FastqPuri** is most informative on Illumina sequence data containing tile information in the sequence name. If this is missing, plots showing qualities per tile are omitted. **FastqPuri** can also process long reads. Read length longer than 400 nt require passing the maximum read length while compiling **FastqPuri**. For read length of several kilobases, however, it might be inconvenient to inspect the plots per base position.

We compared **FastqPuri** with existing tools, although none of them covered all steps provided by **FastqPuri**. We focused our benchmarkings on tools that were designed to preprocess RNA-seq data, as this was also our intention. Benchmarking against all available tools for each of the individual steps downstream of QC was infeasible, so we focused on the most popular and most efficient ones (cutadapt, trimmomatic). We found that the **FastqPuri** modules for quality control and sequence filtering outperformed existing tools in terms of comprehensiveness, versatility and computational efficiency. For example, **FastqPuri** was the fastest tool to generate a QC report on bz2 files and had the lowest peak memory usage for all input formats. Summarizing over different quality score and compression formats, **FastqPuri** was significantly faster than existing tools in generating QC plots.

**FastqPuri** was substantially faster and more memory-efficient than trimmomatic in removing adapter sequences, while it can also search for and remove reads stemming from contaminating loci or species, such as rRNA or host and pathogen contaminations.

Searching for rRNA contaminations, **FastqPuri** outperformed the Hidden Markov Model approach used in RNA-QC-Chain and allowed more flexibility as the user can decide which sequences (in terms of species and locus) should be filtered out. **FastqPuri** also more efficiently removed contaminating reads, e.g. reads from anywhere within the rRNA while RNA-QC-Chain only searched for particular regions (16/18S, 23/28S). Therefore, RNA-QC-Chain might be better suited to identify potential contaminating species than removing the contaminating sequences from the data. Using the BLOOM method to filter out potential contaminations using larger-sized files (e.g. genomes), **FastqPuri** was faster than BioBloom tools in generating the bloom filter but slightly slower in classifying sequences. Because generating the bloom filter takes more than 90% of the time, the summed time of both steps was shorter for **FastqPuri**. We chose a very challenging scenario by selecting mouse as contaminating (e.g. host) species for a human dataset. Because of high sequence similarity between the two species, perfect separation of the reads cannot be expected, and both tools performed equally well in terms of sensitivity and specificity.

For a complete preprocessing run on dataset 3, **FastqPuri** (with initial QC, adapter and low quality base removal, removal of reads originating from human rRNA, QC on filtered fastq file and a summary QC report), took 3 minutes and 3 seconds. In comparison, sequentially running FastQC, trimmomatic, RNA-QC-chain, and again FastQC on the filtered reads took more than 20 times longer (72 minutes and 15 seconds) and used a higher peak memory. Even if the time-consuming step of filtering rRNA was omitted, FastqPuri was still substantially faster, using 66 seconds, while the pipeline of existing tools took 3 minutes and 27 seconds. Therefore, we anticipate that **FastqPuri** will facilitate QC and preprocessing of RNA-seq data and speed-up the analysis of both small and large datasets.

## Methods

### Benchmarking details

#### Data sets

We benchmarked **FastqPuri** and existing tools with the following datasets: Dataset 1: single end reads generated from a human RNA sample. Dataset 2: paired end reads from *Arabidopsis thaliana*. Dataset 3: paired end reads from *Nannochloropsis oceanica* [18]. Dataset 4: paired end reads from *Homo sapiens* (SRA run SRR1216135). Dataset 5. simulated reads from *Homo sapiens* and *Mus musculus*. We generated 20 reads of length 100 nt for each transcript of the human and mouse transcriptomes (ensembl GRCh38 (human) and GRCm38 (mouse)) using the R package ‘polyester’ [9]. This resulted in approximately 2.3 million mouse and 3.7 million human reads which were assigned an arbitrary quality string with individual Q scores being larger than 27, and concatenated and shuffled before generating a fastq file. The mouse reads were considered contamination. The core properties of the datasets used for benchmarking are shown in Table 3.

**Table 3:**
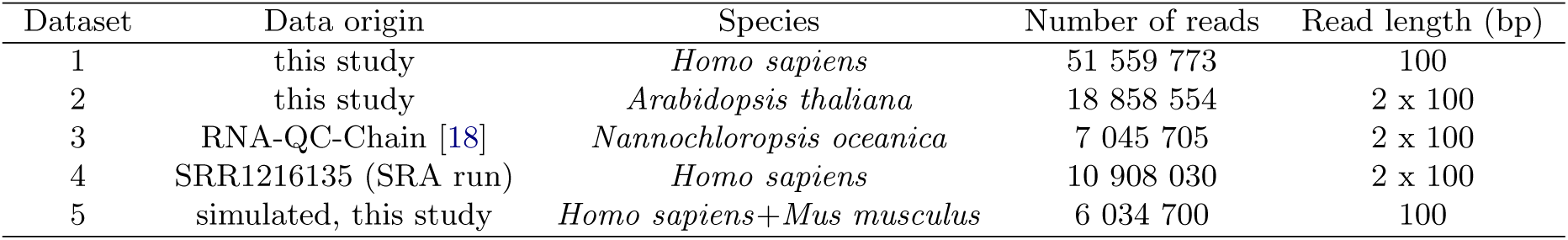
Datasets used for benchmarking.

| Dataset | Data origin | Species | Number of reads | Read length (bp) |
| --- | --- | --- | --- | --- |
| 1 | this study | *Homo sapiens* | 51 559 773 | 100 |
| 2 | this study | *Arabidopsis thaliana* | 18 858 554 | 2 x 100 |
| 3 | RNA-QC-Chain [18] | *Nannochloropsis oceanica* | 7 045 705 | 2 x 100 |
| 4 | SRR1216135 (SRA run) | *Homo sapiens* | 10 908 030 | 2 x 100 |
| 5 | simulated, this study | *Homo sapiens*+*Mus musculus* | 6 034 700 | 100 |

#### Tool settings

Tools were run with default parameters unless stated otherwise. Trimmomatic adapter trimming was performed with the adapter sequences provided by trimmomatic (TruSeq2-PE.fa for paired end data, TruSeq2-SE.fa for single end data). Trimmomatic was run with the following mismatch and score settings: ‘ILLUMINACLIP:TruSeq2-PE.fa:2:8:8’ for paired end data and ‘ILLUMINACLIP:TruSeq2-PE.fa:2:8:8’ for single-end data. `trimFilterPE` of **FastqPuri** was run with the same adapter sequences, allowing at most two mismatches and requiring an alignment score of at least 8 (TruSeq2-PE.fa:TruSeq2-PE.fa:2:8).

To filter reads originating from rRNA transcripts, we took the complete human ribosomal repeating unit (GenBank accession U13369.1), removed lines that contained non-{A, C, G, T} characters (8 out of 616 lines) and invoked **FastqPuri**’s `trimFilterPE` with –method TREE providing the rRNA sequence, a score threshold of 0.4 and an l-mer length of 25.

RNA-QC-chain searches against an internal database of rRNA sequences and because we wanted to remove human rRNA, we only searched against the 18S and 28S parts of the database.

To filter contaminations with the bloom filter approach, bloom filters of the mouse genome (mm10) were generated with a false-positive rate of 0.0075 and k-mers of length 25 nt for both biobloommaker (BioBloom) and makeBloom (**FastqPuri**). Reads of the simulated dataset were then classified setting the score threshold at 0.15 for both tools.

#### Computing infrastructure

All tests were run on a Debian Linux Server, with Linux kernel version 3.16.43–2+deb8u2, with 2 Intel(R) Xeon(R) X5650 CPUs (12 cores, 2.67GHz) and 144GB RAM.

Time was measured using the ‘time’ command of bash. If not stated otherwise, we reported the sum of user and system (CPU) time. Peak memory usage of FastqPuri, RNA-QC-chain, and AfterQC was assessed with valgrind [13]. Tools that used scripts to invoke their executables were profiled with a custom script based on monitoring memory usage of the active process with the bash command ‘ps’ every second. We used the later approach for FastQC, SolexaQA++, trimmomatic, and BioBloomTools.

## Conclusions

We presented a light-weight high-throughput sequence data preprocessing tool, **FastqPuri**. **FastqPuri** was designed for RNA-seq data intended for transcript counting, but it is also applicable to other kinds of fastq data. **FastqPuri** is fast and has a low memory footprint, can be used in pipelines or stand-alone, combines all preprocessing steps needed to apply transcript counting: QC, adapter and quality filtering and filtering biological contaminations as well as QC on the filtered data. **FastqPuri** provides a range of useful graphics, including novel ones, to make informative choices for sequence quality-based read trimming and filtering, which is performed by **FastqPuri** subsequently. In comparison to existing tools which cover parts of the steps performed by **FastqPuri**, **FastqPuri** was more time and memory efficient over a range of currently used quality encoding and compression formats. Therefore, **FastqPuri** widens the bottleneck of time- and memory consuming preprocessing steps in RNA-seq data analysis, allowing higher throughput for large datasets and speeding up preprocessing for all datasets.

## Supporting information

## Availability and requirements

**Project name: FastqPuri**

**Project home page:** https://github.com/jengelmann/FastqPuri

**Programming language:** C, R (for the html reports)

**Operating systems:** Unix/Linux, Mac OS, OpenBSD

**Licence:** GPL v3

**Other requirements:** cmake (at least version 2.8), a C compiler supporting the c11 standard (change the compiler flags otherwise), pandoc (optional), Rscript (optional), R packages pheatmap, knitr, rmarkdown (optional).

**Data availability:** The datasets used for benchmarking are available in NCBI’s SRA (Sequence Read Archive), run number SRR1216135 (dataset 4), the website of RNA-QC-chain (http://bioinfo.single-cell.cn/Released_Software/rna-qc-chain/data.tar.gz, dataset 3), or from the corresponding author on request (dataset 1, 2 and 5).

## Ethics approval and consent to participate

Sampling of human cells has been carried out in accordance with The Code of Ethics of the World Medical Association (Declaration of Helsinki).

## Consent for publication

Not applicable.

## Author’s contributions

PPR and JCE conceived and designed **FastqPuri**. PPR implemented the tool. CL evaluated the software. PPR and JCE wrote the manuscript.

## Acknowledgements

We thank Maria Attenberger and Phu Tran for proof-reading the user manual and testing the software.

## Additional Files

### Additional file 1 — Supplementary text

Supplementary text with details on feature implementation and benchmarking. PDF file.

### Additional file 2 — Archive of FastqPuri

Archive containing all files needed to install and run **FastqPuri**. Date stamp October 12, 2018.

