## Supplementary material for "FastqPuri: high-performance preprocessing of RNA-seq data"

[1] *Institute of Functional Genomics, University of Regensburg, 93053 Regensburg, Germany*
[2] *NIOZ, Royal Netherlands Institute for Sea Research, Department of Marine Microbiology and Biogeochemistry, and Utrecht University, P.O. Box 59, NL-1790 AB Den Burg, The Netherlands.*
(Dated: October 12, 2018)

### Overview

FastqPuri consists of 6 executables which can be run sequentially or individually. Some of them are alternatives for one another, some are optional.

- **Qreport** creates a quality report in `html` format. From a fastq input file and a quality threshold minQ $\in [0, 45]$ under which bases are considered low quality, by default, this threshold is set to 27. Graphs with the following information are created: (1) sequence quality per base position, (2) number of reads with at least $m$ low quality base calls, (3) low quality base calling proportion per nucleotide, per tile and per lane, (4) average quality per position per tile per lane, (5) low quality base calling proportion per position per tile per lane, (6) low quality base calling proportion per position for all tiles and lanes, (7) nucleotide content per position. Graphs (5) and (6) indicate how many reads will be discarded or trimmed at a certain quality threshold, information that cannot be retrieved from graphics showing averages. The data shown in the graphs is stored in a binary file used by **Sreport** for the summary report.
- **makeTree** takes a fasta file containing potential contaminating sequences, creates a tree data structure and saves it to a file to be used by trimFilter/trimFilterPE. Intended for small fasta files.
- **makeBloom** creates a bloom filter from potential contaminating sequences, and stores it in a file for use by trimFilter/trimFilterPE. Suited for large fasta files.
- **trimFilter** takes a fastq file as an input and filters out or trims reads according to the presence of: (1) adapter remnants, (2) contaminations from unwanted genomes, (3) low quality base calls, (4) N's. The output consists of a fastq file containing the reads classified as *good* (left unmodified or trimmed), a fastq files with discarded reads (one per filter option selected) and a binary file with summary statistics.
- **trimFilterPE** filters paired end data in an analogous way to **trimFilter**. It takes two fastq files as input, and applies the selected filters on each read. If one of the reads of a pair does not pass a given filter, both reads are discarded. The output is a fastq file of trimmed reads for each input file and a binary output file as in **trimFilter**, with the only difference that the number of trimmed reads is counted once per each input file.
- **Sreport** generates summary reports in `html` format from a list of **Qreport** binary output files or **trimFilter/trimFilterPE** binary output files. This produces an overview of all samples in a dataset.

We suggest to run FastqPuri on single-end data with a small file of potential contaminations, e.g. rRNA, like so: Qreport-Sreport-makeTree-trimFilter-Qreport-Sreport; and on paired-end data with a large sequence file of potential contaminations, e.g. several genomes, like so: Qreport-Sreport-makeBloom-trimFilterPE-Qreport-Sreport. Of course, FastqPuri can also filter single-end data with a bloom filter and paired-end data with a tree-based filter. Figure 1 of the main paper shows a scheme of the workflow of FastqPuri.

### QUALITY REPORT

The quality report is obtained byexecuting **Qreport**. It reads a fastq file and generates a html report with the information/graphs described in FIG. 1, FIG. 2. It accepts raw fastq files and files that were already filtered. A

| Var | Value |
| --- | --- |
| Input file name | test_output.bin |
| Read length | 51 |
| Min good quality | 27 |
| Number of reads | 500000 |
| Number of highQ reads | 336682 |
| Number of tiles | 20 |
| Number of lanes | 2 |
| Qualities | 2 (#), 14 (/), 22 (7), 27 (<), 33 (B), 37 (F) |
| Reads with N's | 812 |
| Number of N's | 1114 |

(a)General information

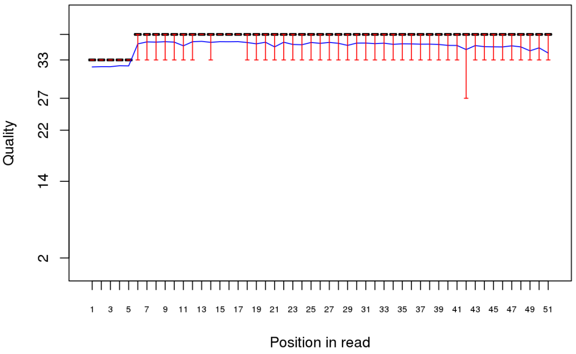


(b)Per base sequence quality box plots. The blue line corresponds to the quality mean value.

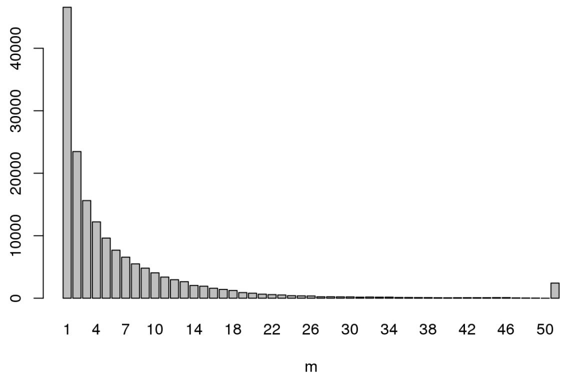


(c)Number of reads with $m$ low quality cycles

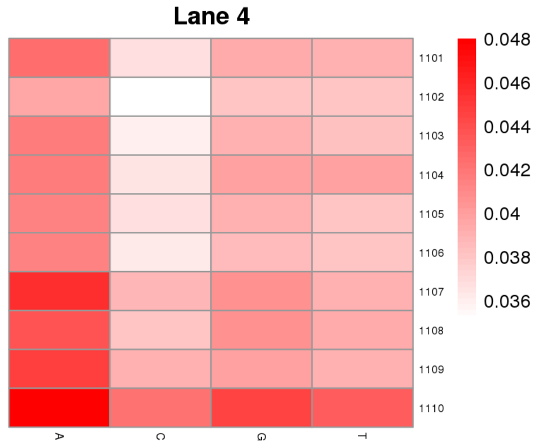


(d)Fraction of low quality bases {A, C, G, T} per position, per tile and per lane

FIG. 1: **Qreport** results (I)

nucleotide in a read is considered to have low quality if it lies below a user defined threshold. See `README_Qreport.md` for more details on the input parameters. The quality values take the Phred+33 convention.

We have tested the performance of **Qreport** vs **fastQC** [1] using an input fastq file with 51559773 reads, of read length 51, compressed with `bzip2`. Since **fastQC** does not accept this format as an input, we compare the performance running both programs on the uncompressed file and **Qreport** running on the compressed file versus the time taken for the file to be uncompressed plus the time taken by **fastQC** to run. The performance and peak memory usage are collected in TABLE I.

### CONTAMINATION OF TECHNICAL SEQUENCES

The presence of technical sequences, such as adapter remnants, in next generation sequencing (NGS) data can affect the alignment and pseudo-alignment/quasi-mapping. This tool offers the possibility of identifying such remnants through the option `--adapter` in the executables **trimFilter** and **trimFilterPE** for single-end and paired-end data, respectively. The approach used is very similar to trimmomatic [2]. Figure FIG. 3 sketches the possible situations for single-end data.

Reads are scanned from the 3' end to the 5' end, with a 16 nucleotides long seed. If only up to a user defined number of mismatches is encountered, the seed is accepted and a local alignment is performed. A score is constructed by adding $\log_{10}(4)$ when the bases match and penalizing with $-Q/10$ for mismatches. If the score exceeds a user

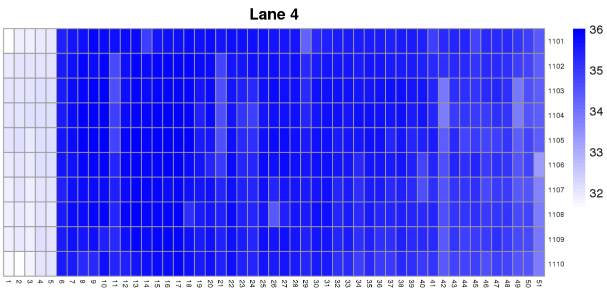

(a)Cycle average quality, per tile, per lane

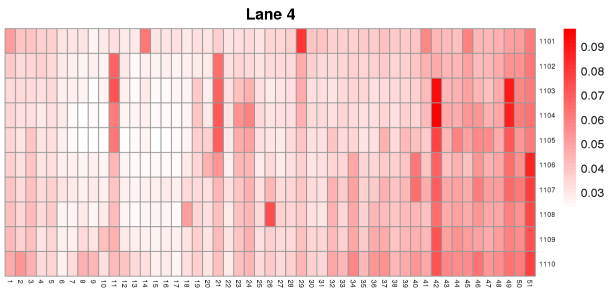

(b)Low quality cycle proportion, per tile, per lane

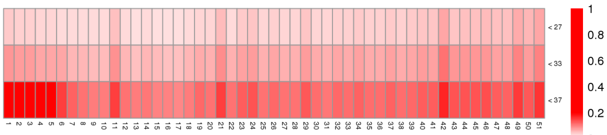
(c)Proportion of cycles lying below a certain quality, for all tiles, all lanes

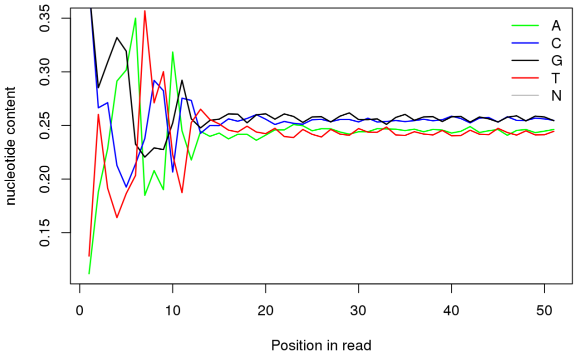

(d)Nucleotide content per position

FIG. 2: **Qreport** results (II)

| | Input | |
|---|---|---|
| real time | Uncompressed | Compressed |
| **Qreport** | 1m35s | 4m43s |
| **fastQC** | 3m16s | 7m27s |
| CPU time | Uncompressed | Compressed |
| **Qreport** | 1m32s | 1m37s |
| **fastQC** | 3m11 | 3m11 |
| mem peak | Uncompressed | Compressed |
| **Qreport** | 9.2MB | 9.2MB |
| **fastQC** | 1.4GB | 1.4GB |

TABLE I: Performance and peak memory usage of **Qreport** vs **fastQC**

defined threshold, it is accepted as adapter contamination. At the ends of the read, a seed of 8 nucleotides is also allowed, with the same score and number of mismatches restrictions. However, an alignment of at least 12 nucleotides is always demanded. If the read presents adapter contamination, it is trimmed if the remaining sequence length is at least the user defined minimum allowed length and discarded otherwise.

The alignments tested in paired-end data are illustrated in FIG. 4. As for single-end data, a 'seed-and-extend' method is applied. We use 16 nucleotide long seeds and when a valid one is found, i.e. a seed with at most the user defined number of mismatches, a score is computed, in the very same way as in the single-end case. If the score exceeds the user defined threshold, both reads are trimmed if the remaining subsequences lengths are allowed and discarded otherwise.

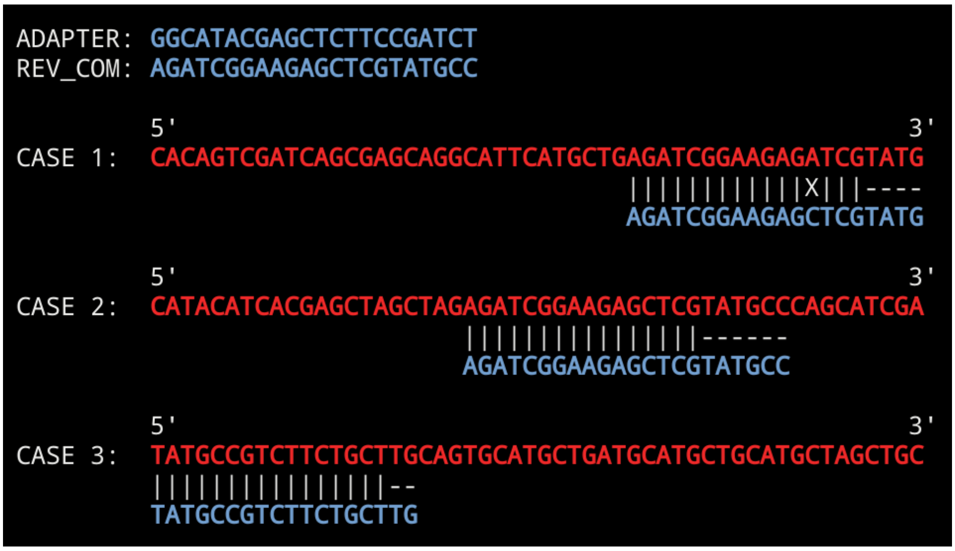


FIG. 3: Adapter contaminations in single-end mode: (1) partial overlap at the 3' end of the read, (2) full overlap at all positions, (3) partial overlap at the 5' end of the read.

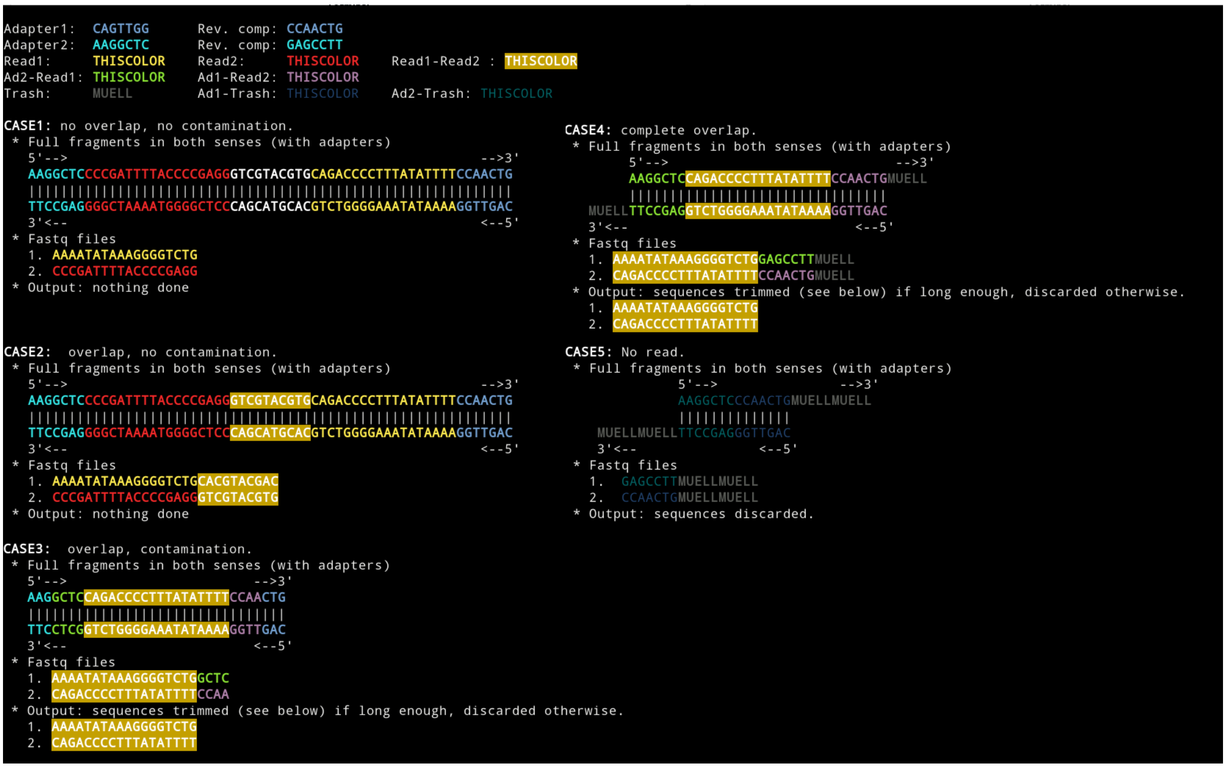


FIG. 4: Adapter contaminations in paired-end mode. The adapters are included in the fragments: (1) read 1 and read 2 have no overlap, (2) the overlap shows no read-through into the adapters, (3) the overlap between the reads present a partial adapter read through, (4) reads overlap including a whole read-through into the adapters, (5) there are no reads, only adapters, fragments contain no useful information.

To check the performance of the code, we have compared our tool with trimmomatic [2] for both single and paired-end data using fastq file(s) with 51559773 reads, of read length 51, measuring the real time, the CPU time and the peak memory usage. Results are listed in TABLE II. We see that most of the time is actually invested in compressing the output file.

| | Compressed output | | Unompressed output | |
|---|---|---|---|---|
| real time | Single-end | Paired-end | Single-end | Paired-end |
| **trimFilter(PE)** | 9m13s | 9m51s | 1m50s | 3m05s |
| **trimmomatic** | 11m13s | 21m53s | 3m27s | 5m40s |
| CPU time | Single-end | Paired-end | Single-end | Paired-end |
| **trimFilter(PE)** | 54s | 2m15s | 46s | 2m52s |
| **trimmomatic** | 10m50s | 21m21s | 3m24s | 10m50s |
| peak mem | Single-end | Paired-end | Single-end | Paired-end |
| **trimFilter(PE)** | 9.2MB | 9.2MB | 9.2MB | 9.2MB |
| **trimmomatic** | 34GB | 34GB | 34GB | 34GB |

TABLE II: Performance (minutes, seconds) and memory usage of **trimFilter(PE)** looking for adapter contaminations vs **trimmomatic** on single and paired-end data. Benchmarks were taken for both compressed and uncompressed output.

### CONTAMINATION OF UNWANTED GENOMES

Sometimes we want to remove reads belonging to undesired biological sequences, such as pathogen or host genomes, or potential contaminations that might interfere with subsequent analysis. This is provided as an option in **trimFilter** and **trimFilterPE**; we offer two options depending on the length of unwanted sequences exceeding 10MB.

#### Short contaminations sequence: 4-ary tree

If the unwanted sequences fasta file is smaller than 10MB, then one can look for contaminations by constructing a 4-ary tree out of the fasta file. One could use the executable **makeTree** to construct a tree and save it to disk, if one wanted to run the code on several samples. However, constructing the tree is in general a relatively cheap task for the sequence lengths under consideration and it does not represent a large overhead. Therefore, we can call **trimFilter(PE)** with the option `--method TREE` and it will be generated "on the flight".

When constructing a tree, the user not only has to pass the fasta file as an input, but also the depth of the tree and, when looking for contaminations, a score threshold is needed as well. A 4-tree, with {A, C, G, T} as leaves and a user defined depth $k$ is constructed including all $k$-mers in the unwanted genome. Then, all $k$-mers in a given read are tested to be in the tree. A read will be considered to be a contamination if the score computed as the proportion of $k$-mers found in the tree is larger than the score threshold. This method is very fast, every search is $O(2k*(L-k+1))$, where $L$ is the read length and the factor 2 appears because the reverse complement has to be checked also. However, it has the drawback that is very memory intensive. This is why we have limited the construction of a tree to sequences with sizes below 10MB.

#### Long contaminating sequences: bloom filter

This method is designed for large sequences up to 4GB. In this case, it is sensible to construct the bloom filter and store it in a file. This is achieved with the executable **makeBloom**. Then, **trimFilter(PE)** can be called with the option `--method BLOOM` and the code looks for contaminations provided a score threshold is passed as an input.

##### *Creating a bloom filter and checking elements*

A bloom filter is a probabilistic data structure used to test if an element is a member of a set. False positive matches are possible but false negative are not. Let $n$ be the number of elements in the set $S$, which in our case is the number of all possible $k$-mer in the sequence. Then, it is proceeded as follows:

- Decide on the number $g$ of hash functions we will use for the construction of the filter and the length of the filter $m$ (number of bits). This choice will be made based on the false positive rate we want to achieve (see Parameter optimization),
- We create an empty bloom filter, $B$, i.e. an array of $m$‘ bits set to 0‘. For every element in the set, $s_\alpha \in S$, compute the $g$ hash functions, $H_i(s_\alpha) \forall i \in \{1, \ldots, g\}$ and set the corresponding bits to 1 in the filter, i.e.,

$$B[H_i(s_\alpha) \mod m] = \begin{cases} 1 & \forall i \in \{1, \ldots, g\} \\ 0 & \text{otherwise} \end{cases}$$

- Then, if we want to check whether an element (a $k$-mer from a fastq read) $s_\beta$ is in the set $S$, we compute $H_i(s_\beta) \forall i \in \{1, \ldots, g\}$ and check whether all coresponding positions in the filter are set to 1, in which case we can say that $s_\beta$ might be in the set. Otherwise it is definitely not in the set.
- A read is discarded if the fraction of its $k$-mers that gave a positive result when looking for them in the filter exceeds the user defined threshold.

In this process, we do not consider $k$-mers with N's.

*Parameter optimization*

We choose the parameters so that the desired false positive rate is achieved. Alternatively, we can pass the filter size, and then the number of hash functions to be used is tuned so that the false positive rate is minimized.

We assume the hash functions select all positions with the same probability. The probability that a bit in the filter $B$ is not set to 1 after inserting an element using $g$ hash functions is $(1 - \frac{1}{m})^g$. If we insert $n$ elements, the probability that an element is still 0 is $(1 - \frac{1}{m})^{gn}$, and the probability of a bit being 1 is $1 - (1 - \frac{1}{m})^{gn}$. The false positive rate, i.e. the probability that all positions computed from the hash functions for a given element are zero without it being in the set is given by,

$$p(g, n, m) = \left[1 - (1 - \tfrac{1}{m})^{gn}\right]^g \sim \left(1 - e^{\frac{gn}{m}}\right)^g$$

For a given $n$ and $m$, the value of $g$ that minimizes $p$ is,

$$\frac{dp(g,n,m)}{dg} = 0 \quad \Rightarrow \quad g = \frac{m}{n} \log(2)$$

The optimal number of bits $m$ per element assuming the optimal number of hash functions being used is,

$$\frac{m}{n} = -\frac{\log(p)}{\log^2(2)}.$$

The memory usage will be determined then by $m$. In FIG. 5, we can see both, the optimal number of bits per element and the optimal number of hash functions as a function of the false positive rate.

As an example, let's assume we want to look for contaminations in a genome of ∼3Gbps and want to keep the false positive rate at 2%. Then, we will need a filter of ∼3.05GB.

The sensitivity can be increased by decreasing the $k$-mer size and the score threshold. False negatives only occur in the presence of mismatches due to variants, or errors in the base calling procedure, since the filter itself does not allow for false negatives. To increase specificity one can increase the score threshold or, obviously reduce the positive rate as input parameter.

*Benchmarkings*

In addition to the human dataset used in the main manuscript, we also checked the performance of filtering out reads from contaminating species by using **dgwsim** simulated data. We generated $10e5$ 150bp *Escherichia coli* reads and $10e5$ 150bp human reads, and created several bloom filters for the *Escherichia coli* genome with $k$-mer size 25 and false positive rate $p = 0.075$. Then, we run **trimFilter** on the E.coli-human data to look for contaminations using scores ranging from 0.05 to 0.2. FIG. 6 displays a ROC curve summarizing the results.

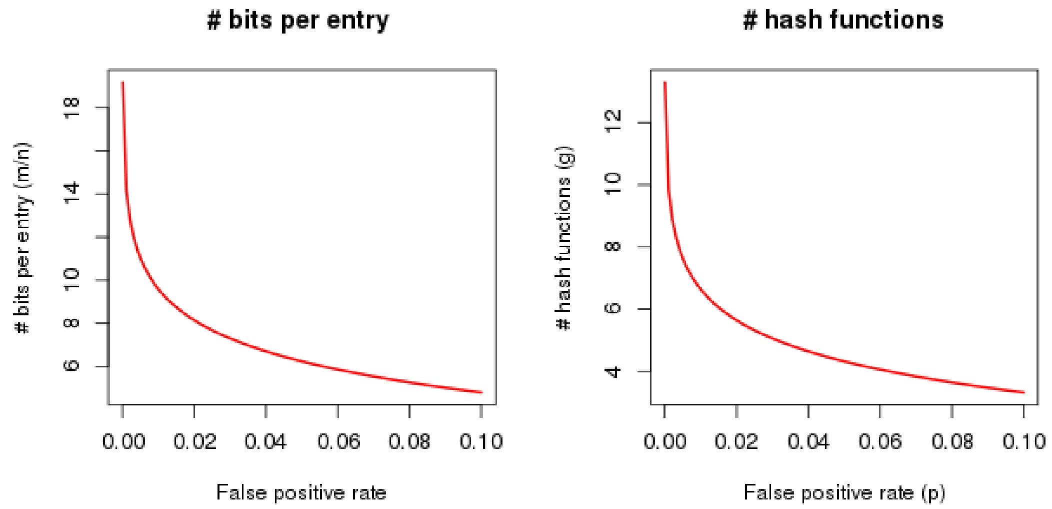


FIG. 5: Optimal number of bits per element (left) and number of hash function (right) in a bloom filter as a function of the false positive rate

### QUALITY FILTER

Fastq files may contain reads with low quality cycles that might lead to alignment of the read at the wrong place or counting the read for the wrong transcript. This tool allows the user to pass a quality threshold as an input, under which cycle qualities are considered to be poor, and the flag `--lowQ` followed by one of the options,

- `NO:` (or flag absent): nothing is done to the reads with low quality,
- `ALL:` all reads containing at least one low quality nucleotide are discarded,
- `ENDS:` look for low quality base callings at the beginning and at the end of the read. Trim them at both ends until the quality is above the threshold. Keep the read if the length of the remaining part is at least the minimum allowed. Discard it otherwise,
- `FRAC [--percent p]:` discard the read if there are more than p% nucleotides whose quality lies below the threshold,
- `ENDSFRAC [--percent p]:` first trim the ends as in the `ENDS` option. Accept the trimmed read if the number of low quality nucleotides does not exceed p%, discard it otherwise.
- `GLOBAL --global n1:n2`: cut all reads globally `n1` nucleotides from the left and `n2` from the right.

Sometimes, we want to discard or trim reads containing nucleotides tagged as N's. This can be done by our tool by passing the flag `--NNN` and one of the following options,

- `NO:` (or flag absent): nothing is done to the reads containing N's,
- `ALL:` all reads containing at least one N are discarded,
- `ENDS:` N's are trimmed if found at the ends, left "as is" otherwise. If the trimmed read length is smaller than the minimal allowed read length, it is discarded.
- `STRIP:` Obtain the largest N free subsequence of the read. Accept it if its length is at least the minimal allowed length, discard it otherwise.

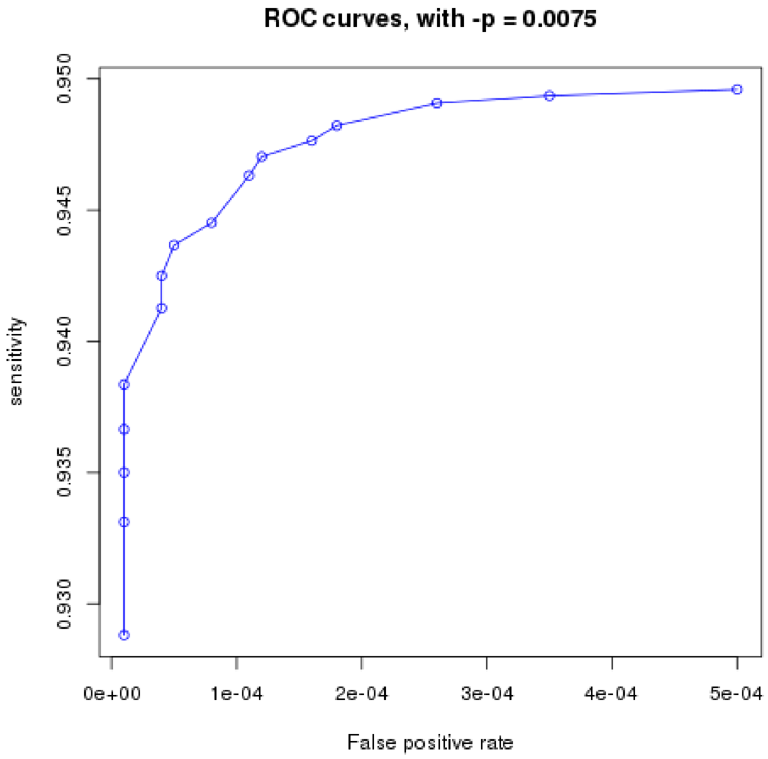


FIG. 6:

### SUMMARY REPORTS

The executable **SReport** creates three types of summary reports in html format:

**-t Q Quality summary report:** takes a folder containing `*.bin` files outputted by **Qreport** as input, and generates an html report with a table containing, for all samples: number of reads, number of tiles, percentage of reads with low quality cycles, percentage of reads with cycles tagged as N, and a heatmap showing the average quality per position for all samples.

**-t F Filtered data summary report, single-end:** takes a folder containing `*.bin` files outputted by **trimFilter** and generates an html report with a table specifying the setup of the run, i.e., the filters that were applied to the files if they were the same for all samples. If the files were generated with different setups, then the table is ambiguous and is therefore not generated. It also contains a table containing, for all samples (rows), the total number of reads, the number of accepted reads, percentage of reads discarded due to: adapter contaminations, unwanted genomes contaminations, low quality issues, presence of Ns, percentage of reads trimmed due to: adapter contaminations, low quality issues and presence of N's.

**-t D Filtered data summary report, paired-end:** it takes a folder containing `*bin` files outputted by **trimFilterPE** and generates an html report very similar to the one for single-end data, with the only difference that there are two columns containing trimmed percentages, i.e., there is a column for read 1 and another one for read 2.

#### *Data sets*

We benchmarked FastqPuri and existing tools with the following datasets: 1. Dataset 1 (same as in the main manuscript): 51 million paired-end reads generated from a human RNA sample. Read 1 of the dataset was also used to serve as a single-end library dataset. 2. We simulated $10e5$ 150nt single-end reads from the human genome (hg19) and simulated 10$e$5 150nt single-end *Escherichia coli* reads from the E.coli genome (HUSEC2011CHR1) with **dgwsim** (https://github.com/nh13/DWGSIM), leaving parameters not mentioned on default values.

### BENCHMARKING DETAILS

All tests were run on a Debian Linux Server, with Linux kernel version 3.16.43–2+deb8u2, with 2 Intel(R) Xeon(R) X5650 CPUs (12 cores, 2.67GHz) and 144GB RAM.

[1] S. Andrews, available online at http://www.bioinformatics.babraham.ac.uk/projects/fastqc (2010).
[2] A. M. Bolger, M. Lohse, and B. Usadel, Bioinformatics **30**, 2114 (2014), URL +http://dx.doi.org/10.1093/bioinformatics/btu170.
