## Supplementary material for "FastqPuri: high-performance preprocessing of RNA-seq data": Doxyfile

# Doxyfile 1.8.8

# This file describes the settings to be used by the documentation system
# doxygen (www.doxygen.org) for a project.
#
# All text after a double hash (##) is considered a comment and is placed in
# front of the TAG it is preceding.
#
# All text after a single hash (#) is considered a comment and will be ignored.
# The format is:
# TAG = value [value, ...]
# For lists, items can also be appended using:
# TAG += value [value, ...]
# Values that contain spaces should be placed between quotes (\" \").

#---------------------------------------------------------------------------
# Project related configuration options
#---------------------------------------------------------------------------

# This tag specifies the encoding used for all characters in the config file
# that follow. The default is UTF-8 which is also the encoding used for all text
# before the first occurrence of this tag. Doxygen uses libiconv (or the iconv
# built into libc) for the transcoding. See http://www.gnu.org/software/libiconv
# for the list of possible encodings.
# The default value is: UTF-8.

DOXYFILE_ENCODING      = UTF-8

# The PROJECT_NAME tag is a single word (or a sequence of words surrounded by
# double-quotes, unless you are using Doxywizard) that should identify the
# project for which the documentation is generated. This name is used in the
# title of most generated pages and in a few other places.
# The default value is: My Project.

PROJECT_NAME           = "FastqPuri"

# The PROJECT_NUMBER tag can be used to enter a project or revision number. This
# could be handy for archiving the generated documentation or if some version
# control system is used.

PROJECT_NUMBER         =

# Using the PROJECT_BRIEF tag one can provide an optional one line description
# for a project that appears at the top of each page and should give viewer a
# quick idea about the purpose of the project. Keep the description short.

PROJECT_BRIEF          =

# With the PROJECT_LOGO tag one can specify an logo or icon that is included in
# the documentation. The maximum height of the logo should not exceed 55 pixels
# and the maximum width should not exceed 200 pixels. Doxygen will copy the logo
# to the output directory.

PROJECT_LOGO           =

# The OUTPUT_DIRECTORY tag is used to specify the (relative or absolute) path
# into which the generated documentation will be written. If a relative path is
# entered, it will be relative to the location where doxygen was started. If
# left blank the current directory will be used.

OUTPUT_DIRECTORY       =

# If the CREATE_SUBDIRS tag is set to YES, then doxygen will create 4096 sub-
# directories (in 2 levels) under the output directory of each output format and
# will distribute the generated files over these directories. Enabling this
# option can be useful when feeding doxygen a huge amount of source files, where
# putting all generated files in the same directory would otherwise causes
# performance problems for the file system.
# The default value is: NO.

CREATE_SUBDIRS         = NO

# If the ALLOW_UNICODE_NAMES tag is set to YES, doxygen will allow non-ASCII
# characters to appear in the names of generated files. If set to NO, non-ASCII
# characters will be escaped, for example _xE3_x81_x84 will be used for Unicode
# U+3044.
# The default value is: NO.

ALLOW_UNICODE_NAMES    = NO

# The OUTPUT_LANGUAGE tag is used to specify the language in which all
# documentation generated by doxygen is written. Doxygen will use this
# information to generate all constant output in the proper language.
# Possible values are: Afrikaans, Arabic, Armenian, Brazilian, Catalan, Chinese,
# Chinese-Traditional, Croatian, Czech, Danish, Dutch, English (United States),
# Esperanto, Farsi (Persian), Finnish, French, German, Greek, Hungarian,
# Indonesian, Italian, Japanese, Japanese-en (Japanese with English messages),
# Korean, Korean-en (Korean with English messages), Latvian, Lithuanian,
# Macedonian, Norwegian, Persian (Farsi), Polish, Portuguese, Romanian, Russian,
# Serbian, Serbian-Cyrillic, Slovak, Slovene, Spanish, Swedish, Turkish,
# Ukrainian and Vietnamese.
# The default value is: English.

OUTPUT_LANGUAGE        = English

# If the BRIEF_MEMBER_DESC tag is set to YES doxygen will include brief member
# descriptions after the members that are listed in the file and class
# documentation (similar to Javadoc). Set to NO to disable this.
# The default value is: YES.

BRIEF_MEMBER_DESC      = YES

# If the REPEAT_BRIEF tag is set to YES doxygen will prepend the brief
# description of a member or function before the detailed description
#
# Note: If both HIDE_UNDOC_MEMBERS and BRIEF_MEMBER_DESC are set to NO, the
# brief descriptions will be completely suppressed.
# The default value is: YES.

REPEAT_BRIEF           = YES

# This tag implements a quasi-intelligent brief description abbreviator that is
# used to form the text in various listings. Each string in this list, if found
# as the leading text of the brief description, will be stripped from the text
# and the result, after processing the whole list, is used as the annotated
# text. Otherwise, the brief description is used as-is. If left blank, the
# following values are used ($name is automatically replaced with the name of
# the entity):The $name class, The $name widget, The $name file, is, provides,
# specifies, contains, represents, a, an and the.

ABBREVIATE_BRIEF       =

# If the ALWAYS_DETAILED_SEC and REPEAT_BRIEF tags are both set to YES then
# doxygen will generate a detailed section even if there is only a brief
# description.
# The default value is: NO.

ALWAYS_DETAILED_SEC    = NO

# If the INLINE_INHERITED_MEMB tag is set to YES, doxygen will show all
# inherited members of a class in the documentation of that class as if those
# members were ordinary class members. Constructors, destructors and assignment
# operators of the base classes will not be shown.
# The default value is: NO.

INLINE_INHERITED_MEMB  = NO

# If the FULL_PATH_NAMES tag is set to YES doxygen will prepend the full path
# before files name in the file list and in the header files. If set to NO the
# shortest path that makes the file name unique will be used
# The default value is: YES.

FULL_PATH_NAMES        = YES

# The STRIP_FROM_PATH tag can be used to strip a user-defined part of the path.
# Stripping is only done if one of the specified strings matches the left-hand
# part of the path. The tag can be used to show relative paths in the file list.
# If left blank the directory from which doxygen is run is used as the path to
# strip.
#
# Note that you can specify absolute paths here, but also relative paths, which
# will be relative from the directory where doxygen is started.
# This tag requires that the tag FULL_PATH_NAMES is set to YES.

STRIP_FROM_PATH        =

# The STRIP_FROM_INC_PATH tag can be used to strip a user-defined part of the
# path mentioned in the documentation of a class, which tells the reader which
# header file to include in order to use a class. If left blank only the name of
# the header file containing the class definition is used. Otherwise one should
# specify the list of include paths that are normally passed to the compiler
# using the -I flag.

STRIP_FROM_INC_PATH    =

# If the SHORT_NAMES tag is set to YES, doxygen will generate much shorter (but
# less readable) file names. This can be useful is your file systems doesn't
# support long names like on DOS, Mac, or CD-ROM.
# The default value is: NO.

SHORT_NAMES            = NO

# If the JAVADOC_AUTOBRIEF tag is set to YES then doxygen will interpret the
# first line (until the first dot) of a Javadoc-style comment as the brief
# description. If set to NO, the Javadoc-style will behave just like regular Qt-
# style comments (thus requiring an explicit @brief command for a brief
# description.)
# The default value is: NO.

JAVADOC_AUTOBRIEF      = NO

# If the QT_AUTOBRIEF tag is set to YES then doxygen will interpret the first
# line (until the first dot) of a Qt-style comment as the brief description. If
# set to NO, the Qt-style will behave just like regular Qt-style comments (thus
# requiring an explicit \brief command for a brief description.)
# The default value is: NO.

QT_AUTOBRIEF           = NO

# The MULTILINE_CPP_IS_BRIEF tag can be set to YES to make doxygen treat a
# multi-line C++ special comment block (i.e. a block of //! or /// comments) as
# a brief description. This used to be the default behavior. The new default is
# to treat a multi-line C++ comment block as a detailed description. Set this
# tag to YES if you prefer the old behavior instead.
#
# Note that setting this tag to YES also means that rational rose comments are
# not recognized any more.
# The default value is: NO.

MULTILINE_CPP_IS_BRIEF = NO

# If the INHERIT_DOCS tag is set to YES then an undocumented member inherits the
# documentation from any documented member that it re-implements.
# The default value is: YES.

INHERIT_DOCS           = YES

# If the SEPARATE_MEMBER_PAGES tag is set to YES, then doxygen will produce a
# new page for each member. If set to NO, the documentation of a member will be
# part of the file/class/namespace that contains it.
# The default value is: NO.

SEPARATE_MEMBER_PAGES  = NO

# The TAB_SIZE tag can be used to set the number of spaces in a tab. Doxygen
# uses this value to replace tabs by spaces in code fragments.
# Minimum value: 1, maximum value: 16, default value: 4.

TAB_SIZE               = 4

# This tag can be used to specify a number of aliases that act as commands in
# the documentation. An alias has the form:
# name=value
# For example adding
# "sideeffect=@par Side Effects:\n"
# will allow you to put the command \sideeffect (or @sideeffect) in the
# documentation, which will result in a user-defined paragraph with heading
# "Side Effects:". You can put \n's in the value part of an alias to insert
# newlines.

ALIASES                =

# This tag can be used to specify a number of word-keyword mappings (TCL only).
# A mapping has the form "name=value". For example adding "class=itcl::class"
# will allow you to use the command class in the itcl::class meaning.

TCL_SUBST              =

# Set the OPTIMIZE_OUTPUT_FOR_C tag to YES if your project consists of C sources
# only. Doxygen will then generate output that is more tailored for C. For
# instance, some of the names that are used will be different. The list of all
# members will be omitted, etc.
# The default value is: NO.

OPTIMIZE_OUTPUT_FOR_C  = NO

# Set the OPTIMIZE_OUTPUT_JAVA tag to YES if your project consists of Java or
# Python sources only. Doxygen will then generate output that is more tailored
# for that language. For instance, namespaces will be presented as packages,
# qualified scopes will look different, etc.
# The default value is: NO.

OPTIMIZE_OUTPUT_JAVA   = NO

# Set the OPTIMIZE_FOR_FORTRAN tag to YES if your project consists of Fortran
# sources. Doxygen will then generate output that is tailored for Fortran.
# The default value is: NO.

OPTIMIZE_FOR_FORTRAN   = NO

# Set the OPTIMIZE_OUTPUT_VHDL tag to YES if your project consists of VHDL
# sources. Doxygen will then generate output that is tailored for VHDL.
# The default value is: NO.

OPTIMIZE_OUTPUT_VHDL   = NO

# Doxygen selects the parser to use depending on the extension of the files it
# parses. With this tag you can assign which parser to use for a given
# extension. Doxygen has a built-in mapping, but you can override or extend it
# using this tag. The format is ext=language, where ext is a file extension, and
# language is one of the parsers supported by doxygen: IDL, Java, Javascript,
# C#, C, C++, D, PHP, Objective-C, Python, Fortran (fixed format Fortran:
# FortranFixed, free formatted Fortran: FortranFree, unknown formatted Fortran:
# Fortran. In the later case the parser tries to guess whether the code is fixed
# or free formatted code, this is the default for Fortran type files), VHDL. For
# instance to make doxygen treat .inc files as Fortran files (default is PHP),
# and .f files as C (default is Fortran), use: inc=Fortran f=C.
#
# Note For files without extension you can use no_extension as a placeholder.
#
# Note that for custom extensions you also need to set FILE_PATTERNS otherwise
# the files are not read by doxygen.

EXTENSION_MAPPING      =

# If the MARKDOWN_SUPPORT tag is enabled then doxygen pre-processes all comments
# according to the Markdown format, which allows for more readable
# documentation. See http://daringfireball.net/projects/markdown/ for details.
# The output of markdown processing is further processed by doxygen, so you can
# mix doxygen, HTML, and XML commands with Markdown formatting. Disable only in
# case of backward compatibilities issues.
# The default value is: YES.

MARKDOWN_SUPPORT       = YES

# When enabled doxygen tries to link words that correspond to documented
# classes, or namespaces to their corresponding documentation. Such a link can
# be prevented in individual cases by by putting a % sign in front of the word
# or globally by setting AUTOLINK_SUPPORT to NO.
# The default value is: YES.

AUTOLINK_SUPPORT       = YES

# If you use STL classes (i.e. std::string, std::vector, etc.) but do not want
# to include (a tag file for) the STL sources as input, then you should set this
# tag to YES in order to let doxygen match functions declarations and
# definitions whose arguments contain STL classes (e.g. func(std::string);
# versus func(std::string) {}). This also make the inheritance and collaboration
# diagrams that involve STL classes more complete and accurate.
# The default value is: NO.

BUILTIN_STL_SUPPORT    = NO

# If you use Microsoft's C++/CLI language, you should set this option to YES to
# enable parsing support.
# The default value is: NO.

CPP_CLI_SUPPORT        = NO

# Set the SIP_SUPPORT tag to YES if your project consists of sip (see:
# http://www.riverbankcomputing.co.uk/software/sip/intro) sources only. Doxygen
# will parse them like normal C++ but will assume all classes use public instead
# of private inheritance when no explicit protection keyword is present.
# The default value is: NO.

SIP_SUPPORT            = NO

# For Microsoft's IDL there are propget and propput attributes to indicate
# getter and setter methods for a property. Setting this option to YES will make
# doxygen to replace the get and set methods by a property in the documentation.
# This will only work if the methods are indeed getting or setting a simple
# type. If this is not the case, or you want to show the methods anyway, you
# should set this option to NO.
# The default value is: YES.

IDL_PROPERTY_SUPPORT   = YES

# If member grouping is used in the documentation and the DISTRIBUTE_GROUP_DOC
# tag is set to YES, then doxygen will reuse the documentation of the first
# member in the group (if any) for the other members of the group. By default
# all members of a group must be documented explicitly.
# The default value is: NO.

DISTRIBUTE_GROUP_DOC   = NO

# Set the SUBGROUPING tag to YES to allow class member groups of the same type
# (for instance a group of public functions) to be put as a subgroup of that
# type (e.g. under the Public Functions section). Set it to NO to prevent
# subgrouping. Alternatively, this can be done per class using the
# \nosubgrouping command.
# The default value is: YES.

SUBGROUPING            = YES

# When the INLINE_GROUPED_CLASSES tag is set to YES, classes, structs and unions
# are shown inside the group in which they are included (e.g. using \ingroup)
# instead of on a separate page (for HTML and Man pages) or section (for LaTeX
# and RTF).
#
# Note that this feature does not work in combination with
# SEPARATE_MEMBER_PAGES.
# The default value is: NO.

INLINE_GROUPED_CLASSES = NO

# When the INLINE_SIMPLE_STRUCTS tag is set to YES, structs, classes, and unions
# with only public data fields or simple typedef fields will be shown inline in
# the documentation of the scope in which they are defined (i.e. file,
# namespace, or group documentation), provided this scope is documented. If set
# to NO, structs, classes, and unions are shown on a separate page (for HTML and
# Man pages) or section (for LaTeX and RTF).
# The default value is: NO.

INLINE_SIMPLE_STRUCTS  = NO

# When TYPEDEF_HIDES_STRUCT tag is enabled, a typedef of a struct, union, or
# enum is documented as struct, union, or enum with the name of the typedef. So
# typedef struct TypeS {} TypeT, will appear in the documentation as a struct
# with name TypeT. When disabled the typedef will appear as a member of a file,
# namespace, or class. And the struct will be named TypeS. This can typically be
# useful for C code in case the coding convention dictates that all compound
# types are typedef'ed and only the typedef is referenced, never the tag name.
# The default value is: NO.

TYPEDEF_HIDES_STRUCT   = NO

# The size of the symbol lookup cache can be set using LOOKUP_CACHE_SIZE. This
# cache is used to resolve symbols given their name and scope. Since this can be
# an expensive process and often the same symbol appears multiple times in the
# code, doxygen keeps a cache of pre-resolved symbols. If the cache is too small
# doxygen will become slower. If the cache is too large, memory is wasted. The
# cache size is given by this formula: 2^(16+LOOKUP_CACHE_SIZE). The valid range
# is 0..9, the default is 0, corresponding to a cache size of 2^16=65536
# symbols. At the end of a run doxygen will report the cache usage and suggest
# the optimal cache size from a speed point of view.
# Minimum value: 0, maximum value: 9, default value: 0.

LOOKUP_CACHE_SIZE      = 0

#---------------------------------------------------------------------------
# Build related configuration options
#---------------------------------------------------------------------------

# If the EXTRACT_ALL tag is set to YES doxygen will assume all entities in
# documentation are documented, even if no documentation was available. Private
# class members and static file members will be hidden unless the
# EXTRACT_PRIVATE respectively EXTRACT_STATIC tags are set to YES.
# Note: This will also disable the warnings about undocumented members that are
# normally produced when WARNINGS is set to YES.
# The default value is: NO.

EXTRACT_ALL            = NO

# If the EXTRACT_PRIVATE tag is set to YES all private members of a class will
# be included in the documentation.
# The default value is: NO.

EXTRACT_PRIVATE        = NO

# If the EXTRACT_PACKAGE tag is set to YES all members with package or internal
# scope will be included in the documentation.
# The default value is: NO.

EXTRACT_PACKAGE        = NO

# If the EXTRACT_STATIC tag is set to YES all static members of a file will be
# included in the documentation.
# The default value is: NO.

EXTRACT_STATIC         = YES

# If the EXTRACT_LOCAL_CLASSES tag is set to YES classes (and structs) defined
# locally in source files will be included in the documentation. If set to NO
# only classes defined in header files are included. Does not have any effect
# for Java sources.
# The default value is: YES.

EXTRACT_LOCAL_CLASSES  = YES

# This flag is only useful for Objective-C code. When set to YES local methods,
# which are defined in the implementation section but not in the interface are
# included in the documentation. If set to NO only methods in the interface are
# included.
# The default value is: NO.

EXTRACT_LOCAL_METHODS  = NO

# If this flag is set to YES, the members of anonymous namespaces will be
# extracted and appear in the documentation as a namespace called
# 'anonymous_namespace{file}', where file will be replaced with the base name of
# the file that contains the anonymous namespace. By default anonymous namespace
# are hidden.
# The default value is: NO.

EXTRACT_ANON_NSPACES   = NO

# If the HIDE_UNDOC_MEMBERS tag is set to YES, doxygen will hide all
# undocumented members inside documented classes or files. If set to NO these
# members will be included in the various overviews, but no documentation
# section is generated. This option has no effect if EXTRACT_ALL is enabled.
# The default value is: NO.

HIDE_UNDOC_MEMBERS     = NO

# If the HIDE_UNDOC_CLASSES tag is set to YES, doxygen will hide all
# undocumented classes that are normally visible in the class hierarchy. If set
# to NO these classes will be included in the various overviews. This option has
# no effect if EXTRACT_ALL is enabled.
# The default value is: NO.

HIDE_UNDOC_CLASSES     = NO

# If the HIDE_FRIEND_COMPOUNDS tag is set to YES, doxygen will hide all friend
# (class|struct|union) declarations. If set to NO these declarations will be
# included in the documentation.
# The default value is: NO.

HIDE_FRIEND_COMPOUNDS  = NO

# If the HIDE_IN_BODY_DOCS tag is set to YES, doxygen will hide any
# documentation blocks found inside the body of a function. If set to NO these
# blocks will be appended to the function's detailed documentation block.
# The default value is: NO.

HIDE_IN_BODY_DOCS      = NO

# The INTERNAL_DOCS tag determines if documentation that is typed after a
# \internal command is included. If the tag is set to NO then the documentation
# will be excluded. Set it to YES to include the internal documentation.
# The default value is: NO.

INTERNAL_DOCS          = NO

# If the CASE_SENSE_NAMES tag is set to NO then doxygen will only generate file
# names in lower-case letters. If set to YES upper-case letters are also
# allowed. This is useful if you have classes or files whose names only differ
# in case and if your file system supports case sensitive file names. Windows
# and Mac users are advised to set this option to NO.
# The default value is: system dependent.

CASE_SENSE_NAMES       = YES

# If the HIDE_SCOPE_NAMES tag is set to NO then doxygen will show members with
# their full class and namespace scopes in the documentation. If set to YES the
# scope will be hidden.
# The default value is: NO.

HIDE_SCOPE_NAMES       = NO

# If the SHOW_INCLUDE_FILES tag is set to YES then doxygen will put a list of
# the files that are included by a file in the documentation of that file.
# The default value is: YES.

SHOW_INCLUDE_FILES     = YES

# If the SHOW_GROUPED_MEMB_INC tag is set to YES then Doxygen will add for each
# grouped member an include statement to the documentation, telling the reader
# which file to include in order to use the member.
# The default value is: NO.

SHOW_GROUPED_MEMB_INC  = NO

# If the FORCE_LOCAL_INCLUDES tag is set to YES then doxygen will list include
# files with double quotes in the documentation rather than with sharp brackets.
# The default value is: NO.

FORCE_LOCAL_INCLUDES   = NO

# If the INLINE_INFO tag is set to YES then a tag [inline] is inserted in the
# documentation for inline members.
# The default value is: YES.

INLINE_INFO            = YES

# If the SORT_MEMBER_DOCS tag is set to YES then doxygen will sort the
# (detailed) documentation of file and class members alphabetically by member
# name. If set to NO the members will appear in declaration order.
# The default value is: YES.

SORT_MEMBER_DOCS       = YES

# If the SORT_BRIEF_DOCS tag is set to YES then doxygen will sort the brief
# descriptions of file, namespace and class members alphabetically by member
# name. If set to NO the members will appear in declaration order. Note that
# this will also influence the order of the classes in the class list.
# The default value is: NO.

SORT_BRIEF_DOCS        = NO

# If the SORT_MEMBERS_CTORS_1ST tag is set to YES then doxygen will sort the
# (brief and detailed) documentation of class members so that constructors and
# destructors are listed first. If set to NO the constructors will appear in the
# respective orders defined by SORT_BRIEF_DOCS and SORT_MEMBER_DOCS.
# Note: If SORT_BRIEF_DOCS is set to NO this option is ignored for sorting brief
# member documentation.
# Note: If SORT_MEMBER_DOCS is set to NO this option is ignored for sorting
# detailed member documentation.
# The default value is: NO.

SORT_MEMBERS_CTORS_1ST = NO

# If the SORT_GROUP_NAMES tag is set to YES then doxygen will sort the hierarchy
# of group names into alphabetical order. If set to NO the group names will
# appear in their defined order.
# The default value is: NO.

SORT_GROUP_NAMES       = NO

# If the SORT_BY_SCOPE_NAME tag is set to YES, the class list will be sorted by
# fully-qualified names, including namespaces. If set to NO, the class list will
# be sorted only by class name, not including the namespace part.
# Note: This option is not very useful if HIDE_SCOPE_NAMES is set to YES.
# Note: This option applies only to the class list, not to the alphabetical
# list.
# The default value is: NO.

SORT_BY_SCOPE_NAME     = NO

# If the STRICT_PROTO_MATCHING option is enabled and doxygen fails to do proper
# type resolution of all parameters of a function it will reject a match between
# the prototype and the implementation of a member function even if there is
# only one candidate or it is obvious which candidate to choose by doing a
# simple string match. By disabling STRICT_PROTO_MATCHING doxygen will still
# accept a match between prototype and implementation in such cases.
# The default value is: NO.

STRICT_PROTO_MATCHING  = NO

# The GENERATE_TODOLIST tag can be used to enable ( YES) or disable ( NO) the
# todo list. This list is created by putting \todo commands in the
# documentation.
# The default value is: YES.

GENERATE_TODOLIST      = YES

# The GENERATE_TESTLIST tag can be used to enable ( YES) or disable ( NO) the
# test list. This list is created by putting \test commands in the
# documentation.
# The default value is: YES.

GENERATE_TESTLIST      = YES

# The GENERATE_BUGLIST tag can be used to enable ( YES) or disable ( NO) the bug
# list. This list is created by putting \bug commands in the documentation.
# The default value is: YES.

GENERATE_BUGLIST       = YES

# The GENERATE_DEPRECATEDLIST tag can be used to enable ( YES) or disable ( NO)
# the deprecated list. This list is created by putting \deprecated commands in
# the documentation.
# The default value is: YES.

GENERATE_DEPRECATEDLIST= YES

# The ENABLED_SECTIONS tag can be used to enable conditional documentation
# sections, marked by \if <section_label> ... \endif and \cond <section_label>
# ... \endcond blocks.

ENABLED_SECTIONS       =

# The MAX_INITIALIZER_LINES tag determines the maximum number of lines that the
# initial value of a variable or macro / define can have for it to appear in the
# documentation. If the initializer consists of more lines than specified here
# it will be hidden. Use a value of 0 to hide initializers completely. The
# appearance of the value of individual variables and macros / defines can be
# controlled using \showinitializer or \hideinitializer command in the
# documentation regardless of this setting.
# Minimum value: 0, maximum value: 10000, default value: 30.

MAX_INITIALIZER_LINES  = 30

# Set the SHOW_USED_FILES tag to NO to disable the list of files generated at
# the bottom of the documentation of classes and structs. If set to YES the list
# will mention the files that were used to generate the documentation.
# The default value is: YES.

SHOW_USED_FILES        = YES

# Set the SHOW_FILES tag to NO to disable the generation of the Files page. This
# will remove the Files entry from the Quick Index and from the Folder Tree View
# (if specified).
# The default value is: YES.

SHOW_FILES             = YES

# Set the SHOW_NAMESPACES tag to NO to disable the generation of the Namespaces
# page. This will remove the Namespaces entry from the Quick Index and from the
# Folder Tree View (if specified).
# The default value is: YES.

SHOW_NAMESPACES        = YES

# The FILE_VERSION_FILTER tag can be used to specify a program or script that
# doxygen should invoke to get the current version for each file (typically from
# the version control system). Doxygen will invoke the program by executing (via
# popen()) the command command input-file, where command is the value of the
# FILE_VERSION_FILTER tag, and input-file is the name of an input file provided
# by doxygen. Whatever the program writes to standard output is used as the file
# version. For an example see the documentation.

FILE_VERSION_FILTER    =

# The LAYOUT_FILE tag can be used to specify a layout file which will be parsed
# by doxygen. The layout file controls the global structure of the generated
# output files in an output format independent way. To create the layout file
# that represents doxygen's defaults, run doxygen with the -l option. You can
# optionally specify a file name after the option, if omitted DoxygenLayout.xml
# will be used as the name of the layout file.
#
# Note that if you run doxygen from a directory containing a file called
# DoxygenLayout.xml, doxygen will parse it automatically even if the LAYOUT_FILE
# tag is left empty.

LAYOUT_FILE            =

# The CITE_BIB_FILES tag can be used to specify one or more bib files containing
# the reference definitions. This must be a list of .bib files. The .bib
# extension is automatically appended if omitted. This requires the bibtex tool
# to be installed. See also http://en.wikipedia.org/wiki/BibTeX for more info.
# For LaTeX the style of the bibliography can be controlled using
# LATEX_BIB_STYLE. To use this feature you need bibtex and perl available in the
# search path. See also \cite for info how to create references.

CITE_BIB_FILES         =

#---------------------------------------------------------------------------
# Configuration options related to warning and progress messages
#---------------------------------------------------------------------------

# The QUIET tag can be used to turn on/off the messages that are generated to
# standard output by doxygen. If QUIET is set to YES this implies that the
# messages are off.
# The default value is: NO.

QUIET                  = NO

# The WARNINGS tag can be used to turn on/off the warning messages that are
# generated to standard error ( stderr) by doxygen. If WARNINGS is set to YES
# this implies that the warnings are on.
#
# Tip: Turn warnings on while writing the documentation.
# The default value is: YES.

WARNINGS               = YES

# If the WARN_IF_UNDOCUMENTED tag is set to YES, then doxygen will generate
# warnings for undocumented members. If EXTRACT_ALL is set to YES then this flag
# will automatically be disabled.
# The default value is: YES.

WARN_IF_UNDOCUMENTED   = YES

# If the WARN_IF_DOC_ERROR tag is set to YES, doxygen will generate warnings for
# potential errors in the documentation, such as not documenting some parameters
# in a documented function, or documenting parameters that don't exist or using
# markup commands wrongly.
# The default value is: YES.

WARN_IF_DOC_ERROR      = YES

# This WARN_NO_PARAMDOC option can be enabled to get warnings for functions that
# are documented, but have no documentation for their parameters or return
# value. If set to NO doxygen will only warn about wrong or incomplete parameter
# documentation, but not about the absence of documentation.
# The default value is: NO.

WARN_NO_PARAMDOC       = NO

# The WARN_FORMAT tag determines the format of the warning messages that doxygen
# can produce. The string should contain the $file, $line, and $text tags, which
# will be replaced by the file and line number from which the warning originated
# and the warning text. Optionally the format may contain $version, which will
# be replaced by the version of the file (if it could be obtained via
# FILE_VERSION_FILTER)
# The default value is: $file:$line: $text.

WARN_FORMAT            = "$file:$line: $text"

# The WARN_LOGFILE tag can be used to specify a file to which warning and error
# messages should be written. If left blank the output is written to standard
# error (stderr).

WARN_LOGFILE           =

#---------------------------------------------------------------------------
# Configuration options related to the input files
#---------------------------------------------------------------------------

# The INPUT tag is used to specify the files and/or directories that contain
# documented source files. You may enter file names like myfile.cpp or
# directories like /usr/src/myproject. Separate the files or directories with
# spaces.
# Note: If this tag is empty the current directory is searched.

INPUT                  = ./src ./include

# This tag can be used to specify the character encoding of the source files
# that doxygen parses. Internally doxygen uses the UTF-8 encoding. Doxygen uses
# libiconv (or the iconv built into libc) for the transcoding. See the libiconv
# documentation (see: http://www.gnu.org/software/libiconv) for the list of
# possible encodings.
# The default value is: UTF-8.

INPUT_ENCODING         = UTF-8

# If the value of the INPUT tag contains directories, you can use the
# FILE_PATTERNS tag to specify one or more wildcard patterns (like *.cpp and
# *.h) to filter out the source-files in the directories. If left blank the
# following patterns are tested:*.c, *.cc, *.cxx, *.cpp, *.c++, *.java, *.ii,
# *.ixx, *.ipp, *.i++, *.inl, *.idl, *.ddl, *.odl, *.h, *.hh, *.hxx, *.hpp,
# *.h++, *.cs, *.d, *.php, *.php4, *.php5, *.phtml, *.inc, *.m, *.markdown,
# *.md, *.mm, *.dox, *.py, *.f90, *.f, *.for, *.tcl, *.vhd, *.vhdl, *.ucf,
# *.qsf, *.as and *.js.

FILE_PATTERNS          =

# The RECURSIVE tag can be used to specify whether or not subdirectories should
# be searched for input files as well.
# The default value is: NO.

RECURSIVE              = NO

# The EXCLUDE tag can be used to specify files and/or directories that should be
# excluded from the INPUT source files. This way you can easily exclude a
# subdirectory from a directory tree whose root is specified with the INPUT tag.
#
# Note that relative paths are relative to the directory from which doxygen is
# run.

EXCLUDE                =

# The EXCLUDE_SYMLINKS tag can be used to select whether or not files or
# directories that are symbolic links (a Unix file system feature) are excluded
# from the input.
# The default value is: NO.

EXCLUDE_SYMLINKS       = NO

# If the value of the INPUT tag contains directories, you can use the
# EXCLUDE_PATTERNS tag to specify one or more wildcard patterns to exclude
# certain files from those directories.
#
# Note that the wildcards are matched against the file with absolute path, so to
# exclude all test directories for example use the pattern */test/*

EXCLUDE_PATTERNS       =

# The EXCLUDE_SYMBOLS tag can be used to specify one or more symbol names
# (namespaces, classes, functions, etc.) that should be excluded from the
# output. The symbol name can be a fully qualified name, a word, or if the
# wildcard * is used, a substring. Examples: ANamespace, AClass,
# AClass::ANamespace, ANamespace::*Test
#
# Note that the wildcards are matched against the file with absolute path, so to
# exclude all test directories use the pattern */test/*

EXCLUDE_SYMBOLS        =

# The EXAMPLE_PATH tag can be used to specify one or more files or directories
# that contain example code fragments that are included (see the \include
# command).

EXAMPLE_PATH           =

# If the value of the EXAMPLE_PATH tag contains directories, you can use the
# EXAMPLE_PATTERNS tag to specify one or more wildcard pattern (like *.cpp and
# *.h) to filter out the source-files in the directories. If left blank all
# files are included.

EXAMPLE_PATTERNS       =

# If the EXAMPLE_RECURSIVE tag is set to YES then subdirectories will be
# searched for input files to be used with the \include or \dontinclude commands
# irrespective of the value of the RECURSIVE tag.
# The default value is: NO.

EXAMPLE_RECURSIVE      = NO

# The IMAGE_PATH tag can be used to specify one or more files or directories
# that contain images that are to be included in the documentation (see the
# \image command).

IMAGE_PATH             =

# The INPUT_FILTER tag can be used to specify a program that doxygen should
# invoke to filter for each input file. Doxygen will invoke the filter program
# by executing (via popen()) the command:
#
# <filter> <input-file>
#
# where <filter> is the value of the INPUT_FILTER tag, and <input-file> is the
# name of an input file. Doxygen will then use the output that the filter
# program writes to standard output. If FILTER_PATTERNS is specified, this tag
# will be ignored.
#
# Note that the filter must not add or remove lines; it is applied before the
# code is scanned, but not when the output code is generated. If lines are added
# or removed, the anchors will not be placed correctly.

INPUT_FILTER           =

# The FILTER_PATTERNS tag can be used to specify filters on a per file pattern
# basis. Doxygen will compare the file name with each pattern and apply the
# filter if there is a match. The filters are a list of the form: pattern=filter
# (like *.cpp=my_cpp_filter). See INPUT_FILTER for further information on how
# filters are used. If the FILTER_PATTERNS tag is empty or if none of the
# patterns match the file name, INPUT_FILTER is applied.

FILTER_PATTERNS        =

# If the FILTER_SOURCE_FILES tag is set to YES, the input filter (if set using
# INPUT_FILTER ) will also be used to filter the input files that are used for
# producing the source files to browse (i.e. when SOURCE_BROWSER is set to YES).
# The default value is: NO.

FILTER_SOURCE_FILES    = NO

# The FILTER_SOURCE_PATTERNS tag can be used to specify source filters per file
# pattern. A pattern will override the setting for FILTER_PATTERN (if any) and
# it is also possible to disable source filtering for a specific pattern using
# *.ext= (so without naming a filter).
# This tag requires that the tag FILTER_SOURCE_FILES is set to YES.

FILTER_SOURCE_PATTERNS =

# If the USE_MDFILE_AS_MAINPAGE tag refers to the name of a markdown file that
# is part of the input, its contents will be placed on the main page
# (index.html). This can be useful if you have a project on for instance GitHub
# and want to reuse the introduction page also for the doxygen output.

USE_MDFILE_AS_MAINPAGE =

#---------------------------------------------------------------------------
# Configuration options related to source browsing
#---------------------------------------------------------------------------

# If the SOURCE_BROWSER tag is set to YES then a list of source files will be
# generated. Documented entities will be cross-referenced with these sources.
#
# Note: To get rid of all source code in the generated output, make sure that
# also VERBATIM_HEADERS is set to NO.
# The default value is: NO.

SOURCE_BROWSER         = NO

# Setting the INLINE_SOURCES tag to YES will include the body of functions,
# classes and enums directly into the documentation.
# The default value is: NO.

INLINE_SOURCES         = NO

# Setting the STRIP_CODE_COMMENTS tag to YES will instruct doxygen to hide any
# special comment blocks from generated source code fragments. Normal C, C++ and
# Fortran comments will always remain visible.
# The default value is: YES.

STRIP_CODE_COMMENTS    = YES

# If the REFERENCED_BY_RELATION tag is set to YES then for each documented
# function all documented functions referencing it will be listed.
# The default value is: NO.

REFERENCED_BY_RELATION = NO

# If the REFERENCES_RELATION tag is set to YES then for each documented function
# all documented entities called/used by that function will be listed.
# The default value is: NO.

REFERENCES_RELATION    = NO

# If the REFERENCES_LINK_SOURCE tag is set to YES and SOURCE_BROWSER tag is set
# to YES, then the hyperlinks from functions in REFERENCES_RELATION and
# REFERENCED_BY_RELATION lists will link to the source code. Otherwise they will
# link to the documentation.
# The default value is: YES.

REFERENCES_LINK_SOURCE = YES

# If SOURCE_TOOLTIPS is enabled (the default) then hovering a hyperlink in the
# source code will show a tooltip with additional information such as prototype,
# brief description and links to the definition and documentation. Since this
# will make the HTML file larger and loading of large files a bit slower, you
# can opt to disable this feature.
# The default value is: YES.
# This tag requires that the tag SOURCE_BROWSER is set to YES.

SOURCE_TOOLTIPS        = YES

# If the USE_HTAGS tag is set to YES then the references to source code will
# point to the HTML generated by the htags(1) tool instead of doxygen built-in
# source browser. The htags tool is part of GNU's global source tagging system
# (see http://www.gnu.org/software/global/global.html). You will need version
# 4.8.6 or higher.
#
# To use it do the following:
# - Install the latest version of global
# - Enable SOURCE_BROWSER and USE_HTAGS in the config file
# - Make sure the INPUT points to the root of the source tree
# - Run doxygen as normal
#
# Doxygen will invoke htags (and that will in turn invoke gtags), so these
# tools must be available from the command line (i.e. in the search path).
#
# The result: instead of the source browser generated by doxygen, the links to
# source code will now point to the output of htags.
# The default value is: NO.
# This tag requires that the tag SOURCE_BROWSER is set to YES.

USE_HTAGS              = NO

# If the VERBATIM_HEADERS tag is set the YES then doxygen will generate a
# verbatim copy of the header file for each class for which an include is
# specified. Set to NO to disable this.
# See also: Section \class.
# The default value is: YES.

VERBATIM_HEADERS       = YES

# If the CLANG_ASSISTED_PARSING tag is set to YES, then doxygen will use the
# clang parser (see: http://clang.llvm.org/) for more accurate parsing at the
# cost of reduced performance. This can be particularly helpful with template
# rich C++ code for which doxygen's built-in parser lacks the necessary type
# information.
# Note: The availability of this option depends on whether or not doxygen was
# compiled with the --with-libclang option.
# The default value is: NO.

CLANG_ASSISTED_PARSING = NO

# If clang assisted parsing is enabled you can provide the compiler with command
# line options that you would normally use when invoking the compiler. Note that
# the include paths will already be set by doxygen for the files and directories
# specified with INPUT and INCLUDE_PATH.
# This tag requires that the tag CLANG_ASSISTED_PARSING is set to YES.

CLANG_OPTIONS          =

#---------------------------------------------------------------------------
# Configuration options related to the alphabetical class index
#---------------------------------------------------------------------------

# If the ALPHABETICAL_INDEX tag is set to YES, an alphabetical index of all
# compounds will be generated. Enable this if the project contains a lot of
# classes, structs, unions or interfaces.
# The default value is: YES.

ALPHABETICAL_INDEX     = YES

# The COLS_IN_ALPHA_INDEX tag can be used to specify the number of columns in
# which the alphabetical index list will be split.
# Minimum value: 1, maximum value: 20, default value: 5.
# This tag requires that the tag ALPHABETICAL_INDEX is set to YES.

COLS_IN_ALPHA_INDEX    = 5

# In case all classes in a project start with a common prefix, all classes will
# be put under the same header in the alphabetical index. The IGNORE_PREFIX tag
# can be used to specify a prefix (or a list of prefixes) that should be ignored
# while generating the index headers.
# This tag requires that the tag ALPHABETICAL_INDEX is set to YES.

IGNORE_PREFIX          =

#---------------------------------------------------------------------------
# Configuration options related to the HTML output
#---------------------------------------------------------------------------

# If the GENERATE_HTML tag is set to YES doxygen will generate HTML output
# The default value is: YES.

GENERATE_HTML          = YES

# The HTML_OUTPUT tag is used to specify where the HTML docs will be put. If a
# relative path is entered the value of OUTPUT_DIRECTORY will be put in front of
# it.
# The default directory is: html.
# This tag requires that the tag GENERATE_HTML is set to YES.

HTML_OUTPUT            = html

# The HTML_FILE_EXTENSION tag can be used to specify the file extension for each
# generated HTML page (for example: .htm, .php, .asp).
# The default value is: .html.
# This tag requires that the tag GENERATE_HTML is set to YES.

HTML_FILE_EXTENSION    = .html

# The HTML_HEADER tag can be used to specify a user-defined HTML header file for
# each generated HTML page. If the tag is left blank doxygen will generate a
# standard header.
#
# To get valid HTML the header file that includes any scripts and style sheets
# that doxygen needs, which is dependent on the configuration options used (e.g.
# the setting GENERATE_TREEVIEW). It is highly recommended to start with a
# default header using
# doxygen -w html new_header.html new_footer.html new_stylesheet.css
# YourConfigFile
# and then modify the file new_header.html. See also section "Doxygen usage"
# for information on how to generate the default header that doxygen normally
# uses.
# Note: The header is subject to change so you typically have to regenerate the
# default header when upgrading to a newer version of doxygen. For a description
# of the possible markers and block names see the documentation.
# This tag requires that the tag GENERATE_HTML is set to YES.

HTML_HEADER            =

# The HTML_FOOTER tag can be used to specify a user-defined HTML footer for each
# generated HTML page. If the tag is left blank doxygen will generate a standard
# footer. See HTML_HEADER for more information on how to generate a default
# footer and what special commands can be used inside the footer. See also
# section "Doxygen usage" for information on how to generate the default footer
# that doxygen normally uses.
# This tag requires that the tag GENERATE_HTML is set to YES.

HTML_FOOTER            =

# The HTML_STYLESHEET tag can be used to specify a user-defined cascading style
# sheet that is used by each HTML page. It can be used to fine-tune the look of
# the HTML output. If left blank doxygen will generate a default style sheet.
# See also section "Doxygen usage" for information on how to generate the style
# sheet that doxygen normally uses.
# Note: It is recommended to use HTML_EXTRA_STYLESHEET instead of this tag, as
# it is more robust and this tag (HTML_STYLESHEET) will in the future become
# obsolete.
# This tag requires that the tag GENERATE_HTML is set to YES.

HTML_STYLESHEET        =

# The HTML_EXTRA_STYLESHEET tag can be used to specify additional user-defined
# cascading style sheets that are included after the standard style sheets
# created by doxygen. Using this option one can overrule certain style aspects.
# This is preferred over using HTML_STYLESHEET since it does not replace the
# standard style sheet and is therefor more robust against future updates.
# Doxygen will copy the style sheet files to the output directory.
# Note: The order of the extra stylesheet files is of importance (e.g. the last
# stylesheet in the list overrules the setting of the previous ones in the
# list). For an example see the documentation.
# This tag requires that the tag GENERATE_HTML is set to YES.

HTML_EXTRA_STYLESHEET  =

# The HTML_EXTRA_FILES tag can be used to specify one or more extra images or
# other source files which should be copied to the HTML output directory. Note
# that these files will be copied to the base HTML output directory. Use the
# $relpath^ marker in the HTML_HEADER and/or HTML_FOOTER files to load these
# files. In the HTML_STYLESHEET file, use the file name only. Also note that the
# files will be copied as-is; there are no commands or markers available.
# This tag requires that the tag GENERATE_HTML is set to YES.

HTML_EXTRA_FILES       =

# The HTML_COLORSTYLE_HUE tag controls the color of the HTML output. Doxygen
# will adjust the colors in the stylesheet and background images according to
# this color. Hue is specified as an angle on a colorwheel, see
# http://en.wikipedia.org/wiki/Hue for more information. For instance the value
# 0 represents red, 60 is yellow, 120 is green, 180 is cyan, 240 is blue, 300
# purple, and 360 is red again.
# Minimum value: 0, maximum value: 359, default value: 220.
# This tag requires that the tag GENERATE_HTML is set to YES.

HTML_COLORSTYLE_HUE    = 220

# The HTML_COLORSTYLE_SAT tag controls the purity (or saturation) of the colors
# in the HTML output. For a value of 0 the output will use grayscales only. A
# value of 255 will produce the most vivid colors.
# Minimum value: 0, maximum value: 255, default value: 100.
# This tag requires that the tag GENERATE_HTML is set to YES.

HTML_COLORSTYLE_SAT    = 100

# The HTML_COLORSTYLE_GAMMA tag controls the gamma correction applied to the
# luminance component of the colors in the HTML output. Values below 100
# gradually make the output lighter, whereas values above 100 make the output
# darker. The value divided by 100 is the actual gamma applied, so 80 represents
# a gamma of 0.8, The value 220 represents a gamma of 2.2, and 100 does not
# change the gamma.
# Minimum value: 40, maximum value: 240, default value: 80.
# This tag requires that the tag GENERATE_HTML is set to YES.

HTML_COLORSTYLE_GAMMA  = 80

# If the HTML_TIMESTAMP tag is set to YES then the footer of each generated HTML
# page will contain the date and time when the page was generated. Setting this
# to NO can help when comparing the output of multiple runs.
# The default value is: YES.
# This tag requires that the tag GENERATE_HTML is set to YES.

HTML_TIMESTAMP         = YES

# If the HTML_DYNAMIC_SECTIONS tag is set to YES then the generated HTML
# documentation will contain sections that can be hidden and shown after the
# page has loaded.
# The default value is: NO.
# This tag requires that the tag GENERATE_HTML is set to YES.

HTML_DYNAMIC_SECTIONS  = NO

# With HTML_INDEX_NUM_ENTRIES one can control the preferred number of entries
# shown in the various tree structured indices initially; the user can expand
# and collapse entries dynamically later on. Doxygen will expand the tree to
# such a level that at most the specified number of entries are visible (unless
# a fully collapsed tree already exceeds this amount). So setting the number of
# entries 1 will produce a full collapsed tree by default. 0 is a special value
# representing an infinite number of entries and will result in a full expanded
# tree by default.
# Minimum value: 0, maximum value: 9999, default value: 100.
# This tag requires that the tag GENERATE_HTML is set to YES.

HTML_INDEX_NUM_ENTRIES = 100

# If the GENERATE_DOCSET tag is set to YES, additional index files will be
# generated that can be used as input for Apple's Xcode 3 integrated development
# environment (see: http://developer.apple.com/tools/xcode/), introduced with
# OSX 10.5 (Leopard). To create a documentation set, doxygen will generate a
# Makefile in the HTML output directory. Running make will produce the docset in
# that directory and running make install will install the docset in
# ~/Library/Developer/Shared/Documentation/DocSets so that Xcode will find it at
# startup. See http://developer.apple.com/tools/creatingdocsetswithdoxygen.html
# for more information.
# The default value is: NO.
# This tag requires that the tag GENERATE_HTML is set to YES.

GENERATE_DOCSET        = NO

# This tag determines the name of the docset feed. A documentation feed provides
# an umbrella under which multiple documentation sets from a single provider
# (such as a company or product suite) can be grouped.
# The default value is: Doxygen generated docs.
# This tag requires that the tag GENERATE_DOCSET is set to YES.

DOCSET_FEEDNAME        = "Doxygen generated docs"

# This tag specifies a string that should uniquely identify the documentation
# set bundle. This should be a reverse domain-name style string, e.g.
# com.mycompany.MyDocSet. Doxygen will append .docset to the name.
# The default value is: org.doxygen.Project.
# This tag requires that the tag GENERATE_DOCSET is set to YES.

DOCSET_BUNDLE_ID       = org.doxygen.Project

# The DOCSET_PUBLISHER_ID tag specifies a string that should uniquely identify
# the documentation publisher. This should be a reverse domain-name style
# string, e.g. com.mycompany.MyDocSet.documentation.
# The default value is: org.doxygen.Publisher.
# This tag requires that the tag GENERATE_DOCSET is set to YES.

DOCSET_PUBLISHER_ID    = org.doxygen.Publisher

# The DOCSET_PUBLISHER_NAME tag identifies the documentation publisher.
# The default value is: Publisher.
# This tag requires that the tag GENERATE_DOCSET is set to YES.

DOCSET_PUBLISHER_NAME  = Publisher

# If the GENERATE_HTMLHELP tag is set to YES then doxygen generates three
# additional HTML index files: index.hhp, index.hhc, and index.hhk. The
# index.hhp is a project file that can be read by Microsoft's HTML Help Workshop
# (see: http://www.microsoft.com/en-us/download/details.aspx?id=21138) on
# Windows.
#
# The HTML Help Workshop contains a compiler that can convert all HTML output
# generated by doxygen into a single compiled HTML file (.chm). Compiled HTML
# files are now used as the Windows 98 help format, and will replace the old
# Windows help format (.hlp) on all Windows platforms in the future. Compressed
# HTML files also contain an index, a table of contents, and you can search for
# words in the documentation. The HTML workshop also contains a viewer for
# compressed HTML files.
# The default value is: NO.
# This tag requires that the tag GENERATE_HTML is set to YES.

GENERATE_HTMLHELP      = NO

# The CHM_FILE tag can be used to specify the file name of the resulting .chm
# file. You can add a path in front of the file if the result should not be
# written to the html output directory.
# This tag requires that the tag GENERATE_HTMLHELP is set to YES.

CHM_FILE               =

# The HHC_LOCATION tag can be used to specify the location (absolute path
# including file name) of the HTML help compiler ( hhc.exe). If non-empty
# doxygen will try to run the HTML help compiler on the generated index.hhp.
# The file has to be specified with full path.
# This tag requires that the tag GENERATE_HTMLHELP is set to YES.

HHC_LOCATION           =

# The GENERATE_CHI flag controls if a separate .chi index file is generated (
# YES) or that it should be included in the master .chm file ( NO).
# The default value is: NO.
# This tag requires that the tag GENERATE_HTMLHELP is set to YES.

GENERATE_CHI           = NO

# The CHM_INDEX_ENCODING is used to encode HtmlHelp index ( hhk), content ( hhc)
# and project file content.
# This tag requires that the tag GENERATE_HTMLHELP is set to YES.

CHM_INDEX_ENCODING     =

# The BINARY_TOC flag controls whether a binary table of contents is generated (
# YES) or a normal table of contents ( NO) in the .chm file. Furthermore it
# enables the Previous and Next buttons.
# The default value is: NO.
# This tag requires that the tag GENERATE_HTMLHELP is set to YES.

BINARY_TOC             = NO

# The TOC_EXPAND flag can be set to YES to add extra items for group members to
# the table of contents of the HTML help documentation and to the tree view.
# The default value is: NO.
# This tag requires that the tag GENERATE_HTMLHELP is set to YES.

TOC_EXPAND             = NO

# If the GENERATE_QHP tag is set to YES and both QHP_NAMESPACE and
# QHP_VIRTUAL_FOLDER are set, an additional index file will be generated that
# can be used as input for Qt's qhelpgenerator to generate a Qt Compressed Help
# (.qch) of the generated HTML documentation.
# The default value is: NO.
# This tag requires that the tag GENERATE_HTML is set to YES.

GENERATE_QHP           = NO

# If the QHG_LOCATION tag is specified, the QCH_FILE tag can be used to specify
# the file name of the resulting .qch file. The path specified is relative to
# the HTML output folder.
# This tag requires that the tag GENERATE_QHP is set to YES.

QCH_FILE               =

# The QHP_NAMESPACE tag specifies the namespace to use when generating Qt Help
# Project output. For more information please see Qt Help Project / Namespace
# (see: http://qt-project.org/doc/qt-4.8/qthelpproject.html#namespace).
# The default value is: org.doxygen.Project.
# This tag requires that the tag GENERATE_QHP is set to YES.

QHP_NAMESPACE          = org.doxygen.Project

# The QHP_VIRTUAL_FOLDER tag specifies the namespace to use when generating Qt
# Help Project output. For more information please see Qt Help Project / Virtual
# Folders (see: http://qt-project.org/doc/qt-4.8/qthelpproject.html#virtual-
# folders).
# The default value is: doc.
# This tag requires that the tag GENERATE_QHP is set to YES.

QHP_VIRTUAL_FOLDER     = doc

# If the QHP_CUST_FILTER_NAME tag is set, it specifies the name of a custom
# filter to add. For more information please see Qt Help Project / Custom
# Filters (see: http://qt-project.org/doc/qt-4.8/qthelpproject.html#custom-
# filters).
# This tag requires that the tag GENERATE_QHP is set to YES.

QHP_CUST_FILTER_NAME   =

# The QHP_CUST_FILTER_ATTRS tag specifies the list of the attributes of the
# custom filter to add. For more information please see Qt Help Project / Custom
# Filters (see: http://qt-project.org/doc/qt-4.8/qthelpproject.html#custom-
# filters).
# This tag requires that the tag GENERATE_QHP is set to YES.

QHP_CUST_FILTER_ATTRS  =

# The QHP_SECT_FILTER_ATTRS tag specifies the list of the attributes this
# project's filter section matches. Qt Help Project / Filter Attributes (see:
# http://qt-project.org/doc/qt-4.8/qthelpproject.html#filter-attributes).
# This tag requires that the tag GENERATE_QHP is set to YES.

QHP_SECT_FILTER_ATTRS  =

# The QHG_LOCATION tag can be used to specify the location of Qt's
# qhelpgenerator. If non-empty doxygen will try to run qhelpgenerator on the
# generated .qhp file.
# This tag requires that the tag GENERATE_QHP is set to YES.

QHG_LOCATION           =

# If the GENERATE_ECLIPSEHELP tag is set to YES, additional index files will be
# generated, together with the HTML files, they form an Eclipse help plugin. To
# install this plugin and make it available under the help contents menu in
# Eclipse, the contents of the directory containing the HTML and XML files needs
# to be copied into the plugins directory of eclipse. The name of the directory
# within the plugins directory should be the same as the ECLIPSE_DOC_ID value.
# After copying Eclipse needs to be restarted before the help appears.
# The default value is: NO.
# This tag requires that the tag GENERATE_HTML is set to YES.

GENERATE_ECLIPSEHELP   = NO

# A unique identifier for the Eclipse help plugin. When installing the plugin
# the directory name containing the HTML and XML files should also have this
# name. Each documentation set should have its own identifier.
# The default value is: org.doxygen.Project.
# This tag requires that the tag GENERATE_ECLIPSEHELP is set to YES.

ECLIPSE_DOC_ID         = org.doxygen.Project

# If you want full control over the layout of the generated HTML pages it might
# be necessary to disable the index and replace it with your own. The
# DISABLE_INDEX tag can be used to turn on/off the condensed index (tabs) at top
# of each HTML page. A value of NO enables the index and the value YES disables
# it. Since the tabs in the index contain the same information as the navigation
# tree, you can set this option to YES if you also set GENERATE_TREEVIEW to YES.
# The default value is: NO.
# This tag requires that the tag GENERATE_HTML is set to YES.

DISABLE_INDEX          = NO

# The GENERATE_TREEVIEW tag is used to specify whether a tree-like index
# structure should be generated to display hierarchical information. If the tag
# value is set to YES, a side panel will be generated containing a tree-like
# index structure (just like the one that is generated for HTML Help). For this
# to work a browser that supports JavaScript, DHTML, CSS and frames is required
# (i.e. any modern browser). Windows users are probably better off using the
# HTML help feature. Via custom stylesheets (see HTML_EXTRA_STYLESHEET) one can
# further fine-tune the look of the index. As an example, the default style
# sheet generated by doxygen has an example that shows how to put an image at
# the root of the tree instead of the PROJECT_NAME. Since the tree basically has
# the same information as the tab index, you could consider setting
# DISABLE_INDEX to YES when enabling this option.
# The default value is: NO.
# This tag requires that the tag GENERATE_HTML is set to YES.

GENERATE_TREEVIEW      = NO

# The ENUM_VALUES_PER_LINE tag can be used to set the number of enum values that
# doxygen will group on one line in the generated HTML documentation.
#
# Note that a value of 0 will completely suppress the enum values from appearing
# in the overview section.
# Minimum value: 0, maximum value: 20, default value: 4.
# This tag requires that the tag GENERATE_HTML is set to YES.

ENUM_VALUES_PER_LINE   = 4

# If the treeview is enabled (see GENERATE_TREEVIEW) then this tag can be used
# to set the initial width (in pixels) of the frame in which the tree is shown.
# Minimum value: 0, maximum value: 1500, default value: 250.
# This tag requires that the tag GENERATE_HTML is set to YES.

TREEVIEW_WIDTH         = 250

# When the EXT_LINKS_IN_WINDOW option is set to YES doxygen will open links to
# external symbols imported via tag files in a separate window.
# The default value is: NO.
# This tag requires that the tag GENERATE_HTML is set to YES.

EXT_LINKS_IN_WINDOW    = NO

# Use this tag to change the font size of LaTeX formulas included as images in
# the HTML documentation. When you change the font size after a successful
# doxygen run you need to manually remove any form_*.png images from the HTML
# output directory to force them to be regenerated.
# Minimum value: 8, maximum value: 50, default value: 10.
# This tag requires that the tag GENERATE_HTML is set to YES.

FORMULA_FONTSIZE       = 10

# Use the FORMULA_TRANPARENT tag to determine whether or not the images
# generated for formulas are transparent PNGs. Transparent PNGs are not
# supported properly for IE 6.0, but are supported on all modern browsers.
#
# Note that when changing this option you need to delete any form_*.png files in
# the HTML output directory before the changes have effect.
# The default value is: YES.
# This tag requires that the tag GENERATE_HTML is set to YES.

FORMULA_TRANSPARENT    = YES

# Enable the USE_MATHJAX option to render LaTeX formulas using MathJax (see
# http://www.mathjax.org) which uses client side Javascript for the rendering
# instead of using prerendered bitmaps. Use this if you do not have LaTeX
# installed or if you want to formulas look prettier in the HTML output. When
# enabled you may also need to install MathJax separately and configure the path
# to it using the MATHJAX_RELPATH option.
# The default value is: NO.
# This tag requires that the tag GENERATE_HTML is set to YES.

USE_MATHJAX            = NO

# When MathJax is enabled you can set the default output format to be used for
# the MathJax output. See the MathJax site (see:
# http://docs.mathjax.org/en/latest/output.html) for more details.
# Possible values are: HTML-CSS (which is slower, but has the best
# compatibility), NativeMML (i.e. MathML) and SVG.
# The default value is: HTML-CSS.
# This tag requires that the tag USE_MATHJAX is set to YES.

MATHJAX_FORMAT         = HTML-CSS

# When MathJax is enabled you need to specify the location relative to the HTML
# output directory using the MATHJAX_RELPATH option. The destination directory
# should contain the MathJax.js script. For instance, if the mathjax directory
# is located at the same level as the HTML output directory, then
# MATHJAX_RELPATH should be ../mathjax. The default value points to the MathJax
# Content Delivery Network so you can quickly see the result without installing
# MathJax. However, it is strongly recommended to install a local copy of
# MathJax from http://www.mathjax.org before deployment.
# The default value is: http://cdn.mathjax.org/mathjax/latest.
# This tag requires that the tag USE_MATHJAX is set to YES.

MATHJAX_RELPATH        = http://cdn.mathjax.org/mathjax/latest

# The MATHJAX_EXTENSIONS tag can be used to specify one or more MathJax
# extension names that should be enabled during MathJax rendering. For example
# MATHJAX_EXTENSIONS = TeX/AMSmath TeX/AMSsymbols
# This tag requires that the tag USE_MATHJAX is set to YES.

MATHJAX_EXTENSIONS     =

# The MATHJAX_CODEFILE tag can be used to specify a file with javascript pieces
# of code that will be used on startup of the MathJax code. See the MathJax site
# (see: http://docs.mathjax.org/en/latest/output.html) for more details. For an
# example see the documentation.
# This tag requires that the tag USE_MATHJAX is set to YES.

MATHJAX_CODEFILE       =

# When the SEARCHENGINE tag is enabled doxygen will generate a search box for
# the HTML output. The underlying search engine uses javascript and DHTML and
# should work on any modern browser. Note that when using HTML help
# (GENERATE_HTMLHELP), Qt help (GENERATE_QHP), or docsets (GENERATE_DOCSET)
# there is already a search function so this one should typically be disabled.
# For large projects the javascript based search engine can be slow, then
# enabling SERVER_BASED_SEARCH may provide a better solution. It is possible to
# search using the keyboard; to jump to the search box use <access key> + S
# (what the <access key> is depends on the OS and browser, but it is typically
# <CTRL>, <ALT>/<option>, or both). Inside the search box use the <cursor down
# key> to jump into the search results window, the results can be navigated
# using the <cursor keys>. Press <Enter> to select an item or <escape> to cancel
# the search. The filter options can be selected when the cursor is inside the
# search box by pressing <Shift>+<cursor down>. Also here use the <cursor keys>
# to select a filter and <Enter> or <escape> to activate or cancel the filter
# option.
# The default value is: YES.
# This tag requires that the tag GENERATE_HTML is set to YES.

SEARCHENGINE           = YES

# When the SERVER_BASED_SEARCH tag is enabled the search engine will be
# implemented using a web server instead of a web client using Javascript. There
# are two flavors of web server based searching depending on the EXTERNAL_SEARCH
# setting. When disabled, doxygen will generate a PHP script for searching and
# an index file used by the script. When EXTERNAL_SEARCH is enabled the indexing
# and searching needs to be provided by external tools. See the section
# "External Indexing and Searching" for details.
# The default value is: NO.
# This tag requires that the tag SEARCHENGINE is set to YES.

SERVER_BASED_SEARCH    = NO

# When EXTERNAL_SEARCH tag is enabled doxygen will no longer generate the PHP
# script for searching. Instead the search results are written to an XML file
# which needs to be processed by an external indexer. Doxygen will invoke an
# external search engine pointed to by the SEARCHENGINE_URL option to obtain the
# search results.
#
# Doxygen ships with an example indexer ( doxyindexer) and search engine
# (doxysearch.cgi) which are based on the open source search engine library
# Xapian (see: http://xapian.org/).
#
# See the section "External Indexing and Searching" for details.
# The default value is: NO.
# This tag requires that the tag SEARCHENGINE is set to YES.

EXTERNAL_SEARCH        = NO

# The SEARCHENGINE_URL should point to a search engine hosted by a web server
# which will return the search results when EXTERNAL_SEARCH is enabled.
#
# Doxygen ships with an example indexer ( doxyindexer) and search engine
# (doxysearch.cgi) which are based on the open source search engine library
# Xapian (see: http://xapian.org/). See the section "External Indexing and
# Searching" for details.
# This tag requires that the tag SEARCHENGINE is set to YES.

SEARCHENGINE_URL       =

# When SERVER_BASED_SEARCH and EXTERNAL_SEARCH are both enabled the unindexed
# search data is written to a file for indexing by an external tool. With the
# SEARCHDATA_FILE tag the name of this file can be specified.
# The default file is: searchdata.xml.
# This tag requires that the tag SEARCHENGINE is set to YES.

SEARCHDATA_FILE        = searchdata.xml

# When SERVER_BASED_SEARCH and EXTERNAL_SEARCH are both enabled the
# EXTERNAL_SEARCH_ID tag can be used as an identifier for the project. This is
# useful in combination with EXTRA_SEARCH_MAPPINGS to search through multiple
# projects and redirect the results back to the right project.
# This tag requires that the tag SEARCHENGINE is set to YES.

EXTERNAL_SEARCH_ID     =

# The EXTRA_SEARCH_MAPPINGS tag can be used to enable searching through doxygen
# projects other than the one defined by this configuration file, but that are
# all added to the same external search index. Each project needs to have a
# unique id set via EXTERNAL_SEARCH_ID. The search mapping then maps the id of
# to a relative location where the documentation can be found. The format is:
# EXTRA_SEARCH_MAPPINGS = tagname1=loc1 tagname2=loc2 ...
# This tag requires that the tag SEARCHENGINE is set to YES.

EXTRA_SEARCH_MAPPINGS  =

#---------------------------------------------------------------------------
# Configuration options related to the LaTeX output
#---------------------------------------------------------------------------

# If the GENERATE_LATEX tag is set to YES doxygen will generate LaTeX output.
# The default value is: YES.

GENERATE_LATEX         = YES

# The LATEX_OUTPUT tag is used to specify where the LaTeX docs will be put. If a
# relative path is entered the value of OUTPUT_DIRECTORY will be put in front of
# it.
# The default directory is: latex.
# This tag requires that the tag GENERATE_LATEX is set to YES.

LATEX_OUTPUT           = latex

# The LATEX_CMD_NAME tag can be used to specify the LaTeX command name to be
# invoked.
#
# Note that when enabling USE_PDFLATEX this option is only used for generating
# bitmaps for formulas in the HTML output, but not in the Makefile that is
# written to the output directory.
# The default file is: latex.
# This tag requires that the tag GENERATE_LATEX is set to YES.

LATEX_CMD_NAME         = latex

# The MAKEINDEX_CMD_NAME tag can be used to specify the command name to generate
# index for LaTeX.
# The default file is: makeindex.
# This tag requires that the tag GENERATE_LATEX is set to YES.

MAKEINDEX_CMD_NAME     = makeindex

# If the COMPACT_LATEX tag is set to YES doxygen generates more compact LaTeX
# documents. This may be useful for small projects and may help to save some
# trees in general.
# The default value is: NO.
# This tag requires that the tag GENERATE_LATEX is set to YES.

COMPACT_LATEX          = NO

# The PAPER_TYPE tag can be used to set the paper type that is used by the
# printer.
# Possible values are: a4 (210 x 297 mm), letter (8.5 x 11 inches), legal (8.5 x
# 14 inches) and executive (7.25 x 10.5 inches).
# The default value is: a4.
# This tag requires that the tag GENERATE_LATEX is set to YES.

PAPER_TYPE             = a4

# The EXTRA_PACKAGES tag can be used to specify one or more LaTeX package names
# that should be included in the LaTeX output. To get the times font for
# instance you can specify
# EXTRA_PACKAGES=times
# If left blank no extra packages will be included.
# This tag requires that the tag GENERATE_LATEX is set to YES.

EXTRA_PACKAGES         =

# The LATEX_HEADER tag can be used to specify a personal LaTeX header for the
# generated LaTeX document. The header should contain everything until the first
# chapter. If it is left blank doxygen will generate a standard header. See
# section "Doxygen usage" for information on how to let doxygen write the
# default header to a separate file.
#
# Note: Only use a user-defined header if you know what you are doing! The
# following commands have a special meaning inside the header: $title,
# $datetime, $date, $doxygenversion, $projectname, $projectnumber,
# $projectbrief, $projectlogo. Doxygen will replace $title with the empy string,
# for the replacement values of the other commands the user is refered to
# HTML_HEADER.
# This tag requires that the tag GENERATE_LATEX is set to YES.

LATEX_HEADER           =

# The LATEX_FOOTER tag can be used to specify a personal LaTeX footer for the
# generated LaTeX document. The footer should contain everything after the last
# chapter. If it is left blank doxygen will generate a standard footer. See
# LATEX_HEADER for more information on how to generate a default footer and what
# special commands can be used inside the footer.
#
# Note: Only use a user-defined footer if you know what you are doing!
# This tag requires that the tag GENERATE_LATEX is set to YES.

LATEX_FOOTER           =

# The LATEX_EXTRA_FILES tag can be used to specify one or more extra images or
# other source files which should be copied to the LATEX_OUTPUT output
# directory. Note that the files will be copied as-is; there are no commands or
# markers available.
# This tag requires that the tag GENERATE_LATEX is set to YES.

LATEX_EXTRA_FILES      =

# If the PDF_HYPERLINKS tag is set to YES, the LaTeX that is generated is
# prepared for conversion to PDF (using ps2pdf or pdflatex). The PDF file will
# contain links (just like the HTML output) instead of page references. This
# makes the output suitable for online browsing using a PDF viewer.
# The default value is: YES.
# This tag requires that the tag GENERATE_LATEX is set to YES.

PDF_HYPERLINKS         = YES

# If the USE_PDFLATEX tag is set to YES, doxygen will use pdflatex to generate
# the PDF file directly from the LaTeX files. Set this option to YES to get a
# higher quality PDF documentation.
# The default value is: YES.
# This tag requires that the tag GENERATE_LATEX is set to YES.

USE_PDFLATEX           = YES

# If the LATEX_BATCHMODE tag is set to YES, doxygen will add the \batchmode
# command to the generated LaTeX files. This will instruct LaTeX to keep running
# if errors occur, instead of asking the user for help. This option is also used
# when generating formulas in HTML.
# The default value is: NO.
# This tag requires that the tag GENERATE_LATEX is set to YES.

LATEX_BATCHMODE        = NO

# If the LATEX_HIDE_INDICES tag is set to YES then doxygen will not include the
# index chapters (such as File Index, Compound Index, etc.) in the output.
# The default value is: NO.
# This tag requires that the tag GENERATE_LATEX is set to YES.

LATEX_HIDE_INDICES     = NO

# If the LATEX_SOURCE_CODE tag is set to YES then doxygen will include source
# code with syntax highlighting in the LaTeX output.
#
# Note that which sources are shown also depends on other settings such as
# SOURCE_BROWSER.
# The default value is: NO.
# This tag requires that the tag GENERATE_LATEX is set to YES.

LATEX_SOURCE_CODE      = NO

# The LATEX_BIB_STYLE tag can be used to specify the style to use for the
# bibliography, e.g. plainnat, or ieeetr. See
# http://en.wikipedia.org/wiki/BibTeX and \cite for more info.
# The default value is: plain.
# This tag requires that the tag GENERATE_LATEX is set to YES.

LATEX_BIB_STYLE        = plain

#---------------------------------------------------------------------------
# Configuration options related to the RTF output
#---------------------------------------------------------------------------

# If the GENERATE_RTF tag is set to YES doxygen will generate RTF output. The
# RTF output is optimized for Word 97 and may not look too pretty with other RTF
# readers/editors.
# The default value is: NO.

GENERATE_RTF           = NO

# The RTF_OUTPUT tag is used to specify where the RTF docs will be put. If a
# relative path is entered the value of OUTPUT_DIRECTORY will be put in front of
# it.
# The default directory is: rtf.
# This tag requires that the tag GENERATE_RTF is set to YES.

RTF_OUTPUT             = rtf

# If the COMPACT_RTF tag is set to YES doxygen generates more compact RTF
# documents. This may be useful for small projects and may help to save some
# trees in general.
# The default value is: NO.
# This tag requires that the tag GENERATE_RTF is set to YES.

COMPACT_RTF            = NO

# If the RTF_HYPERLINKS tag is set to YES, the RTF that is generated will
# contain hyperlink fields. The RTF file will contain links (just like the HTML
# output) instead of page references. This makes the output suitable for online
# browsing using Word or some other Word compatible readers that support those
# fields.
#
# Note: WordPad (write) and others do not support links.
# The default value is: NO.
# This tag requires that the tag GENERATE_RTF is set to YES.

RTF_HYPERLINKS         = NO

# Load stylesheet definitions from file. Syntax is similar to doxygen's config
# file, i.e. a series of assignments. You only have to provide replacements,
# missing definitions are set to their default value.
#
# See also section "Doxygen usage" for information on how to generate the
# default style sheet that doxygen normally uses.
# This tag requires that the tag GENERATE_RTF is set to YES.

RTF_STYLESHEET_FILE    =

# Set optional variables used in the generation of an RTF document. Syntax is
# similar to doxygen's config file. A template extensions file can be generated
# using doxygen -e rtf extensionFile.
# This tag requires that the tag GENERATE_RTF is set to YES.

RTF_EXTENSIONS_FILE    =

#---------------------------------------------------------------------------
# Configuration options related to the man page output
#---------------------------------------------------------------------------

# If the GENERATE_MAN tag is set to YES doxygen will generate man pages for
# classes and files.
# The default value is: NO.

GENERATE_MAN           = NO

# The MAN_OUTPUT tag is used to specify where the man pages will be put. If a
# relative path is entered the value of OUTPUT_DIRECTORY will be put in front of
# it. A directory man3 will be created inside the directory specified by
# MAN_OUTPUT.
# The default directory is: man.
# This tag requires that the tag GENERATE_MAN is set to YES.

MAN_OUTPUT             = man

# The MAN_EXTENSION tag determines the extension that is added to the generated
# man pages. In case the manual section does not start with a number, the number
# 3 is prepended. The dot (.) at the beginning of the MAN_EXTENSION tag is
# optional.
# The default value is: .3.
# This tag requires that the tag GENERATE_MAN is set to YES.

MAN_EXTENSION          = .3

# The MAN_SUBDIR tag determines the name of the directory created within
# MAN_OUTPUT in which the man pages are placed. If defaults to man followed by
# MAN_EXTENSION with the initial . removed.
# This tag requires that the tag GENERATE_MAN is set to YES.

MAN_SUBDIR             =

# If the MAN_LINKS tag is set to YES and doxygen generates man output, then it
# will generate one additional man file for each entity documented in the real
# man page(s). These additional files only source the real man page, but without
# them the man command would be unable to find the correct page.
# The default value is: NO.
# This tag requires that the tag GENERATE_MAN is set to YES.

MAN_LINKS              = NO

#---------------------------------------------------------------------------
# Configuration options related to the XML output
#---------------------------------------------------------------------------

# If the GENERATE_XML tag is set to YES doxygen will generate an XML file that
# captures the structure of the code including all documentation.
# The default value is: NO.

GENERATE_XML           = NO

# The XML_OUTPUT tag is used to specify where the XML pages will be put. If a
# relative path is entered the value of OUTPUT_DIRECTORY will be put in front of
# it.
# The default directory is: xml.
# This tag requires that the tag GENERATE_XML is set to YES.

XML_OUTPUT             = xml

# If the XML_PROGRAMLISTING tag is set to YES doxygen will dump the program
# listings (including syntax highlighting and cross-referencing information) to
# the XML output. Note that enabling this will significantly increase the size
# of the XML output.
# The default value is: YES.
# This tag requires that the tag GENERATE_XML is set to YES.

XML_PROGRAMLISTING     = YES

#---------------------------------------------------------------------------
# Configuration options related to the DOCBOOK output
#---------------------------------------------------------------------------

# If the GENERATE_DOCBOOK tag is set to YES doxygen will generate Docbook files
# that can be used to generate PDF.
# The default value is: NO.

GENERATE_DOCBOOK       = NO

# The DOCBOOK_OUTPUT tag is used to specify where the Docbook pages will be put.
# If a relative path is entered the value of OUTPUT_DIRECTORY will be put in
# front of it.
# The default directory is: docbook.
# This tag requires that the tag GENERATE_DOCBOOK is set to YES.

DOCBOOK_OUTPUT         = docbook

# If the DOCBOOK_PROGRAMLISTING tag is set to YES doxygen will include the
# program listings (including syntax highlighting and cross-referencing
# information) to the DOCBOOK output. Note that enabling this will significantly
# increase the size of the DOCBOOK output.
# The default value is: NO.
# This tag requires that the tag GENERATE_DOCBOOK is set to YES.

DOCBOOK_PROGRAMLISTING = NO

#---------------------------------------------------------------------------
# Configuration options for the AutoGen Definitions output
#---------------------------------------------------------------------------

# If the GENERATE_AUTOGEN_DEF tag is set to YES doxygen will generate an AutoGen
# Definitions (see http://autogen.sf.net) file that captures the structure of
# the code including all documentation. Note that this feature is still
# experimental and incomplete at the moment.
# The default value is: NO.

GENERATE_AUTOGEN_DEF   = NO

#---------------------------------------------------------------------------
# Configuration options related to the Perl module output
#---------------------------------------------------------------------------

# If the GENERATE_PERLMOD tag is set to YES doxygen will generate a Perl module
# file that captures the structure of the code including all documentation.
#
# Note that this feature is still experimental and incomplete at the moment.
# The default value is: NO.

GENERATE_PERLMOD       = NO

# If the PERLMOD_LATEX tag is set to YES doxygen will generate the necessary
# Makefile rules, Perl scripts and LaTeX code to be able to generate PDF and DVI
# output from the Perl module output.
# The default value is: NO.
# This tag requires that the tag GENERATE_PERLMOD is set to YES.

PERLMOD_LATEX          = NO

# If the PERLMOD_PRETTY tag is set to YES the Perl module output will be nicely
# formatted so it can be parsed by a human reader. This is useful if you want to
# understand what is going on. On the other hand, if this tag is set to NO the
# size of the Perl module output will be much smaller and Perl will parse it
# just the same.
# The default value is: YES.
# This tag requires that the tag GENERATE_PERLMOD is set to YES.

PERLMOD_PRETTY         = YES

# The names of the make variables in the generated doxyrules.make file are
# prefixed with the string contained in PERLMOD_MAKEVAR_PREFIX. This is useful
# so different doxyrules.make files included by the same Makefile don't
# overwrite each other's variables.
# This tag requires that the tag GENERATE_PERLMOD is set to YES.

PERLMOD_MAKEVAR_PREFIX =

#---------------------------------------------------------------------------
# Configuration options related to the preprocessor
#---------------------------------------------------------------------------

# If the ENABLE_PREPROCESSING tag is set to YES doxygen will evaluate all
# C-preprocessor directives found in the sources and include files.
# The default value is: YES.

ENABLE_PREPROCESSING   = YES

# If the MACRO_EXPANSION tag is set to YES doxygen will expand all macro names
# in the source code. If set to NO only conditional compilation will be
# performed. Macro expansion can be done in a controlled way by setting
# EXPAND_ONLY_PREDEF to YES.
# The default value is: NO.
# This tag requires that the tag ENABLE_PREPROCESSING is set to YES.

MACRO_EXPANSION        = NO

# If the EXPAND_ONLY_PREDEF and MACRO_EXPANSION tags are both set to YES then
# the macro expansion is limited to the macros specified with the PREDEFINED and
# EXPAND_AS_DEFINED tags.
# The default value is: NO.
# This tag requires that the tag ENABLE_PREPROCESSING is set to YES.

EXPAND_ONLY_PREDEF     = NO

# If the SEARCH_INCLUDES tag is set to YES the includes files in the
# INCLUDE_PATH will be searched if a #include is found.
# The default value is: YES.
# This tag requires that the tag ENABLE_PREPROCESSING is set to YES.

SEARCH_INCLUDES        = YES

# The INCLUDE_PATH tag can be used to specify one or more directories that
# contain include files that are not input files but should be processed by the
# preprocessor.
# This tag requires that the tag SEARCH_INCLUDES is set to YES.

INCLUDE_PATH           = ./include


# You can use the INCLUDE_FILE_PATTERNS tag to specify one or more wildcard
# patterns (like *.h and *.hpp) to filter out the header-files in the
# directories. If left blank, the patterns specified with FILE_PATTERNS will be
# used.
# This tag requires that the tag ENABLE_PREPROCESSING is set to YES.

INCLUDE_FILE_PATTERNS  = *h

# The PREDEFINED tag can be used to specify one or more macro names that are
# defined before the preprocessor is started (similar to the -D option of e.g.
# gcc). The argument of the tag is a list of macros of the form: name or
# name=definition (no spaces). If the definition and the "=" are omitted, "=1"
# is assumed. To prevent a macro definition from being undefined via #undef or
# recursively expanded use the := operator instead of the = operator.
# This tag requires that the tag ENABLE_PREPROCESSING is set to YES.

PREDEFINED             =

# If the MACRO_EXPANSION and EXPAND_ONLY_PREDEF tags are set to YES then this
# tag can be used to specify a list of macro names that should be expanded. The
# macro definition that is found in the sources will be used. Use the PREDEFINED
# tag if you want to use a different macro definition that overrules the
# definition found in the source code.
# This tag requires that the tag ENABLE_PREPROCESSING is set to YES.

EXPAND_AS_DEFINED      =

# If the SKIP_FUNCTION_MACROS tag is set to YES then doxygen's preprocessor will
# remove all references to function-like macros that are alone on a line, have
# an all uppercase name, and do not end with a semicolon. Such function macros
# are typically used for boiler-plate code, and will confuse the parser if not
# removed.
# The default value is: YES.
# This tag requires that the tag ENABLE_PREPROCESSING is set to YES.

SKIP_FUNCTION_MACROS   = YES

#---------------------------------------------------------------------------
# Configuration options related to external references
#---------------------------------------------------------------------------

# The TAGFILES tag can be used to specify one or more tag files. For each tag
# file the location of the external documentation should be added. The format of
# a tag file without this location is as follows:
# TAGFILES = file1 file2 ...
# Adding location for the tag files is done as follows:
# TAGFILES = file1=loc1 "file2 = loc2" ...
# where loc1 and loc2 can be relative or absolute paths or URLs. See the
# section "Linking to external documentation" for more information about the use
# of tag files.
# Note: Each tag file must have a unique name (where the name does NOT include
# the path). If a tag file is not located in the directory in which doxygen is
# run, you must also specify the path to the tagfile here.

TAGFILES               =

# When a file name is specified after GENERATE_TAGFILE, doxygen will create a
# tag file that is based on the input files it reads. See section "Linking to
# external documentation" for more information about the usage of tag files.

GENERATE_TAGFILE       =

# If the ALLEXTERNALS tag is set to YES all external class will be listed in the
# class index. If set to NO only the inherited external classes will be listed.
# The default value is: NO.

ALLEXTERNALS           = NO

# If the EXTERNAL_GROUPS tag is set to YES all external groups will be listed in
# the modules index. If set to NO, only the current project's groups will be
# listed.
# The default value is: YES.

EXTERNAL_GROUPS        = YES

# If the EXTERNAL_PAGES tag is set to YES all external pages will be listed in
# the related pages index. If set to NO, only the current project's pages will
# be listed.
# The default value is: YES.

EXTERNAL_PAGES         = YES

# The PERL_PATH should be the absolute path and name of the perl script
# interpreter (i.e. the result of 'which perl').
# The default file (with absolute path) is: /usr/bin/perl.

PERL_PATH              = /usr/bin/perl

#---------------------------------------------------------------------------
# Configuration options related to the dot tool
#---------------------------------------------------------------------------

# If the CLASS_DIAGRAMS tag is set to YES doxygen will generate a class diagram
# (in HTML and LaTeX) for classes with base or super classes. Setting the tag to
# NO turns the diagrams off. Note that this option also works with HAVE_DOT
# disabled, but it is recommended to install and use dot, since it yields more
# powerful graphs.
# The default value is: YES.

CLASS_DIAGRAMS         = YES

# You can define message sequence charts within doxygen comments using the \msc
# command. Doxygen will then run the mscgen tool (see:
# http://www.mcternan.me.uk/mscgen/)) to produce the chart and insert it in the
# documentation. The MSCGEN_PATH tag allows you to specify the directory where
# the mscgen tool resides. If left empty the tool is assumed to be found in the
# default search path.

MSCGEN_PATH            =

# You can include diagrams made with dia in doxygen documentation. Doxygen will
# then run dia to produce the diagram and insert it in the documentation. The
# DIA_PATH tag allows you to specify the directory where the dia binary resides.
# If left empty dia is assumed to be found in the default search path.

DIA_PATH               =

# If set to YES, the inheritance and collaboration graphs will hide inheritance
# and usage relations if the target is undocumented or is not a class.
# The default value is: YES.

HIDE_UNDOC_RELATIONS   = YES

# If you set the HAVE_DOT tag to YES then doxygen will assume the dot tool is
# available from the path. This tool is part of Graphviz (see:
# http://www.graphviz.org/), a graph visualization toolkit from AT&T and Lucent
# Bell Labs. The other options in this section have no effect if this option is
# set to NO
# The default value is: YES.

HAVE_DOT               = YES

# The DOT_NUM_THREADS specifies the number of dot invocations doxygen is allowed
# to run in parallel. When set to 0 doxygen will base this on the number of
# processors available in the system. You can set it explicitly to a value
# larger than 0 to get control over the balance between CPU load and processing
# speed.
# Minimum value: 0, maximum value: 32, default value: 0.
# This tag requires that the tag HAVE_DOT is set to YES.

DOT_NUM_THREADS        = 0

# When you want a differently looking font in the dot files that doxygen
# generates you can specify the font name using DOT_FONTNAME. You need to make
# sure dot is able to find the font, which can be done by putting it in a
# standard location or by setting the DOTFONTPATH environment variable or by
# setting DOT_FONTPATH to the directory containing the font.
# The default value is: Helvetica.
# This tag requires that the tag HAVE_DOT is set to YES.

DOT_FONTNAME           = Helvetica

# The DOT_FONTSIZE tag can be used to set the size (in points) of the font of
# dot graphs.
# Minimum value: 4, maximum value: 24, default value: 10.
# This tag requires that the tag HAVE_DOT is set to YES.

DOT_FONTSIZE           = 10

# By default doxygen will tell dot to use the default font as specified with
# DOT_FONTNAME. If you specify a different font using DOT_FONTNAME you can set
# the path where dot can find it using this tag.
# This tag requires that the tag HAVE_DOT is set to YES.

DOT_FONTPATH           =

# If the CLASS_GRAPH tag is set to YES then doxygen will generate a graph for
# each documented class showing the direct and indirect inheritance relations.
# Setting this tag to YES will force the CLASS_DIAGRAMS tag to NO.
# The default value is: YES.
# This tag requires that the tag HAVE_DOT is set to YES.

CLASS_GRAPH            = YES

# If the COLLABORATION_GRAPH tag is set to YES then doxygen will generate a
# graph for each documented class showing the direct and indirect implementation
# dependencies (inheritance, containment, and class references variables) of the
# class with other documented classes.
# The default value is: YES.
# This tag requires that the tag HAVE_DOT is set to YES.

COLLABORATION_GRAPH    = YES

# If the GROUP_GRAPHS tag is set to YES then doxygen will generate a graph for
# groups, showing the direct groups dependencies.
# The default value is: YES.
# This tag requires that the tag HAVE_DOT is set to YES.

GROUP_GRAPHS           = YES

# If the UML_LOOK tag is set to YES doxygen will generate inheritance and
# collaboration diagrams in a style similar to the OMG's Unified Modeling
# Language.
# The default value is: NO.
# This tag requires that the tag HAVE_DOT is set to YES.

UML_LOOK               = NO

# If the UML_LOOK tag is enabled, the fields and methods are shown inside the
# class node. If there are many fields or methods and many nodes the graph may
# become too big to be useful. The UML_LIMIT_NUM_FIELDS threshold limits the
# number of items for each type to make the size more manageable. Set this to 0
# for no limit. Note that the threshold may be exceeded by 50% before the limit
# is enforced. So when you set the threshold to 10, up to 15 fields may appear,
# but if the number exceeds 15, the total amount of fields shown is limited to
# 10.
# Minimum value: 0, maximum value: 100, default value: 10.
# This tag requires that the tag HAVE_DOT is set to YES.

UML_LIMIT_NUM_FIELDS   = 10

# If the TEMPLATE_RELATIONS tag is set to YES then the inheritance and
# collaboration graphs will show the relations between templates and their
# instances.
# The default value is: NO.
# This tag requires that the tag HAVE_DOT is set to YES.

TEMPLATE_RELATIONS     = NO

# If the INCLUDE_GRAPH, ENABLE_PREPROCESSING and SEARCH_INCLUDES tags are set to
# YES then doxygen will generate a graph for each documented file showing the
# direct and indirect include dependencies of the file with other documented
# files.
# The default value is: YES.
# This tag requires that the tag HAVE_DOT is set to YES.

INCLUDE_GRAPH          = YES

# If the INCLUDED_BY_GRAPH, ENABLE_PREPROCESSING and SEARCH_INCLUDES tags are
# set to YES then doxygen will generate a graph for each documented file showing
# the direct and indirect include dependencies of the file with other documented
# files.
# The default value is: YES.
# This tag requires that the tag HAVE_DOT is set to YES.

INCLUDED_BY_GRAPH      = YES

# If the CALL_GRAPH tag is set to YES then doxygen will generate a call
# dependency graph for every global function or class method.
#
# Note that enabling this option will significantly increase the time of a run.
# So in most cases it will be better to enable call graphs for selected
# functions only using the \callgraph command.
# The default value is: NO.
# This tag requires that the tag HAVE_DOT is set to YES.

CALL_GRAPH             = NO

# If the CALLER_GRAPH tag is set to YES then doxygen will generate a caller
# dependency graph for every global function or class method.
#
# Note that enabling this option will significantly increase the time of a run.
# So in most cases it will be better to enable caller graphs for selected
# functions only using the \callergraph command.
# The default value is: NO.
# This tag requires that the tag HAVE_DOT is set to YES.

CALLER_GRAPH           = NO

# If the GRAPHICAL_HIERARCHY tag is set to YES then doxygen will graphical
# hierarchy of all classes instead of a textual one.
# The default value is: YES.
# This tag requires that the tag HAVE_DOT is set to YES.

GRAPHICAL_HIERARCHY    = YES

# If the DIRECTORY_GRAPH tag is set to YES then doxygen will show the
# dependencies a directory has on other directories in a graphical way. The
# dependency relations are determined by the #include relations between the
# files in the directories.
# The default value is: YES.
# This tag requires that the tag HAVE_DOT is set to YES.

DIRECTORY_GRAPH        = YES

# The DOT_IMAGE_FORMAT tag can be used to set the image format of the images
# generated by dot.
# Note: If you choose svg you need to set HTML_FILE_EXTENSION to xhtml in order
# to make the SVG files visible in IE 9+ (other browsers do not have this
# requirement).
# Possible values are: png, png:cairo, png:cairo:cairo, png:cairo:gd, png:gd,
# png:gd:gd, jpg, jpg:cairo, jpg:cairo:gd, jpg:gd, jpg:gd:gd, gif, gif:cairo,
# gif:cairo:gd, gif:gd, gif:gd:gd and svg.
# The default value is: png.
# This tag requires that the tag HAVE_DOT is set to YES.

DOT_IMAGE_FORMAT       = png

# If DOT_IMAGE_FORMAT is set to svg, then this option can be set to YES to
# enable generation of interactive SVG images that allow zooming and panning.
#
# Note that this requires a modern browser other than Internet Explorer. Tested
# and working are Firefox, Chrome, Safari, and Opera.
# Note: For IE 9+ you need to set HTML_FILE_EXTENSION to xhtml in order to make
# the SVG files visible. Older versions of IE do not have SVG support.
# The default value is: NO.
# This tag requires that the tag HAVE_DOT is set to YES.

INTERACTIVE_SVG        = NO

# The DOT_PATH tag can be used to specify the path where the dot tool can be
# found. If left blank, it is assumed the dot tool can be found in the path.
# This tag requires that the tag HAVE_DOT is set to YES.

DOT_PATH               =

# The DOTFILE_DIRS tag can be used to specify one or more directories that
# contain dot files that are included in the documentation (see the \dotfile
# command).
# This tag requires that the tag HAVE_DOT is set to YES.

DOTFILE_DIRS           =

# The MSCFILE_DIRS tag can be used to specify one or more directories that
# contain msc files that are included in the documentation (see the \mscfile
# command).

MSCFILE_DIRS           =

# The DIAFILE_DIRS tag can be used to specify one or more directories that
# contain dia files that are included in the documentation (see the \diafile
# command).

DIAFILE_DIRS           =

# When using plantuml, the PLANTUML_JAR_PATH tag should be used to specify the
# path where java can find the plantuml.jar file. If left blank, it is assumed
# PlantUML is not used or called during a preprocessing step. Doxygen will
# generate a warning when it encounters a \startuml command in this case and
# will not generate output for the diagram.
# This tag requires that the tag HAVE_DOT is set to YES.

PLANTUML_JAR_PATH      =

# The DOT_GRAPH_MAX_NODES tag can be used to set the maximum number of nodes
# that will be shown in the graph. If the number of nodes in a graph becomes
# larger than this value, doxygen will truncate the graph, which is visualized
# by representing a node as a red box. Note that doxygen if the number of direct
# children of the root node in a graph is already larger than
# DOT_GRAPH_MAX_NODES then the graph will not be shown at all. Also note that
# the size of a graph can be further restricted by MAX_DOT_GRAPH_DEPTH.
# Minimum value: 0, maximum value: 10000, default value: 50.
# This tag requires that the tag HAVE_DOT is set to YES.

DOT_GRAPH_MAX_NODES    = 50

# The MAX_DOT_GRAPH_DEPTH tag can be used to set the maximum depth of the graphs
# generated by dot. A depth value of 3 means that only nodes reachable from the
# root by following a path via at most 3 edges will be shown. Nodes that lay
# further from the root node will be omitted. Note that setting this option to 1
# or 2 may greatly reduce the computation time needed for large code bases. Also
# note that the size of a graph can be further restricted by
# DOT_GRAPH_MAX_NODES. Using a depth of 0 means no depth restriction.
# Minimum value: 0, maximum value: 1000, default value: 0.
# This tag requires that the tag HAVE_DOT is set to YES.

MAX_DOT_GRAPH_DEPTH    = 0

# Set the DOT_TRANSPARENT tag to YES to generate images with a transparent
# background. This is disabled by default, because dot on Windows does not seem
# to support this out of the box.
#
# Warning: Depending on the platform used, enabling this option may lead to
# badly anti-aliased labels on the edges of a graph (i.e. they become hard to
# read).
# The default value is: NO.
# This tag requires that the tag HAVE_DOT is set to YES.

DOT_TRANSPARENT        = NO

# Set the DOT_MULTI_TARGETS tag to YES allow dot to generate multiple output
# files in one run (i.e. multiple -o and -T options on the command line). This
# makes dot run faster, but since only newer versions of dot (>1.8.10) support
# this, this feature is disabled by default.
# The default value is: NO.
# This tag requires that the tag HAVE_DOT is set to YES.

DOT_MULTI_TARGETS      = NO

# If the GENERATE_LEGEND tag is set to YES doxygen will generate a legend page
# explaining the meaning of the various boxes and arrows in the dot generated
# graphs.
# The default value is: YES.
# This tag requires that the tag HAVE_DOT is set to YES.

GENERATE_LEGEND        = YES

# If the DOT_CLEANUP tag is set to YES doxygen will remove the intermediate dot
# files that are used to generate the various graphs.
# The default value is: YES.
# This tag requires that the tag HAVE_DOT is set to YES.

DOT_CLEANUP            = YES
