## Supplementary material for "FastqPuri: high-performance preprocessing of RNA-seq data": bf_performance.sh

```
# Create bloom filters and construct ROC curves 

# Executables: 

makeBloom=../../bin/makeBloom
trimFilter=../../bin/trimFilter

##################################################################
# STEP 0: Generate Fastq files                                   #
##################################################################

# human_reads.fq.gz : 10e5 human reads created with dgwin 
# dwgsim -1 150 -2 0 -N 100000 <HUMAN_GENOME> <OUTPUT_PREFIX>
# EColi_reads.fq.gz : 10e5 ECOLI reads created with dgwin 
# dwgsim -1 150 -2 0 -N 100000 <ECOLI_GENOME> <OUTPUT_PREFIX>

##################################################################
# STEP1: Create bloom filters. FPR: 0.005, 0.0075, 0.01, 0.02    #
##################################################################
$makeBloom -o EColi_0p02   -f ../fa_fq_files/EColi_genome.fa -p 0.02 
$makeBloom -o EColi_0p01   -f ../fa_fq_files/EColi_genome.fa -p 0.01
$makeBloom -o EColi_0p0075 -f ../fa_fq_files/EColi_genome.fa -p 0.0075
$makeBloom -o EColi_0p005  -f ../fa_fq_files/EColi_genome.fa -p 0.005


##################################################################
# STEP2: run trimFilter on the data                              #
##################################################################
pattern1="\- Reads accepted as good:";
pattern2="\- Discarded due to cont"
EColi_fq="../fa_fq_files/EColi_reads.fq.gz"
human_fq="../fa_fq_files/human_reads.fq.gz"

for FPR in 0p02 0p01 0p0075 0p005; do  
   for s in $(seq 0.05 0.01 0.2) ; do 
      $trimFilter -f "${EColi_fq}"  -l 150 --method BLOOM \
         --idx EColi_${FPR}.bf:"$s" 2>&1 | egrep "$pattern1|$pattern2" | \
      awk -v score="$s" ' 
       BEGIN {line = 1}
       { arr[line] = $6 ; line ++}
       END {
       FN = arr[1]/(arr[1] + arr[2])
           TP = arr[2]/(arr[1] + arr[2])
           printf("%1.8f %1.8f %1.8f \n", score,FN,TP)
       }'
   done > ooo
   
   for s in $(seq 0.05 0.01 0.2) ; do 
      $trimFilter -f "${human_fq}" -l 150 --method BLOOM \
         --idx EColi_${FPR}.bf:"$s" 2>&1 | egrep "$pattern1|$pattern2" | \
      awk -v score="$s" ' 
       BEGIN {line = 1}
       {arr[line] = $6 ; line ++}
       END {
           TN = arr[1]/(arr[1] + arr[2])
           FP = arr[2]/(arr[1] + arr[2])
           printf("%1.8f %1.8f %1.8f \n", score,TN,FP)
       }'
   done > iii
   
   echo "FN,TP,TN,FP" > ROC_${FPR}_bloom.csv                                       
   paste ooo iii | awk '{printf("%f,%f,%f,%f,%f\n",$1,$2,$3,$5,$6)}' \
    >> ROC_${FPR}_bloom.csv
done

rm ooo iii 
rm out*

##################################################################
# STEP3: run the R script to create the plots                    #
##################################################################
R -f create_ROC.R
```
