## Supplementary material for "FastqPuri: high-performance preprocessing of RNA-seq data": create_ROC.R

```
# Generates ROC curves.
tags <-  c("0p02", "0p01", "0p0075", "0p005")
for (FPR_text in tags) {
   bloom <- read.csv(paste0("ROC_",FPR_text,"_bloom.csv"))
   FPR = bloom[,4]
   TPR = bloom[,2]
   pdf(paste0("ROC_",FPR_text,"_bloom.pdf"))
   plot(FPR,TPR, main="ROC curves", 
     xlab="False positive rate",ylab="sensitivity", 
     xlim = c(min(FPR),max(FPR)),
     ylim = c(min(TPR),max(TPR)),
     type="o", col="blue")
   dev.off()
}
```
