## Supplementary material for "FastqPuri: high-performance preprocessing of RNA-seq data": README

################################################################################
# Testing bloom filter performance                                             #
################################################################################
################################################################################
# Execute the script bf_performance.sh to test the bloom filter performance.   #
#                                                                              #
# - We analyse the fastq files:                                                # 
#     * human_reads.fq.gz                                                      #
#     * EColi_reads.fq.gz                                                      #
#   they contain 10e5 reads each generated with dgwin                          #
# - STEP1: Create bloom filters for EColi genome  with FPR:                    #
#          [0.005 0.0075 0.01 0.02]                                            #
#                                                                              #
# - STEP2: Run trimFilter on both data looking for contaminations from EColi   #
#          using all filters generated, with kmersize = 25 and scores ranging  #
#          from 0.05 to 0.2 by 0.01 intervals. We obtain false/true positive/  #
#          negative rates:                                                     #
#          * FPR: % contaminations detected in human_reads.fq.gz               #
#          * TNR (specificity): % good reads detected in human_reads.fq.gz     #
#          * FNR: % good reads detected in EColi_reads.fq.gz                   #
#          * TPR (sensitivity): % contaminations detected in EColi_reads.fq.gz #
#                                                                              #
# - STEP3: Create ROC curves for all filters (sensitivity vs FPR)              #
#                                                                              #
################################################################################
