## Supplementary material for "FastqPuri: high-performance preprocessing of RNA-seq data": simReads.pl

```
#!/usr/bin/perl

#
# simRead.pl
#
# Generate reads from a target sequence
#
# written by Juliane Dohm and Claudio Lottaz
# Max Planck Institute for Molecular Genetics (Berlin, Germany)
# November - December 2006
#
# Modified by Paula Perez Rubio: september 2017
#
if (@ARGV != 5) {
    print("Please, enter 5 arguments:\n",
	  "1. sequence file in fasta format\n",
	  "2. number of reads (in mio)\n", 
	  "3. % error rate per base call\n",
	  "4. length of reads.\n",
	  "5. 1: genome is cyclic\n");
    print "Example:\n";
    print "perl simReads.pl seq.fasta 3 1.4 25 0\n\n";
    exit;
}


$sequenz = $ARGV[0];
$anznmere = $ARGV[1] * 1000000;
$prozfehler = $ARGV[2];
$readlen = $ARGV[3];
$cyclic = $ARGV[4];

######### lies sequenz

#open(SEQ, "< $sequenz") or die "$!\n";
if ($sequenz =~ /.gz$/) {
   open(SEQ, "gunzip -c $sequenz |") or die "$!\n";
}
else {
   open(SEQ, $sequenz) or die "$!\n";
}


@seq = <SEQ>;
close(SEQ);
chomp(@seq);
shift(@seq);		#das war die description-Zeile

foreach $zeile (@seq) { if (not($zeile =~ /^>/)){  $langezeile .= $zeile;} }
$langezeile =~ s/\r//g; # remove LF as used in Windows
$seqlen = length($langezeile);
print STDERR "Sequence length is $seqlen\n";

############ haenge Start ans Ende f�r cyklisches Genom

if ($cyclic == "1") {
    print STDERR "Consider the genome to be cyclic\n";
    $langezeile = $langezeile . substr($langezeile, 0, $readlen);
}


############ erstelle den Gegenstrang

$revlangezeile = reverse($langezeile);
$revlangezeile =~ tr/ATCG/TAGC/;

############ erstelle alle n-mere
$i = 0;
$fehlerzaehler = 0;
$anz_zaehler = 1;
while ($anz_zaehler <= $anznmere) {
    $i = int(rand($seqlen - $readlen));
    if (rand(1) > 0.5) { $mer = substr($langezeile,$i,$readlen); 
       push(@pos,$i); 
    }
    else { 
       $mer = substr($revlangezeile,$i,$readlen);
       push(@pos,-$i); 
    }
    $alt_mer = $mer;

    #### mit Fehler ####
    (@basen) = split(//,$mer);
    $mer = "";
    foreach $base (@basen) {
	$zufall = rand(100);
    	if ($zufall > $prozfehler) { 
	    $mer .= $base; 
	} else{
	    %moeglich = ("A"=>1, "C"=>1, "G"=>1, "T"=>1);
	    delete $moeglich{$base};
	    @moeglich = keys(%moeglich);
	    if ($zufall < $prozfehler/3) { 
		$mer .= @moeglich[0];
	    } elsif ($zufall < 2*$prozfehler/3) { 
		$mer .= @moeglich[1];
	    } else {
		$mer .= @moeglich[2];
	    }
	}
    }
    if ($alt_mer ne $mer) { ++$fehlerzaehler; }
    
    push (@mers,$mer);
    $anz_zaehler++;
    print STDERR "$anz_zaehler.." if ($anz_zaehler % 100000 == 0);	
}
print STDERR "\n";

########### n-mere ausgeben

$i = 1;
foreach (@mers) {
    print "@ read $i\n$_\n+position: $pos[$i-1] \n";
    print "I"x$readlen; print "\n";
    $i++;
}

$len = @mers;
print STDERR "$prozfehler% error per base: $fehlerzaehler faulty n-mers (total: $len n-mers)\n";

####### end of file
```
