## Supplementary material for "FastqPuri: high-performance preprocessing of RNA-seq data": README

################################################################################
#  Qreport example:                                                            #
################################################################################
#                                                                              # 
# Run the example again and compare it with                                    #
# the available results running the following commands                         #
#                                                                              #
# 1. Create a new folder and perform there the tests                           #
#    mkdir run_test                                                            #
#    cd run_test                                                               # 
#                                                                              #
# 2. Run the C executable                                                      #
#  ../../../bin/Qreport -i ../test.fq.bz2  -l 51 -o my_test_output             #
#                                                                              #
# 3. Compare the files:                                                        #
#  - ../test_output.bin      <-> my_test_output.bin                            #
#  - ../test_output.info <-> my_test_output.info                       #
#  - ../test_output.html     <-> my_test_output.html                           #
################################################################################
################################################################################
#  Sreport example:                                                            #
################################################################################
#                                                                              #
# Run the example again in your created folder.                                #
#                                                                              #
# 1. Run the C executable                                                      #
#  ../../../bin/Sreport -i ./ -o my_test_summary_report -t Q                   # 
#                                                                              #
# 2. Compare the files                                                         #
#  - ../test_summary_report.html <-> my_test_summary_report.html               #
#                                                                              #
################################################################################
