## Supplementary material for "FastqPuri: high-performance preprocessing of RNA-seq data": test_output.html

Assessing the quality of the reads


### Assessing the quality of the reads

#### General information

Running on version 1.0

| Var | Value |
| --- | --- |
| Input file name | test\_output.bin |
| Read length | 51 |
| Min good quality | 27 |
| Number of reads | 500000 |
| Number of highQ reads | 336682 |
| Number of tiles | 20 |
| Number of lanes | 2 |
| Qualities | 2 (#), 14 (/), 22 (7), 27 (<), 33 (B), 37 (F) |
| Reads with N’s | 812 |
| Number of N’s | 1114 |

#### Per base sequence quality

#### # reads with at least `m` low Q nucleotides

#### Low Q nucleotide proportion per tile per lane

#### Average quality per position per tile per lane

#### Low Q nucleotides proportion per position per tile per lane

#### Low Q nucleotides proportion per position for all tiles

#### Nucleotide content per position
