## Supplementary material for "FastqPuri: high-performance preprocessing of RNA-seq data": test_summary_report.html

Summary quality report


### Summary quality report

Running on version 1.0

#### General data

|  | # reads | # tiles | % lowQ reads | % reads with N’s |
| --- | --- | --- | --- | --- |
| test\_output | 5e+05 | 20 | 32.6636 | 0.1624 |

#### Mean quality
