## Supplementary material for "FastqPuri: high-performance preprocessing of RNA-seq data": README

################################################################################
#  trimFilter examples (adapters contamination):                               #
################################################################################
#                                                                              # 
# The script `run_examples.sh` runs trimFilter with the option `--adapter`     # 
# for 8 different `fq` files, covering all combinations of the following       #
# options:                                                                     #
#  - read length even/odd.                                                     #
#  - adapter length even/odd.                                                  #
#  - adapter length >=16(long)/<16(short)                                      #
#                                                                              # 
# In all files, the reads inserted artificially try to cover (almost) all      #
# possible cases that can arise.                                               #
#                                                                              # 
# The file `contaminant_list_10032016.txt` contains a list of possible adapter #
# sequences.                                                                   #
#                                                                              # 
################################################################################
