## Supplementary material for "FastqPuri: high-performance preprocessing of RNA-seq data": run_examples.sh

```
#!/bin/bash

# Set variables
trimFilter="../../../bin/trimFilter"
adevL="../../fa_fq_files/adapter_even_long.fa"
adevS="../../fa_fq_files/adapter_even_short.fa"
adodL="../../fa_fq_files/adapter_odd_long.fa"
adodS="../../fa_fq_files/adapter_odd_short.fa"
fqev_adevL="../../fa_fq_files/human_even_wad_even_long.fq.gz"
fqev_adevS="../../fa_fq_files/human_even_wad_even_short.fq.gz"
fqev_adodL="../../fa_fq_files/human_even_wad_odd_long.fq.gz"
fqev_adodS="../../fa_fq_files/human_even_wad_odd_short.fq.gz"
fqod_adevL="../../fa_fq_files/human_odd_wad_even_long.fq.gz"
fqod_adevS="../../fa_fq_files/human_odd_wad_even_short.fq.gz"
fqod_adodL="../../fa_fq_files/human_odd_wad_odd_long.fq.gz"
fqod_adodS="../../fa_fq_files/human_odd_wad_odd_short.fq.gz"


# Run even fq with long even adapter
$trimFilter --ifq ${fqev_adevL}  --length 150 --adapter ${adevL}:2:5.3 \
            --output evev_long
$trimFilter --ifq ${fqod_adevL}  --length 149 --adapter ${adevL}:2:5.3 \
            --output odev_long
$trimFilter --ifq ${fqev_adodL}  --length 150 --adapter ${adodL}:2:5.3 \
            --output evod_long
$trimFilter --ifq ${fqod_adodL}  --length 149 --adapter ${adodL}:2:5.3 \
            --output odod_long
$trimFilter --ifq ${fqev_adevS}  --length 150 --adapter ${adevS}:2:5.3 \
            --output evev_short
$trimFilter --ifq ${fqod_adevS}  --length 149 --adapter ${adevS}:2:5.3 \
            --output odev_short
$trimFilter --ifq ${fqev_adodS}  --length 150 --adapter ${adodS}:2:5.3 \
            --output evod_short
$trimFilter --ifq ${fqod_adodS}  --length 151 --adapter ${adodS}:2:5.3 \
            --output odod_short
```
