## Supplementary material for "FastqPuri: high-performance preprocessing of RNA-seq data": create_fake_bins.R

```
# this small script creates fake *bin data
# as if they were ./trimFilter output 
# so that we can run an example of ./Sreport 
# on ./trimFiltr output data. 


filter <- c(0,1,4,2)
for (i in 1:30) { 
   nreads <- sample(100000:200000,1)
   all_discarded <- sample(20000:40000,1)
   ngood <- nreads  - all_discarded
   trimmed <- c(0, 0, sample(10000:20000,1), sample(1000:2000,1))
   NNNN_disc <- sample(1000:2000, 1)
   lowQ_disc <- sample(5000:10000, 1)
   cont_disc <- all_discarded - lowQ_disc - NNNN_disc
   discarded <- c(0,cont_disc, lowQ_disc, NNNN_disc)
   filename <- paste0("example_",sprintf("%02d",i),"_summary.bin")
   f <- file(filename, "wb")
   writeBin(as.integer(filter),f)
   writeBin(as.integer(trimmed),f)
   writeBin(as.integer(discarded),f)
   writeBin(as.integer(ngood),f)
   writeBin(as.integer(nreads),f)
   close(f)

}
```
