## Supplementary material for "FastqPuri: high-performance preprocessing of RNA-seq data": filter_Sreport_example.html

Summary filtering report


### Summary filtering report

Running on version 1.0

#### Filter set up

|  | Applied | Method |
| --- | --- | --- |
| ADAPTERS | NO | NONE |
| CONTAMINATIONS | YES | TREE |
| LOW Q | YES | ENDSFRAC |
| N’s | YES | ENDS |

#### Statistics summary

|  | Nreads | Naccepted | %disc Ad | %cont | %disc lowQ | %disc N’s | %trim Ad | %trim N’s | %trim lowQ |
| --- | --- | --- | --- | --- | --- | --- | --- | --- | --- |
| example\_01 | 161171 | 131143 | 0 | 12.473 | 5.450 | 0.708 | 0 | 11.240 | 0.810 |
| example\_02 | 194182 | 165370 | 0 | 9.213 | 4.614 | 1.010 | 0 | 5.663 | 0.733 |
| example\_03 | 145192 | 124016 | 0 | 7.048 | 6.812 | 0.725 | 0 | 12.240 | 1.282 |
| example\_04 | 155844 | 123435 | 0 | 15.154 | 4.710 | 0.932 | 0 | 6.623 | 1.143 |
| example\_05 | 137726 | 116735 | 0 | 8.168 | 6.162 | 0.911 | 0 | 12.318 | 1.380 |
| example\_06 | 193116 | 160304 | 0 | 13.248 | 2.809 | 0.934 | 0 | 6.353 | 1.035 |
| example\_07 | 107152 | 73610 | 0 | 23.816 | 5.769 | 1.718 | 0 | 9.632 | 1.610 |
| example\_08 | 182981 | 145860 | 0 | 16.451 | 3.093 | 0.742 | 0 | 8.525 | 0.840 |
| example\_09 | 169124 | 143522 | 0 | 10.957 | 3.518 | 0.663 | 0 | 7.246 | 0.961 |
| example\_10 | 176085 | 154005 | 0 | 7.100 | 4.443 | 0.997 | 0 | 10.779 | 0.847 |
| example\_11 | 174368 | 147833 | 0 | 9.407 | 4.747 | 1.064 | 0 | 6.084 | 0.575 |
| example\_12 | 159828 | 133455 | 0 | 10.301 | 5.259 | 0.941 | 0 | 10.951 | 0.669 |
| example\_13 | 129096 | 94351 | 0 | 19.939 | 6.197 | 0.778 | 0 | 12.615 | 1.141 |
| example\_14 | 121365 | 94100 | 0 | 16.624 | 4.224 | 1.617 | 0 | 9.289 | 1.519 |
| example\_15 | 115981 | 87362 | 0 | 15.142 | 8.163 | 1.371 | 0 | 11.532 | 1.712 |
| example\_16 | 104454 | 77073 | 0 | 18.673 | 5.950 | 1.590 | 0 | 11.768 | 1.635 |
| example\_17 | 163332 | 130422 | 0 | 14.338 | 5.170 | 0.641 | 0 | 8.932 | 0.710 |
| example\_18 | 181867 | 153990 | 0 | 10.989 | 3.545 | 0.794 | 0 | 8.040 | 1.010 |
| example\_19 | 166298 | 132487 | 0 | 14.538 | 4.905 | 0.889 | 0 | 9.968 | 1.007 |
| example\_20 | 109288 | 71374 | 0 | 27.526 | 5.499 | 1.666 | 0 | 17.423 | 1.148 |
| example\_21 | 155584 | 117551 | 0 | 18.448 | 5.115 | 0.882 | 0 | 10.250 | 0.938 |
| example\_22 | 166093 | 131346 | 0 | 16.511 | 3.408 | 1.001 | 0 | 8.303 | 0.877 |
| example\_23 | 185029 | 154488 | 0 | 11.716 | 3.830 | 0.960 | 0 | 10.573 | 0.674 |
| example\_24 | 181964 | 147261 | 0 | 15.539 | 2.894 | 0.639 | 0 | 10.350 | 0.645 |
| example\_25 | 173692 | 137076 | 0 | 16.586 | 3.572 | 0.923 | 0 | 6.691 | 0.634 |
| example\_26 | 193440 | 160564 | 0 | 13.162 | 3.072 | 0.761 | 0 | 9.609 | 0.825 |
| example\_27 | 125846 | 86014 | 0 | 23.823 | 6.638 | 1.190 | 0 | 11.676 | 1.141 |
| example\_28 | 108117 | 73567 | 0 | 25.897 | 5.008 | 1.052 | 0 | 14.595 | 1.314 |
| example\_29 | 159934 | 130573 | 0 | 11.786 | 5.857 | 0.715 | 0 | 11.018 | 0.750 |
| example\_30 | 164678 | 144598 | 0 | 6.150 | 4.879 | 1.165 | 0 | 7.035 | 1.092 |
