## Supplementary material for "FastqPuri: high-performance preprocessing of RNA-seq data": create_fq.sh

```
#!/bin/bash


################################ FUNCTIONS #####################################
# Check --trimQ: add low quality reads (low quality 9 <-> 25)
# check also the low quality threshold
# 9 reads:
# Option ENDSFRAC (-p 5): 4 discarded, 4 trimmed, 1 left "as is"
function trimQ {
echo "@ read LOWQ TRIMQ:5:44
CAGTTCTTTGGTGTAGATGGAGCAGGACGGGATACCCATACCGTGACCCA
+position: -19377
99999IIIIIIIIIIIIIIIIIIIIIIIIIIIIIIIIIIIIIIII99999
@ read 108110 LOWQ TRIMQ:1:48
CAGATAAATGCGAATGAATGGGTAACACTGGAGCTTTTAATGGCTGTAAC
+position: -4142524
9IIIIIIIIIIIIIIIIIIIIIIIIIIIIIIIIIIIIIIIIIIIIIIII9
@ read 108111 LOWQ DISCARDED
AGCGTGACGGTGTAACGCCCGCCTTTTGATGACTGGGTTTCAAAGAAACG
+position: -3336785
99IIIIIIIIIIII9IIIIIIII9IIIIIIIII9IIIIIIIIIIIIII99
@ read 108112  LOWQ TRIM:4:49
ACCGGTAATGATACCCGCCAGTACACCCATGTTTATTTCTGGGTTGATGG
+position: -4548423
9999IIIIIIIIIIIIIIIIIIIIIIIIIIIIIIIIIIIIIIIIIIIIII
@ read 1081133 LOWQ TRIMQ:0:48
GTACATTGATGAGCAACATGGGGCTTGAACTGGCGCTGAAACAGTTAGGA
+position: -144740
IIIIIIIIIIIIIIIIIIIIIIIIIIIIIIIIIIIIIIIIIIIIIIIII9
@ read 108114 LOWQ DISCARDED
CTTAACCTGAACATTGAAGAGTTTAAAGGCGAGCAGTTACGCGTGACGGG
+position: 5075572
IIIIIIIIIIIIIIIIIIIII9999999IIIIIIIIIIIIIIIIIIIIII
@ read 108116 LOWQ DISCARDED
TCAGGGTGCAGCGGTAGCGCCAGTTTCCAGTCCATCGCCATTTTGTAGAC
+position: 1750468
IIIIIIIIIIII9III9IIIIII9IIIIIIIIIIIIIIIIIIIIIIIIII
@ read 108117 LOWQ UNCHANGED
ATTCACTGTAAGCGCGGTCTGGTATGGGGCGAGCTGGAATTTGTACTGCC
+position: 1091320
IIIIIIIIIIIIIIIIIIIIIII9IIIIIIIIIIIIIIIIIIIIIIIIII
@ read 108118 LOWQ DISCARDED
CAGCAAAATCATGCCGAGGAAGGCAATGCAAAGATAGTTGCCGAACGGAT
+position: -4651833
9IIIIIIIIIII9IIIIIIIIIII9IIIIIIII9IIIIIIIIIIIIIII9"  >> $1
}

# Check --trimN: add reads with N's, at the ends and in the middle
# 10 reads.
# Option STRIP: 7 trimmed, 3 discarded.
function trimN {
echo "@ read 1037  STRIP, ENDS -> TRIMN:2:47
NNTCGACACAAGAAAATGCGCCAATTTTGAGCCAGACCCCAGTTACGCNN
+position: -2044610
IIIIIIIIIIIIIIIIIIIIIIIIIIIIIIIIIIIIIIIIIIIIIIIIII
@ read 1038 STRIP -> TRIMN:17:49, ENDS -> UNCHANGED
AAAAAACGGTCGTGGTNGGCTACTGTTATTAAAGCGTTGGCTACAAAAAG
+position: 1361068
IIIIIIIIIIIIIIIIIIIIIIIIIIIIIIIIIIIIIIIIIIIIIIIIII
@ read 1039 STRIP -> DISCARDED N, ENDS -> UNCHANGED
CCAACGGACNATGGAAGTGCTNCGGCTGGNGTTTTTATCCTCCGGCATTC
+position: -4282223
IIIIIIIIIIIIIIIIIIIIIIIIIIIIIIIIIIIIIIIIIIIIIIIIII
@ read 1040 STRIP -> DISCARDED N, ENDS -> UNCHANGED
GCAAACGTATTTATGGCGACNNNNNNNCCGGTCATCAGGCCGATAAACGC
+position: 3171482
IIIIIIIIIIIIIIIIIIIIIIIIIIIIIIIIIIIIIIIIIIIIIIIIII
@ read 1041 STRIP, ENDS -> TRIMN:2:49
NNCATGCGCGTTCACTGCCCGCGATCCAGGATATTTTTTCCAGGGGGACC
+position: -86494
IIIIIIIIIIIIIIIIIIIIIIIIIIIIIIIIIIIIIIIIIIIIIIIIII
@ read 1042 STRIP, ENDS -> TRIMN:0:46
TGTCTGGCAACCTGATTCATCAGGCTTTGCGTAAGCGGAACGATTAANNN
+position: -3861809
IIIIIIIIIIIIIIIIIIIIIIIIIIIIIIIIIIIIIIIIIIIIIIIIII
@ read 1043 STRIP -> DISCARDED, ENDS -> TRIMN:0:48
AATCTGAAACGAAANCTGACAGCGNNCCCCGCTTCTGACAAAATAGGCGN
+position: -4685876
IIIIIIIIIIIIIIIIIIIIIIIIIIIIIIIIIIIIIIIIIIIIIIIIII
@ read 1044 STRIP -> TRIMN:4:46, ENDS -> TRIMN:2:46
NNANATGATCCTGGTCTGCAATAATCGTAAAGGGGCCGTCAGCGTGTNNN
+position: 1377584
IIIIIIIIIIIIIIIIIIIIIIIIIIIIIIIIIIIIIIIIIIIIIIIIII
@ read 1045 STRIP -> TRIMN:12:43, ENDS -> UNCHANGED
CGCCATNCCGANAATGATTGCTTTCGCCATATGGGTACGTCCGGNGATCC
+position: -2919586
IIIIIIIIIIIIIIIIIIIIIIIIIIIIIIIIIIIIIIIIIIIIIIIIII
@ read 1046 STRIP -> TRIMN:15:49, ENDS -> UNCHANGED
CCCCTCTGCANNNCNCAATAATTTCATCGAGGAATCCGTTGCTTCGGGCT
+position: -3548964
IIIIIIIIIIIIIIIIIIIIIIIIIIIIIIIIIIIIIIIIIIIIIIIIII
@ read 1046 STRIP -> TRIMN:15:49, ENDS -> UNCHANGED
CCCCTCTGCANNNCNCAATAATTTCATCGAGGAATCCGTTGCTTCGGGCT
+position: -3548964
IIIIIIIIIIIIIIIIIIIIIIIIIIIIIIIIIIIIIIIIIIIIIIIIII
@ read 1046 STRIP, ENDS -> TRIMX:4:46
CCNNCTCTGCACAATCAATAATTTCATCGAGGAATCCGTTGCTTCGGNNT
+position: -3548964
55IIIIIIIIIIIIIIIIIIIIIIIIIIIIIIIIIIIIIIIIIIIIIII5" >> $1
}

#################################### CODE ######################################

simReads=../fa_fq_files/simReads.pl
out_fq=../fa_fq_files/EColi_rRNA.fq

# Create 2e5 fq reads of length 50  from EColi genome with NO errors
$simReads ../fa_fq_files/EColi_genome.fa 0.02 0 50  0  > ${out_fq}

# Create 5e4 fq reads of length 50  from rRNA_modified with NO errors
# rRNA_modified is the sequence rRNA_CRUnit with the lines containing
# N's removed
$simReads ../fa_fq_files/rRNA_modified.fa 0.005 0 50 0  >>  ${out_fq}

# Create reads with low quality issues and with N's
trimQ ${out_fq}
trimN ${out_fq}

# Zip the output
gzip ${out_fq}
```
