## Supplementary material for "FastqPuri: high-performance preprocessing of RNA-seq data": README

################################################################################
#  trimFilter examples:                                                        #
################################################################################
#                                                                              # 
# The examples work in the  following way:                                     #
#                                                                              #
# 1. See folder `fa_fq_files`. The file `EColi_rRNA.fq.gz` was created with    #
#    `create_fq.sh` and contains:                                              # 
#    * 2e4 reads of length 50 from `EColi_genome.fa` with NO errors.           #
#    * 5e3 reads of length 50 from `rRNA_modified.fa` with NO errrors          #
#      (rRNA contaminations).                                                  #
#    * Artificially generated reads with low quality score (see `create_fq.sh`)#
#    * Artificially generated reads with Ns (see `create_fq.sh`).              #
# 2. `run_example_TREE.sh`: the code was tested with flags:                    # 
#    `../../bin/trimFilter -l 50 --ifq ../fa_fq_files/EColi_rRNA.fq.gz\        #
#     --method TREE --ifa ../fa_fq_files/rRNA_modified.fa:0.9:50 \             #
#     --trimQ ENDSFRAC --trimN STRIP -o tree`                                  #
#    i.e., we check for contaminations from rRNA, trim reads with lowQ at      #
#    the ends and less than 5% in the remaining part, and strip reads          #
#    containing N's.                                                           # 
#    The output should coincide with the files `example_TREE*`                 #
# 3. `run_example_BLOOM.sh`:                                                   # 
#    a) a bloom filter is generated for `rRNA_modified.fa` with FPR = 0.0075   #
#       and `kmersize=25`. The output should coincide with `rRNA_example.bf*`. # 
#    b) trimFilter is run like in 2. but passing a bloom filter to look for    # 
#       contaminations with `score=0.4`.                                       #
# 4. With this set up, it is possible to run further customized tests.         #
# 5. See the folder `adapters` for examples on adapter contaminations.         #
#                                                                              # 
# NOTE: `rRNA_modified.fa` is the `rRNA_CRUnit.fa` sequence, where we have     #
#        removed the lines containing N's for testing purposes.                #
#                                                                              # 
################################################################################
################################################################################
#  Sreport example:                                                            #
################################################################################
#                                                                              #
# In folder `./bin_files`, 30 fake `./trimFilter` binary output files were     #
# generated (with the `R` script `create_fake_bins.R`). An html output was     #
# created out of them (`./bin_files/filter_Sreport_example.html`). It can      # 
# re reproduced if you run:                                                    #
#   ../../bin/Sreport -i ./bin_files/ -o ./bin_files/filter_Sreport_new -t F    #
#                                                                              #
################################################################################
