## Supplementary material for "FastqPuri: high-performance preprocessing of RNA-seq data": run_example_BLOOM.sh

```
#!/bin/bash

# This script creates a bloom filter from rRNA (FPR 0.0075), and runs trimFilter 
# as in the example trimFilter_Sreport/ but using the bloom filter with a score
# of 0.4 instead of the tree.

makeBloom=../../bin/makeBloom 
trimFilter=../../bin/trimFilter
sequence=../fa_fq_files/rRNA_modified.fa
fastq=../fa_fq_files/EColi_rRNA.fq.gz

# STEP1: generate the bloom Filter
$makeBloom -f $sequence -o rRNA -p 0.0075

# STEP2: run bloom Filter on EColu+rRNA.gq.gz data
$trimFilter -f $fastq -l 50 --method BLOOM --idx rRNA.bf:0.4\
           --trimQ ENDSFRAC --trimN ENDS -o bloom 
```
