## Supplementary material for "FastqPuri: high-performance preprocessing of RNA-seq data": run_example_TREE.sh

```
#!/bin/bash 

# This script calls the executable and saves the output 
fastq=../fa_fq_files/EColi_rRNA.fq.gz
fasta=../fa_fq_files/rRNA_modified.fa
trimFilter=../../bin/trimFilter

# Run trimFilter using a tree
$trimFilter -l 50 --ifq "${fastq}" --ifa "$fasta":0.9:50 \
 --method TREE --trimQ ENDSFRAC --trimN STRIP -o tree
```
