## Supplementary material for "FastqPuri: high-performance preprocessing of RNA-seq data": create_fq.sh

```
# This script calls fq_gen.sh to generate the fq files that serve as 
# probe for trimFilterDS with the --adapter option

# Input path and adapter files
pathI="../../fa_fq_files/"
Ad1=$pathI"ad_read1.fa"
Ad2=$pathI"ad_read2.fa"
fq_gen="./fq_gen.sh"

# Create DS fastq files to check the adapters option. EVEN read length 
fq1=$pathI"evenL70_r1.fq"
fq2=$pathI"evenL70_r2.fq"
f1fw="AACCCAGCTACGATCTAGCGAGCGGCGAGGCATGCGCTGATGCAGCCCGATCGCAGCGTCGCCCATATGT"
f2fw="ACAGTTTATGCATCATCTTCTGCGTTGCTCGCTAGCTCAGTCGATTCGATGGCATCTATCGAATTCCCGT"
${fq_gen} $fq1 $fq2 $Ad1 $Ad2 $f1fw $f2fw

# Create DS fastq files to check the adapters option. ODD read length 
fq1=$pathI"oddL71_r1.fq"
fq2=$pathI"oddL71_r2.fq"
f1fw="AACCCAGGCTACGATCTAGCGAGCGGCGAGGCATGCGCTGATGCAGCCCGATCGCAGCGTCGCCCATATGT"
f2fw="ACAGTTGTATGCATCATCTTCTGCGTTGCTCGCTAGCTCAGTCGATTCGATGGCATCTATCGAATTCCCGT"
${fq_gen} $fq1 $fq2 $Ad1 $Ad2 $f1fw $f2fw
```
