## Supplementary material for "FastqPuri: high-performance preprocessing of RNA-seq data": fq_gen.sh

```
#!/bin/bash

# In this script, we generate artificial data to test trimFilterDS with adapters

# Obtains reverse complement of a sequence 
# - $1: sequence
function revcomp {
   echo $1 | rev | tr ACGT TGCA
}
 
# Obtains reverse of a sequence 
# - $1: sequence
function reverse {
   echo $1 | rev 
}

# Cut string from position $2 to position $3 (starting to count in 1)
# $1 string
# $2 start cutting at this position
# $3 end cutting at this position
function cutstr {
   echo $1 | cut -c$2-$3
}

function adsubst {
  tmp=$1
  for i in ${@:2}
  do
     j=$(($i-1))
     if [[ ${1:$j:1} != "A" ]] ; then 
        tmp=`echo $tmp | sed s/./A/$i`
     else 
        tmp=`echo $tmp | sed s/./T/$i`
     fi
  done
  echo $tmp
}

#Create a fastq read
# $1  first line 
# $2  second line
# $3  third line 
# $4  fourth line
function fqread {
   eval l1="$1"
   eval l2="$2"
   eval l3="$3"
   eval l4="$4"
   printf "$l1\n$l2\n$l3\n$l4\n" 
}

#Usage 
function my_usage {
  echo "Usage: ./fq_gen.sh INPUT_R1.fq INPUT_R2.fq  AD1.fa AD2.fa seq1 seq2"
}


# Check the number of arguments
if [ $# -ne 6 ]; then 
   echo "Illegal number of parameters"
   my_usage
   exit
fi

# Reads and adapters. 
file1=$1; file2=$2
Ad1file=$3; Ad2file=$4
f1fw=$5; f2fw=$6

Ad1fw=`tail -n +2 $Ad1file`
Ad2fw=`tail -n +2 $Ad2file`
N=`echo ${#f1fw}`     # Read length
Nad1=`echo ${#Ad1fw}` # ad1 length
Nad2=`echo ${#Ad2fw}` # ad2 length

# Writing info to  stdout
echo "Creating DS files: $file1, $file2"
echo "Reading Adapter files: $Ad1file, $Ad2file"
echo "Creating reads of length: $N"

# Reverse complements and reverse sequences 
f1bw=`revcomp $f1fw`
f2bw=`revcomp $f2fw`
Ad1bw=`revcomp $Ad1fw`
Ad2bw=`revcomp $Ad2fw`


# Remove files if existing and open them
if [ -f $file1 ] ; then rm $file1; fi
if [ -f $file2 ] ; then rm $file2; fi
touch $file1
touch $file2

# Initialize line3 and line4 since they will kept the same for the whole file
l3="+"
l4=`for i in \`seq 1 $N\`; do printf "I"; done`

# Read 1, CASE 1 no overlap, UNCHANGED
l1="@READ01, CASE 1, no overlap  UNCHANGED"
l2_r1=$f1fw
l2_r2=$f2fw
fqread "\${l1}" "\${l2_r1}" "\${l3}" "\${l4}" >> $file1
fqread "\${l1}" "\${l2_r2}" "\${l3}" "\${l4}" >> $file2

# Read 2, CASE 2 overlap in read but no contamination, UNCHANGED.
N1=50
N2=$(($N - $N1))
l1="@READ02, CASE 2, overlap in read, no contamination, UNCHANGED"
l2_r1=$f1fw
l2_r2=`cutstr $f2fw 1 $N1``cutstr $f1bw 1 $N2`
fqread "\${l1}" "\${l2_r1}" "\${l3}" "\${l4}" >> $file1
fqread "\${l1}" "\${l2_r2}" "\${l3}" "\${l4}" >> $file2

# Read 3, CASE 3 overlap in read and contamination, TRIM
N1=50
N2=$(($N - $N1)) 
l1="@READ03, CASE 3, overlap in read, contamination, TRIMA:0:$(($N1-1))"
l2_r1=`cutstr $f1fw 1 $N1``cutstr $Ad2bw 1 $N2`
l2_r2=`cutstr $f1bw $(($N2+1)) $N``cutstr $Ad1bw 1 $N2`
fqread "\${l1}" "\${l2_r1}" "\${l3}" "\${l4}" >> $file1
fqread "\${l1}" "\${l2_r2}" "\${l3}" "\${l4}" >> $file2

# Read 4, CASE 4 full overlap + garbage, DISCARDED
N1=29
L1=$(($N-$N1-$Nad2-1))
L2=$(($N-$N1-$Nad1-1))
l1="@READ04, CASE 4, full overlap + garbage, contamination, TRIMA:0:$(($N1-1))"
l2_r1=`cutstr $f1fw 1 $N1`$Ad2bw`cutstr $f1fw 31 $((31+$L1))`
l2_r2=`cutstr $f1bw $(($N - $N1 + 1)) $N`$Ad1bw`cutstr $f2fw 1 $((1+$L2))`
fqread "\${l1}" "\${l2_r1}" "\${l3}" "\${l4}" >> $file1
fqread "\${l1}" "\${l2_r2}" "\${l3}" "\${l4}" >> $file2

# Read 5, CASE 4 full overlap + garbage, too short, DISCARDED
N1=19
L1=$(($N-$N1-$Nad2-1))
L2=$(($N-$N1-$Nad1-1))
l1="@READ05, CASE 4, full overlap + garbage, contamination, DISCARDED"
l2_r1=`cutstr $f1fw 1 $N1`$Ad2bw`cutstr $f1fw 21 $((21 + $L1))`
l2_r2=`cutstr $f1bw $(($N - $N1 + 1)) $N`$Ad1bw`cutstr $f2fw 1 $((1 + $L2))`
fqread "\${l1}" "\${l2_r1}" "\${l3}" "\${l4}" >> $file1
fqread "\${l1}" "\${l2_r2}" "\${l3}" "\${l4}" >> $file2

# Read 6, CASE 5 full overlap + garbage, DISCARDED
N1=$(($N -$Nad2))
N2=$(($N -$Nad1))
l1="@READ06, CASE 5, No read,  DISCARDED"
l2_r1=$Ad2bw`cutstr $f1fw 1 $N1`
l2_r2=$Ad1bw`cutstr $f2fw 1 $N2`
fqread "\${l1}" "\${l2_r1}" "\${l3}" "\${l4}" >> $file1
fqread "\${l1}" "\${l2_r2}" "\${l3}" "\${l4}" >> $file2

# Read 7, CASE 3 overlap, contamination, with 1 error in both reads, TRIM 
N1=50
N2=$(($N - $N1)) 
l1="@READ07, CASE 3, overlap in read, contamination, with 1 error in both reads TRIMA:0:$(($N1-1))"
l2_r1=`cutstr $f1fw 1 $N1``cutstr $Ad2bw 1 $N2`
l2_r1=`adsubst $l2_r1 64`
l2_r2=`cutstr $f1bw $(($N2+1)) $N``cutstr $Ad1bw 1 $N2`
l2_r2=`adsubst $l2_r2 67`
fqread "\${l1}" "\${l2_r1}" "\${l3}" "\${l4}" >> $file1
fqread "\${l1}" "\${l2_r2}" "\${l3}" "\${l4}" >> $file2

# Read 8, CASE 3 overlap, contamination, with 2 errors in read2 (seed), TRIM
N1=50
N2=$(($N - $N1)) 
l1="@READ08, CASE 3, overlap in read, contamination,  with 2 error in read2 (seed) TRIMA:0:$(($N1-1))"
l2_r1=`cutstr $f1fw 1 $N1``cutstr $Ad2bw 1 $N2`
l2_r2=`cutstr $f1bw $(($N2+1)) $N``cutstr $Ad1bw 1 $N2`
l2_r2=`adsubst $l2_r2 60 64`
fqread "\${l1}" "\${l2_r1}" "\${l3}" "\${l4}" >> $file1
fqread "\${l1}" "\${l2_r2}" "\${l3}" "\${l4}" >> $file2

# Read 9, CASE 3 overlap, contamination, with 3 errors in read2 (seed), UNCHANGED
N1=50
N2=$(($N - $N1)) 
l1="@READ09, CASE 3, overlap in read, contamination, UNCHANGED"
l2_r1=`cutstr $f1fw 1 $N1``cutstr $Ad2bw 1 $N2`
l2_r2=`cutstr $f1bw $(($N2+1)) $N``cutstr $Ad1bw 1 $N2`
l2_r2=`adsubst $l2_r2 64 66 69`
fqread "\${l1}" "\${l2_r1}" "\${l3}" "\${l4}" >> $file1
fqread "\${l1}" "\${l2_r2}" "\${l3}" "\${l4}" >> $file2

# Read 10, CASE 4 full overlap + garbage 1 error both reads, TRIM
N1=29
L1=$(($N-$N1-$Nad2-1))
L2=$(($N-$N1-$Nad1-1))
l1="@READ10, CASE 4, full overlap + garbage, contamination, 1 error in read2 (seed) TRIMA:0:$(($N1-1))"
l2_r1=`cutstr $f1fw 1 $N1`$Ad2bw`cutstr $f1fw 31 $((31+$L1))`
l2_r1=`adsubst $l2_r1 56`
l2_r2=`cutstr $f1bw $(($N - $N1 + 1)) $N`$Ad1bw`cutstr $f2fw 1 $((1+$L2))`
l2_r2=`adsubst $l2_r2 32`
fqread "\${l1}" "\${l2_r1}" "\${l3}" "\${l4}" >> $file1
fqread "\${l1}" "\${l2_r2}" "\${l3}" "\${l4}" >> $file2

# Read 11, CASE 4 full overlap + garbage 2 errors in read2 (seed), TRIM
N1=29
L1=$(($N-$N1-$Nad2-1))
L2=$(($N-$N1-$Nad1-1))
l1="@READ11, CASE 4, full overlap + garbage, contamination, 2 errors in read2 (seed) TRIMA:0:$(($N1-1))"
l2_r1=`cutstr $f1fw 1 $N1`$Ad2bw`cutstr $f1fw 31 $((31+$L1))`
l2_r2=`cutstr $f1bw $(($N - $N1 + 1)) $N`$Ad1bw`cutstr $f2fw 1 $((1+$L2))`
l2_r2=`adsubst $l2_r2 56 55`
fqread "\${l1}" "\${l2_r1}" "\${l3}" "\${l4}" >> $file1
fqread "\${l1}" "\${l2_r2}" "\${l3}" "\${l4}" >> $file2

# Read 12, CASE 4 full overlap + garbage 3 errors in read2(seed), UNCHANGED
N1=29
L1=$(($N-$N1-$Nad2-1))
L2=$(($N-$N1-$Nad1-1))
l1="@READ12, CASE 4, full overlap + garbage, contamination, 3 errors in read2 (seed) UNCHANGED"
l2_r1=`cutstr $f1fw 1 $N1`$Ad2bw`cutstr $f1fw 31 $((31+$L1))`
l2_r2=`cutstr $f1bw $(($N - $N1 + 1)) $N`$Ad1bw`cutstr $f2fw 1 $((1+$L2))`
l2_r2=`adsubst $l2_r2   56 58 57`
fqread "\${l1}" "\${l2_r1}" "\${l3}" "\${l4}" >> $file1
fqread "\${l1}" "\${l2_r2}" "\${l3}" "\${l4}" >> $file2

# Read 13, CASE 4 full overlap + garbage 1 error in read2 (seed), DISCARDED
N1=19
L1=$(($N-$N1-$Nad2-1))
L2=$(($N-$N1-$Nad1-1))
l1="@READ13, CASE 4, full overlap + garbage, contamination, 1 error in read2 (seed) DISCARDED"
l2_r1=`cutstr $f1fw 1 $N1`$Ad2bw`cutstr $f1fw 21 $((21 + $L1))`
l2_r2=`cutstr $f1bw $(($N - $N1 + 1)) $N`$Ad1bw`cutstr $f2fw 1 $((1 + $L2))`
l2_r2=`adsubst $l2_r2 46`
fqread "\${l1}" "\${l2_r1}" "\${l3}" "\${l4}" >> $file1
fqread "\${l1}" "\${l2_r2}" "\${l3}" "\${l4}" >> $file2

# Read 14, CASE 4 full overlap + garbage 2 errors in read2(seed), DISCARDED
N1=19
L1=$(($N-$N1-$Nad2-1))
L2=$(($N-$N1-$Nad1-1))
l1="@READ14, CASE 4, full overlap + garbage, contamination, 2 errors in read2 (seed) DISCARDED"
l2_r1=`cutstr $f1fw 1 $N1`$Ad2bw`cutstr $f1fw 21 $((21 + $L1))`
l2_r2=`cutstr $f1bw $(($N - $N1 + 1)) $N`$Ad1bw`cutstr $f2fw 1 $((1 + $L2))`
l2_r2=`adsubst $l2_r2 46 47`
fqread "\${l1}" "\${l2_r1}" "\${l3}" "\${l4}" >> $file1
fqread "\${l1}" "\${l2_r2}" "\${l3}" "\${l4}" >> $file2

# Read 15, CASE 4 full overlap + garbage 3 errors in read2(seed) UNCHANGED
N1=19
L1=$(($N-$N1-$Nad2-1))
L2=$(($N-$N1-$Nad1-1))
l1="@READ15, CASE 4, full overlap + garbage, contamination, 3 errors en read2 (seed) UNCHANGED"
l2_r1=`cutstr $f1fw 1 $N1`$Ad2bw`cutstr $f1fw 21 $((21 + $L1))`
l2_r2=`cutstr $f1bw $(($N - $N1 + 1)) $N`$Ad1bw`cutstr $f2fw 1 $((1 + $L2))`
l2_r2=`adsubst $l2_r2 46 47 48 `
fqread "\${l1}" "\${l2_r1}" "\${l3}" "\${l4}" >> $file1
fqread "\${l1}" "\${l2_r2}" "\${l3}" "\${l4}" >> $file2

# Read 16, CASE 5 full overlap + garbage, 1 error in read2(seed), DISCARDED
N1=$(($N -$Nad2))
N2=$(($N -$Nad1))
l1="@READ16, CASE 5, No read, 1 error in read 2 (seed) DISCARDED"
l2_r1=$Ad2bw`cutstr $f1fw 1 $N1`
l2_r2=$Ad1bw`cutstr $f2fw 1 $N2`
l2_r2=`adsubst $l2_r2 28 `
fqread "\${l1}" "\${l2_r1}" "\${l3}" "\${l4}" >> $file1
fqread "\${l1}" "\${l2_r2}" "\${l3}" "\${l4}" >> $file2

# Read 17, CASE 5 full overlap + garbage, 2 errors in read2(seed), DISCARDED
N1=$(($N -$Nad2))
N2=$(($N -$Nad1))
l1="@READ17, CASE 5, No read, 2 errors in read 2 (seed) DISCARDED"
l2_r1=$Ad2bw`cutstr $f1fw 1 $N1`
l2_r2=$Ad1bw`cutstr $f2fw 1 $N2`
l2_r2=`adsubst $l2_r2 28 29`
fqread "\${l1}" "\${l2_r1}" "\${l3}" "\${l4}" >> $file1
fqread "\${l1}" "\${l2_r2}" "\${l3}" "\${l4}" >> $file2

# Read 18, CASE 5 full overlap + garbage, 3 errors in read2(seed), UNCHANGED
N1=$(($N -$Nad2))
N2=$(($N -$Nad1))
l1="@READ18, CASE 5, No read, 3 errors in read 2 (seed) UNCHANGED"
l2_r1=$Ad2bw`cutstr $f1fw 1 $N1`
l2_r2=$Ad1bw`cutstr $f2fw 5 $(($N2+4))`

l2_r2=`adsubst $l2_r2 25 26 27`
fqread "\${l1}" "\${l2_r1}" "\${l3}" "\${l4}" >> $file1
fqread "\${l1}" "\${l2_r2}" "\${l3}" "\${l4}" >> $file2
```
