## Supplementary material for "FastqPuri: high-performance preprocessing of RNA-seq data": README

################################################################################
#  trimFilterDS examples (adapters contamination):                             #
################################################################################
#                                                                              # 
#  RUN: ./run_examples.sh and compare the created files with the existing files#
#       in this directory.                                                     #
#                                                                              # 
# The script `run_examples.sh` runs trimFilterDS with the option `--adapter`   # 
# for 2 different `fq` files, once for an even read length and once for an     #
# odd read length. These reads try to cover all(most) possible ways in which   #
# adapters contaminations can be present in double stranded data:              #
#                                                                              # 
# - `trimFilterDS` is run with the following input for the `--adapter` option: #
#   * Adapter read 1: ../../fa_fq_files/ad_read1.fa                            #
#   * Adapter read 2: ../../fa_fq_files/ad_read2.fa                            #
#   * mismatches: 2 (for the SEED)                                             #
#   * score: 20                                                                #
#                                                                              # 
# - Input files can be generated with `create_fq.sh`, a script that calls      # 
#   `gen_fq.sh` and creates the following files:                               #
#   * ../../fa_fq_files/evenL70_r1.fq                                          #
#   * ../../fa_fq_files/evenL70_r2.fq                                          #
#   * ../../fa_fq_files/oddL71_r1.fq                                           #
#   * ../../fa_fq_files/oddL71_r2.fq                                           #
#                                                                              #
# - 18 reads are created with the following characteristics:                   #
#   * 01, CASE 1 no overlap, UNCHANGED                                         # 
#   * 02, CASE 2 overlap in read but no contamination, UNCHANGED               # 
#   * 03, CASE 3 overlap in read and contamination, TRIM                       #
#   * 04, CASE 4 full overlap + garbage, DISCARDED                             #
#   * 05, CASE 4 full overlap + garbage, too short, DISCARDED                  #
#   * 06, CASE 5 full overlap + garbage, DISCARDED                             #
#   * 07, CASE 3 overlap, contamination, 1 error in both reads, TRIM           #
#   * 08, CASE 3 overlap, contamination, 2 errors in read2 (seed), TRIM        #
#   * 09, CASE 3 overlap, contamination, 3 errors in read2 (seed), UNCHANGED   #
#   * 10, CASE 4 full overlap + garbage, 1 error both reads, TRIM              #
#   * 11, CASE 4 full overlap + garbage, 2 errors in read2 (seed), TRIM        #
#   * 12, CASE 4 full overlap + garbage, 3 errors in read2(seed), UNCHANGED    #
#   * 13, CASE 4 full overlap + garbage, 1 error in read2 (seed), DISCARDED    #
#   * 14, CASE 4 full overlap + garbage, 2 errors in read2(seed), DISCARDED    #
#   * 15, CASE 4 full overlap + garbage, 3 errors in read2(seed) UNCHANGED     #
#   * 16, CASE 5 full overlap + garbage, 1 error in read2(seed), DISCARDED     #
#   * 17, CASE 5 full overlap + garbage, 2 errors in read2(seed), DISCARDED    #
#   * 18, CASE 5 full overlap + garbage, 3 errors in read2(seed), UNCHANGED    #
#                                                                              # 
################################################################################
