## Supplementary material for "FastqPuri: high-performance preprocessing of RNA-seq data": run_examples.sh

```
# This script calls the executable and saves the output 
pathI="../../fa_fq_files/"
ad1=$pathI"ad_read1.fa"
ad2=$pathI"ad_read2.fa"
trimFilterDS="../../../bin/trimFilterDS"

# EVEN 
fq1=$pathI"evenL70_r1.fq"
fq2=$pathI"evenL70_r2.fq"
$trimFilterDS -l 70 --ifq "$fq1:$fq2" --adapter $ad1:$ad2:2:20 -o even

# ODD 
fq1=$pathI"oddL71_r1.fq"
fq2=$pathI"oddL71_r2.fq"
$trimFilterDS -l 71 --ifq "$fq1:$fq2" --adapter $ad1:$ad2:2:20 -o odd
```
