## Supplementary material for "FastqPuri: high-performance preprocessing of RNA-seq data": DS_Sreport_example.html

Summary filtering report, DS data


### Summary filtering report, DS data

Running on version 1.0

#### Filter set up

|  | Applied | Method |
| --- | --- | --- |
| ADAPTERS | NO | NONE |
| CONTAMINATIONS | YES | TREE |
| LOW Q | YES | ENDSFRAC |
| N’s | YES | ENDS |

#### Statistics summary

|  | Nreads | Naccepted | %disc Ad | %cont | %disc lowQ | %disc N’s | %trim Ad | %trim1 N’s | %trim1 lowQ | %trim2 N’s | %trim2 lowQ |
| --- | --- | --- | --- | --- | --- | --- | --- | --- | --- | --- | --- |
| example\_01 | 121652 | 90907 | 0 | 19.566 | 4.575 | 1.133 | 0 | 10.908 | 1.054 | 10.533 | 1.356 |
| example\_02 | 179044 | 146789 | 0 | 12.349 | 4.956 | 0.710 | 0 | 10.025 | 0.692 | 7.588 | 1.104 |
| example\_03 | 111925 | 83034 | 0 | 16.482 | 8.289 | 1.042 | 0 | 9.188 | 1.591 | 15.552 | 1.396 |
| example\_04 | 160478 | 122173 | 0 | 17.915 | 5.185 | 0.770 | 0 | 9.679 | 0.986 | 12.432 | 1.015 |
| example\_05 | 187950 | 161029 | 0 | 10.330 | 3.301 | 0.692 | 0 | 6.536 | 0.847 | 5.449 | 0.659 |
| example\_06 | 104451 | 83360 | 0 | 9.179 | 9.164 | 1.849 | 0 | 18.503 | 1.415 | 16.151 | 1.761 |
| example\_07 | 140241 | 118681 | 0 | 8.386 | 5.710 | 1.278 | 0 | 12.071 | 0.913 | 14.248 | 1.209 |
| example\_08 | 137459 | 103630 | 0 | 17.425 | 5.842 | 1.343 | 0 | 10.750 | 0.750 | 10.649 | 1.409 |
| example\_09 | 145631 | 121555 | 0 | 10.121 | 5.112 | 1.299 | 0 | 11.305 | 0.903 | 12.688 | 1.306 |
| example\_10 | 124099 | 90850 | 0 | 20.297 | 5.200 | 1.296 | 0 | 11.849 | 1.151 | 10.145 | 1.446 |
| example\_11 | 118915 | 91525 | 0 | 16.826 | 4.966 | 1.241 | 0 | 16.324 | 1.360 | 15.126 | 1.281 |
| example\_12 | 174284 | 140579 | 0 | 14.185 | 4.566 | 0.588 | 0 | 9.026 | 0.722 | 7.695 | 1.113 |
| example\_13 | 129475 | 95100 | 0 | 19.550 | 5.501 | 1.498 | 0 | 8.717 | 1.216 | 11.970 | 1.240 |
| example\_14 | 147033 | 122598 | 0 | 10.945 | 4.828 | 0.845 | 0 | 10.784 | 0.738 | 9.398 | 0.909 |
| example\_15 | 138450 | 105065 | 0 | 19.393 | 3.965 | 0.755 | 0 | 13.112 | 1.186 | 11.246 | 0.977 |
| example\_16 | 128305 | 95995 | 0 | 17.904 | 5.788 | 1.490 | 0 | 15.474 | 1.533 | 9.450 | 1.433 |
| example\_17 | 149742 | 120309 | 0 | 15.317 | 3.650 | 0.689 | 0 | 7.363 | 0.852 | 7.606 | 0.874 |
| example\_18 | 158509 | 121584 | 0 | 16.192 | 5.861 | 1.242 | 0 | 6.581 | 1.066 | 11.339 | 1.038 |
| example\_19 | 190205 | 152170 | 0 | 15.994 | 3.382 | 0.621 | 0 | 7.514 | 0.679 | 9.889 | 0.816 |
| example\_20 | 113068 | 77270 | 0 | 23.116 | 7.075 | 1.469 | 0 | 14.162 | 1.205 | 10.417 | 1.099 |
| example\_21 | 108813 | 81213 | 0 | 14.939 | 9.091 | 1.334 | 0 | 9.729 | 1.372 | 10.793 | 1.593 |
| example\_22 | 192724 | 166745 | 0 | 9.691 | 2.842 | 0.946 | 0 | 5.915 | 0.560 | 10.178 | 0.551 |
| example\_23 | 160483 | 139741 | 0 | 6.075 | 5.766 | 1.084 | 0 | 8.258 | 0.990 | 10.481 | 0.999 |
| example\_24 | 196258 | 158458 | 0 | 13.280 | 5.010 | 0.970 | 0 | 8.900 | 0.730 | 9.471 | 0.921 |
| example\_25 | 110934 | 80257 | 0 | 21.053 | 4.980 | 1.621 | 0 | 11.510 | 1.154 | 15.834 | 1.467 |
| example\_26 | 157759 | 134488 | 0 | 9.070 | 4.842 | 0.839 | 0 | 9.930 | 0.850 | 9.791 | 1.173 |
| example\_27 | 183640 | 157752 | 0 | 10.731 | 2.780 | 0.586 | 0 | 8.316 | 0.882 | 6.130 | 0.850 |
| example\_28 | 106286 | 66759 | 0 | 29.796 | 5.592 | 1.802 | 0 | 18.350 | 1.507 | 14.883 | 1.584 |
| example\_29 | 163284 | 133461 | 0 | 11.659 | 5.887 | 0.719 | 0 | 10.979 | 0.791 | 6.989 | 0.978 |
| example\_30 | 147604 | 119370 | 0 | 12.655 | 5.213 | 1.260 | 0 | 12.967 | 1.211 | 11.441 | 1.058 |
