## Supplementary material for "FastqPuri: high-performance preprocessing of RNA-seq data": create_fq.sh

```
#!/bin/bash

################################ FUNCTIONS #####################################

# Check --trimQ: add low quality reads (low quality 9 <-> 25)
# check also the low quality threshold. In the other read 
# nothing really happens (second argument)
# 9 reads: 
# Option ENDSFRAC (-p 5): 4 discarded, 4 trimmed, 1 left "as is"
function trimQ {
echo "@ read LOWQ TRIMQ:5:44
CAGTTCTTTGGTGTAGATGGAGCAGGACGGGATACCCATACCGTGACCCA
+position: -19377
99999IIIIIIIIIIIIIIIIIIIIIIIIIIIIIIIIIIIIIIII99999
@ read 108110 LOWQ TRIMQ:1:48
CAGATAAATGCGAATGAATGGGTAACACTGGAGCTTTTAATGGCTGTAAC
+position: -4142524
9IIIIIIIIIIIIIIIIIIIIIIIIIIIIIIIIIIIIIIIIIIIIIIII9
@ read 108111 LOWQ DISCARDED
AGCGTGACGGTGTAACGCCCGCCTTTTGATGACTGGGTTTCAAAGAAACG
+position: -3336785
99IIIIIIIIIIII9IIIIIIII9IIIIIIIII9IIIIIIIIIIIIII99
@ read 108112  LOWQ TRIM:4:49
ACCGGTAATGATACCCGCCAGTACACCCATGTTTATTTCTGGGTTGATGG
+position: -4548423
9999IIIIIIIIIIIIIIIIIIIIIIIIIIIIIIIIIIIIIIIIIIIIII
@ read 1081133 LOWQ TRIMQ:0:48
GTACATTGATGAGCAACATGGGGCTTGAACTGGCGCTGAAACAGTTAGGA
+position: -144740
IIIIIIIIIIIIIIIIIIIIIIIIIIIIIIIIIIIIIIIIIIIIIIIII9
@ read 108114 LOWQ DISCARDED 
CTTAACCTGAACATTGAAGAGTTTAAAGGCGAGCAGTTACGCGTGACGGG
+position: 5075572
IIIIIIIIIIIIIIIIIIIII9999999IIIIIIIIIIIIIIIIIIIIII
@ read 108116 LOWQ DISCARDED
TCAGGGTGCAGCGGTAGCGCCAGTTTCCAGTCCATCGCCATTTTGTAGAC
+position: 1750468
IIIIIIIIIIII9III9IIIIII9IIIIIIIIIIIIIIIIIIIIIIIIII
@ read 108117 LOWQ UNCHANGED
ATTCACTGTAAGCGCGGTCTGGTATGGGGCGAGCTGGAATTTGTACTGCC
+position: 1091320
IIIIIIIIIIIIIIIIIIIIIII9IIIIIIIIIIIIIIIIIIIIIIIIII
@ read 108118 LOWQ DISCARDED
CAGCAAAATCATGCCGAGGAAGGCAATGCAAAGATAGTTGCCGAACGGAT
+position: -4651833
9IIIIIIIIIII9IIIIIIIIIII9IIIIIIII9IIIIIIIIIIIIIII9" >> $1 

echo "@ read LOWQ (the other read) UNCHANGED
CAGTTCTTTGGTGTAGATGGAGCAGGACGGGATACCCATACCGTGACCCA
+position: -19377
IIIIIIIIIIIIIIIIIIIIIIIIIIIIIIIIIIIIIIIIIIIIIIIIII
@ read 108110 LOWQ (the other read) UNCHANGED
CAGATAAATGCGAATGAATGGGTAACACTGGAGCTTTTAATGGCTGTAAC
+position: -4142524
IIIIIIIIIIIIIIIIIIIIIIIIIIIIIIIIIIIIIIIIIIIIIIIIII
@ read 108111 LOWQ DISCARDED (slave)
AGCGTGACGGTGTAACGCCCGCCTTTTGATGACTGGGTTTCAAAGAAACG
+position: -3336785
IIIIIIIIIIIIIIIIIIIIIIIIIIIIIIIIIIIIIIIIIIIIIIIIII
@ read 108112  LOWQ (the other read) UNCHANGED
ACCGGTAATGATACCCGCCAGTACACCCATGTTTATTTCTGGGTTGATGG
+position: -4548423
IIIIIIIIIIIIIIIIIIIIIIIIIIIIIIIIIIIIIIIIIIIIIIIIII
@ read 1081133  LOWQ (the other read) UNCHANGED
GTACATTGATGAGCAACATGGGGCTTGAACTGGCGCTGAAACAGTTAGGA
+position: -144740
IIIIIIIIIIIIIIIIIIIIIIIIIIIIIIIIIIIIIIIIIIIIIIIIII
@ read 108114 LOWQ DISCARDED (slave)
CTTAACCTGAACATTGAAGAGTTTAAAGGCGAGCAGTTACGCGTGACGGG
+position: 5075572
IIIIIIIIIIIIIIIIIIIIIIIIIIIIIIIIIIIIIIIIIIIIIIIIII
@ read 108116 LOWQ DISCARDED (slave)
TCAGGGTGCAGCGGTAGCGCCAGTTTCCAGTCCATCGCCATTTTGTAGAC
+position: 1750468
IIIIIIIIIIIIIIIIIIIIIIIIIIIIIIIIIIIIIIIIIIIIIIIIII
@ read 108117 LOWQ UNCHANGED
ATTCACTGTAAGCGCGGTCTGGTATGGGGCGAGCTGGAATTTGTACTGCC
+position: 1091320
IIIIIIIIIIIIIIIIIIIIIIIIIIIIIIIIIIIIIIIIIIIIIIIIII
@ read 108118 LOWQ DISCARDED (slave)
CAGCAAAATCATGCCGAGGAAGGCAATGCAAAGATAGTTGCCGAACGGAT
+position: -4651833
IIIIIIIIIIIIIIIIIIIIIIIIIIIIIIIIIIIIIIIIIIIIIIIIII" >> $2 

}


function trimN {
# Check --trimN: add reads with N's, at the ends and in the middle
# 10 reads.
# Option STRIP/ENDS: 7 trimmed, 3 discarded.
echo "@ read 1037  STRIP, ENDS -> TRIMN:2:47
NNTCGACACAAGAAAATGCGCCAATTTTGAGCCAGACCCCAGTTACGCNN
+position: -2044610
IIIIIIIIIIIIIIIIIIIIIIIIIIIIIIIIIIIIIIIIIIIIIIIIII
@ read 1038 STRIP -> TRIMN:17:49, ENDS -> UNCHANGED
AAAAAACGGTCGTGGTNGGCTACTGTTATTAAAGCGTTGGCTACAAAAAG
+position: 1361068
IIIIIIIIIIIIIIIIIIIIIIIIIIIIIIIIIIIIIIIIIIIIIIIIII
@ read 1039 STRIP -> DISCARDED N, ENDS -> UNCHANGED
CCAACGGACNATGGAAGTGCTNCGGCTGGNGTTTTTATCCTCCGGCATTC
+position: -4282223
IIIIIIIIIIIIIIIIIIIIIIIIIIIIIIIIIIIIIIIIIIIIIIIIII
@ read 1040 STRIP -> DISCARDED N, ENDS -> UNCHANGED
GCAAACGTATTTATGGCGACNNNNNNNCCGGTCATCAGGCCGATAAACGC
+position: 3171482
IIIIIIIIIIIIIIIIIIIIIIIIIIIIIIIIIIIIIIIIIIIIIIIIII
@ read 1041 STRIP, ENDS -> TRIMN:2:49
NNCATGCGCGTTCACTGCCCGCGATCCAGGATATTTTTTCCAGGGGGACC
+position: -86494
IIIIIIIIIIIIIIIIIIIIIIIIIIIIIIIIIIIIIIIIIIIIIIIIII
@ read 1042 STRIP, ENDS -> TRIMN:0:46
TGTCTGGCAACCTGATTCATCAGGCTTTGCGTAAGCGGAACGATTAANNN
+position: -3861809
IIIIIIIIIIIIIIIIIIIIIIIIIIIIIIIIIIIIIIIIIIIIIIIIII
@ read 1043 STRIP -> DISCARDED, ENDS -> TRIMN:0:48 
AATCTGAAACGAAANCTGACAGCGNNCCCCGCTTCTGACAAAATAGGCGN
+position: -4685876
IIIIIIIIIIIIIIIIIIIIIIIIIIIIIIIIIIIIIIIIIIIIIIIIII
@ read 1044 STRIP -> TRIMN:4:46, ENDS -> TRIMN:2:46 
NNANATGATCCTGGTCTGCAATAATCGTAAAGGGGCCGTCAGCGTGTNNN
+position: 1377584
IIIIIIIIIIIIIIIIIIIIIIIIIIIIIIIIIIIIIIIIIIIIIIIIII
@ read 1045 STRIP -> TRIMN:12:43, ENDS -> UNCHANGED
CGCCATNCCGANAATGATTGCTTTCGCCATATGGGTACGTCCGGNGATCC
+position: -2919586
IIIIIIIIIIIIIIIIIIIIIIIIIIIIIIIIIIIIIIIIIIIIIIIIII
@ read 1046 STRIP -> TRIMN:15:49, ENDS -> UNCHANGED
CCCCTCTGCANNNCNCAATAATTTCATCGAGGAATCCGTTGCTTCGGGCT
+position: -3548964
IIIIIIIIIIIIIIIIIIIIIIIIIIIIIIIIIIIIIIIIIIIIIIIIII
@ read 1046 STRIP, ENDS -> TRIMX:4:46
CCNNCTCTGCACAATCAATAATTTCATCGAGGAATCCGTTGCTTCGGNNT
+position: -3548964
55IIIIIIIIIIIIIIIIIIIIIIIIIIIIIIIIIIIIIIIIIIIIIII5" >> $1

echo "@ read 1037 STRIP, ENDS -> UNCHANGED
AATCGACACAAGAAAATGCGCCAATTTTGAGCCAGACCCCAGTTACGCAA
+position: -2044610
IIIIIIIIIIIIIIIIIIIIIIIIIIIIIIIIIIIIIIIIIIIIIIIIII
@ read 1038 STRIP, ENDS -> UNCHANGED
AAAAAACGGTCGTGGTAGGCTACTGTTATTAAAGCGTTGGCTACAAAAAG
+position: 1361068
IIIIIIIIIIIIIIIIIIIIIIIIIIIIIIIIIIIIIIIIIIIIIIIIII
@ read 1039 STRIP -> DISCARDED (slave) ENDS -> UNCHANGED 
CCAACGGACAATGGAAGTGCTACGGCTGGAGTTTTTATCCTCCGGCATTC
+position: -4282223
IIIIIIIIIIIIIIIIIIIIIIIIIIIIIIIIIIIIIIIIIIIIIIIIII
@ read 1040 STRIP -> DISCARDED (slave) ENDS -> UNCHANGED 
GCAAACGTATTTATGGCGACAAAAAAACCGGTCATCAGGCCGATAAACGC
+position: 3171482
IIIIIIIIIIIIIIIIIIIIIIIIIIIIIIIIIIIIIIIIIIIIIIIIII
@ read 1041 STRIP, ENDS -> UNCHANGED
AACATGCGCGTTCACTGCCCGCGATCCAGGATATTTTTTCCAGGGGGACC
+position: -86494
IIIIIIIIIIIIIIIIIIIIIIIIIIIIIIIIIIIIIIIIIIIIIIIIII
@ read 1042 STRIP, ENDS -> UNCHANGED 
TGTCTGGCAACCTGATTCATCAGGCTTTGCGTAAGCGGAACGATTAAAAA
+position: -3861809
IIIIIIIIIIIIIIIIIIIIIIIIIIIIIIIIIIIIIIIIIIIIIIIIII
@ read 1043 STRIP -> DISCARDED, ENDS -> UNCHANGED 
AATCTGAAACGAAAACTGACAGCGAACCCCGCTTCTGACAAAATAGGCGA
+position: -4685876
IIIIIIIIIIIIIIIIIIIIIIIIIIIIIIIIIIIIIIIIIIIIIIIIII
@ read 1044 STRIP, ENDS -> UNCHANGED 
AAAAATGATCCTGGTCTGCAATAATCGTAAAGGGGCCGTCAGCGTGTAAA
+position: 1377584
IIIIIIIIIIIIIIIIIIIIIIIIIIIIIIIIIIIIIIIIIIIIIIIIII
@ read 1045 STRIP, ENDS -> UNCHANGED  
CGCCATACCGAAAATGATTGCTTTCGCCATATGGGTACGTCCGGAGATCC
+position: -2919586
IIIIIIIIIIIIIIIIIIIIIIIIIIIIIIIIIIIIIIIIIIIIIIIIII
@ read 1046 STRIP, ENDS -> UNCHANGED 
CCCCTCTGCAAAACACAATAATTTCATCGAGGAATCCGTTGCTTCGGGCT
+position: -3548964
IIIIIIIIIIIIIIIIIIIIIIIIIIIIIIIIIIIIIIIIIIIIIIIIII
@ read 1046 STRIP, ENDS -> UNCHANGED 
CCCCTCTGCAAAACACAATAATTTCATCGAGGAATCCGTTGCTTCGGGCT
+position: -3548964
IIIIIIIIIIIIIIIIIIIIIIIIIIIIIIIIIIIIIIIIIIIIIIIIII" >> $2
}

#################################### CODE ######################################

EColi_gen="../fa_fq_files/EColi_genome.fa"
rRNA_seq="../fa_fq_files/rRNA_modified.fa"
name="../fa_fq_files/EColi_rRNA_DS" # prefix of the output file
suf1=".read1.fq" # suffix output1
suf2=".read2.fq" # suffix output2 

# USE Claudios tool 
# Generate EColi reads
../fa_fq_files/simReads.pl ${EColi_gen} 0.01 0 50 0 > ${name}$suf1
../fa_fq_files/simReads.pl ${EColi_gen} 0.01 0 50 0 > ${name}$suf2

# Generates rRNA reads
../fa_fq_files/simReads.pl ${rRNA_seq} 0.005 0 50 0 >> ${name}$suf1
../fa_fq_files/simReads.pl ${rRNA_seq} 0.005 0 50 0 >> ${name}$suf2

# Create reads with low quality issues and with N's 
trimQ  ${name}$suf1 $name$suf2
trimQ  ${name}$suf2 $name$suf1
trimN  ${name}$suf1 $name$suf2
trimN  ${name}$suf2 $name$suf1

# Zip the output
gzip $name$suf1
gzip $name$suf2
```
