## Supplementary material for "FastqPuri: high-performance preprocessing of RNA-seq data": README

################################################################################
#  trimFilterDS examples:                                                      #
################################################################################
#                                                                              # 
# The examples work in the  following way:                                     #
#                                                                              #
# 1. See folder `fa_fq_files`. The files `EColi_rRNA_DS.read1.fq.gz` and       #
#    `EColi_rRNA_DS.read2.fq.gz` were created with `create_fq.sh` and contain: #
#    * 1000 reads of length 50 from `EColi_genome.fa` with NO errors.          #
#    * 5000 reads of length 50 from `rRNA_modified.fa` with NO errrors         #
#      (rRNA contaminations).                                                  #
#    * Artificially generated reads with low quality score (see `create_fq.sh`)#
#    * Artificially generated reads with Ns (see `create_fq.sh`).              #
# 2. `run_example_TREE.sh`: the code was tested with flags:                    # 
#    `../../bin/trimFilter -l 50 --ifq\                                        #
#      --ifq ../fa_fq_files/EColi_rRNA_DS.read1.fq.gz:../fa_fq_files/EColi_rRNA_DS.read1.fq.gz \#
#     --method TREE --ifa ../fa_fq_files/rRNA_modified.fa:0.2:30 \             #
#     --trimQ ENDSFRAC --trimN ENDS -o treeDS --adapters \                     #
#     ../fa_fq_files/ad_read1.fa:../fa_fq_files/ad_read2.fa:2:40               #
#    i.e., we check for contaminations from rRNA, trim reads with lowQ at      #
#    the ends and less than 5% in the remaining part, and strip reads          #
#    containing N's at the ends.                                               # 
# 3. `run_example_BLOOM.sh`:                                                   # 
#    trimFilterDS is run like in 2. but passing a bloom filter to look for     # 
#       contaminations with `score=0.4` and the --trimN STRIP option.          #
# 4. With this set up, it is possible to run further customized tests.         #
# 5. See the folder `adapters` for examples on adapter contaminations.         #
#                                                                              # 
# NOTE: `rRNA_modified.fa` is the `rRNA_CRUnit.fa` sequence, where we have     #
#        removed the lines containing N's for testing purposes.                #
#                                                                              # 
################################################################################
################################################################################
#  Sreport example:                                                            #
################################################################################
#                                                                              #
# In folder `./bin_files`, 30 fake `./trimFilterDS` binary output files were   #
# generated (with the `R` script `create_fake_bins.R`). An html output was     #
# created out of them (`./bin_files/filter_Sreport_example.html`). It can      # 
# re reproduced if you run:                                                    #
#   ../../bin/Sreport -i .bin_files/ -o ./bin_files/filter_Sreport_new -t D    #
#                                                                              #
################################################################################
