## Supplementary material for "FastqPuri: high-performance preprocessing of RNA-seq data": run_example_BLOOM.sh

```
#!/bin/bash

# This script calls the executable and saves the output. Uses a bloom filter 
fastq1="../fa_fq_files/EColi_rRNA_DS.read1.fq.gz"
fastq2="../fa_fq_files/EColi_rRNA_DS.read2.fq.gz"
trimFilterDS=../../bin/trimFilterPE
bf=../trimFilter_Sreport/rRNA_example.bf
ad1=../fa_fq_files/ad_read1.fa
ad2=../fa_fq_files/ad_read2.fa

# Run trimFilterDS using a tree
$trimFilterDS -l 50 --ifq "${fastq1}:${fastq2}" --idx  $bf:0.4 \
 --method BLOOM --trimQ ENDSFRAC --trimN STRIP -o bloomDS --adapter \
$ad1:$ad2:2:40
```
