## Supplementary material for "FastqPuri: high-performance preprocessing of RNA-seq data": run_example_TREE.sh

```
#!/bin/bash

# This script calls the executable and saves the output 
fastq1=../fa_fq_files/EColi_rRNA_DS.read1.fq.gz
fastq2=../fa_fq_files/EColi_rRNA_DS.read2.fq.gz
fasta=../fa_fq_files/rRNA_modified.fa
trimFilterDS=../../bin/trimFilterPE
ad1=../fa_fq_files/ad_read1.fa
ad2=../fa_fq_files/ad_read2.fa

# Run trimFilterDS using a tree
$trimFilterDS -l 50 --ifq "${fastq1}:${fastq2}" --ifa "$fasta":0.2:30 \
 --method TREE --trimQ ENDSFRAC --trimN ENDS -o treeDS  --adapter \
$ad1:$ad2:2:40
```
