## Supplementary material for "FastqPuri: high-performance preprocessing of RNA-seq data": adapters_8c.html

FastqPuri: src/adapters.c File Reference

FastqPuri

- src

Functions |
Variables

adapters.c File Reference

sequence manipulation for alignment
More...

`#include <stdlib.h>`  
`#include <string.h>`  
`#include <stdio.h>`  
`#include "adapters.h"`  
`#include "Lmer.h"`

Include dependency graph for adapters.c:

|  |  |
| --- | --- |
| Functions | |
| void | init\_alLUTs () |
|  | look up table initialization for alignment (used for adapters) More... |
| int | process\_seq (unsigned char \*packed, unsigned char \*sequence, int L, bool shift, bool isreverse) |
|  | Packs a sequence using alfw0, alfw1, albw0, albw1. More... |
| Ad\_seq \* | pack\_adapter (Fa\_data \*ptr\_fa) |
|  | reads a **Fa\_data** with adapters and stores them in an array of **Ad\_seq** structs. More... |
| double | obtain\_score (Fq\_read \*seq, int pos\_seq, Ad\_seq \*ptr\_adap, int pos\_ad) |
|  | computes score of a possible alignment, after having found a seed. More... |

|  |  |
| --- | --- |
| Variables | |
| static uint8\_t | alfw0 [256] |
| static uint8\_t | alfw1 [256] |
| static uint8\_t | albw0 [256] |
| static uint8\_t | albw1 [256] |
| uint8\_t | fw\_1B [256] |
| uint8\_t | bw\_1B [256] |

### Detailed Description

sequence manipulation for alignment

Author
:   Paula Perez paula.nosp@@m.o@@m.om

Date
:   23.09.2017

### Function Documentation

### ◆ init\_alLUTs()

|  |  |  |  |
| --- | --- | --- | --- |
| void init\_alLUTs | ( |  | ) |

look up table initialization for alignment (used for adapters)

It initializes: fw\_1B, bw\_1B. They are uint8\_t arrays with 256 elements. All elements are set to 0xFF excepting the ones corresponding to 'a', 'A', 'c', 'C', 'g', 'G', 't', 'T':

| Var | a,A | c,C | g,G | t,T | Var | a,A | c,C | g,G | t,T |
| --- | --- | --- | --- | --- | --- | --- | --- | --- | --- |
| alfw0 | 0x01 | 0x02 | 0x04 | 0x08 | albw0 | 0x08 | 0x04 | 0x02 | 0x01 |
| alfw1 | 0x10 | 0x20 | 0x40 | 0x80 | albw1 | 0x80 | 0x40 | 0x20 | 0x10 |

With this variables we will encode sequences that can be compared later on. Using the bitwise XOR operator, every mismatch will amount to two bits set to 1.

### ◆ obtain\_score()

|  |  |  |  |
| --- | --- | --- | --- |
| double obtain\_score | ( | Fq\_read \* | *seq*, |
|  |  | int | *pos\_seq*, |
|  |  | Ad\_seq \* | *ptr\_adap*, |
|  |  | int | *pos\_ad* |
|  | ) |  |  |

computes score of a possible alignment, after having found a seed.

The score is computed as follows:

- matching bases: score += log\_10(4)
- unmatching bases: score -= Q/10, where Q is the quality score.

Parameters
:   |  |  |
    | --- | --- |
    | seq | pointer to **Fq\_read**. |
    | pos\_seq | read starting position of the alignment |
    | ptr\_adap | pointer to **Ad\_seq**, contains the adapter info |
    | pos\_ad | adapter starting position of the alignment (reverse) |

Returns
:   score of the alignment

### ◆ pack\_adapter()

|  |  |  |  |  |
| --- | --- | --- | --- | --- |
| Ad\_seq\* pack\_adapter | ( | Fa\_data \* | *ptr\_fa* | ) |

reads a **Fa\_data** with adapters and stores them in an array of **Ad\_seq** structs.

It reads the fasta structure. For every entry, an **Ad\_seq** structure is allocated and the sequences are processed to create the packed sequences.

Parameters
:   |  |  |
    | --- | --- |
    | ptr\_fa | pointer to **Fa\_data** structure |

Returns
:   pointer to **Ad\_seq**, where the information is stored.

### ◆ process\_seq()

|  |  |  |  |
| --- | --- | --- | --- |
| int process\_seq | ( | unsigned char \* | *packed*, |
|  |  | unsigned char \* | *sequence*, |
|  |  | int | *L*, |
|  |  | bool | *shift*, |
|  |  | bool | *isreverse* |
|  | ) |  |  |

Packs a sequence using alfw0, alfw1, albw0, albw1.

It takes a sequence of length L and packs it using the look up tables into an unsigned char array, where every bytes corresponds to 2 nucleotides. One can encode the reverse complement or the sequence shifted by 1/2 byte.

Parameters
:   |  |  |
    | --- | --- |
    | packed | packed sequence |
    | sequence | original sequence |
    | L | original sequence length |
    | shift | 0 if taken as is we want to shift the output 1/2 byte (>>4) |
    | isreverse | 0 if we want the forward sequence, 1 reverse complement |

Returns
:   Lhalf, length in Bytes of the packed sequence

### Variable Documentation

### ◆ albw0

uint8\_t albw0[256]

static

variable for brackward packing, first half

### ◆ albw1

uint8\_t albw1[256]

static

variable for brackward packing, second half

### ◆ alfw0

uint8\_t alfw0[256]

static

variable for forward packing, first half

### ◆ alfw1

uint8\_t alfw1[256]

static

variable for forward packing, second half

## ◆ bw\_1B

uint8\_t bw\_1B[256]

global variable. Lookup table.

## ◆ fw\_1B

uint8\_t fw\_1B[256]

global variable. Lookup table.

---

Generated on Mon Mar 19 2018 23:42:01 for FastqPuri by  

 1.8.14
