## Supplementary material for "FastqPuri: high-performance preprocessing of RNA-seq data": adapters_8h_source.html

FastqPuri: include/adapters.h Source File


FastqPuri


- include

adapters.h

Go to the documentation of this file.

1 /\*\*\*\*\*\*\*\*\*\*\*\*\*\*\*\*\*\*\*\*\*\*\*\*\*\*\*\*\*\*\*\*\*\*\*\*\*\*\*\*\*\*\*\*\*\*\*\*\*\*\*\*\*\*\*\*\*\*\*\*\*\*\*\*\*\*\*\*\*\*\*\*\*\*\*\*

2  \* Copyright (C) 2017 by Paula Perez Rubio \*

3  \* \*

4  \* This file is part of FastqPuri. \*

5  \* \*

6  \* FastqPuri is free software: you can redistribute it and/or modify \*

7  \* it under the terms of the GNU General Public License as \*

8  \* published by the Free Software Foundation, either version 3 of the \*

9  \* License, or (at your option) any later version. \*

10  \* \*

11  \* FastqPuri is distributed in the hope that it will be useful, \*

12  \* but WITHOUT ANY WARRANTY; without even the implied warranty of \*

13  \* MERCHANTABILITY or FITNESS FOR A PARTICULAR PURPOSE. See the \*

14  \* GNU General Public License for more details. \*

15  \* \*

16  \* You should have received a copy of the GNU General Public License \*

17  \* along with FastqPuri. \*

18  \* If not, see <http://www.gnu.org/licenses/>. \*

19  \*\*\*\*\*\*\*\*\*\*\*\*\*\*\*\*\*\*\*\*\*\*\*\*\*\*\*\*\*\*\*\*\*\*\*\*\*\*\*\*\*\*\*\*\*\*\*\*\*\*\*\*\*\*\*\*\*\*\*\*\*\*\*\*\*\*\*\*\*\*\*\*\*\*\*\*/

20

28 #ifndef ADAPTERS\_H\_

29 #define ADAPTERS\_H\_

30

31 #include "fq\_read.h"

32 #include "fa\_read.h"

33 #include "defines.h"

34

38 typedef struct \_ad\_seq {

39  int L;

40  char seq[READ\_MAXLEN];

41  int Lpack;

42  int Lpack\_sh;

43  unsigned char pack[(READ\_MAXLEN+1)/2];

44  unsigned char pack\_sh[(READ\_MAXLEN+1)/2];

45 } Ad\_seq;

46

47 void init\_alLUTs();

48

49 int process\_seq(unsigned char \*packed, unsigned char \*read, int L, bool shift,

50  bool isreverse);

51

52 Ad\_seq \*pack\_adapter(Fa\_data \*ptr\_fa);

53

54 double obtain\_score(Fq\_read \*seq, int pos\_seq, Ad\_seq \*ptr\_adap, int pos\_ad);

55

56 #endif // endif INIT\_ALIGNER\_H\_

\_ad\_seq::L

int L

**Definition:** adapters.h:39

\_ad\_seq::pack\_sh

unsigned char pack\_sh[(READ\_MAXLEN+1)/2]

**Definition:** adapters.h:44

\_fq\_read

stores a fastq entry

**Definition:** fq\_read.h:37

\_fa\_data

stores sequences of a fasta file

**Definition:** fa\_read.h:46

Ad\_seq

struct \_ad\_seq Ad\_seq

stores an adapter entry

pack\_adapter

Ad\_seq \* pack\_adapter(Fa\_data \*ptr\_fa)

reads a Fa\_data with adapters and stores them in an array of Ad\_seq structs.

**Definition:** adapters.c:129

fa\_read.h

reads in and stores fasta files

\_ad\_seq::Lpack

int Lpack

**Definition:** adapters.h:41

defines.h

Macro definitions.

\_ad\_seq::Lpack\_sh

int Lpack\_sh

**Definition:** adapters.h:42

\_ad\_seq

stores an adapter entry

**Definition:** adapters.h:38

process\_seq

int process\_seq(unsigned char \*packed, unsigned char \*read, int L, bool shift, bool isreverse)

Packs a sequence using alfw0, alfw1, albw0, albw1.

**Definition:** adapters.c:92

fq\_read.h

fastq entries manipulations (read/write)

obtain\_score

double obtain\_score(Fq\_read \*seq, int pos\_seq, Ad\_seq \*ptr\_adap, int pos\_ad)

computes score of a possible alignment, after having found a seed.

**Definition:** adapters.c:155

\_ad\_seq::pack

unsigned char pack[(READ\_MAXLEN+1)/2]

**Definition:** adapters.h:43

\_ad\_seq::seq

char seq[READ\_MAXLEN]

**Definition:** adapters.h:40

init\_alLUTs

void init\_alLUTs()

look up table initialization for alignment (used for adapters)

**Definition:** adapters.c:59


---

Generated on Mon Mar 19 2018 23:42:01 for FastqPuri by  

 1.8.14
