## Supplementary material for "FastqPuri: high-performance preprocessing of RNA-seq data": annotated.html

FastqPuri: Class List


FastqPuri


Class List

Here are the classes, structs, unions and interfaces with brief descriptions:

|  |  |
| --- | --- |
| C\_ad\_seq | Stores an adapter entry |
| C\_adapter |  |
| C\_bfilter | Bloom filter structure |
| C\_bfkmer | Stores a processed kmer (2 bits pro nucleotide) |
| C\_ds\_adap | Structure containing an adapter pair (for read 1 and read 2) |
| C\_fa\_data | Stores sequences of a fasta file |
| C\_fa\_entry | Fasta entry |
| C\_fq\_read | Stores a fastq entry |
| C\_iparam\_makeBloom | MakeBloom input parameters |
| C\_iparam\_makeTree | MakeTree input parameters |
| C\_iparam\_Qreport | Qreport input parameters |
| C\_iparam\_Sreport | Sreport input parameters |
| C\_iparam\_trimFilter | TrimFilter input parameters |
| C\_node | Node structure: formed out of T\_ACGT pointers to Node structure |
| C\_split | Splitted string and the number or splitted fields |
| C\_stats\_TF | Collects stats info from the filtering procedure |
| C\_stats\_TFDS | Collects stats info from the filtering procedure |
| C\_tree | Structure containing a T\_ACGT-tree |
| C\_uint128 |  |
| Cstatsinfo | Stores info needed to create the summary graphs |


---

Generated on Mon Mar 19 2018 23:42:01 for FastqPuri by  

 1.8.14
