## Supplementary material for "FastqPuri: high-performance preprocessing of RNA-seq data": bloom_8c.html

FastqPuri: src/bloom.c File Reference

FastqPuri

- src

Functions |
Variables

bloom.c File Reference

functions that implement the bloom filter
More...

`#include "bloom.h"`  
`#include <string.h>`  
`#include <stdio.h>`

Include dependency graph for bloom.c:

|  |  |
| --- | --- |
| Functions | |
| void | init\_LUTs () |
|  | look up table initialization More... |
| Bfilter \* | init\_Bfilter (int kmersize, uint64\_t bfsizeBits, int hashNum, double falsePosRate, uint64\_t nelem) |
|  | initialization of a Bfilter structure More... |
| void | free\_Bfilter (Bfilter \*ptr\_bf) |
|  | free Bfilter memory |
| Bfkmer \* | init\_Bfkmer (int kmersize, int hashNum) |
|  | initializes a Bfkmer structure, given the kmersize and the number of hash functions More... |
| void | free\_Bfkmer (Bfkmer \*ptr\_bfkmer) |
|  | free Bfkmer |
| int | compact\_kmer (const unsigned char \*sequence, uint64\_t position, Bfkmer \*ptr\_bfkmer) |
|  | compactifies a kmer for insertion in the bloomfilter More... |
| void | multiHash (Bfkmer \*ptr\_bfkmer) |
|  | obtains the hashNum hashvalues for a compactified kmer More... |
| bool | insert\_and\_fetch (Bfilter \*ptr\_bf, Bfkmer \*ptr\_bfkmer) |
|  | inserts the hashvalues of a kmer in filter More... |
| bool | contains (Bfilter \*ptr\_bf, Bfkmer \*ptr\_bfkmer) |
|  | check if kmer is contained in the filter More... |
| Bfilter \* | create\_Bfilter (Fa\_data \*ptr\_fasta, int kmersize, uint64\_t bfsizeBits, int hashNum, double falsePosRate, uint64\_t nelem) |
|  | creates a bloom filter from a fasta structure. More... |
| void | save\_Bfilter (Bfilter \*ptr\_bf, char \*filterfile, char \*paramfile) |
|  | saves a bloomfilter to disk More... |
| Bfilter \* | read\_Bfilter (char \*filterfile, char \*paramfile) |
|  | reads a bloom filter from a file More... |

|  |  |
| --- | --- |
| Variables | |
| static uint8\_t | fw0 [256] |
|  | Global variables (lookup table) Used to compactify kmers. |
| static uint8\_t | **fw1** [256] |
| static uint8\_t | **fw2** [256] |
| static uint8\_t | **fw3** [256] |
| static uint8\_t | **bw0** [256] |
| static uint8\_t | **bw1** [256] |
| static uint8\_t | **bw2** [256] |
| static uint8\_t | **bw3** [256] |
| static const unsigned char | bitMask [0x08] |
|  | bitMask, ith bit set to 1 in position i More... |
| uint64\_t | alloc\_mem |

### Detailed Description

functions that implement the bloom filter

Author
:   Paula Perez paula.nosp@@m.o@@m.om

Date
:   04.09.2017

### Function Documentation

### ◆ compact\_kmer()

|  |  |  |  |
| --- | --- | --- | --- |
| int compact\_kmer | ( | const unsigned char \* | *sequence*, |
|  |  | uint64\_t | *position*, |
|  |  | Bfkmer \* | *ptr\_bfkmer* |
|  | ) |  |  |

compactifies a kmer for insertion in the bloomfilter

Parameters
:   |  |  |
    | --- | --- |
    | sequence | unsigned char DNA sequence (or cDNA) |
    | position | position in the sequence where the kmer starts |
    | ptr\_bfkmer | initialized Bfkmer |

The compactified sequence is computed in the following way:

- We start compactifying both, the forward and backward (reverse complement). The outer loop covers up until half of the sequence.
- As soon as one of the two is lexicographically smaller, we continue only with it. In that way, the "smaller" sequence is consistently returned.
- If the sequence is palindromic, we continue with the forward sequence.
- kmersize should be > 3.

  We illustrate the compactification with an example:

  kmer = TTTT|GGAT

  m\_fw = 00000000 | 00000000 // 2 bytes

  m\_bw = 00000000 | 00000000 // 2 bytes

  m\_fw[0] |= fw0['T'] = 0xC0|0x00; m\_bw[0] |= bw0['T'] = 0x00|0x00;

  m\_fw[0] |= fw1['T'] = 0xF0|0x00; m\_bw[0] |= bw1['A'] = 0x30|0x00;

  m\_fw[0] |= fw2['T'] = 0xFC|0x00; m\_bw[0] |= bw2['G'] = 0x34|0x00;

  m\_fw[0] |= fw3['T'] = 0xFF|0x00; m\_bw[0] |= bw3['G'] = 0x35|0x00;

  m\_fw[1] |= fw0['G'] = 0xC0|0x80; m\_bw[1] |= bw0['T'] = 0x35|0x00;

  m\_fw[1] |= fw1['G'] = 0xF0|0xA0; m\_bw[1] |= bw1['T'] = 0x35|0x00;

  m\_fw[1] |= fw2['A'] = 0xFC|0xA0; m\_bw[1] |= bw2['T'] = 0x35|0x00;

  m\_fw[1] |= fw3['T'] = 0xFF|0xA3; m\_bw[1] |= bw3['T'] = 0x35|0x00;

  (In this case, we would store m\_bw)

### ◆ contains()

|  |  |  |  |
| --- | --- | --- | --- |
| bool contains | ( | Bfilter \* | *ptr\_bf*, |
|  |  | Bfkmer \* | *ptr\_bfkmer* |
|  | ) |  |  |

check if kmer is contained in the filter

Parameters
:   |  |  |
    | --- | --- |
    | ptr\_bf | pointer to a Bfilter structure, where a bloomfilter is stored |
    | ptr\_bfkmer | pointer to a Bfkmer structure containing the hash values |

Returns
:   true if all corresponding bits were set to 1 in the filter

### ◆ create\_Bfilter()

|  |  |  |  |
| --- | --- | --- | --- |
| Bfilter\* create\_Bfilter | ( | Fa\_data \* | *ptr\_fasta*, |
|  |  | int | *kmersize*, |
|  |  | uint64\_t | *bfsizeBits*, |
|  |  | int | *hashNum*, |
|  |  | double | *falsePosRate*, |
|  |  | uint64\_t | *nelem* |
|  | ) |  |  |

creates a bloom filter from a fasta structure.

Parameters
:   |  |  |
    | --- | --- |
    | ptr\_fasta | pointer to fasta structure |
    | kmersize | length of kmers to be inserted in the filter |
    | bfsizeBits | size of Bloom filter in bits |
    | hashNum | number of hash functions to be used |
    | falsePosRate | false positive rate |
    | nelem | number of elemens (kmers in the sequece) contained in the filter |

Returns
:   pointer to Bloom filter structure, where the fasta file was encoded.

### ◆ init\_Bfilter()

|  |  |  |  |
| --- | --- | --- | --- |
| Bfilter\* init\_Bfilter | ( | int | *kmersize*, |
|  |  | uint64\_t | *bfsizeBits*, |
|  |  | int | *hashNum*, |
|  |  | double | *falsePosRate*, |
|  |  | uint64\_t | *nelem* |
|  | ) |  |  |

initialization of a Bfilter structure

Parameters
:   |  |  |
    | --- | --- |
    | kmersize | number of elements of the kmer |
    | bfsizeBits | size of the bloomfilter (in Bits) |
    | hashNum | number of hash functions to be computed |
    | falsePosRate | false positive rate |
    | nelem | number of elemens (kmers in the sequece) contained in the filter |

Returns
:   pointer to initialized Bfilter structure

Given a kmersize, bfsizeBits, number of hash functions, we assign these values to the struture and the two additional values: kmersizeBytes = (kmersize + BASESINCHAR - 1 )/BASESINCHAR

### ◆ init\_Bfkmer()

|  |  |  |  |
| --- | --- | --- | --- |
| Bfkmer\* init\_Bfkmer | ( | int | *kmersize*, |
|  |  | int | *hashNum* |
|  | ) |  |  |

initializes a Bfkmer structure, given the kmersize and the number of hash functions

Parameters
:   |  |  |
    | --- | --- |
    | kmersize | number of elements of the kmer |
    | hashNum | number of hash functions to be computed |

Returns
:   pointer to a Bfkmer structure

kmersizeBytes, halfsizeBytes, hangingBases, hasOverhead hashNum are assigned and memory is allocated and set to 0 for compact and hashValues

### ◆ init\_LUTs()

|  |  |  |  |
| --- | --- | --- | --- |
| void init\_LUTs | ( |  | ) |

look up table initialization

It initializes: fw0, fw1, fw2, fw3, bw0, bw2, bw3, bw4. They are uint8\_t arrays with 256 elements. All elements are set to 0xFF excepting the ones corresponding to 'a', 'A', 'c', 'C', 'g', 'G', 't', 'T':

| Var | a,A | c,C | g,G | t,T | Var | a,A | c,C | g,G | t,T |
| --- | --- | --- | --- | --- | --- | --- | --- | --- | --- |
| fw0 | 0x00 | 0x40 | 0x80 | 0xC0 | bw0 | 0xC0 | 0x80 | 0x40 | 0x00 |
| fw1 | 0x00 | 0x10 | 0x20 | 0x30 | bw1 | 0x30 | 0x20 | 0x10 | 0x00 |
| fw2 | 0x00 | 0x04 | 0x08 | 0x0C | bw2 | 0x0C | 0x08 | 0x04 | 0x00 |
| fw3 | 0x00 | 0x01 | 0x02 | 0x03 | bw3 | 0x03 | 0x02 | 0x01 | 0x00 |

With these variables, we will be able to encode a Sequence using 2 bits per nucleotide.

### ◆ insert\_and\_fetch()

|  |  |  |  |
| --- | --- | --- | --- |
| bool insert\_and\_fetch | ( | Bfilter \* | *ptr\_bf*, |
|  |  | Bfkmer \* | *ptr\_bfkmer* |
|  | ) |  |  |

inserts the hashvalues of a kmer in filter

Parameters
:   |  |  |
    | --- | --- |
    | ptr\_bf | pointer to Bfilter structure, where we will include the new entry |
    | ptr\_bfkmer | pointer to Bfkmer structure, where the hashvalues are stored |

Returns
:   true if the positions of the hash values were already set to one previously.

The hash values are inserted in the following way.

- modValue = hashvalue mod(filter size) is calculated.
- the bit in position modValue of the filter is set to 1.

### ◆ multiHash()

|  |  |  |  |  |
| --- | --- | --- | --- | --- |
| void multiHash | ( | Bfkmer \* | *ptr\_bfkmer* | ) |

obtains the hashNum hashvalues for a compactified kmer

The hash values are computed using the CityHash64 hash functions.

### ◆ read\_Bfilter()

|  |  |  |  |
| --- | --- | --- | --- |
| Bfilter\* read\_Bfilter | ( | char \* | *filterfile*, |
|  |  | char \* | *paramfile* |
|  | ) |  |  |

reads a bloom filter from a file

Parameters
:   |  |  |
    | --- | --- |
    | filterfile | path to file containing the filter |
    | paramfile | path to file containing the filter |

Returns
:   a pointer to a filter structure containing the bloomfilter

This function reads two files, the auxiliar inputfile where kmersize, hashNum and bfsizeBits are stored, and the actual filter file. If one of them is missing, the program exits with an error. If successful, a pointer to a Bfilter structure with the bloom filter is return

### ◆ save\_Bfilter()

|  |  |  |  |
| --- | --- | --- | --- |
| void save\_Bfilter | ( | Bfilter \* | *ptr\_bf*, |
|  |  | char \* | *filterfile*, |
|  |  | char \* | *paramfile* |
|  | ) |  |  |

saves a bloomfilter to disk

Parameters
:   |  |  |
    | --- | --- |
    | ptr\_bf | pointer to Bfilter structure (contains the filter) |
    | filterfile | path to file where the output will be stored |
    | paramfile | path to file where the prameters will be stored |

This function will save the bloomfilter in the path filterfile. The paramfile will store the following data:

- kmersize
- hashNum
- bfsizeBits
- falsePosRate
- nelem

### Variable Documentation

### ◆ alloc\_mem

uint64\_t alloc\_mem

allocated memory

global variable. Memory allocated in the heap.

### ◆ bitMask

const unsigned char bitMask[0x08]

static

**Initial value:**

= {0x01, 0x02, 0x04, 0x08,

0x10, 0x20, 0x40, 0x80}

bitMask, ith bit set to 1 in position i

---

Generated on Mon Mar 19 2018 23:42:01 for FastqPuri by  

 1.8.14
