## Supplementary material for "FastqPuri: high-performance preprocessing of RNA-seq data": bloom_8h_source.html

FastqPuri: include/bloom.h Source File


FastqPuri


- include

bloom.h

Go to the documentation of this file.

1 /\*\*\*\*\*\*\*\*\*\*\*\*\*\*\*\*\*\*\*\*\*\*\*\*\*\*\*\*\*\*\*\*\*\*\*\*\*\*\*\*\*\*\*\*\*\*\*\*\*\*\*\*\*\*\*\*\*\*\*\*\*\*\*\*\*\*\*\*\*\*\*\*\*\*\*\*

2  \* Copyright (C) 2017 by Paula Perez Rubio \*

3  \* \*

4  \* This file is part of FastqPuri. \*

5  \* \*

6  \* FastqPuri is free software: you can redistribute it and/or modify \*

7  \* it under the terms of the GNU General Public License as \*

8  \* published by the Free Software Foundation, either version 3 of the \*

9  \* License, or (at your option) any later version. \*

10  \* \*

11  \* FastqPuri is distributed in the hope that it will be useful, \*

12  \* but WITHOUT ANY WARRANTY; without even the implied warranty of \*

13  \* MERCHANTABILITY or FITNESS FOR A PARTICULAR PURPOSE. See the \*

14  \* GNU General Public License for more details. \*

15  \* \*

16  \* You should have received a copy of the GNU General Public License \*

17  \* along with FastqPuri. \*

18  \* If not, see <http://www.gnu.org/licenses/>. \*

19  \*\*\*\*\*\*\*\*\*\*\*\*\*\*\*\*\*\*\*\*\*\*\*\*\*\*\*\*\*\*\*\*\*\*\*\*\*\*\*\*\*\*\*\*\*\*\*\*\*\*\*\*\*\*\*\*\*\*\*\*\*\*\*\*\*\*\*\*\*\*\*\*\*\*\*\*/

20

29 #ifndef BLOOM\_MAKER\_H\_

30 #define BLOOM\_MAKER\_H\_

31

32 #include "city.h"

33 #include "fa\_read.h"

34 #include "defines.h"

35

36

41 typedef struct \_bfilter {

42  int kmersize;

43  int hashNum;

44  int kmersizeBytes;

45  double falsePosRate;

46  uint64\_t bfsizeBits;

47  uint64\_t bfsizeBytes;

48  uint64\_t nelem;

49  unsigned char \*filter;

50 } Bfilter;

51

52

57 typedef struct \_bfkmer {

58  int kmersize;

59  int hashNum;

60  int kmersizeBytes;

61  int halfsizeBytes;

63  int hangingBases;

64  int hasOverhead;

65  unsigned char \*compact;

66  uint64\_t \*hashValues;

67 } Bfkmer;

68

69 void init\_LUTs();

70

71 Bfilter \*init\_Bfilter(int kmersize, uint64\_t bfsizeBits, int hashNum,

72  double falsePosRate, uint64\_t nelem);

73

74 Bfkmer \*init\_Bfkmer(int kmersize, int hashNum);

75

76 void free\_Bfilter(Bfilter \*ptr\_bf);

77

78 void free\_Bfkmer(Bfkmer \*ptr\_bfkmer);

79

80 int compact\_kmer(const unsigned char \*sequence, uint64\_t position,

81  Bfkmer \*ptr\_bfkmer);

82

83 void multiHash(Bfkmer\* ptr\_bfkmer);

84

85 bool insert\_and\_fetch(Bfilter \*pr\_bf, Bfkmer\* ptr\_bfkmer);

86

87 bool contains(Bfilter \*ptr\_bf, Bfkmer\* ptr\_bfkmer);

88

89 Bfilter \*create\_Bfilter(Fa\_data \*ptr\_fasta, int kmersize, uint64\_t bfsizeBits,

90  int hashNum, double falsePosRate, uint64\_t nelem);

91

92 void save\_Bfilter(Bfilter \*ptr\_bf, char \*filterfile, char \*paramfile);

93

94 Bfilter \*read\_Bfilter(char \*filterfile, char \*paramfile);

95

96

97 #endif // endif BLOOM\_MAKER\_H\_

free\_Bfilter

void free\_Bfilter(Bfilter \*ptr\_bf)

free Bfilter memory

**Definition:** bloom.c:143

save\_Bfilter

void save\_Bfilter(Bfilter \*ptr\_bf, char \*filterfile, char \*paramfile)

saves a bloomfilter to disk

**Definition:** bloom.c:544

city.h

functions for hashin strings, C translation of cityhash (C++, google)

init\_Bfilter

Bfilter \* init\_Bfilter(int kmersize, uint64\_t bfsizeBits, int hashNum, double falsePosRate, uint64\_t nelem)

initialization of a Bfilter structure

**Definition:** bloom.c:109

\_fa\_data

stores sequences of a fasta file

**Definition:** fa\_read.h:46

Bfkmer

struct \_bfkmer Bfkmer

stores a processed kmer (2 bits pro nucleotide)

\_bfilter::filter

unsigned char \* filter

**Definition:** bloom.h:49

fa\_read.h

reads in and stores fasta files

\_bfkmer::hashNum

int hashNum

**Definition:** bloom.h:59

init\_Bfkmer

Bfkmer \* init\_Bfkmer(int kmersize, int hashNum)

initializes a Bfkmer structure, given the kmersize and the number of hash functions ...

**Definition:** bloom.c:159

\_bfkmer::hashValues

uint64\_t \* hashValues

**Definition:** bloom.h:66

\_bfkmer::halfsizeBytes

int halfsizeBytes

**Definition:** bloom.h:61

free\_Bfkmer

void free\_Bfkmer(Bfkmer \*ptr\_bfkmer)

free Bfkmer

**Definition:** bloom.c:183

defines.h

Macro definitions.

\_bfkmer::kmersizeBytes

int kmersizeBytes

**Definition:** bloom.h:60

\_bfilter::hashNum

int hashNum

**Definition:** bloom.h:43

\_bfilter

Bloom filter structure.

**Definition:** bloom.h:41

contains

bool contains(Bfilter \*ptr\_bf, Bfkmer \*ptr\_bfkmer)

check if kmer is contained in the filter

**Definition:** bloom.c:477

\_bfkmer::hangingBases

int hangingBases

**Definition:** bloom.h:63

compact\_kmer

int compact\_kmer(const unsigned char \*sequence, uint64\_t position, Bfkmer \*ptr\_bfkmer)

compactifies a kmer for insertion in the bloomfilter

**Definition:** bloom.c:224

\_bfkmer::compact

unsigned char \* compact

**Definition:** bloom.h:65

Bfilter

struct \_bfilter Bfilter

Bloom filter structure.

\_bfilter::kmersizeBytes

int kmersizeBytes

**Definition:** bloom.h:44

create\_Bfilter

Bfilter \* create\_Bfilter(Fa\_data \*ptr\_fasta, int kmersize, uint64\_t bfsizeBits, int hashNum, double falsePosRate, uint64\_t nelem)

creates a bloom filter from a fasta structure.

**Definition:** bloom.c:501

\_bfkmer::hasOverhead

int hasOverhead

**Definition:** bloom.h:64

\_bfilter::bfsizeBits

uint64\_t bfsizeBits

**Definition:** bloom.h:46

\_bfkmer

stores a processed kmer (2 bits pro nucleotide)

**Definition:** bloom.h:57

init\_LUTs

void init\_LUTs()

look up table initialization

**Definition:** bloom.c:66

multiHash

void multiHash(Bfkmer \*ptr\_bfkmer)

obtains the hashNum hashvalues for a compactified kmer

**Definition:** bloom.c:435

read\_Bfilter

Bfilter \* read\_Bfilter(char \*filterfile, char \*paramfile)

reads a bloom filter from a file

**Definition:** bloom.c:587

\_bfilter::kmersize

int kmersize

**Definition:** bloom.h:42

\_bfilter::falsePosRate

double falsePosRate

**Definition:** bloom.h:45

\_bfkmer::kmersize

int kmersize

**Definition:** bloom.h:58

insert\_and\_fetch

bool insert\_and\_fetch(Bfilter \*pr\_bf, Bfkmer \*ptr\_bfkmer)

inserts the hashvalues of a kmer in filter

**Definition:** bloom.c:457

\_bfilter::nelem

uint64\_t nelem

**Definition:** bloom.h:48

\_bfilter::bfsizeBytes

uint64\_t bfsizeBytes

**Definition:** bloom.h:47


---

Generated on Mon Mar 19 2018 23:42:01 for FastqPuri by  

 1.8.14
