## Supplementary material for "FastqPuri: high-performance preprocessing of RNA-seq data": city_8c.html

FastqPuri: src/city.c File Reference

FastqPuri

- src

Macros |
Functions |
Variables

city.c File Reference

functions for hashin strings, C translation of cityhash (C++, google)
More...

`#include <string.h>`  
`#include "city.h"`  
`#include <byteswap.h>`

Include dependency graph for city.c:

|  |  |
| --- | --- |
| Macros | |
| #define | **uint32\_in\_expected\_order**(x)   (x) |
| #define | **uint64\_in\_expected\_order**(x)   (x) |
| #define | **LIKELY**(x)   (x) |
| #define | **PERMUTE3**(a, b, c)   do { std::swap(a, b); std::swap(a, c); } while (0) |
| #define | **PERMUTE3\_32**(a, b, c)   do { uint32\_t t = a; a = c; c = b; b = t;} while (0) |
| #define | **PERMUTE3\_64**(a, b, c)   do { uint64 t = a; a = c; c = b; b = t;} while (0) |

|  |  |
| --- | --- |
| Functions | |
| static uint64 | **UNALIGNED\_LOAD64** (const char \*p) |
| static uint32 | **UNALIGNED\_LOAD32** (const char \*p) |
| static uint64 | **Fetch64** (const char \*p) |
| static uint32 | **Fetch32** (const char \*p) |
| static uint32 | **fmix** (uint32 h) |
| static uint32 | **Rotate32** (uint32 val, int shift) |
| static uint32 | **Mur** (uint32 a, uint32 h) |
| static uint32 | **Hash32Len13to24** (const char \*s, size\_t len) |
| static uint32 | **Hash32Len0to4** (const char \*s, size\_t len) |
| static uint32 | **Hash32Len5to12** (const char \*s, size\_t len) |
| uint32 | **CityHash32** (const char \*s, size\_t len) |
| static uint64 | **Rotate** (uint64 val, int shift) |
| static uint64 | **ShiftMix** (uint64 val) |
| static uint64 | **HashLen16** (uint64 u, uint64 v) |
| static uint64 | **HashLen16\_3a** (uint64 u, uint64 v, uint64 mul) |
| static uint64 | **HashLen0to16** (const char \*s, size\_t len) |
| static uint64 | **HashLen17to32** (const char \*s, size\_t len) |
| uint128 | **WeakHashLen32WithSeeds** (uint64 w, uint64 x, uint64 y, uint64 z, uint64 a, uint64 b) |
| uint128 | **WeakHashLen32WithSeeds\_3a** (const char \*s, uint64 a, uint64 b) |
| static uint64 | **HashLen33to64** (const char \*s, size\_t len) |
| uint64 | **CityHash64** (const char \*s, size\_t len) |
| uint64 | **CityHash64WithSeed** (const char \*s, size\_t len, uint64 seed) |
| uint64 | **CityHash64WithSeeds** (const char \*s, size\_t len, uint64 seed0, uint64 seed1) |
| static uint128 | **CityMurmur** (const char \*s, size\_t len, uint128 seed) |
| uint128 | **CityHash128WithSeed** (const char \*s, size\_t len, uint128 seed) |
| uint128 | **CityHash128** (const char \*s, size\_t len) |

|  |  |
| --- | --- |
| Variables | |
| static const uint64 | **k0** = 0xc3a5c85c97cb3127ULL |
| static const uint64 | **k1** = 0xb492b66fbe98f273ULL |
| static const uint64 | **k2** = 0x9ae16a3b2f90404fULL |
| static const uint32\_t | **c1** = 0xcc9e2d51 |
| static const uint32\_t | **c2** = 0x1b873593 |

### Detailed Description

functions for hashin strings, C translation of cityhash (C++, google)

Author
:   bdnt

See also
:   https://github.com/bdnt/cityhash-c
:   https://github.com/google/cityhash

---

Generated on Mon Mar 19 2018 23:42:01 for FastqPuri by  

 1.8.14
