## Supplementary material for "FastqPuri: high-performance preprocessing of RNA-seq data": city_8h.html

FastqPuri: include/city.h File Reference

FastqPuri

- include

Classes |
Macros |
Typedefs |
Functions

city.h File Reference

functions for hashin strings, C translation of cityhash (C++, google)
More...

`#include <stdlib.h>`  
`#include <stdint.h>`

Include dependency graph for city.h:

This graph shows which files directly or indirectly include this file:

Go to the source code of this file.

|  |  |
| --- | --- |
| Classes | |
| struct | \_uint128 |

|  |  |
| --- | --- |
| Macros | |
| #define | **Uint128Low64**(x)   (x).first |
| #define | **Uint128High64**(x)   (x).second |

|  |  |
| --- | --- |
| Typedefs | |
| typedef uint8\_t | **uint8** |
| typedef uint16\_t | **uint16** |
| typedef uint32\_t | **uint32** |
| typedef uint64\_t | **uint64** |
| typedef struct \_uint128 | **uint128** |

|  |  |
| --- | --- |
| Functions | |
| uint64\_t | **CityHash64** (const char \*buf, size\_t len) |
| uint64\_t | **CityHash64WithSeed** (const char \*buf, size\_t len, uint64\_t seed) |
| uint64\_t | **CityHash64WithSeeds** (const char \*buf, size\_t len, uint64\_t seed0, uint64\_t seed1) |
| uint128 | **CityHash128** (const char \*s, size\_t len) |
| uint128 | **CityHash128WithSeed** (const char \*s, size\_t len, uint128 seed) |
| uint32 | **CityHash32** (const char \*buf, size\_t len) |
| static uint64\_t | **Hash128to64** (const uint128 x) |

### Detailed Description

functions for hashin strings, C translation of cityhash (C++, google)

Author
:   bdnt

See also
:   https://github.com/bdnt/cityhash-c
:   https://github.com/google/cityhash

---

Generated on Mon Mar 19 2018 23:42:01 for FastqPuri by  

 1.8.14
