## Supplementary material for "FastqPuri: high-performance preprocessing of RNA-seq data": city_8h_source.html

FastqPuri: include/city.h Source File


FastqPuri


- include

city.h

Go to the documentation of this file.

1 // Copyright (c) 2011 Google, Inc.

2 //

3 // Permission is hereby granted, free of charge, to any person obtaining a copy

4 // of this software and associated documentation files (the "Software"), to deal

5 // in the Software without restriction, including without limitation the rights

6 // to use, copy, modify, merge, publish, distribute, sublicense, and/or sell

7 // copies of the Software, and to permit persons to whom the Software is

8 // furnished to do so, subject to the following conditions:

9 //

10 // The above copyright notice and this permission notice shall be included in

11 // all copies or substantial portions of the Software.

12 //

13 // THE SOFTWARE IS PROVIDED "AS IS", WITHOUT WARRANTY OF ANY KIND, EXPRESS OR

14 // IMPLIED, INCLUDING BUT NOT LIMITED TO THE WARRANTIES OF MERCHANTABILITY,

15 // FITNESS FOR A PARTICULAR PURPOSE AND NONINFRINGEMENT. IN NO EVENT SHALL THE

16 // AUTHORS OR COPYRIGHT HOLDERS BE LIABLE FOR ANY CLAIM, DAMAGES OR OTHER

17 // LIABILITY, WHETHER IN AN ACTION OF CONTRACT, TORT OR OTHERWISE, ARISING FROM,

18 // OUT OF OR IN CONNECTION WITH THE SOFTWARE OR THE USE OR OTHER DEALINGS IN

19 // THE SOFTWARE.

20 //

21 // CityHash, by Geoff Pike and Jyrki Alakuijala

22 //

23 // http://code.google.com/p/cityhash/

24 //

25 // This file provides a few functions for hashing strings. All of them are

26 // high-quality functions in the sense that they pass standard tests such

27 // as Austin Appleby's SMHasher. They are also fast.

28 //

29 // For 64-bit x86 code, on short strings, we don't know of anything faster than

30 // CityHash64 that is of comparable quality. We believe our nearest competitor

31 // is Murmur3. For 64-bit x86 code, CityHash64 is an excellent choice for hash

32 // tables and most other hashing (excluding cryptography).

33 //

34 // For 64-bit x86 code, on long strings, the picture is more complicated.

35 // On many recent Intel CPUs, such as Nehalem, Westmere, Sandy Bridge, etc.,

36 // CityHashCrc128 appears to be faster than all competitors of comparable

37 // quality. CityHash128 is also good but not quite as fast. We believe our

38 // nearest competitor is Bob Jenkins' Spooky. We don't have great data for

39 // other 64-bit CPUs, but for long strings we know that Spooky is slightly

40 // faster than CityHash on some relatively recent AMD x86-64 CPUs, for example.

41 //

42 // For 32-bit x86 code, we don't know of anything faster than CityHash32 that

43 // is of comparable quality. We believe our nearest competitor is Murmur3A.

44 // (On 64-bit CPUs, it is typically faster to use the other CityHash variants.)

45 //

46 // Functions in the CityHash family are not suitable for cryptography.

47 //

48 // WARNING: This code has been only lightly tested on big-endian platforms!

49 // It is known to work well on little-endian platforms that have a small penalty

50 // for unaligned reads, such as current Intel and AMD moderate-to-high-end CPUs.

51 // It should work on all 32-bit and 64-bit platforms that allow unaligned reads;

52 // bug reports are welcome.

53 //

54 // By the way, for some hash functions, given strings a and b, the hash

55 // of a+b is easily derived from the hashes of a and b. This property

56 // doesn't hold for any hash functions in this file.

57

67 #ifndef CITY\_HASH\_H\_

68 #define CITY\_HASH\_H\_

69

70 #include <stdlib.h> // for size\_t.

71 #include <stdint.h>

72

73

74

75 typedef uint8\_t uint8;

76 typedef uint16\_t uint16;

77 typedef uint32\_t uint32;

78 typedef uint64\_t uint64;

79

80 typedef struct \_uint128 uint128;

81 struct \_uint128 {

82  uint64 first;

83  uint64 second;

84 };

85

86 #define Uint128Low64(x) (x).first

87 #define Uint128High64(x) (x).second

88

89 //inline uint64\_t Uint128Low64(const uint128& x) { return x.first; }

90 //inline uint64\_t Uint128High64(const uint128& x) { return x.second; }

91

92 // Hash function for a byte array.

93 uint64\_t CityHash64(const char \*buf, size\_t len);

94

95 // Hash function for a byte array. For convenience, a 64-bit seed is also

96 // hashed into the result.

97 uint64\_t CityHash64WithSeed(const char \*buf, size\_t len, uint64\_t seed);

98

99 // Hash function for a byte array. For convenience, two seeds are also

100 // hashed into the result.

101 uint64\_t CityHash64WithSeeds(const char \*buf, size\_t len,

102  uint64\_t seed0, uint64\_t seed1);

103

104 // Hash function for a byte array.

105 uint128 CityHash128(const char \*s, size\_t len);

106

107 // Hash function for a byte array. For convenience, a 128-bit seed is also

108 // hashed into the result.

109 uint128 CityHash128WithSeed(const char \*s, size\_t len, uint128 seed);

110

111 // Hash function for a byte array. Most useful in 32-bit binaries.

112 uint32 CityHash32(const char \*buf, size\_t len);

113

114 // Hash 128 input bits down to 64 bits of output.

115 // This is intended to be a reasonably good hash function.

116 static inline uint64\_t Hash128to64(const uint128 x) {

117  // Murmur-inspired hashing.

118  const uint64\_t kMul = 0x9ddfea08eb382d69ULL;

119  uint64\_t a = (Uint128Low64(x) ^ Uint128High64(x)) \* kMul;

120  a ^= (a >> 47);

121  uint64\_t b = (Uint128High64(x) ^ a) \* kMul;

122  b ^= (b >> 47);

123  b \*= kMul;

124  return b;

125 }

126

127 #endif // CITY\_HASH\_H\_

\_uint128

**Definition:** city.h:81


---

Generated on Mon Mar 19 2018 23:42:01 for FastqPuri by  

 1.8.14
