## Supplementary material for "FastqPuri: high-performance preprocessing of RNA-seq data": citycrc_8h.html

FastqPuri: include/citycrc.h File Reference

FastqPuri

- include

Functions

citycrc.h File Reference

functions for hashin strings, C translation of cityhash (C++, google)
More...

`#include "city.h"`

Include dependency graph for citycrc.h:

Go to the source code of this file.

|  |  |
| --- | --- |
| Functions | |
| uint128 | **CityHashCrc128** (const char \*s, size\_t len) |
| uint128 | **CityHashCrc128WithSeed** (const char \*s, size\_t len, uint128 seed) |
| void | **CityHashCrc256** (const char \*s, size\_t len, uint64 \*result) |

### Detailed Description

functions for hashin strings, C translation of cityhash (C++, google)

Author
:   bdnt

See also
:   https://github.com/bdnt/cityhash-c
:   https://github.com/google/cityhash

---

Generated on Mon Mar 19 2018 23:42:01 for FastqPuri by  

 1.8.14
