## Supplementary material for "FastqPuri: high-performance preprocessing of RNA-seq data": citycrc_8h_source.html

FastqPuri: include/citycrc.h Source File


FastqPuri


- include

citycrc.h

Go to the documentation of this file.

1 // Copyright (c) 2011 Google, Inc.

2 //

3 // Permission is hereby granted, free of charge, to any person obtaining a copy

4 // of this software and associated documentation files (the "Software"), to deal

5 // in the Software without restriction, including without limitation the rights

6 // to use, copy, modify, merge, publish, distribute, sublicense, and/or sell

7 // copies of the Software, and to permit persons to whom the Software is

8 // furnished to do so, subject to the following conditions:

9 //

10 // The above copyright notice and this permission notice shall be included in

11 // all copies or substantial portions of the Software.

12 //

13 // THE SOFTWARE IS PROVIDED "AS IS", WITHOUT WARRANTY OF ANY KIND, EXPRESS OR

14 // IMPLIED, INCLUDING BUT NOT LIMITED TO THE WARRANTIES OF MERCHANTABILITY,

15 // FITNESS FOR A PARTICULAR PURPOSE AND NONINFRINGEMENT. IN NO EVENT SHALL THE

16 // AUTHORS OR COPYRIGHT HOLDERS BE LIABLE FOR ANY CLAIM, DAMAGES OR OTHER

17 // LIABILITY, WHETHER IN AN ACTION OF CONTRACT, TORT OR OTHERWISE, ARISING FROM,

18 // OUT OF OR IN CONNECTION WITH THE SOFTWARE OR THE USE OR OTHER DEALINGS IN

19 // THE SOFTWARE.

20 //

21 // CityHash, by Geoff Pike and Jyrki Alakuijala

22 //

23 // This file declares the subset of the CityHash functions that require

24 // \_mm\_crc32\_u64(). See the CityHash README for details.

25 //

26 // Functions in the CityHash family are not suitable for cryptography.

27

36 #ifndef CITY\_HASH\_CRC\_H\_

37 #define CITY\_HASH\_CRC\_H\_

38

39 #include "city.h"

40

41 // Hash function for a byte array.

42 uint128 CityHashCrc128(const char \*s, size\_t len);

43

44 // Hash function for a byte array. For convenience, a 128-bit seed is also

45 // hashed into the result.

46 uint128 CityHashCrc128WithSeed(const char \*s, size\_t len, uint128 seed);

47

48 // Hash function for a byte array. Sets result[0] ... result[3].

49 void CityHashCrc256(const char \*s, size\_t len, uint64 \*result);

50

51 #endif // endif CITY\_HASH\_CRC\_H\_

\_uint128

**Definition:** city.h:81

city.h

functions for hashin strings, C translation of cityhash (C++, google)


---

Generated on Mon Mar 19 2018 23:42:01 for FastqPuri by  

 1.8.14
