## Supplementary material for "FastqPuri: high-performance preprocessing of RNA-seq data": classes.html

FastqPuri: Class Index

FastqPuri

Class Index

\_ | s

|  |  |  |  |  |
| --- | --- | --- | --- | --- |
| \_ | \_bfkmer | \_iparam\_makeBloom | \_node | \_uint128 |
|  | \_ds\_adap | \_iparam\_makeTree | \_split | s |
| \_ad\_seq | \_fa\_data | \_iparam\_Qreport | \_stats\_TF |  |
| \_adapter | \_fa\_entry | \_iparam\_Sreport | \_stats\_TFDS | statsinfo |
| \_bfilter | \_fq\_read | \_iparam\_trimFilter | \_tree |  |

\_ | s

---

Generated on Mon Mar 19 2018 23:42:01 for FastqPuri by  

 1.8.14
