## Supplementary material for "FastqPuri: high-performance preprocessing of RNA-seq data": config_8h_source.html

My Project: config.h Source File


My Project


config.h

1 #define VERSION "1.0"

2 #define HAVE\_RPKG

3 #define RSCRIPT\_EXEC "/usr/local/bin/compdiag/Rscript\_RBioC"

4 #define READ\_MAXLEN 400

5 #define RMD\_QUALITY\_REPORT "/home/loc03475/bighome/projects/FastqPuri/R/quality\_report.Rmd"

6 #define RMD\_SUMMARY\_REPORT "/home/loc03475/bighome/projects/FastqPuri/R/summary\_report.Rmd"

7 #define RMD\_SUMMARY\_FILTER\_REPORT "/home/loc03475/bighome/projects/FastqPuri/R/summary\_filter\_report.Rmd"

8 #define RMD\_SUMMARY\_FILTER\_REPORTDS "/home/loc03475/bighome/projects/FastqPuri/R/summary\_filter\_reportDS.Rmd"


---

Generated by  

 1.8.14
