## Supplementary material for "FastqPuri: high-performance preprocessing of RNA-seq data": defines_8h.html

FastqPuri: include/defines.h File Reference

FastqPuri

- include

Macros

defines.h File Reference

Macro definitions.
More...

`#include <stdint.h>`  
`#include <inttypes.h>`

Include dependency graph for defines.h:

This graph shows which files directly or indirectly include this file:

Go to the source code of this file.

|  |  |
| --- | --- |
| Macros | |
| #define | B\_LEN   131072 |
| #define | MAX\_FILENAME   300 |
| #define | bool   int16\_t |
| #define | true   1 |
| #define | false   0 |
| #define | max(a, b)   (((a) > (b)) ? (a) : (b)) |
| #define | min(a, b)   (((a) < (b)) ? (a) : (b)) |
| #define | mem\_usageMB() |
| #define | mem\_usage() |
| #define | DEFAULT\_MINQ   27 |
| #define | DEFAULT\_NTILES   96 |
| #define | DEFAULT\_NQ   46 |
| #define | ZEROQ   33 |
| #define | N\_ACGT   5 |
| #define | MAX\_RCOMMAND   4000 |
| #define | FA\_ENTRY\_BUF   20 |
| #define | LOG\_4   0.60206 |
| #define | MIN\_NMATCHES   12 |
| #define | T\_ACGT   4 |
| #define | NPOOL\_1D   1048576 |
| #define | NPOOL\_2D   16 |
| #define | MAX\_FASZ\_TREE   1e7 |
| #define | BITSPERCHAR   8 |
| #define | BASESPERCHAR   4 |
| #define | KMER\_LEN   25 |
| #define | FALSE\_POS\_RATE   0.05 |
| #define | ZERO\_POS\_RATE   1e-14 |
| #define | NO   0 |
| #define | ALL   1 |
| #define | ENDS   2 |
| #define | STRIP   3 |
| #define | FRAC   3 |
| #define | ENDSFRAC   4 |
| #define | GLOBAL   5 |
| #define | TREE   1 |
| #define | BLOOM   2 |
| #define | ERROR   1000 |
| #define | DEFAULT\_MINL   25 |
| #define | ADAP   0 |
| #define | CONT   1 |
| #define | LOWQ   2 |
| #define | NNNN   3 |
| #define | GOOD   4 |
| #define | NFILTERS   4 |
| #define | ADAP2   5 |
| #define | CONT2   6 |
| #define | LOWQ2   7 |
| #define | NNNN2   8 |
| #define | GOOD2   9 |
| #define | NFILES\_DS   10 |

### Detailed Description

Macro definitions.

Author
:   Paula Perez paula.nosp@@m.o@@m.om

Date
:   07.08.2017

### Macro Definition Documentation

### ◆ ADAP

#define ADAP   0

Adapter filter

### ◆ ADAP2

#define ADAP2   5

Adapter filter read2

### ◆ ALL

#define ALL   1

Trims if a lowQ base calling | N is found

### ◆ B\_LEN

#define B\_LEN   131072

buffer size

### ◆ BASESPERCHAR

#define BASESPERCHAR   4

number of nucleotides that can fit in a char

### ◆ BITSPERCHAR

#define BITSPERCHAR   8

number of bits in a char

### ◆ BLOOM

#define BLOOM   2

Use a bloom filter to look for contaminations

### ◆ bool

#define bool   int16\_t

define a bool type

### ◆ CONT

#define CONT   1

Contamination filter

### ◆ CONT2

#define CONT2   6

Contamination filter read2

### ◆ DEFAULT\_MINL

#define DEFAULT\_MINL   25

Default minimum length under which we discard the reads

### ◆ DEFAULT\_MINQ

#define DEFAULT\_MINQ   27

Minimum quality threshold

### ◆ DEFAULT\_NQ

#define DEFAULT\_NQ   46

Default number of different quality values

### ◆ DEFAULT\_NTILES

#define DEFAULT\_NTILES   96

Default number of tiles

### ◆ ENDS

#define ENDS   2

Trims at the ends

### ◆ ENDSFRAC

#define ENDSFRAC   4

trims at the ends and discards a read if the remaining part has more than > percent lowQ bases

### ◆ ERROR

#define ERROR   1000

Encodes an error when reading in trimN, trimQ, method options in trimFilter

### ◆ FA\_ENTRY\_BUF

#define FA\_ENTRY\_BUF   20

buffer for fasta entries

### ◆ false

#define false   0

assign false to 0

### ◆ FALSE\_POS\_RATE

#define FALSE\_POS\_RATE   0.05

default false positive rate

### ◆ FRAC

#define FRAC   3

Discards a read if it contains > percent lowQ bases

### ◆ GLOBAL

#define GLOBAL   5

Trims a fixed # bases from e left and right

### ◆ GOOD

#define GOOD   4

Good reads

### ◆ GOOD2

#define GOOD2   9

Good reads read2

### ◆ KMER\_LEN

#define KMER\_LEN   25

default kmer length

### ◆ LOG\_4

#define LOG\_4   0.60206

log\_10(4) for the adapters alignment score

### ◆ LOWQ

#define LOWQ   2

Low quality filter

### ◆ LOWQ2

#define LOWQ2   7

Low quality filter read2

### ◆ max

|  |  |  |  |
| --- | --- | --- | --- |
| #define max | ( |  | a, |
|  |  |  | b |
|  | ) |  | (((a) > (b)) ? (a) : (b)) |

max function

### ◆ MAX\_FASZ\_TREE

#define MAX\_FASZ\_TREE   1e7

Maximum fasta size for constructing a tree. DECIDE A SENSIBLE SIZE

### ◆ MAX\_FILENAME

#define MAX\_FILENAME   300

Maximum # chars in a filename

### ◆ MAX\_RCOMMAND

#define MAX\_RCOMMAND   4000

Maximum # chars in R command

### ◆ mem\_usage

|  |  |  |  |
| --- | --- | --- | --- |
| #define mem\_usage | ( |  | ) |

**Value:**

fprintf(stderr, \

"- Current allocated memory: %" PRIu64 "Bytes.\n", \

alloc\_mem)

alloc\_mem

uint64\_t alloc\_mem

**Definition:** makeBloom.c:42

returns allocated memory in Bytes

### ◆ mem\_usageMB

|  |  |  |  |
| --- | --- | --- | --- |
| #define mem\_usageMB | ( |  | ) |

**Value:**

fprintf(stderr, \

"- Current allocated memory: %" PRIu64 "MB.\n", \

alloc\_mem >> 20)

alloc\_mem

uint64\_t alloc\_mem

**Definition:** makeBloom.c:42

returns allocated memory in MB

### ◆ min

|  |  |  |  |
| --- | --- | --- | --- |
| #define min | ( |  | a, |
|  |  |  | b |
|  | ) |  | (((a) < (b)) ? (a) : (b)) |

min function

### ◆ MIN\_NMATCHES

#define MIN\_NMATCHES   12

minimum number of matches demanded

### ◆ N\_ACGT

#define N\_ACGT   5

Number of different nucleotides in the fq file

### ◆ NFILES\_DS

#define NFILES\_DS   10

number of outputfiles in double stranded case

### ◆ NFILTERS

#define NFILTERS   4

total number of filters

### ◆ NNNN

#define NNNN   3

N's presence filter

### ◆ NNNN2

#define NNNN2   8

N's presence filter read2

## ◆ NO

#define NO   0

No trimming

### ◆ NPOOL\_1D

#define NPOOL\_1D   1048576

Number of Node structs allocated in inner dim

### ◆ NPOOL\_2D

#define NPOOL\_2D   16

Number of \*Node allocated in outer dim

### ◆ STRIP

#define STRIP   3

Looks for the largest N-free sequence

### ◆ T\_ACGT

#define T\_ACGT   4

Number of children per node in tree

### ◆ TREE

#define TREE   1

Use a tree to look for contaminations

### ◆ true

#define true   1

assign true to 1

### ◆ ZERO\_POS\_RATE

#define ZERO\_POS\_RATE   1e-14

0 threshold for a double

### ◆ ZEROQ

#define ZEROQ   33

ASCII code of lowest quality value (!)

---

Generated on Mon Mar 19 2018 23:42:01 for FastqPuri by  

 1.8.14
