## Supplementary material for "FastqPuri: high-performance preprocessing of RNA-seq data": defines_8h_source.html

FastqPuri: include/defines.h Source File


FastqPuri


- include

defines.h

Go to the documentation of this file.

1 /\*\*\*\*\*\*\*\*\*\*\*\*\*\*\*\*\*\*\*\*\*\*\*\*\*\*\*\*\*\*\*\*\*\*\*\*\*\*\*\*\*\*\*\*\*\*\*\*\*\*\*\*\*\*\*\*\*\*\*\*\*\*\*\*\*\*\*\*\*\*\*\*\*\*\*\*

2  \* Copyright (C) 2017 by Paula Perez Rubio \*

3  \* \*

4  \* This file is part of FastqPuri. \*

5  \* \*

6  \* FastqPuri is free software: you can redistribute it and/or modify \*

7  \* it under the terms of the GNU General Public License as \*

8  \* published by the Free Software Foundation, either version 3 of the \*

9  \* License, or (at your option) any later version. \*

10  \* \*

11  \* FastqPuri is distributed in the hope that it will be useful, \*

12  \* but WITHOUT ANY WARRANTY; without even the implied warranty of \*

13  \* MERCHANTABILITY or FITNESS FOR A PARTICULAR PURPOSE. See the \*

14  \* GNU General Public License for more details. \*

15  \* \*

16  \* You should have received a copy of the GNU General Public License \*

17  \* along with FastqPuri. \*

18  \* If not, see <http://www.gnu.org/licenses/>. \*

19  \*\*\*\*\*\*\*\*\*\*\*\*\*\*\*\*\*\*\*\*\*\*\*\*\*\*\*\*\*\*\*\*\*\*\*\*\*\*\*\*\*\*\*\*\*\*\*\*\*\*\*\*\*\*\*\*\*\*\*\*\*\*\*\*\*\*\*\*\*\*\*\*\*\*\*\*/

20

29 #ifndef DEFINES\_H\_

30 #define DEFINES\_H\_

31

32 #include <stdint.h>

33 #include <inttypes.h>

34

35 // General

36 #define B\_LEN 131072

37 #define MAX\_FILENAME 300

38 #define bool int16\_t

39 #define true 1

40 #define false 0

42 #ifndef max

43  #define max(a, b) (((a) > (b)) ? (a) : (b))

44 #endif

45

46 #ifndef min

47  #define min(a, b) (((a) < (b)) ? (a) : (b))

48 #endif

49

50 #ifndef mem\_usageMB

51  #define mem\_usageMB() fprintf(stderr, \

52  "- Current allocated memory: %" PRIu64 "MB.\n", \

53  alloc\_mem >> 20)

54 #endif

55

56 #ifndef mem\_usage

57  #define mem\_usage() fprintf(stderr, \

58  "- Current allocated memory: %" PRIu64 "Bytes.\n", \

59  alloc\_mem)

60 #endif

61

62

63 // Q\_report, S\_report

64 #define DEFAULT\_MINQ 27

65 #define DEFAULT\_NTILES 96

66 #define DEFAULT\_NQ 46

67 #define ZEROQ 33

68 #define N\_ACGT 5

69 #define MAX\_RCOMMAND 4000

72 // Fasta files

73 #define FA\_ENTRY\_BUF 20

75 // Adapters

76 #define LOG\_4 0.60206

77 #define MIN\_NMATCHES 12

79 // Tree

80 #define T\_ACGT 4

81 #define NPOOL\_1D 1048576

82 #define NPOOL\_2D 16

83 #define MAX\_FASZ\_TREE 1e7

85 // BloomFilter

86 #define BITSPERCHAR 8

87 #define BASESPERCHAR 4

88 #define KMER\_LEN 25

89 #define FALSE\_POS\_RATE 0.05

90 #define ZERO\_POS\_RATE 1e-14

92 // Trimming

93 #define NO 0

94 #define ALL 1

95 #define ENDS 2

96 // trimN only

97 #define STRIP 3

98 // trimQ only

99 #define FRAC 3

100 #define ENDSFRAC 4

102 #define GLOBAL 5

104 #define TREE 1

105 #define BLOOM 2

107 #define ERROR 1000

109 #define DEFAULT\_MINL 25

112 // Classification of filters

113 #define ADAP 0

114 #define CONT 1

115 #define LOWQ 2

116 #define NNNN 3

117 #define GOOD 4

119 // Number of filters

120 #define NFILTERS 4

122 // Double stranded: classification of filters

123 #define ADAP2 5

124 #define CONT2 6

125 #define LOWQ2 7

126 #define NNNN2 8

127 #define GOOD2 9

129 // Double stranded: number of outputfiles

130 #define NFILES\_DS 10

132 #endif // endif DEFINES\_H\_

133


---

Generated on Mon Mar 19 2018 23:42:01 for FastqPuri by  

 1.8.14
