## Supplementary material for "FastqPuri: high-performance preprocessing of RNA-seq data": dir_000001_000000.html

FastqPuri: src -> include Relation


FastqPuri


- src

### src → include Relation

| File in src | Includes file in include |
| --- | --- |
| adapters.c | adapters.h |
| adapters.c | Lmer.h |
| bloom.c | bloom.h |
| city.c | city.h |
| fa\_read.c | defines.h |
| fa\_read.c | fa\_read.h |
| fa\_read.c | fopen\_gen.h |
| fopen\_gen.c | fopen\_gen.h |
| fq\_read.c | fq\_read.h |
| fq\_read.c | str\_manip.h |
| init\_makeBloom.c | init\_makeBloom.h |
| init\_makeBloom.c | str\_manip.h |
| init\_makeTree.c | init\_makeTree.h |
| init\_makeTree.c | str\_manip.h |
| init\_Qreport.c | defines.h |
| init\_Qreport.c | init\_Qreport.h |
| init\_Qreport.c | str\_manip.h |
| init\_Sreport.c | init\_Sreport.h |
| init\_Sreport.c | str\_manip.h |
| init\_trimFilter.c | init\_trimFilter.h |
| init\_trimFilter.c | str\_manip.h |
| init\_trimFilterDS.c | init\_trimFilterDS.h |
| init\_trimFilterDS.c | str\_manip.h |
| io\_trimFilter.c | defines.h |
| io\_trimFilter.c | io\_trimFilter.h |
| **io\_trimFilterDS.c** | defines.h |
| **io\_trimFilterDS.c** | io\_trimFilterDS.h |
| Lmer.c | Lmer.h |
| makeBloom.c | bloom.h |
| makeBloom.c | defines.h |
| makeBloom.c | fa\_read.h |
| makeBloom.c | init\_makeBloom.h |
| makeTree.c | defines.h |
| makeTree.c | fa\_read.h |
| makeTree.c | init\_makeTree.h |
| makeTree.c | tree.h |
| Qreport.c | fopen\_gen.h |
| Qreport.c | fq\_read.h |
| Qreport.c | init\_Qreport.h |
| Qreport.c | Rcommand\_Qreport.h |
| Qreport.c | stats\_info.h |
| Rcommand\_Qreport.c | defines.h |
| Rcommand\_Qreport.c | init\_Qreport.h |
| Rcommand\_Qreport.c | Rcommand\_Qreport.h |
| Rcommand\_Sreport.c | defines.h |
| Rcommand\_Sreport.c | init\_Sreport.h |
| Rcommand\_Sreport.c | Rcommand\_Sreport.h |
| Sreport.c | init\_Sreport.h |
| Sreport.c | Rcommand\_Sreport.h |
| stats\_info.c | init\_Qreport.h |
| stats\_info.c | stats\_info.h |
| stats\_info.c | str\_manip.h |
| str\_manip.c | str\_manip.h |
| struct\_trimFilter.c | struct\_trimFilter.h |
| tree.c | fopen\_gen.h |
| tree.c | Lmer.h |
| tree.c | tree.h |
| trim.c | defines.h |
| trim.c | str\_manip.h |
| trim.c | struct\_trimFilter.h |
| trim.c | trim.h |
| trimDS.c | Lmer.h |
| trimDS.c | struct\_trimFilter.h |
| trimDS.c | trim.h |
| trimDS.c | trimDS.h |
| trimFilter.c | bloom.h |
| trimFilter.c | defines.h |
| trimFilter.c | fopen\_gen.h |
| trimFilter.c | fq\_read.h |
| trimFilter.c | init\_trimFilter.h |
| trimFilter.c | io\_trimFilter.h |
| trimFilter.c | tree.h |
| trimFilter.c | trim.h |
| trimFilterDS.c | adapters.h |
| trimFilterDS.c | bloom.h |
| trimFilterDS.c | defines.h |
| trimFilterDS.c | fopen\_gen.h |
| trimFilterDS.c | fq\_read.h |
| trimFilterDS.c | init\_trimFilterDS.h |
| trimFilterDS.c | io\_trimFilterDS.h |
| trimFilterDS.c | Lmer.h |
| trimFilterDS.c | tree.h |
| trimFilterDS.c | trim.h |
| trimFilterDS.c | trimDS.h |


---

Generated on Mon Mar 19 2018 23:42:01 for FastqPuri by  

 1.8.14
