## Supplementary material for "FastqPuri: high-performance preprocessing of RNA-seq data": dir_68267d1309a1af8e8297ef4c3efbcdba.html

FastqPuri: src Directory Reference

FastqPuri

- src

src Directory Reference

Directory dependency graph for src:

|  |  |
| --- | --- |
| Files | |
| file | adapters.c |
|  | sequence manipulation for alignment |
| file | bloom.c |
|  | functions that implement the bloom filter |
| file | city.c |
|  | functions for hashin strings, C translation of cityhash (C++, google) |
| file | fa\_read.c |
|  | reads in and stores fasta files |
| file | fopen\_gen.c |
|  | Uncompress/compress input/output files using pipes. |
| file | fq\_read.c |
|  | fastq entries manipulations (read/write) |
| file | init\_makeBloom.c |
|  | Help dialog for makeBloom and initialization of the command line arguments. |
| file | init\_makeTree.c |
|  | Help dialog for makeTree and initialization of the command line arguments. |
| file | init\_Qreport.c |
|  | Help dialog for Qreport and initialization of the command line arguments. |
| file | init\_Sreport.c |
|  | Help dialog for Sreport and initialization of the command line arguments. |
| file | init\_trimFilter.c |
|  | help dialog for trimFilter and initialization of the command line arguments. |
| file | init\_trimFilterDS.c |
|  | help dialog for trimFilterDS and initialization of the command line arguments. |
| file | io\_trimFilter.c |
|  | buffer fq output, write summary file |
| file | Lmer.c |
|  | Manipulation of Lmers and sequences. |
| file | makeBloom.c |
|  | makeBloom main function |
| file | makeTree.c |
|  | makeTree main function |
| file | Qreport.c |
|  | QReport main function. |
| file | Rcommand\_Qreport.c |
|  | get Rscript command for Qreport |
| file | Rcommand\_Sreport.c |
|  | get Rscript command for Sreport |
| file | Sreport.c |
|  | Sreport main function. |
| file | stats\_info.c |
|  | Construct the quality report variables and update them. |
| file | str\_manip.c |
|  | functions that do string manipulation |
| file | struct\_trimFilter.c |
|  | function that frees the memory of parTF (structure storing the trimFilter/trimFilterDS input arguments). |
| file | tree.c |
|  | Construction of tree, check paths, write tree, read in tree. |
| file | trim.c |
|  | trims/filter sequences after Quality, N's contaminations. |
| file | trimDS.c |
|  | trim adapters from double stranded data |
| file | trimFilter.c |
|  | trimFilter main function |
| file | trimFilterDS.c |
|  | trimFilterDS main function |

---

Generated on Mon Mar 19 2018 23:42:01 for FastqPuri by  

 1.8.14
