## Supplementary material for "FastqPuri: high-performance preprocessing of RNA-seq data": dir_d44c64559bbebec7f509842c48db8b23.html

FastqPuri: include Directory Reference

FastqPuri

- include

include Directory Reference

|  |  |
| --- | --- |
| Files | |
| file | adapters.h [code] |
|  | sequence manipulation for alignment |
| file | bloom.h [code] |
|  | functions that implement the bloom filter |
| file | city.h [code] |
|  | functions for hashin strings, C translation of cityhash (C++, google) |
| file | citycrc.h [code] |
|  | functions for hashin strings, C translation of cityhash (C++, google) |
| file | defines.h [code] |
|  | Macro definitions. |
| file | fa\_read.h [code] |
|  | reads in and stores fasta files |
| file | fopen\_gen.h [code] |
|  | Uncompress/compress input/output files using pipes. |
| file | fq\_read.h [code] |
|  | fastq entries manipulations (read/write) |
| file | init\_makeBloom.h [code] |
|  | Help dialog for makeBloom and initialization of the command line arguments. |
| file | init\_makeTree.h [code] |
|  | Help dialog for makeTree and initialization of the command line arguments. |
| file | init\_Qreport.h [code] |
|  | Header file: help dialog for Qreport and initialization of the command line arguments. |
| file | init\_Sreport.h [code] |
|  | Help dialog for Sreport and initialization of the command line arguments. |
| file | init\_trimFilter.h [code] |
|  | help dialog for trimFilter and initialization of the command line arguments. |
| file | init\_trimFilterDS.h [code] |
|  | help dialog for trimFilterDS and initialization of the command line arguments. |
| file | io\_trimFilter.h [code] |
|  | buffer fq output, write summary file |
| file | io\_trimFilterDS.h [code] |
|  | buffer fq output, write summary file |
| file | Lmer.h [code] |
|  | Manipulation of Lmers and sequences. |
| file | Rcommand\_Qreport.h [code] |
|  | get Rscript command for Qreport |
| file | Rcommand\_Sreport.h [code] |
|  | get Rscript command for Sreport |
| file | stats\_info.h [code] |
|  | Construct the quality report variables and update them. |
| file | str\_manip.h [code] |
|  | functions that do string manipulation |
| file | struct\_trimFilter.h [code] |
|  | structure where the input arguments of trimFilter and trimFilterDS will be stored and function to free the memory of it. |
| file | tree.h [code] |
|  | Construction of tree, check paths, write tree, read in tree. |
| file | trim.h [code] |
|  | trims/filter sequences after Quality, N's contaminations. |
| file | trimDS.h [code] |
|  | trim adapters from double stranded data |

---

Generated on Mon Mar 19 2018 23:42:01 for FastqPuri by  

 1.8.14
