## Supplementary material for "FastqPuri: high-performance preprocessing of RNA-seq data": dynsections.js

```
/*
 @licstart  The following is the entire license notice for the
 JavaScript code in this file.

 Copyright (C) 1997-2017 by Dimitri van Heesch

 This program is free software; you can redistribute it and/or modify
 it under the terms of the GNU General Public License as published by
 the Free Software Foundation; either version 2 of the License, or
 (at your option) any later version.

 This program is distributed in the hope that it will be useful,
 but WITHOUT ANY WARRANTY; without even the implied warranty of
 MERCHANTABILITY or FITNESS FOR A PARTICULAR PURPOSE.  See the
 GNU General Public License for more details.

 You should have received a copy of the GNU General Public License along
 with this program; if not, write to the Free Software Foundation, Inc.,
 51 Franklin Street, Fifth Floor, Boston, MA 02110-1301 USA.

 @licend  The above is the entire license notice
 for the JavaScript code in this file
 */
function toggleVisibility(linkObj)
{
 var base = $(linkObj).attr('id');
 var summary = $('#'+base+'-summary');
 var content = $('#'+base+'-content');
 var trigger = $('#'+base+'-trigger');
 var src=$(trigger).attr('src');
 if (content.is(':visible')===true) {
   content.hide();
   summary.show();
   $(linkObj).addClass('closed').removeClass('opened');
   $(trigger).attr('src',src.substring(0,src.length-8)+'closed.png');
 } else {
   content.show();
   summary.hide();
   $(linkObj).removeClass('closed').addClass('opened');
   $(trigger).attr('src',src.substring(0,src.length-10)+'open.png');
 }
 return false;
}

function updateStripes()
{
  $('table.directory tr').
       removeClass('even').filter(':visible:even').addClass('even');
}

function toggleLevel(level)
{
  $('table.directory tr').each(function() {
    var l = this.id.split('_').length-1;
    var i = $('#img'+this.id.substring(3));
    var a = $('#arr'+this.id.substring(3));
    if (l<level+1) {
      i.removeClass('iconfopen iconfclosed').addClass('iconfopen');
      a.html('&#9660;');
      $(this).show();
    } else if (l==level+1) {
      i.removeClass('iconfclosed iconfopen').addClass('iconfclosed');
      a.html('&#9654;');
      $(this).show();
    } else {
      $(this).hide();
    }
  });
  updateStripes();
}

function toggleFolder(id)
{
  // the clicked row
  var currentRow = $('#row_'+id);

  // all rows after the clicked row
  var rows = currentRow.nextAll("tr");

  var re = new RegExp('^row_'+id+'\\d+_$', "i"); //only one sub

  // only match elements AFTER this one (can't hide elements before)
  var childRows = rows.filter(function() { return this.id.match(re); });

  // first row is visible we are HIDING
  if (childRows.filter(':first').is(':visible')===true) {
    // replace down arrow by right arrow for current row
    var currentRowSpans = currentRow.find("span");
    currentRowSpans.filter(".iconfopen").removeClass("iconfopen").addClass("iconfclosed");
    currentRowSpans.filter(".arrow").html('&#9654;');
    rows.filter("[id^=row_"+id+"]").hide(); // hide all children
  } else { // we are SHOWING
    // replace right arrow by down arrow for current row
    var currentRowSpans = currentRow.find("span");
    currentRowSpans.filter(".iconfclosed").removeClass("iconfclosed").addClass("iconfopen");
    currentRowSpans.filter(".arrow").html('&#9660;');
    // replace down arrows by right arrows for child rows
    var childRowsSpans = childRows.find("span");
    childRowsSpans.filter(".iconfopen").removeClass("iconfopen").addClass("iconfclosed");
    childRowsSpans.filter(".arrow").html('&#9654;');
    childRows.show(); //show all children
  }
  updateStripes();
}


function toggleInherit(id)
{
  var rows = $('tr.inherit.'+id);
  var img = $('tr.inherit_header.'+id+' img');
  var src = $(img).attr('src');
  if (rows.filter(':first').is(':visible')===true) {
    rows.css('display','none');
    $(img).attr('src',src.substring(0,src.length-8)+'closed.png');
  } else {
    rows.css('display','table-row'); // using show() causes jump in firefox
    $(img).attr('src',src.substring(0,src.length-10)+'open.png');
  }
}
/* @license-end */
```
