## Supplementary material for "FastqPuri: high-performance preprocessing of RNA-seq data": fa__read_8c.html

FastqPuri: src/fa\_read.c File Reference

FastqPuri

- src

Functions |
Variables

fa\_read.c File Reference

reads in and stores fasta files
More...

`#include <stdlib.h>`  
`#include <string.h>`  
`#include "fa_read.h"`  
`#include "defines.h"`  
`#include "fopen_gen.h"`

Include dependency graph for fa\_read.c:

|  |  |
| --- | --- |
| Functions | |
| static int | ignore\_line (char \*line) |
|  | ignore header lines. More... |
| static void | init\_fa (Fa\_data \*ptr\_fa) |
|  | Initialization of Fa\_data. More... |
| static void | realloc\_fa (Fa\_data \*ptr\_fa) |
|  | Reallocation of Fa\_data, in case the length of entrylen is exhausted. More... |
| static void | init\_entries (Fa\_data \*ptr\_fa) |
|  | Allocation of Fa\_entries. More... |
| static uint64\_t | sweep\_fa (char \*filename, Fa\_data \*ptr\_fa) |
|  | this function sweeps a fasta file to obtain structure details. More... |
| int | read\_fasta (char \*filename, Fa\_data \*ptr\_fa) |
|  | reads a fasta file and stores the contents in a Fa\_data structure. More... |
| uint64\_t | size\_fasta (Fa\_data \*ptr\_fa) |
|  | computes length of genome in fasta structure More... |
| uint64\_t | nkmers (Fa\_data \*ptr\_fa, int kmersize) |
|  | number of kmers of length kmersize contained in a fasta structure More... |
| void | free\_fasta (Fa\_data \*ptr\_fa) |
|  | free fasta file More... |

|  |  |
| --- | --- |
| Variables | |
| uint64\_t | alloc\_mem |

### Detailed Description

reads in and stores fasta files

Author
:   Paula Perez paula.nosp@@m.o@@m.om

Date
:   18.08.2017

### Function Documentation

### ◆ free\_fasta()

|  |  |  |  |  |
| --- | --- | --- | --- | --- |
| void free\_fasta | ( | Fa\_data \* | *ptr\_fa* | ) |

free fasta file

Parameters
:   |  |  |
    | --- | --- |
    | ptr\_fa | pointer to Fa\_data structure. |

The dynamically allocated memory in a Fa\_data struct is deallocated and counted, so that we can

### ◆ ignore\_line()

|  |  |  |  |  |
| --- | --- | --- | --- | --- |
| static int ignore\_line | ( | char \* | *line* | ) |

static

ignore header lines.

Parameters
:   |  |  |
    | --- | --- |
    | line | string of characters. |

Returns
:   number of characters to jump until a   
    is found.

### ◆ init\_entries()

|  |  |  |  |  |
| --- | --- | --- | --- | --- |
| static void init\_entries | ( | Fa\_data \* | *ptr\_fa* | ) |

static

Allocation of Fa\_entries.

Parameters
:   |  |  |
    | --- | --- |
    | ptr\_fa | pointer to Fa\_data structure. |

When we have sweeped the fasta file once, we can proceed to allocate the memory for the entries (now we have registered their length).

### ◆ init\_fa()

|  |  |  |  |  |
| --- | --- | --- | --- | --- |
| static void init\_fa | ( | Fa\_data \* | *ptr\_fa* | ) |

static

Initialization of Fa\_data.

Parameters
:   |  |  |
    | --- | --- |
    | ptr\_fa | pointer to Fa\_data structure. |

Initializes nlines, linelen, nentries to 0 and allocates memory for entrylen (FA\_ENTRY\_BUF entries).

### ◆ nkmers()

|  |  |  |  |
| --- | --- | --- | --- |
| uint64\_t nkmers | ( | Fa\_data \* | *ptr\_fa*, |
|  |  | int | *kmersize* |
|  | ) |  |  |

number of kmers of length kmersize contained in a fasta structure

Returns
:   number of kmers of length kmersize contained in a fasta structure

### ◆ read\_fasta()

|  |  |  |  |
| --- | --- | --- | --- |
| int read\_fasta | ( | char \* | *filename*, |
|  |  | Fa\_data \* | *ptr\_fa* |
|  | ) |  |  |

reads a fasta file and stores the contents in a Fa\_data structure.

Parameters
:   |  |  |
    | --- | --- |
    | filename | path to a fasta input file. |
    | ptr\_fa | pointer to Fa\_data structure. |

Returns
:   number of entries in the fasta file.

A fasta file is read and stored in a structure Fa\_data The basic problem with reading FASTA files is that there is no end-of-record indicator. When you're reading sequence n, you don't know you're done until you've read the header line for sequence n+1, which you won't parse 'til later (when you're reading in the sequence n+1). The solution implemented here is to read the file twice. The first time, (sweep\_fa), we initialize Fa\_data and store the parameters:

- nlines: number of lines of the fasta file.
- nentries: number of entries in the fasta file.
- linelen: length of a line in the considered fasta file.
- entrylen: array containing the lengths of every entry. With this information, the pointer to Fa\_entry can be allocated and the file is read again and the entries are stored in the structure.

### ◆ realloc\_fa()

|  |  |  |  |  |
| --- | --- | --- | --- | --- |
| static void realloc\_fa | ( | Fa\_data \* | *ptr\_fa* | ) |

static

Reallocation of Fa\_data, in case the length of entrylen is exhausted.

Parameters
:   |  |  |
    | --- | --- |
    | ptr\_fa | pointer to Fa\_data structure. |

### ◆ size\_fasta()

|  |  |  |  |  |
| --- | --- | --- | --- | --- |
| uint64\_t size\_fasta | ( | Fa\_data \* | *ptr\_fa* | ) |

computes length of genome in fasta structure

Parameters
:   |  |  |
    | --- | --- |
    | ptr\_fa | pointer to Fa\_data |

Returns
:   total number of nucleotides

### ◆ sweep\_fa()

|  |  |  |  |
| --- | --- | --- | --- |
| static uint64\_t sweep\_fa | ( | char \* | *filename*, |
|  |  | Fa\_data \* | *ptr\_fa* |
|  | ) |  |  |

static

this function sweeps a fasta file to obtain structure details.

Parameters
:   |  |  |
    | --- | --- |
    | filename | path to a fasta input file. |
    | ptr\_fa | pointer to Fa\_data structure. |

Returns
:   size of fasta file.

This function sweeps over the fasta file once to annotate how many entries there are, how long they are, how many characters there are per line, and how many lines the file has.

### Variable Documentation

### ◆ alloc\_mem

uint64\_t alloc\_mem

global variable. Memory allocated in the heap.

---

Generated on Mon Mar 19 2018 23:42:01 for FastqPuri by  

 1.8.14
