## Supplementary material for "FastqPuri: high-performance preprocessing of RNA-seq data": fa__read_8h_source.html

FastqPuri: include/fa\_read.h Source File


FastqPuri


- include

fa\_read.h

Go to the documentation of this file.

1 /\*\*\*\*\*\*\*\*\*\*\*\*\*\*\*\*\*\*\*\*\*\*\*\*\*\*\*\*\*\*\*\*\*\*\*\*\*\*\*\*\*\*\*\*\*\*\*\*\*\*\*\*\*\*\*\*\*\*\*\*\*\*\*\*\*\*\*\*\*\*\*\*\*\*\*\*

2  \* Copyright (C) 2017 by Paula Perez Rubio \*

3  \* \*

4  \* This file is part of FastqPuri. \*

5  \* \*

6  \* FastqPuri is free software: you can redistribute it and/or modify \*

7  \* it under the terms of the GNU General Public License as \*

8  \* published by the Free Software Foundation, either version 3 of the \*

9  \* License, or (at your option) any later version. \*

10  \* \*

11  \* FastqPuri is distributed in the hope that it will be useful, \*

12  \* but WITHOUT ANY WARRANTY; without even the implied warranty of \*

13  \* MERCHANTABILITY or FITNESS FOR A PARTICULAR PURPOSE. See the \*

14  \* GNU General Public License for more details. \*

15  \* \*

16  \* You should have received a copy of the GNU General Public License \*

17  \* along with FastqPuri. \*

18  \* If not, see <http://www.gnu.org/licenses/>. \*

19  \*\*\*\*\*\*\*\*\*\*\*\*\*\*\*\*\*\*\*\*\*\*\*\*\*\*\*\*\*\*\*\*\*\*\*\*\*\*\*\*\*\*\*\*\*\*\*\*\*\*\*\*\*\*\*\*\*\*\*\*\*\*\*\*\*\*\*\*\*\*\*\*\*\*\*\*/

20

30 #ifndef FA\_READ\_H\_

31 #define FA\_READ\_H\_

32

33 #include <stdint.h>

34

38 typedef struct \_fa\_entry {

39  uint64\_t N;

40  char \*seq;

41 } Fa\_entry;

42

46 typedef struct \_fa\_data {

47  uint64\_t nlines;

48  int nentries;

49  int linelen;

50  uint64\_t \*entrylen;

51  Fa\_entry \*entry;

52 } Fa\_data;

53

54 int read\_fasta(char \*filename, Fa\_data \*ptr\_fa);

55 uint64\_t size\_fasta(Fa\_data \*ptr\_fa);

56 uint64\_t nkmers(Fa\_data \*ptr\_fa, int kmersize);

57 void free\_fasta(Fa\_data \*ptr\_fa);

58

59 // static functions:

60 // static int ignore\_line(char \*line)

61 // static void init\_fa(Fa\_data \*ptr\_fa)

62 // static voiid realloc\_fa(Fa\_data \*ptr\_fa)

63 // static voiid init\_entries(Fa\_data \*ptr\_fa)

64 // static uint64\_t swee\_fa(char \*filename, Fa\_data \*ptr\_fa)

65

66

67 #endif // endif FA\_READ\_H\_

\_fa\_entry::N

uint64\_t N

**Definition:** fa\_read.h:39

\_fa\_data::linelen

int linelen

**Definition:** fa\_read.h:49

\_fa\_data

stores sequences of a fasta file

**Definition:** fa\_read.h:46

\_fa\_data::entry

Fa\_entry \* entry

**Definition:** fa\_read.h:51

\_fa\_data::entrylen

uint64\_t \* entrylen

**Definition:** fa\_read.h:50

\_fa\_data::nlines

uint64\_t nlines

**Definition:** fa\_read.h:47

read\_fasta

int read\_fasta(char \*filename, Fa\_data \*ptr\_fa)

reads a fasta file and stores the contents in a Fa\_data structure.

**Definition:** fa\_read.c:212

Fa\_entry

struct \_fa\_entry Fa\_entry

fasta entry

free\_fasta

void free\_fasta(Fa\_data \*ptr\_fa)

free fasta file

**Definition:** fa\_read.c:311

size\_fasta

uint64\_t size\_fasta(Fa\_data \*ptr\_fa)

computes length of genome in fasta structure

**Definition:** fa\_read.c:281

\_fa\_entry::seq

char \* seq

**Definition:** fa\_read.h:40

\_fa\_entry

fasta entry

**Definition:** fa\_read.h:38

\_fa\_data::nentries

int nentries

**Definition:** fa\_read.h:48

nkmers

uint64\_t nkmers(Fa\_data \*ptr\_fa, int kmersize)

number of kmers of length kmersize contained in a fasta structure

**Definition:** fa\_read.c:295

Fa\_data

struct \_fa\_data Fa\_data

stores sequences of a fasta file


---

Generated on Mon Mar 19 2018 23:42:01 for FastqPuri by  

 1.8.14
