## Supplementary material for "FastqPuri: high-performance preprocessing of RNA-seq data": files.html

FastqPuri: File List


FastqPuri


File List

Here is a list of all documented files with brief descriptions:

[detail level 12]

|  |  |
| --- | --- |
| ▼ include |  |
| adapters.h | Sequence manipulation for alignment |
| bloom.h | Functions that implement the bloom filter |
| city.h | Functions for hashin strings, C translation of cityhash (C++, google) |
| citycrc.h | Functions for hashin strings, C translation of cityhash (C++, google) |
| defines.h | Macro definitions |
| fa\_read.h | Reads in and stores fasta files |
| fopen\_gen.h | Uncompress/compress input/output files using pipes |
| fq\_read.h | Fastq entries manipulations (read/write) |
| init\_makeBloom.h | Help dialog for makeBloom and initialization of the command line arguments |
| init\_makeTree.h | Help dialog for makeTree and initialization of the command line arguments |
| init\_Qreport.h | Header file: help dialog for Qreport and initialization of the command line arguments |
| init\_Sreport.h | Help dialog for Sreport and initialization of the command line arguments |
| init\_trimFilter.h | Help dialog for trimFilter and initialization of the command line arguments |
| init\_trimFilterDS.h | Help dialog for trimFilterDS and initialization of the command line arguments |
| io\_trimFilter.h | Buffer fq output, write summary file |
| io\_trimFilterDS.h | Buffer fq output, write summary file |
| Lmer.h | Manipulation of Lmers and sequences |
| Rcommand\_Qreport.h | Get Rscript command for Qreport |
| Rcommand\_Sreport.h | Get Rscript command for Sreport |
| stats\_info.h | Construct the quality report variables and update them |
| str\_manip.h | Functions that do string manipulation |
| struct\_trimFilter.h | Structure where the input arguments of trimFilter and trimFilterDS will be stored and function to free the memory of it |
| tree.h | Construction of tree, check paths, write tree, read in tree |
| trim.h | Trims/filter sequences after Quality, N's contaminations |
| trimDS.h | Trim adapters from double stranded data |
| ▼ src |  |
| adapters.c | Sequence manipulation for alignment |
| bloom.c | Functions that implement the bloom filter |
| city.c | Functions for hashin strings, C translation of cityhash (C++, google) |
| fa\_read.c | Reads in and stores fasta files |
| fopen\_gen.c | Uncompress/compress input/output files using pipes |
| fq\_read.c | Fastq entries manipulations (read/write) |
| init\_makeBloom.c | Help dialog for makeBloom and initialization of the command line arguments |
| init\_makeTree.c | Help dialog for makeTree and initialization of the command line arguments |
| init\_Qreport.c | Help dialog for Qreport and initialization of the command line arguments |
| init\_Sreport.c | Help dialog for Sreport and initialization of the command line arguments |
| init\_trimFilter.c | Help dialog for trimFilter and initialization of the command line arguments |
| init\_trimFilterDS.c | Help dialog for trimFilterDS and initialization of the command line arguments |
| io\_trimFilter.c | Buffer fq output, write summary file |
| Lmer.c | Manipulation of Lmers and sequences |
| makeBloom.c | MakeBloom main function |
| makeTree.c | MakeTree main function |
| Qreport.c | QReport main function |
| Rcommand\_Qreport.c | Get Rscript command for Qreport |
| Rcommand\_Sreport.c | Get Rscript command for Sreport |
| Sreport.c | Sreport main function |
| stats\_info.c | Construct the quality report variables and update them |
| str\_manip.c | Functions that do string manipulation |
| struct\_trimFilter.c | Function that frees the memory of parTF (structure storing the trimFilter/trimFilterDS input arguments) |
| tree.c | Construction of tree, check paths, write tree, read in tree |
| trim.c | Trims/filter sequences after Quality, N's contaminations |
| trimDS.c | Trim adapters from double stranded data |
| trimFilter.c | TrimFilter main function |
| trimFilterDS.c | TrimFilterDS main function |


---

Generated on Mon Mar 19 2018 23:42:01 for FastqPuri by  

 1.8.14
