## Supplementary material for "FastqPuri: high-performance preprocessing of RNA-seq data": fopen__gen_8c.html

FastqPuri: src/fopen\_gen.c File Reference

FastqPuri

- src

Functions

fopen\_gen.c File Reference

Uncompress/compress input/output files using pipes.
More...

`#include <stdlib.h>`  
`#include <string.h>`  
`#include <unistd.h>`  
`#include <assert.h>`  
`#include <sys/types.h>`  
`#include <fcntl.h>`  
`#include "fopen_gen.h"`

Include dependency graph for fopen\_gen.c:

|  |  |
| --- | --- |
| Functions | |
| static const char \* | **zcatExec** (const char \*path) |
| static const char \* | catExec (const char \*path) |
|  | Commands to compress files. To be done in output. |
| static int | uncompress (const char \*path) |
|  | Open a pipe to uncompress file. Open a pipe to uncompress the specified file. Not thread safe. More... |
| static int | compress (const char \*path) |
|  | Open a pipe to compress output. Open a pipe to uncompress the specified file. Not thread safe. More... |
| int | **setCloexec** (int fd) |
| static FILE \* | funcompress (const char \*path) |
|  | Open a pipe to uncompress the specified file. More... |
| static FILE \* | fcompress (const char \*path) |
|  | Open a pipe to compress the specified file. More... |
| FILE \* | fopen\_gen (const char \*path, const char \*mode) |
|  | Generalized fopen function. fopen\_gen is to be used as fopen. Can be used in read and in write mode. When used in read mode with a compressed extension, the file will be first decompressed and then read. When used in write mode with a compressed extension, the output will be compressed. More... |

### Detailed Description

Uncompress/compress input/output files using pipes.

Hook the standard file opening functions, open, fopen and fopen64. If the extension of the file being opened indicates the file is compressed (.gz, .bz2, .xz), when opening in the reading mode a pipe to a program is opened that decompresses that file (gunzip, bunzip2 or xzdec) and return a handle to the open pipe. When opening in the writing mode (only for .gz, .bam), a pipe to a program is opened that compresses the output.

Author
:   Paula Perez paula.nosp@@m.o@@m.om

Date
:   03.08.2017

Warning
:   vfork vs fork to be checked!

Note
:   - original copyright note - (reading mode, original C++ code) author: Shaun Jackman sjack.nosp@m.man@.nosp@m.bcgsc.nosp@m..ca, https://github.com/bcgsc,   
    filename: Uncompress.cpp

### Function Documentation

### ◆ compress()

|  |  |  |  |  |
| --- | --- | --- | --- | --- |
| static int compress | ( | const char \* | *path* | ) |

static

Open a pipe to compress output. Open a pipe to uncompress the specified file. Not thread safe.

Returns
:   a file descriptor

### ◆ fcompress()

|  |  |  |  |  |
| --- | --- | --- | --- | --- |
| static FILE\* fcompress | ( | const char \* | *path* | ) |

static

Open a pipe to compress the specified file.

Returns
:   a FILE pointer

### ◆ fopen\_gen()

|  |  |  |  |
| --- | --- | --- | --- |
| FILE\* fopen\_gen | ( | const char \* | *path*, |
|  |  | const char \* | *mode* |
|  | ) |  |  |

Generalized fopen function. fopen\_gen is to be used as fopen. Can be used in read and in write mode. When used in read mode with a compressed extension, the file will be first decompressed and then read. When used in write mode with a compressed extension, the output will be compressed.

Returns
:   a FILE pointer

### ◆ funcompress()

|  |  |  |  |  |
| --- | --- | --- | --- | --- |
| static FILE\* funcompress | ( | const char \* | *path* | ) |

static

Open a pipe to uncompress the specified file.

Returns
:   a FILE pointer

### ◆ uncompress()

|  |  |  |  |  |
| --- | --- | --- | --- | --- |
| static int uncompress | ( | const char \* | *path* | ) |

static

Open a pipe to uncompress file. Open a pipe to uncompress the specified file. Not thread safe.

Returns
:   a file descriptor

---

Generated on Mon Mar 19 2018 23:42:01 for FastqPuri by  

 1.8.14
