## Supplementary material for "FastqPuri: high-performance preprocessing of RNA-seq data": fopen__gen_8h_source.html

FastqPuri: include/fopen\_gen.h Source File


FastqPuri


- include

fopen\_gen.h

Go to the documentation of this file.

1 /\*\*\*\*\*\*\*\*\*\*\*\*\*\*\*\*\*\*\*\*\*\*\*\*\*\*\*\*\*\*\*\*\*\*\*\*\*\*\*\*\*\*\*\*\*\*\*\*\*\*\*\*\*\*\*\*\*\*\*\*\*\*\*\*\*\*\*\*\*\*\*\*\*\*\*\*

2  \* Copyright (C) 2017 by Paula Perez Rubio \*

3  \* \*

4  \* This file is part of FastqPuri. \*

5  \* \*

6  \* FastqPuri is free software: you can redistribute it and/or modify \*

7  \* it under the terms of the GNU General Public License as \*

8  \* published by the Free Software Foundation, either version 3 of the \*

9  \* License, or (at your option) any later version. \*

10  \* \*

11  \* FastqPuri is distributed in the hope that it will be useful, \*

12  \* but WITHOUT ANY WARRANTY; without even the implied warranty of \*

13  \* MERCHANTABILITY or FITNESS FOR A PARTICULAR PURPOSE. See the \*

14  \* GNU General Public License for more details. \*

15  \* \*

16  \* You should have received a copy of the GNU General Public License \*

17  \* along with FastqPuri. \*

18  \* If not, see <http://www.gnu.org/licenses/>. \*

19  \*\*\*\*\*\*\*\*\*\*\*\*\*\*\*\*\*\*\*\*\*\*\*\*\*\*\*\*\*\*\*\*\*\*\*\*\*\*\*\*\*\*\*\*\*\*\*\*\*\*\*\*\*\*\*\*\*\*\*\*\*\*\*\*\*\*\*\*\*\*\*\*\*\*\*\*/

20

42 #ifndef FOPEN\_GEN\_H\_

43 #define FOPEN\_GEN\_H\_

44

45 #define READ\_END 0

46 #define WRITE\_END 1

47 #define PERMISSIONS 0640

48

49 #include <stdio.h>

50

51 #ifdef \_\_STDC\_\_

52 FILE\* fdopen(int, const char\*);

53 #endif

54

55 int setCloexec(int fd);

56 FILE\* fopen\_gen(const char \*path, const char \* mode);

57

70 #endif // FOPEN\_GEN\_H\_

fopen\_gen

FILE \* fopen\_gen(const char \*path, const char \*mode)

Generalized fopen function. fopen\_gen is to be used as fopen. Can be used in read and in write mode...

**Definition:** fopen\_gen.c:241


---

Generated on Mon Mar 19 2018 23:42:01 for FastqPuri by  

 1.8.14
