## Supplementary material for "FastqPuri: high-performance preprocessing of RNA-seq data": Lmer_8c.html

FastqPuri: src/Lmer.c File Reference

FastqPuri

- src

Functions |
Variables

Lmer.c File Reference

Manipulation of Lmers and sequences.
More...

`#include "Lmer.h"`  
`#include <string.h>`  
`#include <stdint.h>`  
`#include <stdio.h>`

Include dependency graph for Lmer.c:

|  |  |
| --- | --- |
| Functions | |
| void | init\_map () |
|  | Initialize lookup table fw\_1B. More... |
| void | Lmer\_sLmer (char \*Lmer, int L) |
|  | Transforms an Lmer to the convention stored in the lookup table fw\_1B. |
| void | rev\_comp (char \*sLmer, int L) |
|  | Obtains the reverse complement, for {'\000','\001','\002','\003'}. |

|  |  |
| --- | --- |
| Variables | |
| uint8\_t | fw\_1B [256] |
| uint8\_t | bw\_1B [256] |
| uint8\_t | Nencode |

### Detailed Description

Manipulation of Lmers and sequences.

Author
:   Paula Perez paula.nosp@@m.o@@m.om

Date
:   18.08.2017

### Function Documentation

### ◆ init\_map()

|  |  |  |  |
| --- | --- | --- | --- |
| void init\_map | ( |  | ) |

Initialize lookup table fw\_1B.

{'a','c','g','t'} –> {'\000','\001','\002','\003'}, rest '\004'.

### Variable Documentation

## ◆ bw\_1B

uint8\_t bw\_1B[256]

global variable. Lookup table.

## ◆ fw\_1B

uint8\_t fw\_1B[256]

global variable. Lookup table.

### ◆ Nencode

uint8\_t Nencode

global variable. Encoding for N's(\004)

---

Generated on Mon Mar 19 2018 23:42:01 for FastqPuri by  

 1.8.14
