## Supplementary material for "FastqPuri: high-performance preprocessing of RNA-seq data": Lmer_8h.html

FastqPuri: include/Lmer.h File Reference

FastqPuri

- include

Functions

Lmer.h File Reference

Manipulation of Lmers and sequences.
More...

This graph shows which files directly or indirectly include this file:

Go to the source code of this file.

|  |  |
| --- | --- |
| Functions | |
| void | init\_map () |
|  | Initialize lookup table fw\_1B. More... |
| void | Lmer\_sLmer (char \*Lmer, int L) |
|  | Transforms an Lmer to the convention stored in the lookup table fw\_1B. |
| void | rev\_comp (char \*sLmer, int L) |
|  | Obtains the reverse complement, for {'\000','\001','\002','\003'}. |

### Detailed Description

Manipulation of Lmers and sequences.

Author
:   Paula Perez paula.nosp@@m.o@@m.om

Date
:   18.08.2017

Note
:   I have to try to merge the two versions of conversions!   
    Basically, and depending on the method used, nucleotides {'a', 'c', 'g', 't'} are shifted to the characters {'\000','\001','\002','\003'} or to {'\001','\002','\003','\004'} in a Lmer. A function to provide the reverse complement is also provided.

### Function Documentation

### ◆ init\_map()

|  |  |  |  |
| --- | --- | --- | --- |
| void init\_map | ( |  | ) |

Initialize lookup table fw\_1B.

{'a','c','g','t'} –> {'\000','\001','\002','\003'}, rest '\004'.

---

Generated on Mon Mar 19 2018 23:42:01 for FastqPuri by  

 1.8.14
