## Supplementary material for "FastqPuri: high-performance preprocessing of RNA-seq data": Qreport_8c.html

FastqPuri: src/Qreport.c File Reference

FastqPuri

- src

Functions |
Variables

Qreport.c File Reference

QReport main function.
More...

`#include <stdio.h>`  
`#include <stdlib.h>`  
`#include <string.h>`  
`#include <time.h>`  
`#include "init_Qreport.h"`  
`#include "fopen_gen.h"`  
`#include "fq_read.h"`  
`#include "stats_info.h"`  
`#include "Rcommand_Qreport.h"`

Include dependency graph for Qreport.c:

|  |  |
| --- | --- |
| Functions | |
| int | main (int argc, char \*argv[]) |
|  | Qreport main function. |

|  |  |
| --- | --- |
| Variables | |
| Iparam\_Qreport | par\_QR |

### Detailed Description

QReport main function.

Author
:   Paula Perez paula.nosp@@m.o@@m.om

Date
:   03.08.2017 This file contains the quality report main function. It reads a fastq file and creates a html quality report. See README\_Qreport.md for more details.

### Variable Documentation

### ◆ par\_QR

Iparam\_Qreport par\_QR

global variable: input parameters for Qreport

---

Generated on Mon Mar 19 2018 23:42:01 for FastqPuri by  

 1.8.14
