## Supplementary material for "FastqPuri: high-performance preprocessing of RNA-seq data": Rcommand__Qreport_8c.html

FastqPuri: src/Rcommand\_Qreport.c File Reference

FastqPuri

- src

Functions |
Variables

Rcommand\_Qreport.c File Reference

get Rscript command for Qreport
More...

`#include <stdio.h>`  
`#include <stdlib.h>`  
`#include <unistd.h>`  
`#include "Rcommand_Qreport.h"`  
`#include "init_Qreport.h"`  
`#include "defines.h"`  
`#include "config.h"`

Include dependency graph for Rcommand\_Qreport.c:

|  |  |
| --- | --- |
| Functions | |
| char \* | command\_Qreport () |
|  | returns Rscript command that generates the quality report in html |

|  |  |
| --- | --- |
| Variables | |
| Iparam\_Qreport | par\_QR |

### Detailed Description

get Rscript command for Qreport

Author
:   Paula Perez paula.nosp@@m.o@@m.om

Date
:   07.08.2017

### Variable Documentation

### ◆ par\_QR

Iparam\_Qreport par\_QR

input parameters Qreport

global variable: input parameters for Qreport

---

Generated on Mon Mar 19 2018 23:42:01 for FastqPuri by  

 1.8.14
