## Supplementary material for "FastqPuri: high-performance preprocessing of RNA-seq data": Rcommand__Qreport_8h.html

FastqPuri: include/Rcommand\_Qreport.h File Reference

FastqPuri

- include

Functions

Rcommand\_Qreport.h File Reference

get Rscript command for Qreport
More...

This graph shows which files directly or indirectly include this file:

Go to the source code of this file.

|  |  |
| --- | --- |
| Functions | |
| char \* | command\_Qreport () |
|  | returns Rscript command that generates the quality report in html |

### Detailed Description

get Rscript command for Qreport

Author
:   Paula Perez paula.nosp@@m.o@@m.om

Date
:   09.08.2017

---

Generated on Mon Mar 19 2018 23:42:01 for FastqPuri by  

 1.8.14
