## Supplementary material for "FastqPuri: high-performance preprocessing of RNA-seq data": Rcommand__Qreport_8h_source.html

FastqPuri: include/Rcommand\_Qreport.h Source File


FastqPuri


- include

Rcommand\_Qreport.h

Go to the documentation of this file.

1 /\*\*\*\*\*\*\*\*\*\*\*\*\*\*\*\*\*\*\*\*\*\*\*\*\*\*\*\*\*\*\*\*\*\*\*\*\*\*\*\*\*\*\*\*\*\*\*\*\*\*\*\*\*\*\*\*\*\*\*\*\*\*\*\*\*\*\*\*\*\*\*\*\*\*\*\*

2  \* Copyright (C) 2017 by Paula Perez Rubio \*

3  \* \*

4  \* This file is part of FastqPuri. \*

5  \* \*

6  \* FastqPuri is free software: you can redistribute it and/or modify \*

7  \* it under the terms of the GNU General Public License as \*

8  \* published by the Free Software Foundation, either version 3 of the \*

9  \* License, or (at your option) any later version. \*

10  \* \*

11  \* FastqPuri is distributed in the hope that it will be useful, \*

12  \* but WITHOUT ANY WARRANTY; without even the implied warranty of \*

13  \* MERCHANTABILITY or FITNESS FOR A PARTICULAR PURPOSE. See the \*

14  \* GNU General Public License for more details. \*

15  \* \*

16  \* You should have received a copy of the GNU General Public License \*

17  \* along with FastqPuri. \*

18  \* If not, see <http://www.gnu.org/licenses/>. \*

19  \*\*\*\*\*\*\*\*\*\*\*\*\*\*\*\*\*\*\*\*\*\*\*\*\*\*\*\*\*\*\*\*\*\*\*\*\*\*\*\*\*\*\*\*\*\*\*\*\*\*\*\*\*\*\*\*\*\*\*\*\*\*\*\*\*\*\*\*\*\*\*\*\*\*\*\*/

20

30 #ifndef RCOMMAND\_QREPORT\_H\_

31 #define RCOMMAND\_QREPORT\_H\_

32

33 char \*command\_Qreport();

34

35 #endif // endif RCOMMAND\_QREPORT\_H\_

command\_Qreport

char \* command\_Qreport()

returns Rscript command that generates the quality report in html

**Definition:** Rcommand\_Qreport.c:42


---

Generated on Mon Mar 19 2018 23:42:01 for FastqPuri by  

 1.8.14
