## Supplementary material for "FastqPuri: high-performance preprocessing of RNA-seq data": Rcommand__Sreport_8c.html

FastqPuri: src/Rcommand\_Sreport.c File Reference

FastqPuri

- src

Functions |
Variables

Rcommand\_Sreport.c File Reference

get Rscript command for Sreport
More...

`#include <stdio.h>`  
`#include <stdlib.h>`  
`#include <unistd.h>`  
`#include "Rcommand_Sreport.h"`  
`#include "init_Sreport.h"`  
`#include "defines.h"`  
`#include "config.h"`

Include dependency graph for Rcommand\_Sreport.c:

|  |  |
| --- | --- |
| Functions | |
| char \* | command\_Sreport () |
|  | returns Rscript command that generates the summary report in html More... |

|  |  |
| --- | --- |
| Variables | |
| Iparam\_Sreport | par\_SR |

#### Detailed Description

get Rscript command for Sreport

Author
:   Paula Perez paula.nosp@@m.o@@m.om

Date
:   09.08.2017

#### Function Documentation

#### ◆ command\_Sreport()

|  |  |  |  |
| --- | --- | --- | --- |
| char\* command\_Sreport | ( |  | ) |

returns Rscript command that generates the summary report in html

### To run between quotation marks after: Rscript\_RBioC -e (Rscript)

inputfolder = normalizePath( <par.SR.inputfolder>, mustWork = TRUE);

output = <par\_SR.outputfile>;

output\_file = gsub('.\* /', '', output);

path = gsub('[^/]+$', '', output);

if (path != '') {

outputfile = paste0(normalizePath(path, mustWork = TRUE), '/', outputfile);

} else {

outputfile = paste0(cwd, '/', output\_file); # cwd: current working dir

};

rmarkdown::render(<par\_SR.Rmd\_file>,

params = list(inputfolder = inputfolder, version= VERSION),

output\_file = output\_file)

#### Variable Documentation

#### ◆ par\_SR

Iparam\_Sreport par\_SR

input parameters Sreport

---

Generated on Mon Mar 19 2018 23:42:01 for FastqPuri by  

 1.8.14
