## Supplementary material for "FastqPuri: high-performance preprocessing of RNA-seq data": Sreport_8c.html

FastqPuri: src/Sreport.c File Reference

FastqPuri

- src

Functions |
Variables

Sreport.c File Reference

Sreport main function.
More...

`#include <stdio.h>`  
`#include <stdlib.h>`  
`#include <time.h>`  
`#include "init_Sreport.h"`  
`#include "Rcommand_Sreport.h"`  
`#include "config.h"`

Include dependency graph for Sreport.c:

|  |  |
| --- | --- |
| Functions | |
| int | main (int argc, char \*argv[]) |
|  | Qreport main function. |

|  |  |
| --- | --- |
| Variables | |
| Iparam\_Sreport | par\_SR |

### Detailed Description

Sreport main function.

Author
:   Paula Perez paula.nosp@@m.o@@m.om

Date
:   09.08.2017 This file contains the summary report main function. Given a folder containing \*bin as from Qreport output, Sreport generates a summary report in html format. See README\_Sreport.md for more details.

### Variable Documentation

### ◆ par\_SR

Iparam\_Sreport par\_SR

input parameters Sreport

---

Generated on Mon Mar 19 2018 23:42:01 for FastqPuri by  

 1.8.14
