## Supplementary material for "FastqPuri: high-performance preprocessing of RNA-seq data": PANDOC.md

### Installing Pandoc

#### Windows and Mac OS X

The [pandoc installation](http://johnmacfarlane.net/pandoc/installing.html) page includes easy to use installers for Windows and Mac OS X.

#### Linux

The version of pandoc included in the standard repositories is not recent enough for use with the **rmarkdown** package. You can install a more recent version of pandoc by installing the Haskell Platform and then following these instructions for [building pandoc from source](http://johnmacfarlane.net/pandoc/installing.html#all-platforms).

This method installs a large number of Haskell dependencies so might not be desirable. You can also obtain a standalone version of pandoc without the dependencies as follows:

##### Older Systems (RHEL 5)

For older Linux systems you can obtain a standalone version of pandoc v1.13.1 (with no Haskell dependencies) from <https://copr.fedoraproject.org/coprs/petersen/pandoc-el5/> as follows:

    $ sudo wget -P /etc/yum.repos.d/ https://copr.fedoraproject.org/coprs/petersen/pandoc-el5/repo/epel-5/petersen-pandoc-el5-epel-5.repo
    $ yum install pandoc pandoc-citeproc

##### Newer Systems (Debian/Ubuntu/Fedora/RHEL6+)

For newer Linux systems you can make a standalone version of pandoc v1.12.3 available to the system by soft-linking the binaries included with RStudio:

    $ sudo ln -s /usr/lib/rstudio/bin/pandoc/pandoc /usr/local/bin
    $ sudo ln -s /usr/lib/rstudio/bin/pandoc/pandoc-citeproc /usr/local/bin

If you are running RStudio Server the commands would be:

    $ sudo ln -s /usr/lib/rstudio-server/bin/pandoc/pandoc /usr/local/bin
    $ sudo ln -s /usr/lib/rstudio-server/bin/pandoc/pandoc-citeproc /usr/local/bin

If you aren't running RStudio at all you can simply copy the binaries out of the RStudio `bin/pandoc` directory and locate them within `/usr/local/bin` on your target system.
