## Supplementary material for "FastqPuri: high-performance preprocessing of RNA-seq data": quality_report.Rmd

````
---
title: "Assessing the quality of the reads"
output: 
   html_document:
      number_section: no
      toc_depth: 2
      toc_float: true
css: style.css
params:
    inputfile:
        value: x
    filter:
        value: x
    version:
        value: x
---

## General information

Running on version `r params$version` 

```{r, echo = F}
   warn <- ""
   if (params$filter == 2 )
      warn <- paste0("**WARNING**: file has been filtered with other tool.",
                     " Nucleotide position in read might be misleading")
```

`r warn`

```{r, echo = F}
source("utils.R")
library(pheatmap)
library(knitr)
data <- getQualStats(params$inputfile)
color <- colorRampPalette(c("white", "red"))( 50 )
color2 <- colorRampPalette(c("white", "blue"))( 50 )
qualities <- ""
for (i in data$qual_tags){
   qualities <- paste0(qualities, sprintf("%d (%s), ",i, intToUtf8(i+data$zeroQ) ))
}

qualities <- substr(qualities,1,nchar(qualities)-2) 

df_colnames <- c("Var","Value")
Var <- c("Input file name", "Read length", 
"Min good quality", "Number of reads",  
"Number of highQ reads", "Number of tiles",  
"Number of lanes", "Qualities",
"Reads with N's", "Number of N's")

Value=c(sub('.*\\/', '', params$inputfile),
  data$read_len,  
  data$minQ,
  data$nreads,   
  data$reads_MlowQ[1],   
  ifelse(data$tile_tags[1]==-1, "N/A", data$ntiles),
  ifelse(data$lane_tags[1]==-1, "N/A", length(unique(data$lane_tags))),
  qualities,
  data$reads_wN,
  sum(data$ACGT_pos[5,]))

df <- data.frame(Var, Value)

kable(df)


```

                    
## Per base sequence quality

```{r, echo = F}
  QualAbund <- getQualAbundancies(data)
  qualityPosSums <- apply(QualAbund, 1, sum)
  if (any(qualityPosSums == 0)) {
    cat("Incorrect quality values observed at position", which(qualityPosSums == 0), "\n")
  } else {
    sampleQ <- mimic(QualAbund)
    my_plot(sampleQ)
  }
```

## # reads with at least  `m` low Q nucleotides

```{r, echo =F}
barplot(data$reads_MlowQ[2:length(data$reads_MlowQ)], xlab="m" , names=1:data$read_len)
```

## Low Q nucleotide proportion per tile per lane

```{r, fig.width=4, fig.height=data$ntiles/6, echo = F}
if (data$tile_tags[1] == -1) {
  cat("No tile/lane information found in read headers, tile/lane quality heatmap skipped\n")
} else {
  if(all(data$lowQ_ACGT_tile == 0)){
     cat( "No low quality nucleotides found. tile/lane quality heatmap skipped\n")
  } else {
    lanes <- unique(data$lane_tags)
    for (lpos in lanes) {
      tiles <- which(data$lane_tags == lpos)
      pheatmap((data$lowQ_ACGT_tile/data$ACGT_tile)[tiles,1:4],cluster_cols=FALSE, 
               cluster_rows=FALSE, fontsize_row=6, fontsize_col=6, color = color,
               main = paste0("Lane ", lpos))
    }
  }
} 
```

## Average quality per position per tile per lane

```{r,fig.width=7, fig.height=data$ntiles/6, echo = F }
if (data$tile_tags[1] == -1) {
  cat("No tile/lane information found in read headers, tile/lane average quality heatmap skipped\n")
} else {
  tiles_meanQ <- array(dim=c(data$ntiles,data$read_len), 
                       dimnames = list(data$tile_tags, 1:data$read_len))
  for(i in 1:data$ntiles){
    for(j in 1:data$read_len){
      tiles_meanQ[i,j] <- weighted.mean(data$qual_tags,data$QPosTile_table[j,,i])
    }
  }
  lanes <- unique(data$lane_tags)
  for (lpos in lanes) {
    tiles <- which(data$lane_tags == lpos)
    if (any(is.nan(tiles_meanQ))){
      cat("\t\t TILES WITH MISSING POSITIONS FOUND. PLOTTING THEM BLACK.\n")
      color2[length(color2)] <- "#000000"
      from <- min(tiles_meanQ, na.rm=T); to <- max(tiles_meanQ, na.rm=T)
      breaks <- c(seq(from=from,  to=to, 
                      by = (to - from)/(length(color2) -1 )), to+0.01)
      tiles_meanQ[which(is.nan(tiles_meanQ))] = to+0.01 
      pheatmap(tiles_meanQ[tiles,],cluster_cols=FALSE, cluster_rows=FALSE, 
               fontsize_row=6, fontsize_col=6, color=color2, 
               breaks=breaks, main =  paste0("Lane ",lpos))
    } else {
      pheatmap(tiles_meanQ[tiles,],cluster_cols=FALSE, cluster_rows=FALSE, 
               fontsize_row=6, fontsize_col=6,color=color2, 
               main =  paste0("Lane ",lpos))
    }
  }
}
```

## Low Q nucleotides proportion per position per tile per lane

```{r,fig.width=7, fig.height=data$ntiles/6, echo = F}
if (data$tile_tags[1] == -1) {
  cat("No tile/lane information found in read headers, tile/lane position quality heatmap skipped\n")
} else {
  tiles_lowQ <- array(dim=c(data$ntiles,data$read_len), 
                      dimnames = list(data$tile_tags, 1:data$read_len))
  lowQ_idx <- which(data$qual_tags < data$minQ)
  for(i in 1:data$ntiles){
    for(j in 1:data$read_len){
      tiles_lowQ[i,j] <- sum(data$QPosTile_table[j,lowQ_idx,i])/
      sum(data$QPosTile_table[j,,i])
    }
  }
  if(sum(is.nan(tiles_lowQ)) > 0) {
    cat("\t\t THERE WERE TILES WITH MISSING POSITIONS. REMOVING THESE.\n")
    tiles <- which(rowSums(is.nan(tiles_lowQ)) == 0)
    data$tile_tags <- data$tile_tags[tiles]
    data$lane_tags <- data$lane_tags[tiles]
    tiles_lowQ <- tiles_lowQ[tiles,]
  }
  if (all(tiles_lowQ==0)){
    cat( "\t\t NO LOW QUALITY NUCLEOTIDES FOUND. NOT PLOTTING HEATMAP.\n")
  } else { 
    lanes <- unique(data$lane_tags)
    for (lpos in lanes){
      tiles <- which(data$lane_tags == lpos)
      pheatmap(tiles_lowQ[tiles,],cluster_cols=F, cluster_rows=F, 
               fontsize_row=6, fontsize_col=6,color=color, 
               main =  paste0("Lane ",lpos))
    }
  }
}
```


## Low Q nucleotides proportion per position for all tiles

```{r, echo = F, fig.height=2.5 }
N = length(data$qual_tags)
data$lowQprops <- sort(data$lowQprops)
lowQ_prop <- array(dim=c(length(data$lowQprops),data$read_len), 
                     dimnames = list(paste("<", data$lowQprops), 1:data$read_len))

i = 1
for (lowQ in data$lowQprops){
   lowQ_idx <- which(data$qual_tags < lowQ)
   for (j in 1:data$read_len){
      lowQ_prop[i,j] <- sum(data$QPosTile_table[j,lowQ_idx,])/sum(data$QPosTile_table[j,,])
   }
   i = i + 1
}
breaks <- seq(0,1,length=length(color))
pheatmap(lowQ_prop, cluster_cols=FALSE, cluster_rows=FALSE, fontsize_row=6, fontsize_col=6,color=color, breaks = breaks)
```

## Nucleotide content per position

```{r, echo = F}
   x <- 1:data$read_len
   nreads_pos <- colSums(data$ACGT_pos)
   y_min=min(data$ACGT_pos[1:4,]/nreads_pos)
   y_max=max(data$ACGT_pos[1:4,]/nreads_pos)
   plot(x,data$ACGT_pos[1,]/nreads_pos,type="l",lwd=1.5,col="green",
        ylim=c(y_min, y_max), xlab="Position in read",ylab="nucleotide content" )
   lines(x, data$ACGT_pos[2,]/nreads_pos,type="l",lwd=1.5,col="blue")
   lines(x, data$ACGT_pos[3,]/nreads_pos,type="l",lwd=1.5,col="black")
   lines(x, data$ACGT_pos[4,]/nreads_pos,type="l",lwd=1.5,col="red")
   lines(x, data$ACGT_pos[5,]/nreads_pos,type="l",lwd=1.5,col="grey")
   legend('topright', c("A","C","G", "T", "N "), bty= "n",lwd=1.5, 
          col=c("green","blue","black","red","grey"))
```
````
