## Supplementary material for "FastqPuri: high-performance preprocessing of RNA-seq data": summary_filter_report.Rmd

````
---
title: "Summary filtering report"
output: 
   html_document:
      number_section: no
      toc_depth: 2
      toc_float: true
css: style.css
params:
    inputfolder:
        value: x
    version:
        value: x
---


Running on version `r params$version` 

```{r, echo = F}
   library(knitr)
   source("utils.R")
   data <- getFilterTables(params$inputfolder)
```


## Filter set up

```{r, echo = F}
if ( !is.null(data$setup)) {
   kable(data$setup)
} else {
   print("Data analyzed do not correspond to the same set up conditions")
}
```

## Statistics summary 

```{r, echo = F}
   kable(data$table, digits = 3)
```
````
