## Supplementary material for "FastqPuri: high-performance preprocessing of RNA-seq data": summary_report.Rmd

````
---
title: "Summary quality report"
output: 
   html_document:
      number_section: no
      toc_depth: 2
      toc_float: true
css: style.css
params:
    inputfolder:
        value: x
    version:
        value: x
---

Running on version `r params$version` 

## General data

```{r, echo = F}
source("utils.R")
library(knitr)
library(pheatmap)
color <- colorRampPalette(c("white", "red"))( 50 )
color2 <- colorRampPalette(c("white", "blue"))( 50 )
files <- list.files(params$inputfolder,pattern="bin$")
fsize <- sapply(files, function(f) file.info(paste0(params$inputfolder,"/",f))$size)
files <- files[fsize > 100]
nombres <- gsub('\\.bin$', '', files)
Ns <- length(files)
table <- matrix(nrow = Ns, ncol = 4,
               dimnames = list(nombres,c("# reads","# tiles", 
               "% lowQ reads","% reads with N's")))

i = 1
for (f in files){
  data <- getQualStats(paste0(params$inputfolder,"/",f))
  if (i == 1){
    meanQ <- matrix(nrow = Ns, ncol = data$read_len,
             dimnames = list(nombres, 1:data$read_len))
  }
  table[i,] <- c(data$nreads,data$ntiles,
                 (data$nreads - data$reads_MlowQ[1])/data$nreads*100, 
                 data$reads_wN/data$nreads*100)
  for(j in 1:data$read_len){
     meanQ[i,j] <- weighted.mean(data$qual_tags,
                                 rowSums(data$QPosTile_table[j,,]))
  }
  i = i + 1
}

kable(table)

```

## Mean quality

```{r, fig.width=8, fig.height=max(1,Ns/7), echo = F}

pheatmap(meanQ,cluster_cols=F, cluster_rows=F, 
         fontsize_row=8, fontsize_col=6, color=color2)

```
````
