## Supplementary material for "FastqPuri: high-performance preprocessing of RNA-seq data": utils.R

```
getQualStats <- function(path){

   to.read = file(path,"rb")
   N_ACGT = 5
   res <- list() 
   res$read_len <- readBin(to.read, integer())
   res$ntiles <- readBin(to.read, integer())
   res$minQ <- readBin(to.read, integer())
   res$nLowQprops <- readBin(to.read, integer())
   res$nQ <- readBin(to.read, integer())
   res$zeroQ <- readBin(to.read, integer())
   res$nreads <- readBin(to.read, integer())
   res$reads_wN <- readBin(to.read, integer())
  
   res$sz_lowQ_ACGT_tile <- readBin(to.read, integer())
   res$sz_ACGT_tile <- readBin(to.read, integer())
   res$sz_reads_MlowQ <- readBin(to.read, integer())
   res$sz_QPosTile_table <- readBin(to.read, integer())
   res$sz_ACGT_pos <- readBin(to.read, integer())
    
   res$base_tags <- c("A","C","G","T","N")
   res$lowQprops <- readBin(to.read, integer(), n=res$nLowQprops)
   res$tile_tags <- readBin(to.read, integer(), n=res$ntiles)
   res$lane_tags <- readBin(to.read, integer(), n=res$ntiles)
   res$qual_tags <- readBin(to.read, integer(), n=res$nQ)
   res$lowQ_ACGT_tile <- t(array(readBin(to.read, integer(), 
                           n=res$sz_lowQ_ACGT_tile, size=8, 
                           endian = "little"),
                           dim=c(N_ACGT,res$ntiles), 
                           dimnames=list(res$base_tags,res$tile_tags)))
   res$ACGT_tile <- t(array(readBin(to.read, integer(), 
                       n=res$sz_ACGT_tile,size=8),
                       dim=c(N_ACGT,res$ntiles), 
                       dimnames=list(res$base_tags,res$tile_tags)))
   res$reads_MlowQ <- t(array(readBin(to.read, integer(), 
                        n=res$sz_reads_MlowQ,size=8),
                        dim=res$read_len+1,dimnames = list(0:res$read_len))) 
   res$QPosTile_table <- array(readBin(to.read, integer(), 
                         n=res$sz_QPosTile_table,size=8), 
                         dim=c(res$read_len,res$nQ,res$ntiles),
                         dimnames=list(1:res$read_len,
                                       res$qual_tags,res$tile_tags) )
   
   res$ACGT_pos <- array(readBin(to.read, integer(), 
                        n=res$sz_ACGT_pos,size=8),
                        dim=c(N_ACGT,res$read_len), 
                        dimnames=list(res$base_tags,1:res$read_len))
   close(to.read)
   res
}

getQualAbundancies <- function(data){
   Q_pos <- apply(data$QPosTile_table,2,rowSums)
   MeanQ <- apply(Q_pos,1, function(x) weighted.mean(data$qual_tags,x))
   rows <- nrow(Q_pos)
   quals <- ncol(Q_pos)
   sdQ <- numeric(rows)
   Ns <- rowSums(Q_pos)
   prob <- Q_pos/Ns
   N <- 10000
   simData<- matrix(0,nrow=N,ncol=rows)
   for (i in 1:rows){
      simData[,i] <- sample(data$qual_tags,N,replace=T,prob=prob[i,])
      for (j in 1:quals)
         sdQ[i] <- sdQ[i] + Q_pos[i,j]*((MeanQ[i] - data$qual_tags[j]))^2
      sdQ[i] <- sqrt(sdQ[i]/(Ns[i] -1))
   }

   QualAbund <- matrix(0,nrow = rows, ncol= quals, dimnames = 
                       list(NULL,data$qual_tags))
   i <- 1
   for (tabla in  apply(simData,2,table)){
       QualAbund[i,names(tabla)] = tabla
       i <- i + 1
   }

   return (QualAbund) 
}

mimic <- function(Q){
    L <- nrow(Q)
    N <- 10000
    means <- numeric(L)
    qualities <- as.numeric(colnames(Q))
    Q_boxes <- matrix(0,ncol=L,nrow=N)
    whiskers <- matrix(0,ncol = 2, nrow = L)
    colnames(whiskers) <- c("Q10","Q90")
    for( i in 1:L){
        means[i] <- weighted.mean(qualities,Q[i,])
        Q_boxes[,i] <- sample(qualities, N, replace=TRUE, prob=Q[i,]/sum(Q[i,]))
        sorted <- sort(Q_boxes[,i])
        whiskers[i,] <- sorted[c(round(N*0.1,0), round(N*0.9,0))]
    }
    return(list( Q_boxes =  Q_boxes, means = means, whiskers = whiskers, qualities = qualities))
}

simulateQ <- function(){
   N <- 10000
   boxplotData<- matrix(0,nrow=N,ncol=25)
   means <- c(25,25,25,25,31,32,32,32,32,35,35,36,35,
              36,37,37,37,37,37,37,37,31,29,28,27)
   for (i in 1:25){
      a <- round(rnorm(100000,means[i]),0)
      l <-table(a)
      boxplotData[,i] <- sample(as.numeric(names(l)),10000,replace=T,prob=l/sum(l))
   }
   return(boxplotData)
}

my_boxplot <- function(data,...){
    a <- boxplot(data$Q_boxes, col="grey", outline=FALSE, range=1,...)
    points(data$means,type="l", col="blue")
    d <- 0.2
    for (i in 1:nrow(data$whiskers)){
       lines(c(i,i),data$whiskers[i,],col="red" )
       lines(c(i-d,i+d),rep(data$whiskers[i,1],2),col="red")
       lines(c(i-d,i+d),rep(data$whiskers[i,2],2),col="red")
    }
    return(a)
}

my_plot <- function(data){
    L <- ncol(data$Q_boxes)
    my_boxplot(data, ylim=c(0,42),xlab="Position in read", ylab="Quality", xaxt="n", yaxt="n")
    axis(1,1:L,1:L,cex.axis=0.6)
    axis(2,data$qualities,data$qualities, cex.axis=0.6)

}

# Reads trimFilter output data 
# and stores them in a list.
getFilterStats <- function(path) {
  to.read = file(path, "rb")
  if (file.info(path)$size != 56) {
      stop("In file ", path, " size is ", file.info(path)$size, " instead of 56 bytes as expected. Exiting.")
  }
  NFILTER <- 4
  res <- list()
  res$filter <- readBin(to.read, integer(), n=NFILTER)
  res$trimmed <- readBin(to.read, integer(), n=NFILTER)
  res$discarded <- readBin(to.read, integer(), n=NFILTER)
  res$good <- readBin(to.read, integer())
  res$nreads <- readBin(to.read, integer())
  if (res$nreads > 0) {
     res$trimmed <- res$trimmed/res$nreads*100
     res$discarded <- res$discarded/res$nreads*100
  } else {
     stop("All reads were discarded")
  }
  close(to.read)
  res   
}

# Reads trimFilterDS output data 
# and stores them in a list.
getFilterStatsDS <- function(path) {
  to.read = file(path, "rb")
  if (file.info(path)$size != 72) {
    stop("In file ", path, " size is ", file.info(path)$size, " instead of 72 bytes as expected. Exiting.")
  }
  NFILTER <- 4
  res <- list()
  res$filter <- readBin(to.read, integer(), n=NFILTER)
  res$trimmed1 <- readBin(to.read, integer(), n=NFILTER)
  res$trimmed2 <- readBin(to.read, integer(), n=NFILTER)
  res$discarded <- readBin(to.read, integer(), n=NFILTER)
  res$good <- readBin(to.read, integer())
  res$nreads <- readBin(to.read, integer())
  if (res$nreads > 0) {
     res$trimmed1 <- res$trimmed1/res$nreads*100
     res$trimmed2 <- res$trimmed2/res$nreads*100
     res$discarded <- res$discarded/res$nreads*100
  } else {
     stop("All reads were discarded")
  }
  close(to.read)
  res   
}

# Construct the two tables that will be outputted 
# in the summary_filter_report performing 
# some consistency checks
getFilterTables <- function(inputfolder) {
   NFILTER <- 4
   GRAL_TABLE <- TRUE
   adapters <- c("NONE", "DEFAULT")
   method <- c("NONE", "TREE", "SA", "BLOOM")
   trimQ <- c("NONE", "ALL", "ENDS", "FRAC", "ENDSFRAC", "GLOBAL")
   trimN <- c("NONE", "ALL", "ENDS", "STRIPS")
   applied <- c("NO", "YES", "YES", "YES", "YES", "YES")
   files <- list.files(inputfolder,pattern=".bin$")
   fsizes <- sapply(files, function(f) file.info(paste0(inputfolder,"/", f))$size)
   files <- files[fsizes == 56]
   nombres <- gsub('_summary\\.bin$', '', files)
   Ns <- length(files)
   table <- matrix(nrow = Ns, ncol = 9,
            dimnames = list(nombres,c("Nreads","Naccepted", "%disc Ad", 
                                      "%cont", "%disc lowQ", "%disc N's", 
                                      "%trim Ad", "%trim N's", "%trim lowQ")))
   i = 1
   for (f in files) {
      st_f <- getFilterStats(paste0(inputfolder,"/",f)) 
      if (i == 1) {
         filter = st_f$filter
      } else {
         GRAL_TABLE  <- GRAL_TABLE && all.equal(filter, st_f$filter) 
      }
      table[i,] <- c(st_f$nreads, st_f$good, st_f$discarded, 
                     st_f$trimmed[c(1,3,4)])
      i = i+1
   }
   if (GRAL_TABLE) {
      setup  <- matrix(nrow = NFILTER, ncol =  2, dimnames = 
                       list(c("ADAPTERS", "CONTAMINATIONS", "LOW Q", "N's"),
                           c("Applied", "Method")))
      setup[,1] <- applied[st_f$filter+1]
      setup["ADAPTERS",2] <- adapters[st_f$filter[1] +1]
      setup["CONTAMINATIONS",2] <- method[st_f$filter[2] +1]
      setup["LOW Q",2] <- trimQ[st_f$filter[3] +1]
      setup["N's",2] <- trimN[st_f$filter[4] +1]
   } else  {
      setup <- NULL
   }
   return (list(setup = setup, table = table))
}

# Construct the two tables that will be outputted 
# in the summary_filter_reportDS performing 
# some consistency checks
getFilterTablesDS <- function(inputfolder) {
   NFILTER <- 4
   GRAL_TABLE <- TRUE
   adapters <- c("NONE", "DEFAULT")
   method <- c("NONE", "TREE", "SA", "BLOOM")
   trimQ <- c("NONE", "ALL", "ENDS", "FRAC", "ENDSFRAC", "GLOBAL")
   trimN <- c("NONE", "ALL", "ENDS", "STRIPS")
   applied <- c("NO", "YES", "YES", "YES", "YES", "YES")
   files <- list.files(inputfolder,pattern="_summary\\.bin$")
   fsizes <- sapply(files, function(f) file.info(paste0(inputfolder, "/", f))$size)
   files <- files[fsizes == 72]
   nombres <- gsub('_summary\\.bin$', '', files)
   Ns <- length(files)
   table <- matrix(nrow = Ns, ncol = 11,
            dimnames = list(nombres,c("Nreads","Naccepted", "%disc Ad", 
                                      "%cont", "%disc lowQ", "%disc N's", 
                                      "%trim Ad", "%trim1 lowQ", "%trim1 N's", 
                                      "%trim2 lowQ", "%trim2 N's")))
   i = 1
   for (f in files) {
      st_f <- getFilterStatsDS(paste0(inputfolder,"/",f)) 
      if (i == 1) {
         filter = st_f$filter
      } else {
         GRAL_TABLE  <- GRAL_TABLE && all.equal(filter, st_f$filter) 
      }
      table[i,] <- c(st_f$nreads, st_f$good, st_f$discarded, 
                     st_f$trimmed1[c(1,3,4)], st_f$trimmed2[c(3,4)])
      i = i+1
   }
   if (GRAL_TABLE) {
      setup  <- matrix(nrow = NFILTER, ncol =  2, dimnames = 
                       list(c("ADAPTERS", "CONTAMINATIONS", "LOW Q", "N's"),
                           c("Applied", "Method")))
      setup[,1] <- applied[st_f$filter+1]
      setup["ADAPTERS",2] <- adapters[st_f$filter[1] +1]
      setup["CONTAMINATIONS",2] <- method[st_f$filter[2] +1]
      setup["LOW Q",2] <- trimQ[st_f$filter[3] +1]
      setup["N's",2] <- trimN[st_f$filter[4] +1]
   } else  {
      setup <- NULL
   }
   return (list(setup = setup, table = table))
}
```
