## Supplementary material for "FastqPuri: high-performance preprocessing of RNA-seq data": README.md

FastqPuri, an fq quality control and filter tool 
=========

Software and source code of `FastqPuri`. It creates quality 
reports of `fastq` files and filters them removing low quality 
reads, reads containing too many N's or contamination reads 
(unwanted rRNA reads, impurities coming from another organism, ...).


## Installation

Clone the repository, or download the source. Make sure that 
your system supplies the following dependencies for FastqPuri.

- OS: Linux (clang, gcc), Mac OS (clang, gcc), OpenBSD (clang)
- `cmake` (at least version 2.8), 
- a `C` compiler supporting the `c11` standard 
  (change the compiler flags otherwise),
- pandoc (optional, see documentation in `PANDOC.md`),
- `Rscript` (optional),
- Following `R` packages installed (optional):
   * `pheatmap`
   * `knitr`
   * `rmarkdown`

**NOTE:**  FastqPuri will work without the optional dependencies 
but will skip creating html reports if they are not available.

```
$ cmake -H. -Bbuild/ [-DRSCRIPT=/path/to/my/R/bin/Rscript ... ]
$ cd build 
$ make 
```

When running cmake, there are some variables you can set 
using the option -D followed by the variable name. This variables are, 

- `CMAKE_C_COMPILER`: `C` compiler (default `gcc`)
- `CMAKE_C_FLAGS`: compiler flags (default `-Wall -O3 -march=native -std=c11`).
- `PANDOC`: `pandoc` executable (default `pandoc`),
- `RSCRIPT`:  `Rscript` executable (default `Rscript`), 
- `READ_MAXLEN`: Maximum Illumina read length (default 400),
- `RMD_QUALITY_REPORT`: path to `quality_report.Rmd`,
- `RMD_SUMMARY_REPORT`: path to `summary_report.Rmd`,

The executables will be created in the folder `bin`. 

**WARNING:** do not move the executables that depend on `R` scripts, 
anywhere else, unless you also move the corresponding `R` scripts respecting
the local folder structure. 


## Executables

* `Qreport`: creates a quality report in html format (see `README_Qreport.md`),
* `Sreport`: creates a summary report in html format on a set of samples, 
   regarding either the original files or the filtering process
   (see `README_Sreport.md`),
* `makeBloom`: creates a  bloom filter from a fata file of a certain size,
   and stores it in a file (see `README_makeBloom.md`)
* `makeTree`: creates a tree of a certain depth from a fasta file and stores
 it in a file (see `README_makeTree.md`),
* `trimFilter`: performs the filtering process for single-end data 
   (see `README_trimFilter.md`).
* `trimFilterPE`: performs the filtering process for double stranded data 
   (see `README_trimFilterPE.md`).

An exemplar work flow could be:

* `Qreport`
* `Sreport`
* `makeBloom`
* `trimFilter` or `trimFilterPE`
* `Qreport`
* `Sreport`

## Documentation of the code

A Doxygen documentation of the code is available: 
- `html` version under the folder `html` (open `index.html` with a browser).
- `pdf` version: `latex/refman.pdf`

## Contributors

Paula Pérez Rubio 

## License

GPL v3 (see LICENSE.txt)
