## Supplementary material for "FastqPuri: high-performance preprocessing of RNA-seq data": README_makeBloom.md

# makeBloom user manual

Reads a `fasta` file, creates a bloom filter with a predefined:
 - size (bits). 
 - number of hash functions.
 - false positive rate.
and saves it to a file. 

## Running the program

Usage `C` executable (in folder `bin`): 

```
Usage: ./makeBloom --fasta <FASTA_INPUT> --output <FILTERFILE> --kmersize [KMERSIZE] 
 (--fal_pos_rate [p] | --hashNum [HASHNUM] | --bfsizeBits [SIZEBITS])
Options: 
 -v, --version      Prints package version.
 -h, --help         Prints help dialog.
 -f, --fasta        Fasta input file. Mandatory option.
 -o, --output       Output file, with NO extension. Mandatory option.
 -k, --kmersize     kmer size, number or elements. Optional(default = 25)
 -p, --fal_pos_rate false positive rate. Optional (default = 0.05)
 -g, --hashNum      number of hash functions used. Optional (default
                    value computed from the false positive rate).
 -m, --bfsizeBits   size of the filter in bits. It will be forced to be
                    a multiple of 8. Optional (default value computed
                    from the false positive rate).
NOTE: the options -p, -g, -m are mutually exclusive. The program 
      will give an error if more than one of them are passed as input.
      It is recommended to pass the false positive rate and let the 
      program compute the other variables (excepting singular situations).
      If none of them are passed, the false positive rate will default
      to 0.05 and the other variables will be computed respecting this
      requirement. See the `Doxygen` documentation and supplementary material for 
      more details.
```


## Output description

Two files are created as output: 
 - `<FILTERFILE>.bf`: contains the filter (binary file).
 - `<FILTERFILE>.txt`: contains the following information: 
   * `kmersize`: length of the kmers inserted in the filter, 
   * `bfsizeBits`: size of Bloom filter in bits,
   * `hashNum`: number of hash functions used in the filter, 
   * `falsePosRate:` false positive rate,
   * `nelem`: number of elements (kmers in the sequece) contained in the filter.

For further details, read the `Doxygen` documentation of the files
`bloom.c`, `init_makeBloom.c`, `makeBloom.c`

## Example 

 In the folder `../examples/bloomROC/` an example script can be run to 
test the bloom filter performance.  
 - The fastq files:
   * `human_reads.fq.gz`
   * `EColi_reads.fq.gz`
   are analyzed. They contain 10e5 reads each generated with `dgwsim`.
 - STEP1: Create bloom filters for the E.coli genome  with FPR:
   `[0.005 0.0075 0.01 0.02]`                                            

 - STEP2: Run trimFilter on both data looking for contaminations from E.coli
          using all filters generated with `kmersize = 25` and scores ranging
          from `0.05` to `0.2` in `0.01` intervals. We obtain false/true 
          positive/negative rates:
   * FPR: % contaminations detected in human_reads.fq.gz
   * TNR (specificity): % good reads detected in `human_reads.fq.gz`
   * FNR: % good reads detected in `EColi_reads.fq.gz`
   * TPR (sensitivity): % contaminations detected in `EColi_reads.fq.gz`
                                                                              
 - STEP3: Create ROC curves for all filters (sensitivity vs FPR).            

The results (`*csv`, `*pdf`) can be compared with `example*pdf`, and
`example*csv` in the folder (see example for `-p = 0.0075` below). 

<p align="center">
<img src=./pics/ROC_0p0075_bloom.png alt="noimage" title="ROC plots">
</p>


## Details on bloom filters

**NOTE**: For further details, read the `Doxygen` documentation of the files
`bloom.c`, `init_makeBloom.c`, `makeBloom.c` and the supplementary material.

A bloom filter is a probabilistic data structure used to test if an element
is a member of a set. False positive matches are possible but false negative 
are not. For a given set of `n` elements, we proceed as follows: 

- decide on the number `g` of hash functions we will use for the construction
of the filter and the length of the filter `m` (number of bits). This choice 
will be made based on the false positive rate we want to 
achieve (see Parameters).

- we create an empty bloom filter, `B`, i.e. an array of `m` bits set 
to `0`. For every element in the set,  **s<sub>&alpha;</sub> &isin; S**, compute 
the `g` hash functions,  **H<sub>i</sub> (s<sub>&alpha;</sub>) &forall; 
i  &isin; {1,...,g}** and set the corresponding bits to
`1` in the filter, i.e., 
**B[H<sub>i</sub> (s<sub>&alpha;</sub>) mod m] = 1 &forall; i &isin; 
{1,...,g}** and `0` otherwise. 

Then, if we want to check whether an element **s<sub>&beta;</sub>** is in 
the set **S**, we compute **H<sub>i</sub> (s<sub>&beta;</sub>) &forall;
i &isin; {1,...,g}** and check whether all coresponding positions in the 
filter are set to `1`, in which case we can say that **s<sub>&beta;</sub>** 
might be in the set. Otherwise it is definitely not in the set. 

### Parameters 

We choose the parameters so that the desired false positive rate is 
achieved. Alternatively, we can pass the filter size, and then the 
number of hash functions to be used is tuned so that the false positive
rate is minimized. 

We assume the hash functions select all positions with the same 
probability. The probability that a bit in the filter `B` is not 
set to `1` after inserting an element using `g` hash functions is: 

<p align="center"><b>
(1 - <sup>1</sup>&frasl;<sub>m</sub>)<sup>g</sup>
</b></p>

where `m` is the number of bits of the filter. If we insert `n` elements, 
the probability that an element is still 0 is: 

<p align="center"><b>
p<sub>0</sub> = (1 - <sup>1</sup>&frasl;<sub>m</sub>)<sup>gn</sup>
</b></p>

The probability that a bit is `1` is then, 

<p align="center"><b>
p<sub>1</sub> = 1 - (1 - <sup>1</sup>&frasl;<sub>m</sub>)<sup>gn</sup>
</b></p>

Now, let's compute the false positive rate, i.e., that probability that
an element that is not in the set is classified as belonging to it. This 
is the probability that all positions computed from the hash functions 
being `1` is, 

<p align="center"><b>
p(g, n, m) = (1 - (1 - <sup>1</sup>&frasl;<sub>m</sub>)<sup>gn</sup>)<sup>g</sup>
= (1 - e<sup> - <sup>gn</sup>&frasl;<sub>m</sub> </sup>)<sup>g</sup>
</b></p>

For a given `n` and `m` the value of `g` that minimizes `p` is, 

<p align="center"><b>
<sup>dp(g, n, m)</sup>&frasl;<sub>dg</sub> = 0 &rArr; 
g = <sup>m</sup>&frasl;<sub>n</sub> log(2)
</b></p>

The required number of bits `m` for the desired positive rate given 
`n` number of elements and assuming the optimal number of hash functions 
being used is, 

<p align="center"><b>
m = - <sup>n log (p)</sup>&frasl;<sub>log<sup>2</sup>(2)</sub>.
</b></p>

### Creating a bloom filter from a `fasta` file

Given a `fasta` file, the elements to be inserted in the bloom filter are 
all possible `k`-mers contained in the `fasta` file. The length `k` of the 
`k`-mers can be given by the user as an input parameter and is chosen to 
be `25` by default. All `k`-mers containing nucleotides different from 
`{A,C,G,T}` will not be considered and they are encoded such that every
nucleotide takes only 2-bits memory. We look into the reverse complement 
and insert only the one that is lexicographically smaller. 

Once the `k`-mer has been processed, the hash functions are computed and the 
positions of the output values are set to `1` in the filter. 


### Checking if a read in a `fastq` file is in the filter

To check whether a `fastq` read of length `L` is in the filter, we 
proceed as follows: 

1. Start with a score `score = 0`. 
2. Construct and process all `k`-mers of length `k` in the read, `(L - k + 1)`
   (note that `L `&ge;`k`)
3. For each `k`-mer, compute the `g` hash functions and check whether all 
   bits in the corresponding positions in the filter are set to `1`. If so, 
   add  **<sup>1</sup>&frasl;<sub>(L - k + 1)</sub>** to the score. 

If the score is above the user predefined threshold (`-s`), 
the read is classified as belonging to the set, and not otherwise. 

### Memory usage, sensitivity and specificity

The **memory usage** will be determined by `m`, the size of the filter. The optimal
number of bits per element is

<p align="center"><b>
<sup>m</sup>&frasl;<sub>n</sub> = - <sup> log (p)</sup>&frasl;
<sub>log<sup>2</sup>(2)</sub>.
</b></p>

In the figures below, we can see both, the optimal number of bits 
per element and the optimal number of hash functions as a function 
of the false positive rate. 


<p align="center">
<img src=./pics/bloomfilter.png alt="noimage" title="FDR plots">
</p>

As an example, let's assume we want to look for contaminations in a
genome of `~3GB` and want to keep the false positive rate at `2%`. Then,
we will need a filter of `~3.05GB`. 

**Sensitivity** (true positive rate,  TP/(TP + FN)) can be increased 
by decreasing the `k`-mer size (`-k`) and the score threshold. False 
negatives only occur in the presence of mismatches due to variants, 
or errors in the base calling procedure, since the filter itself
does not allow for false negatives. 

To increase **specificity** (true negative rate, TN/(TN + FP)), you can increase
the score threshold (`-s`) or, obviously reduce the positive rate, (`-p`). 


## Contributors

Paula Pérez Rubio 

## License

GPL v3 (see LICENSE.txt)
