## Supplementary material for "FastqPuri: high-performance preprocessing of RNA-seq data": README_makeTree.md

# makeTree user manual

Reads a `fasta` file, creates a tree structure of a predefined 
depth `DEPTH` and saves it to a file. 

**NOTE**: computing a tree from a fasta file is very memory 
intensive. The program will not compute a tree if the length of the 
sequences in the fasta file exceeds 10 MB. 

## Running the program

Usage `C` executable (in folder `bin`): 

```
Usage: ./makeTree -f|--fasta <FASTA_INPUT> -l|--depth <DEPTH> 
-o, --output <OUTPUT_FILE> Reads a *fa file, constructs a tree of 
depth DEPTH and saves it compressed in OUTPUT_FILE.
Options: 
 -v, --version Prints package version.
 -h, --help    Prints help dialog.
 -f, --fasta   Fasta input file of potential contaminations. Mandatory option.
 -l, --depth   depth of the tree structure.
 -o, --output  Output file. If the extension is not *gz, it is added. Mandatory option.
```


## Output description

Compressed file containing the tree structure. For further details,
read the `Doxygen` documentation of the file `tree.c`, function `save_tree`.

## Example 
 
TODOTODOTODOTODOTODO (It was tested somewhere!!)

## Contributors

Paula Pérez Rubio 

## License

GPL v3 (see LICENSE.txt)
