## Supplementary material for "FastqPuri: high-performance preprocessing of RNA-seq data": README_Qreport.md

# QReport user manual

Obtains statistics from the read qualities of a fastq file. 
The binary output is then read by an `Rmd` script that generates a 
html visualization of the data. 


## Running the program

Usage `C` executable (in folder `bin`): 

```
Usage: ./Qreport -i <INPUT_FILE.fq> -l <READ_LENGTH>
       -o <OUTPUT_FILE> [-t <NUMBER_OF_TILES>] [-q <MINQ>]
        [-n <#_QUALITY_VALUES>] [-f <FILTER_STATUS>]
	[-0 <ZEROQ>] [-Q <quality-values>]
Reads in a fq file (gz, bz2, z formats also accepted) and creates a
quality report (html file) along with the necessary data to create it
stored in binary format.
Options: 
 -v prints package version.
 -h prints help dialog.
 -i Input file [*fq|*fq.gz|*fq.bz2]. Mandatory option.
 -l Read length. Length of the reads. Mandatory option.
 -o Output file prefix (with NO extension). Mandatory option.
 -t Number of tiles. Can be obtained from Illumina's `Run Summary` (see https://support.illumina.com/help/BaseSpace_OLH_009008/Content/Vault/Informatics/Sequencing_Analysis/BS/swSEQ_mBS_ViewRunSamplesList.htm). It is usually 96 for flow cells used on HiSeq and NextSeq machines, and 19 for flow cells for the MiSeq with v3 chemistry. Multiply by the number of lanes, if data comes from more than one lane. Optional (default 96).
 -q Minimum quality allowed. Optional (default 27).
 -n Number of different quality values allowed. Optional (default 46).
 -f Filter status: 0 original file, 1 file filtered with trimFilter,
    2 file filtered with another tool. Optional (default 0).
 -0 ASCII value for quality score 0. Optional (default 33).
 -Q quality values for low quality proportion plots. Optional (default 27,33,37),
    Format is either <int>[,<int>]* or <min-int>:<max-int>.
    
```


## Output description

- Binary output: 
   * `int` (4B) : read length (`read_len`), 
   * `int` (4B) : number of tiles (`ntiles`),
   * `int` (4B) : minimum quality accepted (`minQ`),   
   * `int` (4B) : number of possible qualities (`nQ`), 
   * `int` (4B) : number of reads (`nreads`),
   * `int` (4B) : number of reads containing N's (`reads_wN`),
   * `int` (4B) : size of `lowQ_ACGT_tile`, see below (`sz_lowQ_ACGT_tile`),
   * `int` (4B) : size of `ACGT_tile`, see below (`sz_ACGT_tile`), 
   * `int` (4B) : size of `reads_MlowQ`, see below (`sz_reads_MlowQ`), 
   * `int` (4B) : size of `QPosTile_table`, see below (`sz_QPosTile_table`),
   * `int` (4B) : size of `ACGT_pos`, see below (`sz_ACGT_pos`),
   * `ntiles*int ` (4x`ntiles`B) : tile tags (`tile_tags`),
   * `ntiles*int ` (4x`ntiles`B) : lane tags (`lane_tags`),
   * `nQxint` (4x`nQ`B) : quality tags (`quality tags`),
   * `5 x ntiles x (long int)` (5x`ntiles`x8B) : 
     \# (A,C,G,T,N) per tile with low quality (`lowQ_ACGT_tile`),
   * `5 x ntiles x (long int)` (5x`ntiles`x8B) : 
     \# (A,C,G,T,N) per tile (`ACGT_tile`),
   * `(read_len+1) x (long int)` ((`read_len`+1)x8B) : 
     \# reads with at least `m` low quality base callings (`reads_MlowQ`),
   * `ntiles x read_len x nQ x (long int)` (`ntiles`x`read_len`x`nQ`x8B) : 
     base callings  per tile per position with a given quality (`QposTile_table`).
   * `5 x read_len x (long int)` (5x`read_len`x8B ): 
     \# (A,C,G,T,N) per position, (`ACGT_pos`)

- html output:
   * Table with general information,
   * Plot 1: per base sequence quality, 
   * Plot 2: # reads with at least m low Q base callings,
   * Plot 3: Low Q base calling proportion per nucleotide type per tile per lane,
   * Plot 4: Average quality per position per tile per lane,
   * Plot 5: Low Q base calling proportion per position per tile per lane,
   * Plot 6: Low Q base calling proportion per position for all tiles.
   * Plot 7: Nucleotide content per position

**NOTE** If the data were sequenced on more than one lane, in Plots 3, 4 and 5 
there will be one heatmap per lane.

## Example 

An example is given in the folder `examples/QReport_Sreport`. To run an 
example, type, 

``` 
    $ cd example/Qreport_Sreport
    $ mkdir run_test
    $ cd run_test
    $ ../../../bin/QReport -i ../test.fq.bz2 -l 51 -o my_test_output
```
 and compare it with the provided run example, as specified in the README
 file under `./example/QReport_Sreport`.

  
## Contributors

Paula Pérez Rubio 

## License

GPL v3 (see LICENSE.txt)
