## Supplementary material for "FastqPuri: high-performance preprocessing of RNA-seq data": README_Sreport.md

# SReport user manual

Creates two types of summary reports in html format

 - **Q** (Quality): report of the quality status of a set of `fastq` files.
   Needs the binary output of `Qreport`, for several files. It will analyze
   all files with extension `*.bin` inside the folder specified in
   the input and will expect them to have the appropriate format
   (output from `Qreport`).
 - **F** (Filter): report of the filtering process (`trimFilter`) of a set
   of `fastq` files. It will analyze all files with extension `*bin` inside
   the folder specified in the input and will expect them to have the
   appropriate format, (output from `trimFilter`).
 - **P** (Filter): report of the filtering process (`trimFilterPE`) of a set
   of paired-end `fastq` files. It will analyze all files with extension 
   `*bin` inside the folder specified in the input and will expect them to have 
   the appropriate format, (output from `trimFilterPE`).


## Running the program

Usage `C` executable (in folder `bin`):

```
Usage: ./Sreport -i <INPUT_FOLDER> -t <Q|T|P> -o <OUTPUT_FILE>
Uses all *bin files found in a folder (output of Qreport|trimFilter)
and generates a summary report in html format (of Qreport|trimFilter).
Options:
 -v Prints package version.
 -h Prints help dialog.
 -i Input folder containing *bin data (output from Qreport). Mandatory option.
 -t {Q,F,P} Type of report to be generated: 'Q' for quality summary
    report, 'F' for filter summary report based on sinlge-end reads,
    and 'P' for filtering summary report based on paired-end reads
    data filter summary report. Mandatory option,
 -o Output file (with NO extension). Mandatory option.
```

## Output description

- **Q** (html output):
   * Table with: `# reads , # tiles, % lowQ reads, % reads with N’s` for
     all samples.
   * Heatmap showing the average quality per position for all samples.
- **F** (html output):
   * Table with .trimFilter setup. If there were samples with different
     setups, Sreport only reports summaries generated for single-end filtering
   * Table with : `Nreads`, `Naccepted`, `%disc Ad`, `%cont`, `%disc lowQ`,
     `%disc N’s`, `%trim Ad`,`%trim N’s`,`%trim lowQ` for all samples.
- **P** (html output):
   * Table with .trimFilterPE setup. If there were samples with different
     setups, Sreport only reports summaries generated for paired-end filtering
   * Table with : `Nreads`, `Naccepted`, `%disc Ad`, `%cont`, `%disc lowQ`,
     `%disc N’s`, `%trim Ad`,`%trim1 N’s`,`%trim1 lowQ`, `%trim2 N’s`, 
     `%trim2 lowQ` for all samples.

## Example

###  Option `-t Q`

An example is given in the folder `examples/QReport_Sreport`. To run an
example, type,

```
    $ cd example/Qreport_Sreport/run_test/
    $ ../../../bin/SReport -i ./ -t Q -o my_test_summary_report
```
 and compare it with the provided run example, as specified in the README
 file under `./examples/QReport_Sreport`

**NOTE:** `Sreport` has to be run AFTER the `Qreport` example

###  Option `-t F`

In folder `.examples/trimFilter_Sreport/bin_files`, 30 fake
`./trimFilter` binary output files were generated (with the `R` script
`create_fake_bins.R`). An html output was created from them
(`.examples/trimFilter_Sreport/bin_files/filter_Sreport_example.html`).
It can be reproduced if you run:

```
$ ../../bin/Sreport -i .bin_files/ -o ./bin_files/filter_Sreport_new -t F
```

###  Option `-t P` 

In folder `.examples/trimFilterPE_Sreport/bin_files`, 30 fake
`./trimFilterPE` binary output files were generated (with the `R` script
`create_fake_bins.R`). An html output was created from them
(`.examples/trimFilterPE_Sreport/bin_files/DS_Sreport_example.html`).
It can be reproduced if you run:

```
$ ../../bin/Sreport -i .bin_files/ -o ./bin_files/DS_Sreport_new -t P
```

## Contributors

Paula Pérez Rubio

## License

GPL v3 (see LICENSE.txt)
