## Supplementary material for "FastqPuri: high-performance preprocessing of RNA-seq data": README_trimFilter.md

# trimFilter user manual

 This program reads a `fastq` file as an input, and filters it according
 to the following criteria:
 - Discard/trims reads containing adapter remnants.
 - Discards reads matching contaminations (sequences collected in a `fasta`
   file or in an `idx` file created by `makeTree` or `makeBloom`.
 - Discards/trims low quality reads.
 - Discards/trims reads containing N base callings.

## Running the program

Usage `C` executable (in folder `bin`): 

```
Usage: trimFilter --ifq <INPUT_FILE.fq> --length <READ_LENGTH>
                  --output [O_PREFIX] --gzip [y|n]
                  --adapter [<ADAPTERS.fa>:<mismatches>:<score>]
                  --method [TREE|BLOOM]
                  (--idx [<INDEX_FILE>:<score>:<lmer_len>] |
                   --ifa [<INPUT.fa>:<score>:[lmer_len]])
                  --trimQ [NO|ALL|ENDS|FRAC|ENDSFRAC|GLOBAL]
                  --minL [MINL]  --minQ [MINQ] --zeroQ [ZEROQ]
                  (--percent [percent] | --global [n1:n2])
                  --trimN [NO|ALL|ENDS|STRIP]
Reads in a fq file (gz, bz2, z formats also accepted) and removes:
  * low quality reads,
  * reads containing N base callings,
  * reads representing contaminations, belonging to sequences in INPUT.fa
Outputs 4 [O_PREFIX]_fq.gz files containing: "good" reads, discarded
low Q reads, discarded reads containing N's, discarded contaminations.
Options:
 -v, --version prints package version.
 -h, --help    prints help dialog.
 -f, --ifq     fastq input file [*fq|*fq.gz|*fq.bz2], mandatory option.
 -l, --length  read length: length of the reads, mandatory option.
 -o, --output  output prefix (with path), optional (default ./out).
 -z, --gzip    gzip output files: yes or no (default yes).
 -A, --adapter adapter input. Three fields separated by colons:
               <ADAPTERS.fa>: fasta file containing adapters,
               <mismatches>: maximum mismatch count allowed,
               <score>: score threshold  for the aligner.
 -x, --idx     index input file. To be included with any method. 
               3 fields separated by colons:
               <INDEX_FILE>: output of makeTree, makeBloom,
               <score>: score threshold to accept a match [0,1],
               [lmer_len]: corresponds to the length of the lmers to be
                        looked for in the reads [1,READ_LENGTH].
 -a, --ifa     fasta input file. To be included only with method TREE
               (it excludes the option --idx). Otherwise, an
               index file has to be precomputed and given as parameter
               (see option --idx). 3 fields separated by colons:
               <INPUT.fa>: fasta input file [*fa|*fa.gz|*fa.bz2],
               <score>: score threshold to accept a match [0,1],
               <lmer_len>: depth of the tree: [1,READ_LENGTH]. It will
                        correspond to the length of the lmers to be
                        looked for in the reads.
 -C, --method  method used to look for contaminations:
               TREE:  uses a 4-ary tree. Index file optional,
               BLOOM: uses a bloom filter. Index file mandatory.
 -Q, --trimQ   NO:       does nothing to low quality reads (default),
               ALL:      removes all reads containing at least one low
                         quality nucleotide.
               ENDS:     trims the ends of the read if their quality is
                         below the threshold -q,
               FRAC:     discards a read if the fraction of bases whose
                         quality lies below
                         the threshold -q is over 5 percent or a user
                         defined percentage in -p.
               ENDSFRAC: trims the ends and then discards the read if
                         there are more low quality nucleotides than 
                         allowed by the option -p.
               GLOBAL:   removes n1 bases on the left and n2 on the
                         right, specified by -g.
               All reads are discarded if they are shorter than `minL`.
 -m, --minL    minimum length allowed for a read before it is discarded
               (default 25).
 -q, --minQ    minimum quality allowed (int), optional (default 27).
 -0, --zeroQ   value of ASCII character representing zero quality (int), optional (default 33)
 -p, --percent percentage of low quality bases to be admitted before
               discarding a read (default 5),
 -g, --global  required option if --trimQ GLOBAL is passed. Two int,
               n1:n2, have to be passed specifying the number of bases
               to be globally cut from the left and right, respectively.
 -N, --trimN   NO:     does nothing to reads containing N's,
               ALL:    removes all reads containing N's,
               ENDS:   trims ends of reads with N's,
               STRIPS: looks for the largest substring with no N's.
               All reads are discarded if they are shorter than `minL`.
```

NOTE: the parameters -l or --length are meant to identify the length
of the reads in the input data. Actually, `trimFilter` also copes with
data holding reads with different lengths. The length parameter must
hold the length of the longest read in the dataset.


## Output description

- `O_PREFIX_good.fq.gz`: contains reads that passed all filters (may be trimmed).
- `O_PREFIX_adap.fq.gz`: contains reads discarded due to the presence of adapters.
- `O_PREFIX_cont.fq.gz`: contains contamination reads.
- `O_PREFIX_lowQ.fq.gz`: contains reads discarded due to low quality issues.
- `O_PREFIX_NNNN.fq.gz`: contains reads discarded due to *N*'s issues.
- `O_PREFIX_summary.bin`: binary file where information about the filtering
   process is stored. Structure of the file.
    * filters, `4*sizeof(int)  Bytes`: array of int with entries
       `i = {ADAP(0), CONT(1), LOWQ(2), NNNN(3)}`. A given entry takes
       the value of the filter it was applied to and 0 otherwise.
       `filters[ADAPT] = {0,1}`, `filters[CONT] = {NO(0), TREE(1), BLOOM(2)}`,  
       `filters[LOWQ] = {NO(0), ALL(1), ENDS(2), FRAC(3), 
       ENDSFRAC(4), GLOBAL(5)}`, `filters[trimN] = {NO(0), ALL(1), 
       ENDS(2), STRIPS(2)}`.
    * trimmed, `4*sizeof(int) Bytes`: array of integers with entries
       i = {ADAP(0), CONT(1), LOWQ(2), NNNN(3)}, containing how many
       reads were trimmed due to the corresponding filter.
    * discarded, `4*sizeof(int) Bytes`: array of integers with entries
       i = {ADAP(0), CONT(1), LOWQ(2), NNNN(3)}, containing how many
      reads were discarded due to the corresponding filter.
    * good, `sizeof(int) Bytes`: number of accepted reads (may be trimmed).
    * nreads, `sizeof(int) Bytes`: total number of reads.

## Filters

#### Adapters

Technical sequences within the reads are detected if the option
`--adapters <ADAPTERS.fa>:<mismatches>:<score>` is given. The
adapter(s) sequence(s) are read from the fasta file, and the 
search is done using an 'seed and extend' approach. It starts by looking for
16-nucleotides long seeds, for which a user defined number of mismatches is
allowed (`mismatches`). If found, a score is computed. If the score is larger 
than the user defined threshold (`score`) and the number of matched 
nucleotides exceeds 12, then the read is trimmed if the remaining part is 
longer than `minL` (user defined) and discarded otherwise. If no 
16-nucleotide-long seeds are found, we proceed with 8-nucleotide-long seeds 
and apply the same criteria to trim/discard a read. A list of possible
situations follows, to illustrate how it works (`minL=25`, `mismatches=2`):

```
ADAPTER: GGCATACGAGCTCTTCCGATCT
REV_COM: AGATCGGAAGAGCTCGTATGCC

CASE1A:  CACAGTCGATCAGCGAGCAGGCATTCATGCTGAGATCGGAAGAGATCGTATG
                                         ||||||||||||X|||----
                                         AGATCGGAAGAGCTCGTATG
         - Seed: 16 Nucleotides
         - Return: trimmed, TRIMA:0:31
CASE1B:  CACATCATCGCTAGCTATCGATCGATCGATGCTATGCAAGATCGGAAGAGCT
                                               ||||||||------
                                               AGATCGGAAGAGCT
         - Seed: 8 Nucleotides
         - Return: trimmed, TRIMA:0:37
CASE1C:  CACATCATCGCTAGCTATCGATCGATCGATGCTATGCACGAAGATCGGAAGA
                                                  ||||||||---
                                                  AGATCGGAAGA
         - Seed: 8 Nucleotides
         - Return: nothing done, reason: Match length < 12
CASE2A:  CATACATCACGAGCTAGCTAGAGATCGGAAGAGCTCGTATGCCCAGCATCGA
                              ||||||||||||||||------
                              AGATCGGAAGAGCTCGTATGCC
         - Seed: 16 Nucleotides
         - Return: discarded, reason: remaining read too short.
CASE2B:  CCACAGTACAATACATCACGAGCTAGCTAGAGATCGGAAGAGCTCGTATGCC
                                       ||||||||||||||||||||||
                                       AGATCGGAAGAGCTCGTATGCC
         - Seed: 16 Nucleotides
         - Return: trimmed, TRIMA:0:28
CASE3A:  TATGCCGTCTTCTGCTTGCAGTGCATGCTGATGCATGCTGCATGCTAGCTGC
         ||||||||||||||||--
         TATGCCGTCTTCTGCTTG
         - Seed: 16 Nucleotides
         - Return: discarded, reason: remaining read too short
CASE3B:  CGTCTTCTGCTTGCCGATCGATGCTAGCTACGATCGTCGAGCTAGCTACGTG
         ||||||||-----
         CGTCTTCTGCTTG
         - Seed: 8 Nucleotides
         - Return: discarded, reason: remaining read too short
CASE3C:  TCTTCTGCTTGCCGATCGATGCTAGCTACGATCGTCGAGCTAGCTACGTGCG
         ||||||||---
         TCTTCTGCTTG
         - Seed: 8 Nucleotides
         - Return: nothing done, reason: Match length < 12
```

The score is calculated as follows: 
 *  - matching bases: `score += log_10(4)`
 *  - unmatching bases: `score -= Q/10`, where Q is the quality score.


#### Impurities/biological contaminations

 Biological contaminations are removed if a fasta or an index file are given as an input.
 Two methods have been implemented to check for contaminations:

- **TREE**: this method is designed to identify impurities from
  small sequences, such as rRNA, or E.coli (not larger than 10MB).
  When using this method, one has to pass one of these two options:
   - `--ifa <INPUT.fa>:<score>:<lmer_len>`: the file `INPUT.fa`
     is read and a tree of depth `lmer_len` is constructed on the flight.
   - `--idx <INDEX_FILE>:<score>`: in `INDEX_FILE`  the tree structure
      was stored with `./makeTree`.
  A read will be considered to be a contamination if the score is bigger
  than `score`. The score is computed as the proportion of Lmers of a given
  read found in the tree. This method is very fast, every search is
  `O( 2 * Lmer * (L - Lmer + 1))` (the reverse complement has to be
   checked also), but has the drawback that is very memory intensive.
   This is why we have limited the construction of a tree to sequences
   whose size does not exceed 10 MB. Constructing a tree is very fast,
   that is why most of the times it might not be worth to create the
   index file and then read it for every sample. Nevertheless, both
   options are provided.
- **BLOOM**: this method is designed for large sequences up to 4GB. An
  index is precomputed, using `makeBloom`, so that it is computed only
  once for all samples. The index file name and the threshold score have
  to be specified through the following option:
   - `--idx <INDEX_FILE>:<score>`
  The score is computed as for the previous options. 

#### Low quality

- `--trimQ NO` or flag absent: nothing is done to the reads with low quality.
- `--trimQ ALL`: all reads containing at least one low quality nucleotide are
  redirected to `*_lowq.fq.gz`
- `--trimQ ENDS`: look for low quality (below minQ) base callings at the
  beginning and at the end of the read. Trim them at both ends until the
  quality is above the threshold. Keep the read in `*_good.fq.gz`
  and annotate in the fourth line where the read has been trimmed (starting to
  count from 0) if the length of the remaining part is at least `minL`.
  Redirect the read to `*_lowq.fq.gz` otherwise.

    Examples (-q 27 [<] -m 25):
 ```
 ORIGINAL                                            FILTERED
 @ read 1081133                                      @ read 1081133
 GTACATTGATGAGCAACATGGGGCTTGAACTGGCGCTGAAACAGTTAGGA  GTACATTGATGAGCAACATGGGGCTTGAACTGGCGCTGAAACAGTTAGG
 +position: -144740                                  +position: -144740 TRIMQ:0:48
 IIIIIIIIIIIII9IIIIIIIIIIIIIIIIIIIIIIIIIIIIIIIIIII9  IIIIIIIIIIIII9IIIIIIIIIIIIIIIIIIIIIIIIIIIIIIIIIII
 @ read 3435                                         @ read  3435
 CAGTTCTTTGGTGTAGATGGAGCAGGACGGGATACCCATACCGTGACCCA  CTTTGGTGTAGATGGAGCAGGACGGGATACCCATACCGTG
 +position: -19377                                   +position: -19377  TRIMQ:5:44
 99999IIIIIIIIIIIIIIIIIIIIIIIIIIIIIIIIIIIIIIII99999  IIIIIIIIIIIIIIIIIIIIIIIIIIIIIIIIIIIIIIII
 @ read 108110                                       @ read 108110
 CAGATAAATGCGAATGAATGGGTAACACTGGAGCTTTTAATGGCTGTAAC  AGATAAATGCGAATGAATGGGTAACACTGGAGCTTTTAATGGCTGTAA
 +position: -4142524                                 +position: -4142524  TRIMQ:1:48
 9IIIIIIIIIIIIIIIIIII9IIII9III9IIIIIIIIIIIIIIIIIII9  IIIIIIIIIIIIIIIIIII9IIII9III9IIIIIIIIIIIIIIIIIII
 @ read 108111
 AGCGTGACGGTGTAACGCCCGCCTTTTGATGACTGGGTTTCAAAGAAACG
 +position: -3336785
 999999999999999IIIIIIII9IIIIIIIII9II99999999999999
 ```

- `--trimQ FRAC [--percent p]`: redirect  the read to `*_lowq.fq.gz` if
   there are more than `p%` nucleotides whose quality lies below the threshold.
   `-p 5` per default.

    Examples (-q 27 [<] -m 25 -p 5):
 ```
 ORIGINAL                                            FILTERED
 @ read 1081133                                      @ read 1081133
 GTACATTGATGAGCAACATGGGGCTTGAACTGGCGCTGAAACAGTTAGGA  GTACATTGATGAGCAACATGGGGCTTGAACTGGCGCTGAAACAGTTAGGA
 +position: -144740                                  +position: -144740
 IIIIIIIIIIIII9IIIIIIIIIIIIIIIIIIIIIIIIIIIIIIIIIII9  IIIIIIIIIIIII9IIIIIIIIIIIIIIIIIIIIIIIIIIIIIIIIIII9
 @ read 3435
 CAGTTCTTTGGTGTAGATGGAGCAGGACGGGATACCCATACCGTGACCCA
 +position: -19377
 99999IIIIIIIIIIIIIIIIIIIIIIIIIIIIIIIIIIIIIIII99999
 @ read 108110
 CAGATAAATGCGAATGAATGGGTAACACTGGAGCTTTTAATGGCTGTAAC
 +position: -4142524
 9IIIIIIIIIIIIIIIIIII9IIII9III9IIIIIIIIIIIIIIIIIII9
 @ read 108111
 AGCGTGACGGTGTAACGCCCGCCTTTTGATGACTGGGTTTCAAAGAAACG
 +position: -3336785
 999999999999999IIIIIIII9IIIIIIIII9II99999999999999
 ```
- `--trimQ ENDSFRAC --percent p`: first trim the ends as in the `ENDS` option.
   Accept the trimmed read if the number of low quality nucleotides does not
   exceed `p%` (default  `-p 5`).
   Redirect the read to `*_lowq.fq.gz` otherwise.

   Examples (-q 27 [<] -m 25  -p 5):
   Filtered reads will end up in `*_lowq.fq.gz`.
 ```
 ORIGINAL                                            FILTERED
 @ read 1081133                                      @ read 1081133
 GTACATTGATGAGCAACATGGGGCTTGAACTGGCGCTGAAACAGTTAGGA  GTACATTGATGAGCAACATGGGGCTTGAACTGGCGCTGAAACAGTTAGG
 +position: -144740                                  +position: -144740 TRIMQ:0:48
 IIIIIIIIIIIII9IIIIIIIIIIIIIIIIIIIIIIIIIIIIIIIIIII9  IIIIIIIIIIIII9IIIIIIIIIIIIIIIIIIIIIIIIIIIIIIIIIII
 @ read 3435                                         @ read  3435
 CAGTTCTTTGGTGTAGATGGAGCAGGACGGGATACCCATACCGTGACCCA  CTTTGGTGTAGATGGAGCAGGACGGGATACCCATACCGTG
 +position: -19377                                   +position: -19377  TRIMQ:5:44
 99999IIIIIIIIIIIIIIIIIIIIIIIIIIIIIIIIIIIIIIII99999  IIIIIIIIIIIIIIIIIIIIIIIIIIIIIIIIIIIIIIII
 @ read 108110
 CAGATAAATGCGAATGAATGGGTAACACTGGAGCTTTTAATGGCTGTAAC
 +position: -4142524
 9IIIIIIIIIIIIIIIIIII9IIII9III9IIIIIIIIIIIIIIIIIII9
 @ read 108111
 AGCGTGACGGTGTAACGCCCGCCTTTTGATGACTGGGTTTCAAAGAAACG
 +position: -3336785
 999999999999999IIIIIIII9IIIIIIIII9II99999999999999
 ```

- `--trimQ GLOBAL --global n1:n2`: cut all reads globally `n1` nucleotides from
   the left and `n2` from the right.

**Note:** qualities are evaluated assuming the reads to follow the
L - Illumina 1.8+ Phred+33, convention, see [Wikipedia](https://en.wikipedia.org/wiki/FASTQ_format#Encoding).
Adjust the values for a different convention.


#### N trimming

We allow for the following options:

- `--trimN NO` (or flag absent): Nothing is done to the reads containing N's.
- `--trimN ALL`: All reads containing at least one N are redirected to
  `*_NNNN.fq.gz`
- `--trimN ENDS`: N's are trimmed if found at the ends, left "as is"
  otherwise. If the trimmed read length is smaller than minL, the read is discarded.
  Example:
```
ORIGINAL                                           FILTERED
@ read 1037                                        @ read 1037
NNTCGACACAAGAAAATGCGCCAATTTTGAGCCAGACCCCAGTTACGCNN TCGACACAAGAAAATGCGCCAATTTTGAGCCAGACCCCAGTTACGC
+position: -2044610                                +position: -2044610  TRIMN:2:47
IIIIIIIIIIIIIIIIIIIIIIIIIIIIIIIIIIIIIIIIIIIIIIIIII IIIIIIIIIIIIIIIIIIIIIIIIIIIIIIIIIIIIIIIIIIIIII
@ read 1038
AAAAAACGGTCGTGGTNGGCTACTGTTATTAAAGCGTTGGCTACAAAAAG
+position: 1361068
IIIIIIIIIIIIIIIIIIIIIIIIIIIIIIIIIIIIIIIIIIIIIIIIII
@ read 1039
CCAACGGACNATGGAAGTGCTNCGGCTGGNGTTTTTATCCTCCGGCATTC
+position: -4282223
IIIIIIIIIIIIIIIIIIIIIIIIIIIIIIIIIIIIIIIIIIIIIIIIII
@ read 1043                                        @ read 1043
AATCTGAAACGAAANCTGACAGCGNNCCCCGCTTCTGACAAAATAGGCGN AATCTGAAACGAAANCTGACAGCGNNCCCCGCTTCTGACAAAATAGGCG
+position: -4685876                                +position: -4685876  TRIMN:0:48
IIIIIIIIIIIIIIIIIIIIIIIIIIIIIIIIIIIIIIIIIIIIIIIIII IIIIIIIIIIIIIIIIIIIIIIIIIIIIIIIIIIIIIIIIIIIIIIIII
```


- `--trimN STRIP`: Obtain the largest N free subsequence of the read. Accept it
   if is at least minL nucleotides long, redirect it to
   `*_NNNN.fq.gz` otherwise. Example:
```
ORIGINAL                                           FILTERED
@ read 1037                                        @ read 1037
NNTCGACACAAGAAAATGCGCCAATTTTGAGCCAGACCCCAGTTACGCNN TCGACACAAGAAAATGCGCCAATTTTGAGCCAGACCCCAGTTACGC
+position: -2044610                                +position: -2044610  TRIMN:2:47
IIIIIIIIIIIIIIIIIIIIIIIIIIIIIIIIIIIIIIIIIIIIIIIIII IIIIIIIIIIIIIIIIIIIIIIIIIIIIIIIIIIIIIIIIIIIIII
@ read 1038                                        @ read 1038
AAAAAACGGTCGTGGTNGGCTACTGTTATTAAAGCGTTGGCTACAAAAAG GGCTACTGTTATTAAAGCGTTGGCTACAAAAAG
+position: 1361068                                 +position: 1361068  TRIMN:17:49
IIIIIIIIIIIIIIIIIIIIIIIIIIIIIIIIIIIIIIIIIIIIIIIIII IIIIIIIIIIIIIIIIIIIIIIIIIIIIIIIII
@ read 1039
CCAACGGACNATGGAAGTGCTNCGGCTGGNGTTTTTATCCTCCGGCATTC
+position: -4282223
IIIIIIIIIIIIIIIIIIIIIIIIIIIIIIIIIIIIIIIIIIIIIIIIII
@ read 1043
AATCTGAAACGAAANCTGACAGCGNNCCCCGCTTCTGACAAAATAGGCGN
+position: -4685876
IIIIIIIIIIIIIIIIIIIIIIIIIIIIIIIIIIIIIIIIIIIIIIIIII
```


## Test/examples

 The examples in folder `examples/trimFilter_SReport/` work in the following
 way:

1. See folder `examples/fa_fq_files`. The file `EColi_rRNA.fq` was created with
 `create_fq.sh` and contains:                                               
   * 2e4 reads of length 50 from `EColi_genome.fa` with NO errors. 
   * 5e3 reads of length 50 from `rRNA_modified.fa` with NO errrors 
     (rRNA contaminations).                                                  
   * Artificially generated reads with low quality score (see `create_fq.sh`)
   * Artificially generated reads with Ns (see `create_fq.sh`).
   * Adapter files: `adapter_even_long.fa`, `adapter_odd_long.fa`, 
   `adapter_even_short.fa`, `adapter_odd_short.fa`. Fasta files containing 
    one adapter sequence each, longer/shorter than 16 nucleotides and with 
    an even/odd length. 
   * Example files to test the adapter contamination searches:
   `human_[even/odd]_wad_[even/odd]_[long/short].fq`. Short fastq files where
    adapters/contaminations have been inserted in all possible ways:
    even/odd positions, at the beginning/middle/end of the reads. Read 
    lengths are even or odd as the first suffix indicates. The adapter 
    contaminations included are suggested by the second even/odd suffix, 
    and the long/short suffix. 
2. `adapters/run_example.sh`: runs examples of reads containing adapter
   contaminations. A set of different possibilities is covered. 
   See README file inside the folder `adapters`.

3. `run_example_TREE.sh`: the code was tested with flags:                     
   ```
    $ ../../bin/trimFilter -l 50 --ifq PATH/TO/EColi_rRNA.fq.gz 
    --method TREE --ifa PATH/TO/rRNA_modified.fa:0.9:50 
    --trimQ ENDSFRAC --trimN STRIP -o tree
   ```                                  
   i.e., we check for contaminations from rRNA, trim reads with low qualities at
   the ends and less than 5% in the remaining part, and strip reads
   containing N's. The output should coincide with the files `example_TREE*`                 
4. `run_example_BLOOM.sh`:                                                    
  * a bloom filter is generated for `rRNA_modified.fa` with FPR = 0.0075
    and `kmersize=25`. The output should coincide with `rRNA_example.bf*`.
  * trimFilter is run like in 2. but passing a bloom filter to look for
    contaminations with `score=0.4`. 
5. With this set up, it is possible to run further customized tests.         
                                                                              
**NOTE:** `rRNA_modified.fa` is the `rRNA_CRUnit.fa` sequence, where we have     
        removed the lines containing N's for testing purposes.                
                                                                               

## Contributors

Paula Pérez Rubio

## License

GPL v3 (see LICENSE.txt)
