## Supplementary material for "FastqPuri: high-performance preprocessing of RNA-seq data": README_trimFilterPE.md

# trimFilterPE user manual

 This program reads two paired end `fastq` files as an input and filters them
according to the following criteria:
 - Discard/trims reads containing adapter remnants.
 - Discards reads matching biological contaminations (sequences collected in a `fasta`
   file or in an `idx` file created by `makeTree` or `makeBloom`)
 - Discards/trims low quality reads.
 - Discards/trims reads containing N base callings.

If one of the two reads is discarded, the corresponding paired read is
automatically discarded as well.

## Running the program

Usage `C` executable (in folder `bin`):

```
Usage: trimFilterPE --ifq <INPUT1.fq>:<INPUT2.fq> --length <READ_LENGTH>
                  --output [O_PREFIX] --gzip [y|n]
                  --adapter [<AD1.fa>:<AD2.fa>:<mismatches>:<score>]
                  --method [TREE|BLOOM]
                  (--idx [<INDEX_FILE>:<score>:<lmer_len>] |
                   --ifa [<INPUT.fa>:<score>:[lmer_len]])
                  --trimQ [NO|ALL|ENDS|FRAC|ENDSFRAC|GLOBAL]
                  --minL [MINL]  --minQ [MINQ] --zeroQ [ZEROQ]
                  (--percent [percent] | --global [n1:n2])
                  --trimN [NO|ALL|ENDS|STRIP]
Reads in paired end fq files (gz, bz2, z formats also accepted) and removes:
  * low quality reads,
  * reads containing N base callings,
  * reads representing contaminations, belonging to sequences in INPUT.fa
Outputs 4 [O_PREFIX]_fq.gz files per input fastq file containing: "good" reads,
discarded low quality reads, discarded reads containing N's, discarded contaminations.

If one read is removed, the corresponding paired read is removed as well.

Options:
 -v, --version prints package version.
 -h, --help    prints help dialog.
 -f, --ifq     2 fastq input files [*fq|*fq.gz|*fq.bz2] separated by colons, mandatory option.
 -l, --length  read length: length of the reads, mandatory option.
 -o, --output  output prefix (with path), optional (default ./out). 
 -z, --gzip    gzips output files: yes or no (default yes).
 -A, --adapter adapter input. Four fields separated by colons:
               <AD1.fa>: fasta file containing adapters from read 1,
               <AD2.fa>: fasta file containing adapters from read 2,
               <mismatches>: maximum mismatch count allowed,
               <score>: score threshold  for the aligner.
 -x, --idx     index input file. To be included with any method. 
               3 fields separated by colons:
               <INDEX_FILE>: output of makeTree or makeBloom,
               <score>: score threshold to accept a match [0,1],
               [lmer_len]: corresponds to the length of the lmers to be
                        looked for in the reads [1,READ_LENGTH].
 -a, --ifa     fasta input file. To be included only with method TREE
               (it excludes the option --idx). Otherwise, an
               index file has to be precomputed and given as parameter
               (see option --idx). 3 fields separated by colons:
               <INPUT.fa>: fasta input file [*fa|*fa.gz|*fa.bz2],
               <score>: score threshold to accept a match [0,1],
               <lmer_len>: depth of the tree: [1,READ_LENGTH]. 
                        corresponds to the length of the lmers to be
                        looked for in the reads.
 -C, --method  method used to look for contaminations:
               TREE:  uses a 4-ary tree. Index file optional,
               BLOOM: uses a bloom filter. Index file mandatory.
 -Q, --trimQ   NO:       does nothing to low quality reads (default),
               ALL:      removes all reads containing at least one low
                         quality nucleotide.
               ENDS:     trims the ends of the read if their quality is
                         below the threshold -q,
               FRAC:     discards a read if the fraction of bases whose
                         quality lies below the threshold -q is over 
			 5 percent or a user defined percentage in -p.
               ENDSFRAC: trims the ends and then discards the read if
                         there are more low quality nucleotides than 
                         allowed by the option -p.
               GLOBAL:   removes n1 bases on the left and n2 on the
                         right, specified in -g.
               All reads are discarded if they are shorter than minL.
 -m, --minL    minimum length allowed for a read before it is discarded
               (default 25).
 -q, --minQ    minimum quality allowed (int), optional (default 27).
 -0, --zeroQ   value of ASCII character representing zero quality (int), optional (default 33)
 -p, --percent percentage of low quality bases to be admitted before
               discarding a read (default 5),
 -g, --global  required option if --trimQ GLOBAL is passed. Two int,
               n1:n2, have to be passed specifying the number of bases
               to be globally cut from the left and right, respectively.
 -N, --trimN   NO:     does nothing to reads containing N's,
               ALL:    removes all reads containing N's,
               ENDS:   trims ends of reads with N's,
               STRIPS: looks for the largest substring with no N's.
               All reads are discarded if they are shorter than minL.
```

NOTE: the parameters -l or --length are meant to identify the length
of the reads in the input data.  Actually, `trimFilterPE` also copes with
data holding reads with different lengths. The length parameter must
hold the length of the longest read in the dataset.

## Output description

- `[O_PREFIX1 | O_PREFIX2]_good.fq.gz`: contains reads that passed all filters (may be trimmed).
- `[O_PREFIX1 | O_PREFIX2]_adap.fq.gz`: contains reads discarded due to the presence of adapters.
- `[O_PREFIX1 | O_PREFIX2]_cont.fq.gz`: contains reads from biological contamination.
- `[O_PREFIX1 | O_PREFIX2]_lowQ.fq.gz`: contains reads discarded due to low quality issues.
- `[O_PREFIX1 | O_PREFIX2]_NNNN.fq.gz`: contains reads discarded due to *N*'s issues.
- `[O_PREFIX1 | O_PREFIX2]_summary.bin`: binary file where information about the filtering
   process is stored. Structure of the file:
    * filters, `4*sizeof(int)  Bytes`: array of int with entries
       `i = {ADAP(0), CONT(1), LOWQ(2), NNNN(3)}`. A given entry takes
       the value of the filter it was applied to and 0 otherwise.
       `filters[ADAPT] = {0,1}`, `filters[CONT] = {NO(0), TREE(1),
        BLOOM(2)}`, `filters[LOWQ] = {NO(0), ALL(1), ENDS(2), FRAC(3),
        ENDSFRAC(4), GLOBAL(5)}`, `filters[trimN] = {NO(0), ALL(1),
        ENDS(2), STRIPS(2)}`.
    * trimmed, `4*sizeof(int) Bytes`: array of integers with entries
       i = {ADAP(0), CONT(1), LOWQ(2), NNNN(3)}, containing how many
       reads were trimmed due to the corresponding filter.
    * discarded, `4*sizeof(int) Bytes`: array of integers with entries
       i = {ADAP(0), CONT(1), LOWQ(2), NNNN(3)}, containing how many
      reads were discarded due to the corresponding filter.
    * good, `sizeof(int) Bytes`: number of accepted reads (may be trimmed).
    * nreads, `sizeof(int) Bytes`: total number of reads.

## Filters

#### Adapters

Technical sequences within the reads are detected if the option
`--adapters <ADAPTERS1.fa>:<ADAPTERS2.fa>:<mismatches>:<score>` is given.
The adapter(s) sequence(s) are read from the fasta files, and then prepended
to their respective reads. Then, a 'seed and extend' approach is used
to look for overlaps following the same rules followed in the single end case.
See `README_trimFilter.md` for details on how two matching subsequences are
detected and how the score is computed. The paired reads are correspondingly
trimmed, removed or left as is. The following figure describes possible
cases:

<p align="center">
<img src=./pics/adapters/palindrome_new.png alt="noimage" title="Adapters identification">
</p>


#### Impurities/biological contamination

Contaminations are removed if a fasta file or an index file are given as an
input. The methods provided to look for contaminations work in the very same
way as they work for single end data. If one of the reads is discarded,
then, the other read is discarded as well. See `README_trimFilter.md` for more
details on how the contaminations are handled.

#### Low quality

Again, the detection and trimming/removal of reads containing low quality
nucleotides is done following the same procedure as for single end data.
We list the options below, see `README_trimFilter.md` for more details.

- `--trimQ NO` or flag absent: nothing is done to the reads with low quality.
- `--trimQ ALL`: all reads containing at least one low quality nucleotide are
  redirected to  `*_lowq.fq.gz`.
- `--trimQ ENDS`: look for low quality (below minQ) base callings at the
  beginning and at the end of the read. Trim them at both ends until the
  quality is above the threshold. Keep the read in `*_good.fq.gz`
  and annotate in the fourth line where the read has been trimmed (starting to
  count from 0) if the length of the remaining part is larger than `minL`.
  Redirect the read to `*_lowq.fq.gz` otherwise.
- `--trim FRAC [--percent p]`: redirect  the read to `*_lowq.fq.gz` if
   there are more than `p%` nucleotides whose quality lies below the threshold.
   `-p 5` per default.
- `--trim ENDSFRAC --percent p`: first trim the ends as in the `ENDS` option.
   Accept the trimmed read if the number of low quality nucleotides does not
   exceed `p%` (default  `-p 5`).
   Redirect the read to `*_lowq.fq.gz` otherwise.
- `--trim GLOBAL --global n1:n2`: cut all reads globally `n1` nucleotides from
   the left and `n2` from the right.

**Note:** qualities are evaluated assuming the reads to follow the
L - Illumina 1.8+ Phred+33, convention, see [Wikipedia](https://en.wikipedia.org/wiki/FASTQ_format#Encoding).
Adjust the values for a different convention.


#### N trimming

We allow for the following options (see README_trimFilter.md for
examples and more details):

- `--trimN NO` (or flag absent): Nothing is done to the reads containing N's.
- `--trimN ALL`: All reads containing at least one N are redirected to
  `*_NNNN.fq.gz`
- `--trimN ENDS`: N's are trimmed if found at the ends, left "as is"
  otherwise. If the trimmed read length is smaller than minL, it is discarded.
- `--trimN STRIP`: Obtain the largest N free subsequence of the read. Accept it
   if it is longer than half the original read length, redirect it to
   `*_NNNN.fq.gz` otherwise.

## Test/examples

 The examples in folder `examples/trimFilterPE_SReport/` work in the following
 way:
 1. See folder `fa_fq_files`. The files `EColi_rRNA_DS.read1.fq.gz` and
    `EColi_rRNA_DS.read2.fq.gz` were created with `create_fq.sh` and contain:
    * 1000 reads of length 50 from `EColi_genome.fa` with NO errors.
    * 5000 reads of length 50 from `rRNA_modified.fa` with NO errrors
      (rRNA contaminations).
    * Artificially generated reads with low quality score (see `create_fq.sh`)
    * Artificially generated reads with Ns (see `create_fq.sh`).
 2. `run_example_TREE.sh`: the code was tested with flags:
    `../../bin/trimFilter -l 50 --ifq\
      --ifq ../fa_fq_files/EColi_rRNA_DS.read1.fq.gz:../fa_fq_files/EColi_rRNA_DS.read1.fq.gz
     --method TREE --ifa ../fa_fq_files/rRNA_modified.fa:0.2:30 \
     --trimQ ENDSFRAC --trimN ENDS -o treeDS --adapters \
     ../fa_fq_files/ad_read1.fa:../fa_fq_files/ad_read2.fa:2:40
    i.e., we check for contaminations from rRNA, trim reads with low qualities at
    the ends and less than 5% in the remaining part, and strip reads
    containing N's at the ends.
 3. `run_example_BLOOM.sh`:
    trimFilterPE is run like in 2. but passing a bloom filter to look for
       contaminations with `score=0.4` and the --trimN STRIP option.
 4. With this set up, it is possible to run further customized tests.
 5. See the folder `adapters` for examples on adapter contaminations (and its
    corresponding README file).

 NOTE: `rRNA_modified.fa` is the `rRNA_CRUnit.fa` sequence, where we have
        removed the lines containing N's for testing purposes.


## Contributors

Paula Pérez Rubio

## License

GPL v3 (see LICENSE.txt)
