## Supplementary figures and images for "FastqPuri: high-performance preprocessing of RNA-seq data"

### adapters_8c__incl.png

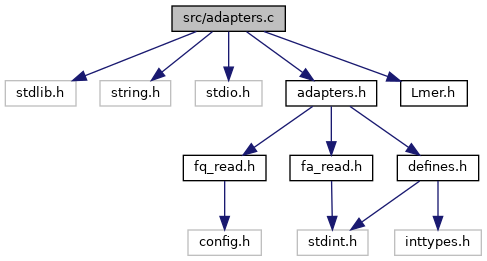

### adapters_8h__dep__incl.png

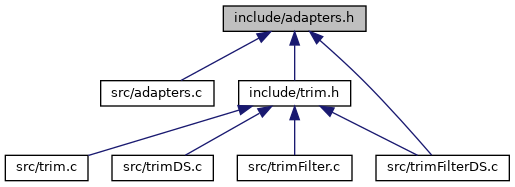

### adapters_8h__incl.png

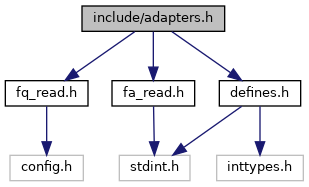

### bc_s.png

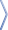

### bdwn.png

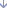

### bloom_8c__incl.png

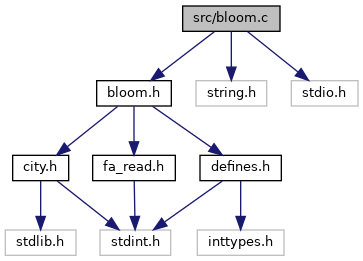

### bloom_8h__dep__incl.png

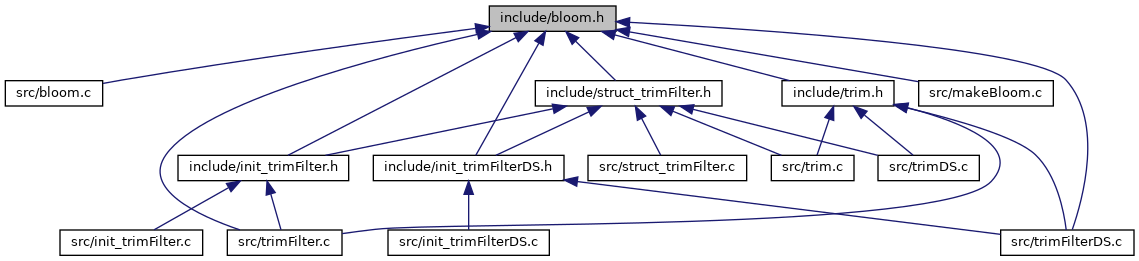

### bloom_8h__incl.png

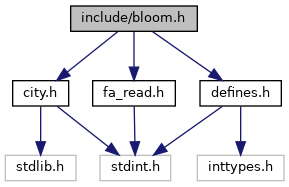

### city_8c__incl.png

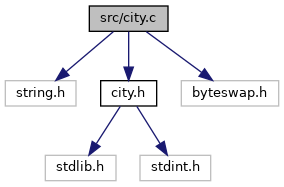

### city_8h__dep__incl.png

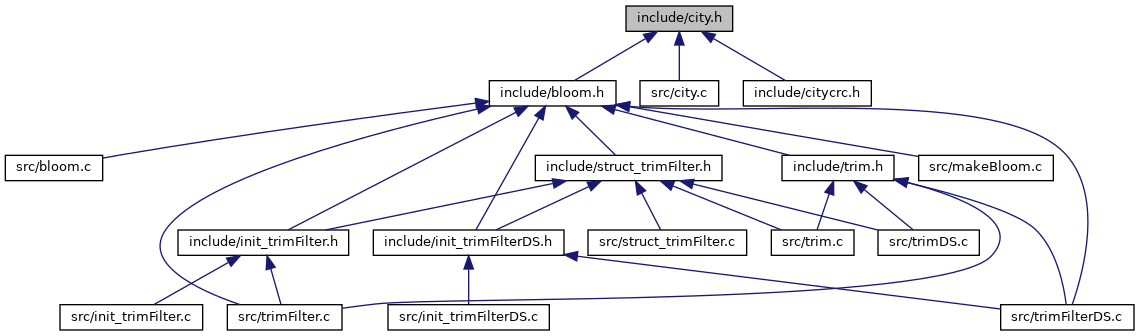

### city_8h__incl.png

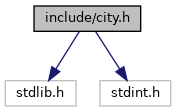

### citycrc_8h__incl.png

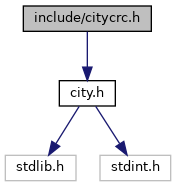

### closed.png

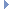

### defines_8h__dep__incl.png

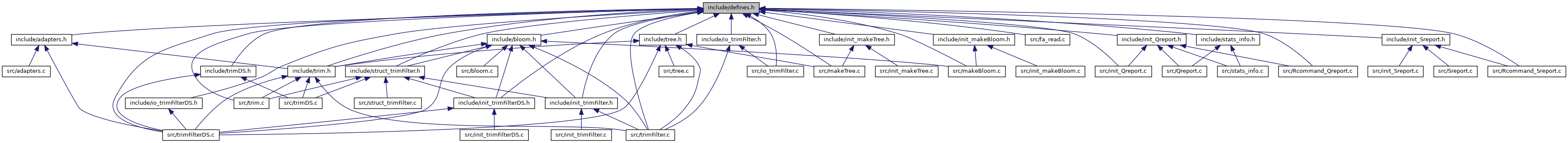

### defines_8h__incl.png

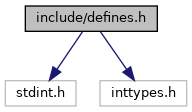

### dir_68267d1309a1af8e8297ef4c3efbcdba_dep.png

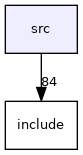

### doc.png

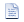

### doxygen.png

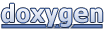

### example_ROC_0p01_bloom.pdf

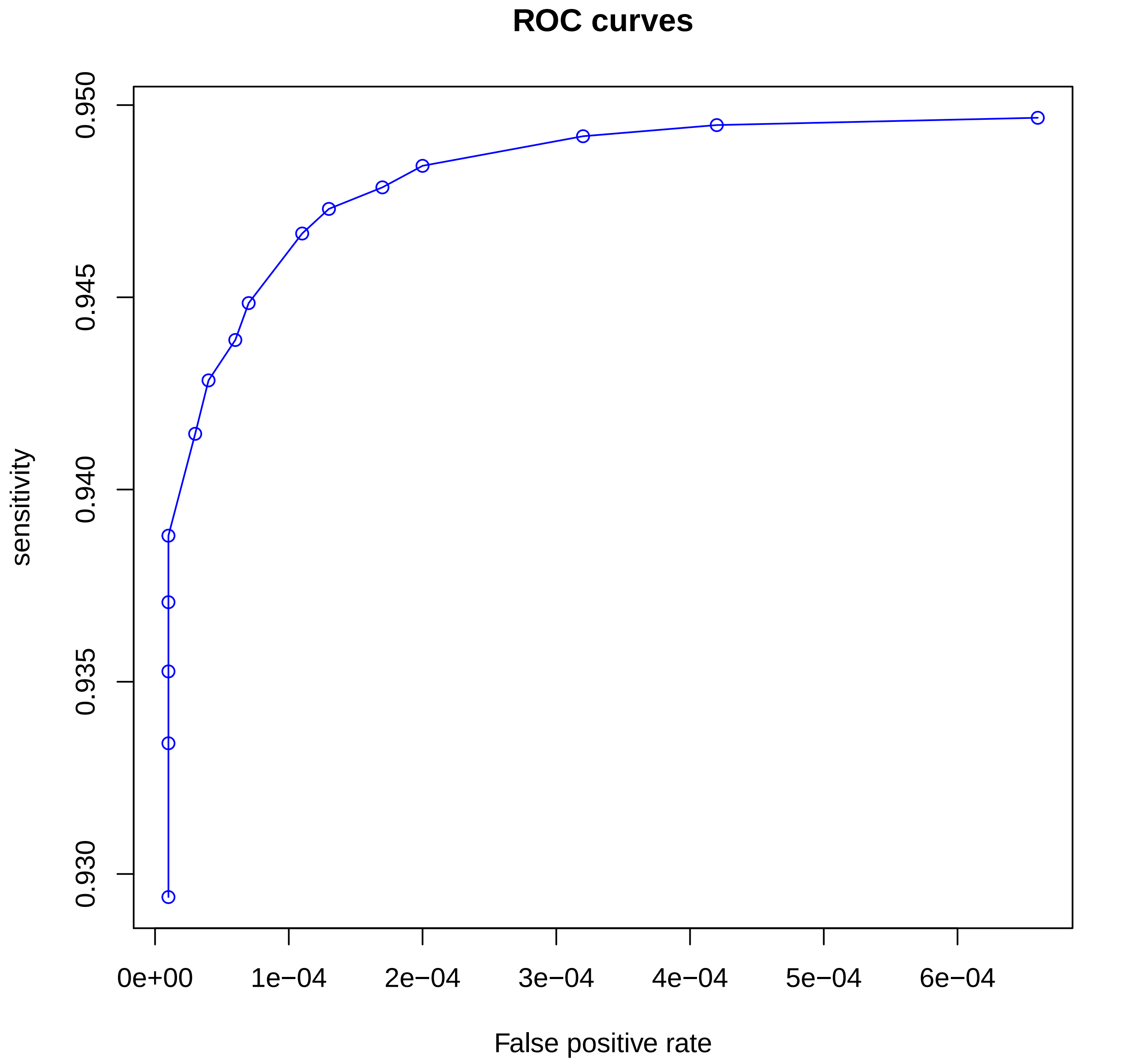

### example_ROC_0p02_bloom.pdf

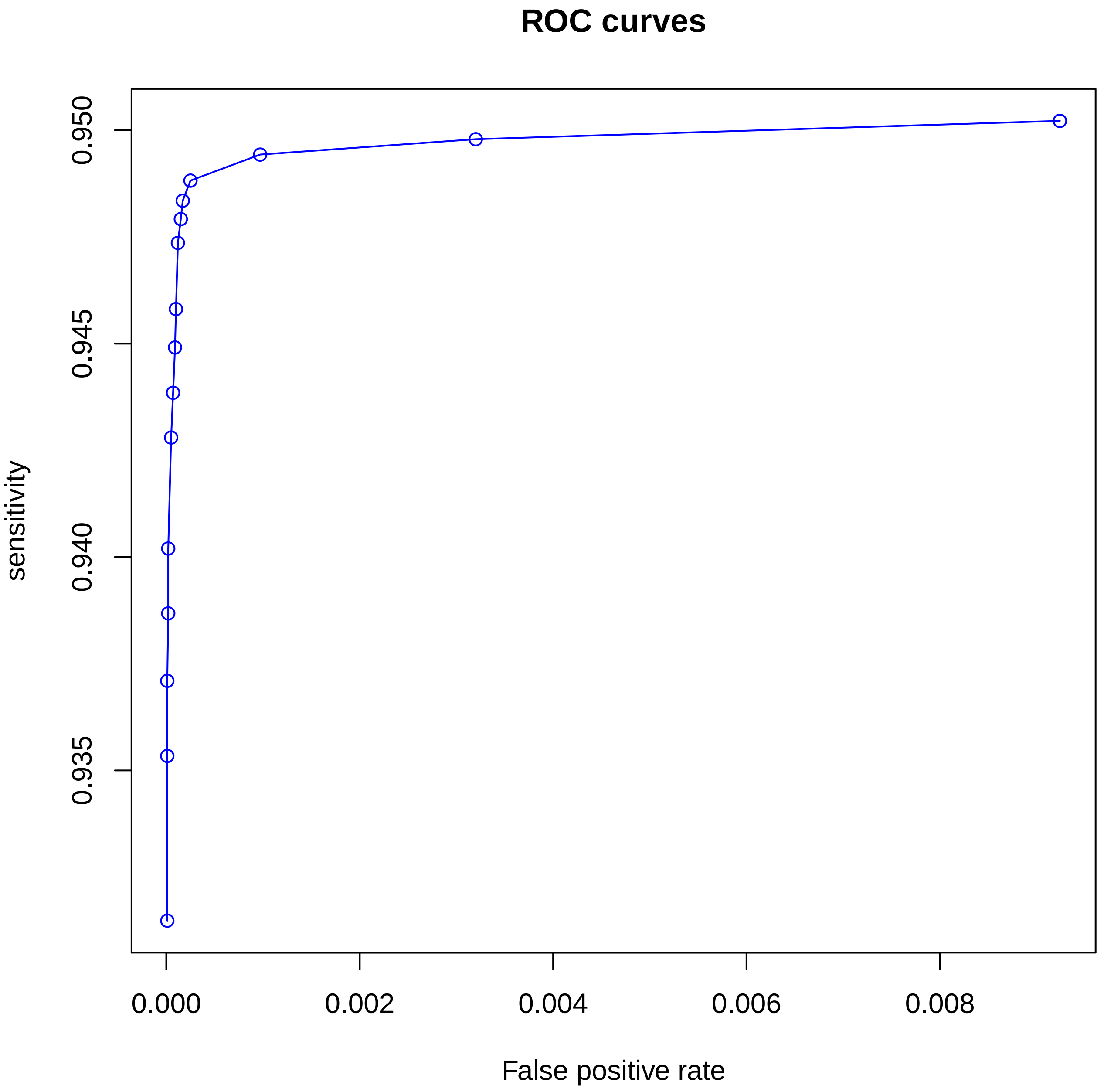

### example_ROC_0p005_bloom.pdf

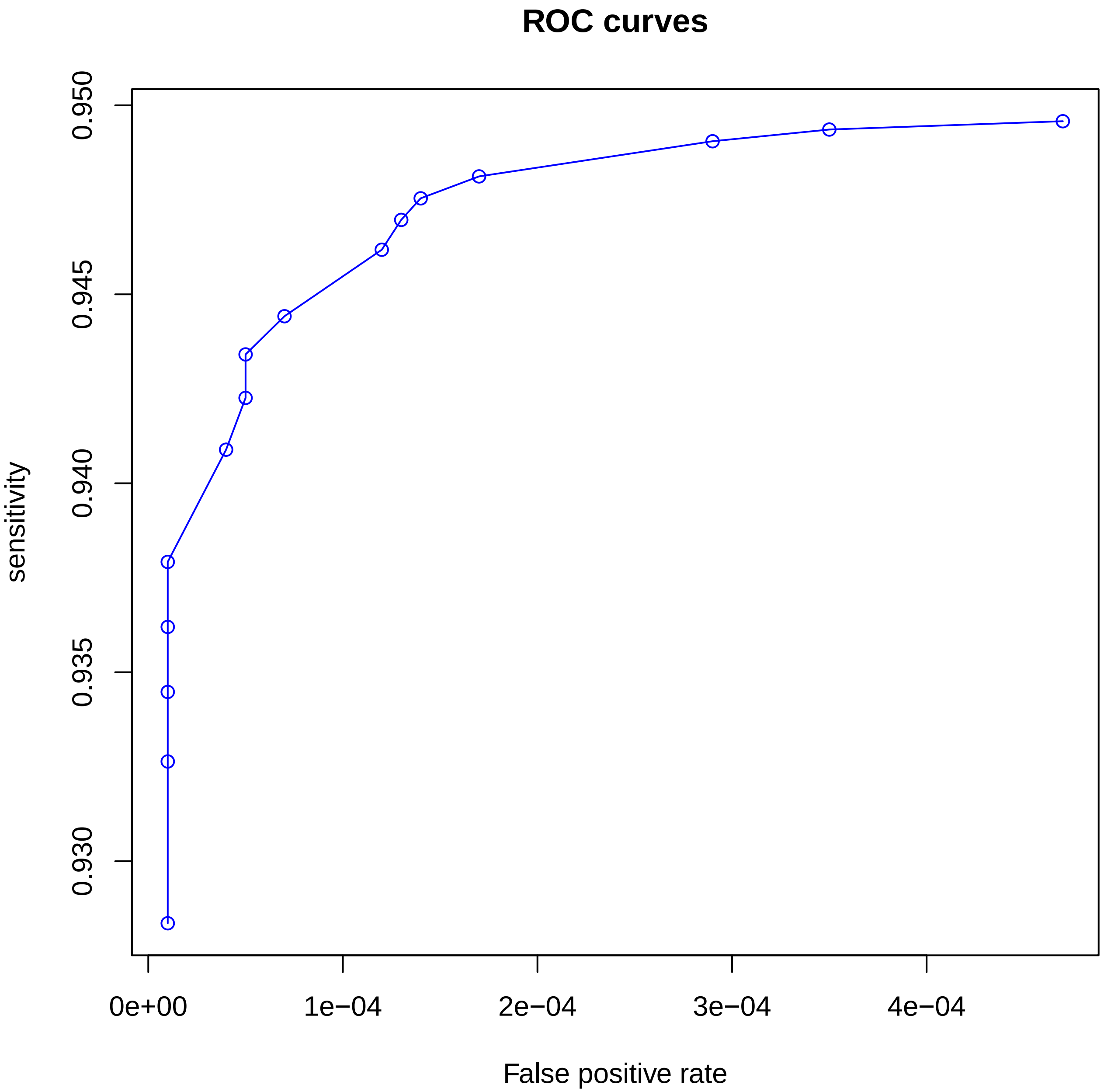

### example_ROC_0p0075_bloom.pdf

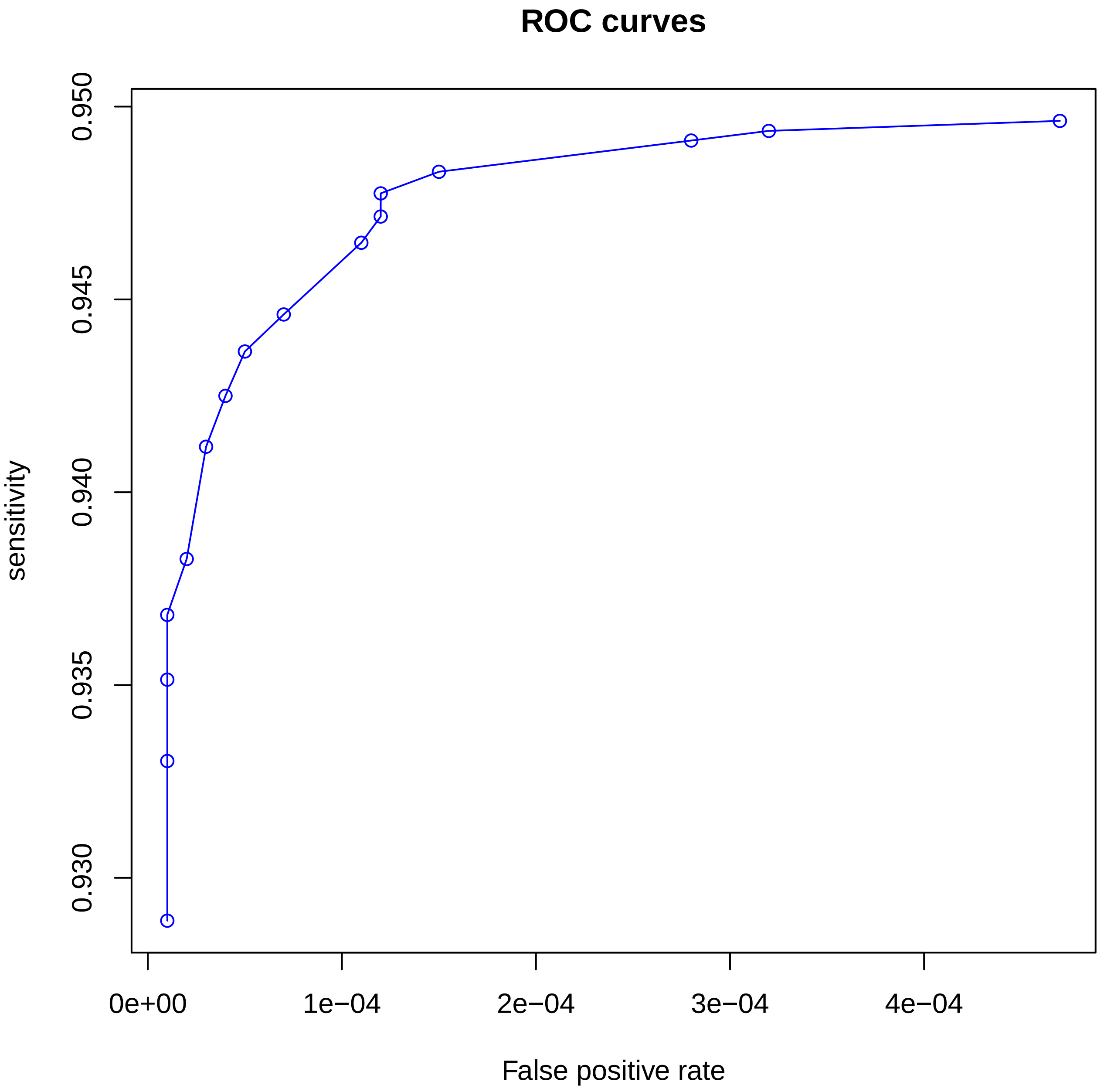

### fa__read_8c__incl.png

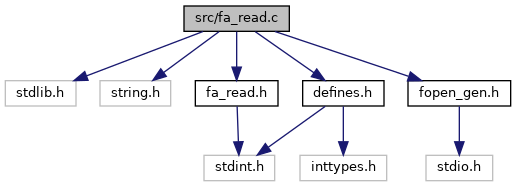

### fa__read_8h__dep__incl.png

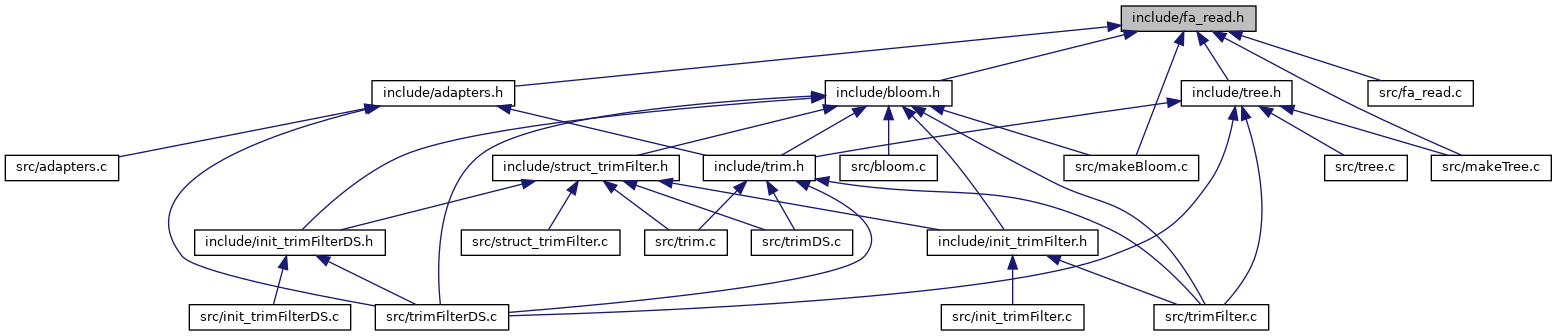

### fa__read_8h__incl.png

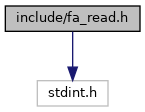

### folderclosed.png

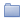

### folderopen.png

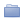

### fopen__gen_8c__incl.png

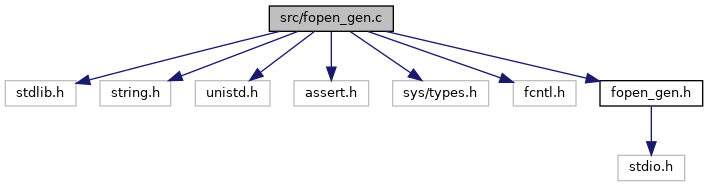

### fopen__gen_8h__dep__incl.png

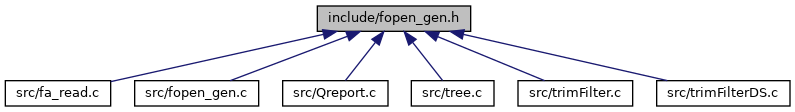

### fopen__gen_8h__incl.png

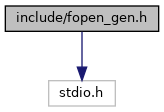

### Lmer_8c__incl.png

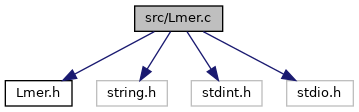

### Lmer_8h__dep__incl.png

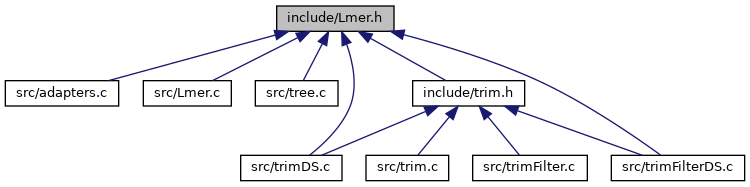

### Qreport_8c__incl.png

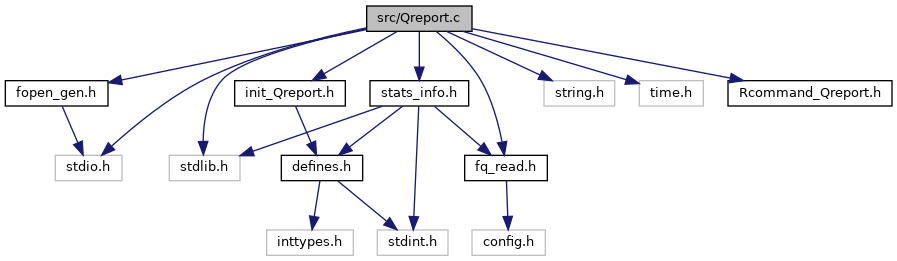

### Rcommand__Qreport_8c__incl.png

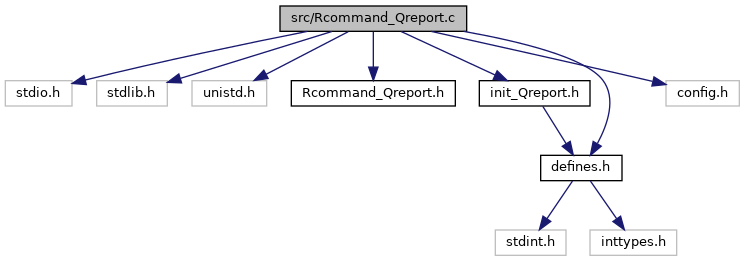

### Rcommand__Qreport_8h__dep__incl.png

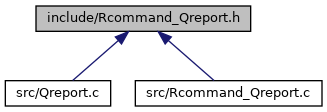

### Rcommand__Sreport_8c__incl.png

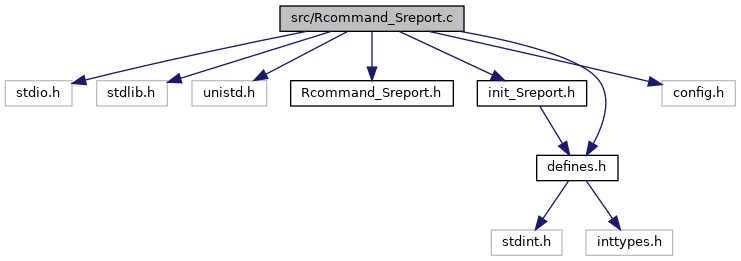

### Rcommand__Sreport_8h__dep__incl.png

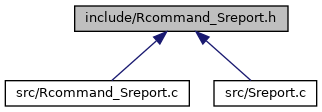

### Sreport_8c__incl.png

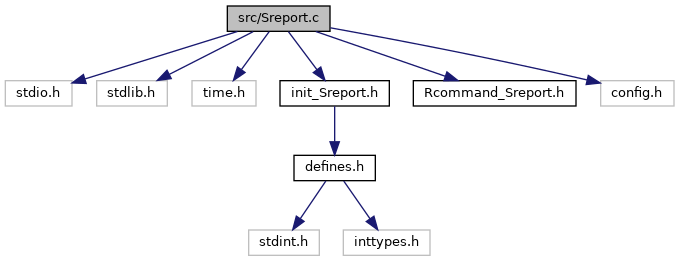
